# Bladder cancer organoids as a functional system to model different disease stages and therapy response

**DOI:** 10.1101/2022.03.31.486514

**Authors:** Martina Minoli, Thomas Cantore, Mirjam Kiener, Tarcisio Fedrizzi, Federico La Manna, Sofia Karkampouna, Vera Genitisch, Antonio Rodriguez, Irena Klima, Paola Gasperini, Bernhard Kiss, Roland Seiler-Blarer, Francesca Demichelis, George N. Thalmann, Marianna Kruithof-de Julio

## Abstract

Bladder Cancer (BLCa) inter-patient heterogeneity is considered the primary cause of tumor reoccurrence and treatment failure, suggesting that BLCa patients could benefit from a more personalized treatment approach. Patient-derived organoids (PDOs) have been successfully used as a functional model for predicting drug response in different cancer types. In our study, we established BLCa PDO cultures from different BLCa stages. BLCa PDOs preserve the histological and molecular heterogeneity of the parental tumors, including their multiclonal genetic landscapes. BLCa PDOs consistently share key genetic alterations detected in parental tumors, mirroring tumor evolution in longitudinal sampling. Our drug screening pipeline was implemented using BLCa PDOs, testing both standard-of-care and additional FDA-approved compounds for other solid tumors. Integrative analysis of drug response profiles with matched PDO genomic analysis was used to determine enrichment thresholds for candidate markers of therapy resistance and sensitivity. By assessing the clinical history of longitudinally sampled cases, the clonal evolution of the disease could be determined and matched with drug response profiles. In conclusion, we have developed a clinically relevant pipeline for drug response profile assessment and discovery of candidate markers of therapy resistance.

## Introduction

Bladder cancer (BLCa) is the tenth most common cancer worldwide, with 573,278 new cases and 212,536 deaths reported in 2020 alone (1). BLCa diagnosis falls into two main classes, non-muscle invasive (NMIBC) and muscle-invasive (MIBC) BLCa, determined by the depth of invasion towards the underlying bladder wall and with very distinct clinical profiles, etiologies, and natural histories. 70-75% of patients are diagnosed with NMIBC which is characterized by a high proliferation rate and frequent local relapses after tumor resection (2). Despite a 5-year overall survival rate of about 90% (3,4), NMIBC patients undergo multiple periodic resections and require life-long monitoring. Within the NMIBC group, approximately 15% will progress to MIBC following the acquisition of deleterious mutations in key driver genes (5). *De novo* MIBC is diagnosed in 25-30% of patients and has a 5-year overall survival rate of 50%. Its genetic instability frequently leads to rapid progression with metastatic dissemination in 5% of patients resulting in a 5-year overall survival rate of <10% (3,4,6,7). Despite the high morbidity and socio-economic costs, BLCa remains relatively understudied compared to other cancers with similar impact (4,8).

As a standard-of-care (SOC), NMIBC patients undergo transurethral resection of the bladder (TUR-B) and receive intravesical instillations of chemotherapy alone (low-risk NMIBC) or in combination with Bacillus Calmette-Guérin (BCG) vaccine (intermediate- and high-risk NMIBC) to promote tumor clearance and reduce the risk of recurrence (9). In contrast, MIBC patients have a high risk of local and systemic progression and, therefore, are treated with systemic neoadjuvant cisplatin-based chemotherapy (NAC) followed by radical cystectomy (10).

The distinct clinical and histological profiles between NMIBC and MIBC can be attributed to their genetic differences (11,12). NMIBC tumors predominantly harbor mutations promoting tumor growth, such as *FGFR3* and *HRAS* gain-of-function mutations (13–15). By contrast, MIBC tumors are characterized by alterations in tumor suppressor genes (*TP53* and *Rb1*) and genetic alterations in *EGFR/ERBB2* genes (13,16,17). Several groups have proposed different molecular classifications of both NMIBC and MIBC based on large-scale transcriptomic and genomic profiling (13,14,18–22). The identification of these molecular classifiers was a significant step forward in the understanding of BLCa heterogeneity. Likely due to the high molecular heterogeneity of BLCa, the overall response to SOC is lower than 30%, resulting in the high recurrence rate of NMIBC and adverse prognosis of MIBC (23,24). Better patient stratification and personalized treatment for both NMIBC and MIBC patients are urgently needed.

In recent years, precision medicine approaches have been explored for their feasibility and clinical effectiveness across numerous cancer types. In this context, patient-derived organoids (PDOs), human-derived 3D models suitable for high-throughput solutions, demonstrated increasing translational potential, offering an interesting tool for precision medicine approaches (25–28). To date, organoids have been derived from a variety of normal and malignant tissues including gastrointestinal (stomach, colorectal, pancreatic, and liver), breast, ovarian, prostate and bladder (29–40). PDOs have been shown to recapitulate key aspects of tissue composition, including architecture and function, and retain organ identity and remain genetically stable in culture (33,41). Most importantly, PDOs preserve patient-specific tissue composition (including tumor heterogeneity) and can predict drug response *in vitro* (31,33,35,36,38–40,42).

Here we successfully establish and culture organoids from NMIBC and MIBC that largely recapitulate key aspects of the original tumor tissue and BLCa mutational mechanisms allowing drug screening and mechanistic studies on drug sensitivity/resistance profiles as well as monitoring of tumor evolution. We demonstrate that BLCa PDOs maintain the histological and molecular heterogeneity of the original tumors. We use PDOs to stratify patients according to their response to SOC chemotherapies and targeted therapies. Moreover, we demonstrate that when organoid drug response profiles are integrated with patient genomic background they can identify biomarkers or signatures that can help to design a treatment regimen that is unique and appropriate to the patient’s genetic profile. These results strengthen the evidence that BLCa PDO can be applied as a platform for precision medicine.

## Results

### Establishment and culture of BLCa PDOs from diverse clinical samples

PDOs were generated from specimens obtained from patients that underwent either TUR-B, radical cystectomy or nephroureterectomy (**Fig. 1A, Tables 1–3**) at the Inselspital, University Hospital in Bern and representing the spectrum of BLCa, ranging from low-grade, non-invasive BLCa to high-grade invasive tumors, including both NMIBC and MIBC. PDOs were derived and grown in suspension (32) from fresh and cryopreserved tissue (viability 88%±6% and 86%±11%, respectively) and cryopreserved single cells (viability 75%±15%) (**Table S1**).

**Figure 1.**
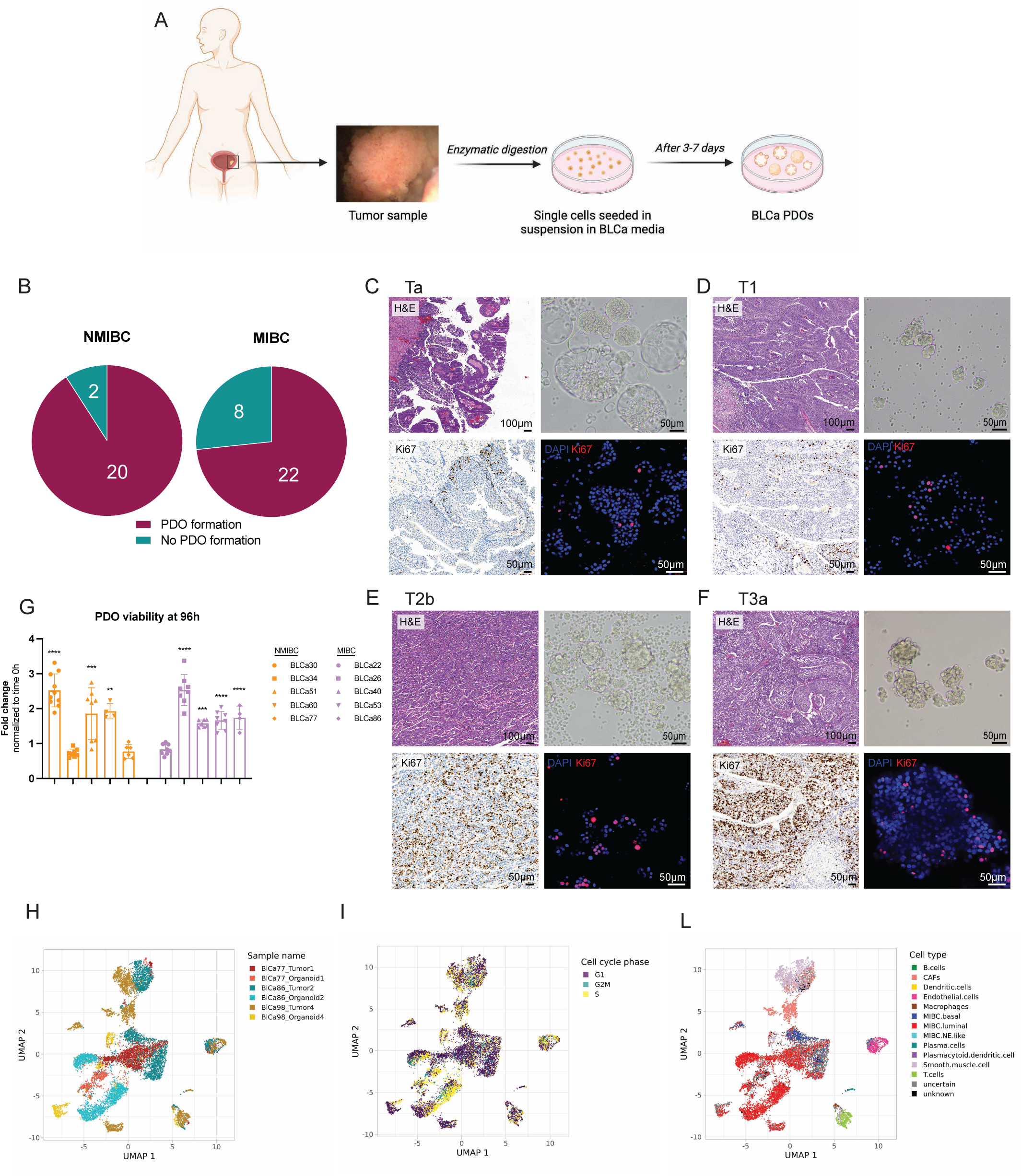
Isolation and culture of patient-derived organoids (PDOs) from non-muscle invasive bladder cancer (NMIBC) and muscle invasive bladder cancer (MIBC) (A) Scheme of the experimental protocol for bladder cancer (BLCa) organoids derivation and culture. (B) Number of PDO formation and no PDO formation over the total samples cultured for NMIBC (n=22) and MIBC (n=30). Fisher’s test; p-value = 0.1608. (C-F) Morphology of primary tumors (PTs) and matched PDOs (passage (p) 1) for four representative cases (BLCa34 transurethral resection of bladder tumor Ta (C); BLCa112 nephroureterectomy T1 (D); BLCa40 cystectomy T2b (E); and BLCa98 cystectomy T3a (F)). H&E and IHC for Ki67 for PT, and brightfield images and whole-mount immunofluorescence staining for Ki67 for PDO. (G) Viability assay of PDOs (p2) derived from 5 NMIBC and 5 MIBC samples after 96h in culture. PDO viability at 96h was normalized to the time 0. Each data point corresponds to one technical replicate (mean± SD). Ordinary two-way ANOVA with Sidak’s multiple comparison between time 0 and 96h * p-value ≤ 0.05, ** p-value < 0.01, *** p-value < 0.001, **** p-value < 0.0001. (H) UMAP plot of cells derived from PTs and PDOs for three sample (BLCa77, BLCa86 and BLCa98), colored by patient and sample type. (I) UMAP plot of cells derived from PTs and PDOs showing clustering by cell cycle phases. (L) UMAP plots showing cell types present in PT and PDOs at p1. Abbreviations: CAFs, cancer-associated fibroblasts; NE-like, neuroendocrine-like.

**Table 1.** List of 52 bladder cancer samples tested to generate organoid derivation. If samples are derived from the same patient at different time points, previous samples are reported
in brackets in the Sample ID column. Abbreviations: CIS, carcinoma in situ; F, female; M, male; N/C, no comment; TUR-B, transurethral
resection of the bladder.

| Sample ID | Specimen | Tumor stage | Age (Years) | Sex | Organoids formation | Comments |
| --- | --- | --- | --- | --- | --- | --- |
| BLCa30 | TUR-B | Ta | 66 | M | yes | N/C |
| BLCa34 | TUR-B | Ta | 77 | M | yes | N/C |
| BLCa35 | TUR-B | Ta | 81 | M | yes | N/C |
| BLCa43 | TUR-B | Ta | 67 | M | yes | N/C |
| BLCa51 | TUR-B | Ta | 67 | M | yes | N/C |
| BLCa57 (BLCa51) | TUR-B | Ta | 68 | M | yes | N/C |
| BLCa60 | TUR-B | Ta | 72 | M | yes | N/C |
| BLCa69 | TUR-B | Ta | 65 | M | yes | N/C |
| BLCa77 | TUR-B | Ta | 69 | M | yes | N/C |
| BLCa81 (BLCa69) | TUR-B | Ta | 66 | M | yes | N/C |
| BLCa113 | TUR-B | Ta | 78 | M | yes | N/C |
| BLCa127 (BLCa113) | Cystectomy | Ta | 78 | M | yes | N/C |
| BLCa116 (BLCa69, BLCa81) | TUR-B | Ta/CIS | 67 | M | yes | N/C |
| BLCa46 | TUR-B | T1 | 77 | F | yes | N/C |
| BLCa50 | TUR-B | T1 | 67 | M | yes | N/C |
| BLCa61 | TUR-B | T1 | 74 | M | yes | N/C |
| BLCa82 | TUR-B | T1 | 71 | M | yes | N/C |
| BLCa85 | TUR-B | T1 | 74 | M | yes | N/C |
| BLCa100 | TUR-B | T1 | 63 | M | yes | N/C |
| BLCa112 (BLCa51, BLCa57) | Nephroureterectomy | T1 | 69 | M | yes | N/C |
| BLCa22 | Cystectomy | T2a | 59 | M | yes | N/C |
| BLCa26 | TUR-B | T2a | 71 | M | yes | N/C |
| BLCa27 | Cystectomy | T2a | 74 | M | yes | N/C |
| BLCa66 | Cystectomy | T2a | 68 | M | yes | N/C |
| BLCa115 | Cystectomy | T2a | 84 | M | yes | N/C |
| BLCa40 (BLCa35) | Cystectomy | T2b | 81 | M | yes | N/C |
| BLCa86 (BLCa85) | Cystectomy | T2b | 74 | M | yes | N/C |
| BLCa92 | Cystectomy | T2b | 69 | M | yes | N/C |
| BLCa31 | Cystectomy | T3a | 62 | M | yes | N/C |
| BLCa38 | Cystectomy | T3a | 77 | M | yes | N/C |
| BLCa41 | Cold biopsy | T3a | 66 | M | yes | N/C |
| BLCa47 | Cystectomy | T3a | 66 | F | yes | N/C |
| BLCa48 (BLCa29) | Cystectomy | T3a | 78 | M | yes | N/C |
| BLCa53 | TUR-B | T3a | 80 | M | yes | N/C |
| LN_BLCa1 | Nephroureterectomy | T3a | 59 | F | yes | N/C |
| BLCa98 | Cystectomy | T3a | 83 | M | yes | N/C |
| BLCa33 | Cystectomy | T3b | 71 | M | yes | N/C |
| BLCa125 | Cystectomy | T3b | 55 | M | yes | N/C |
| BLCa93 | TUR-B | T3b/4a | 72 | M | yes | N/C |
| BLCa111 | Cystectomy | T3b/4a | 91 | M | yes | N/C |
| BLCa63 | TUR-B | T4a | 71 | M | yes | N/C |
| BLCa114 | Cystectomy | T4b | 58 | M | yes | N/C |
| BLCa18 | TUR-B | T1 | 42 | F | no | not enough viable cells |
| BLCa19 (BLCa18) | Cystectomy | T1 | 42 | F | no | not enough viable cells |
| BLCa71 | Cystectomy | T2 | 64 | M | no | contamination |
| BLCa87 | TUR-B | T2 | 62 | M | no | not enough viable cells |
| BLCa23 | TUR-B | T3 | 65 | M | no | not enough viable cells |
| BLCa29 | TUR-B | T3 | 77 | M | no | not enough viable cells |
| BLCa32<br>(BLCa23) | Needle biopsy | T3 | 65 | M | no | not enough viable cells |
| BLCa68<br>(BLCa23,<br>BLCa32) | TUR-B | T3 | 66 | M | no | debris and not enough<br>viable cells |
| BLCa102 | Cystectomy | T3 | 62 | M | no | not enough viable cells |
| BLCa58 | Lymph node dissection | T4a | 68 | M | no | not enough viable cells |

**Table 2.** Summary of patients’ clinical history. If samples are derived from the same patient at different time points, previous samples are reported
in brackets in the Sample ID column. Abbreviations: BCG, Bacillus Calmette-Guérin treatment; Cis/Gem, cisplatin and gemcitabine combination, F, female; M, male; N/A, not available; PY, per year; TUR-B, transurethral resection of the bladder; uns., unsampled; 1x, 1 cycle.

| Sample ID | Gender | Age | Smoker | Comorbidities | Pre-treatment | Post-operative treatment | Treatment response | Progression | Time to progression | Last follow up | Cancer-specific death |
| --- | --- | --- | --- | --- | --- | --- | --- | --- | --- | --- | --- |
| BLCa22 | M | 59 | Yes (30 PY) | Incidental adenocarcinoma of the prostate (T2c) | No | No | N/A | No | N/A | 27.04.21 | No |
| BLCa33 | M | 71 | Until 2013 | Adenocarcinoma of the prostate (T1b) | Neoadjuvant Cis/Gem | No | No | No | N/A | 09.06.21 | No |
| BLCa34 | M | 77 | Until 2009 (70-80 PY) | N/A | No | Intravesical installation Epirubicin | Yes | No | N/A | 02.03.21 | No |
| BLCa35 | M | 77 | N/A | Incidental adenocarcinoma of the prostate (T1a), hepatocellular carcinoma and chronic hepatitis C | No | No | N/A | No | N/A | 22.12.20 | No |
| BLCa40 (BLCa35) | M | 81 | N/A | BLCa35 Neuroendocrine-like tumor in the liver | No | No | N/A | No | N/A | 22.12.20 | No |
| BLCa46 | F | 77 | No | N/A | No | Intravesical installation BCG 6x | N/A | No | N/A | 18.02.20 | No |
| BLCa47 | F | 69 | N/A | N/A | No | No | N/A | No | N/A | 29.09.21 | No |
| BLCa50 | M | 67 | N/A | Prostate cancer (T2a) | No | No | N/A | No | N/A | 19.03.21 | No |
| BLCa57 | M | 68 | N/A | Acinar adenocarcinoma of the prostate (T2c) | Intravesical installation Epirubicin and BCG 6x previous treatments | No | N/A | Yes (BLCa112) | 486 days | 14.01.21 | No |
| BLCa60 | M | 72 | No | Adenocarcinoma of the prostate (T2a) | No | No | N/A | No | N/A | 08.09.21 | No |
| BLCa61 | M | 73 | N/A | N/A | Intravesical installation BCG 1x | No | N/A | No | N/A | 15.09.21 | No |
| BLCa69 | M | 65 | Until 2019 (40 PY) | Acute myeloid leukemia | No | Intravesical installation Epirubicin 1x | No | Yes | 45 days first relapse (uns.), 196 days between baseline and second relapse (BLCa81, relapse 1) | 28.10.21 | No |
| BLCa81 (BLCa69) | M | 66 | BLCa69 | BLCa69 | Intravesical installation Epirubicin 1x (BLCa69) | Intravesical installation BCG 6x | No | Yes | 389 days (uns.), 446 days (sampled, relapse 2), 679 days (uns.) and 699 days (sampled, relapse 3) after the baseline sample<br>(BLCa69) | 28.10.21 | No |
| BLCa82 | M | 71 | Yes<br>(70PY) | N/A | No | Palliative<br>chemotherapy<br>(Carboplatin/Gemcita<br>bine) | No | No | N/A | 23.07.20 | No |
| BLCa86 | M | 74 | N/A | N/A | No | No | N/A | No | N/A | 21.12.20 | No |
| BLCa98 | M | 83 | No | Benigne prostate<br>hyperplasia | No | No | N/A | No | N/A | 18.01.21 | No |
| BCLa112 |  | 69 | N/A | BLCa57 | Intravesical<br>installation<br>Epirubicin and BCG<br>6x previous<br>treatments | No | N/A | No | N/A | 14.01.21 | No |

**Table 3.** Pathological data of the tumor. Tumor stage and pathological classification were evaluated for the parental tumor. If samples are derived from the same patient at different time points, previous samples are reported in brackets in the Sample ID column. Abbreviations: TUR-B, transurethral resection of the bladder; CIS, carcinoma in situ; N/A, not
available; PDO, patient-derived organoid.

| Sample code | Tissue source | Tumor stage | Precursor lesion |  | Histological variant of invasive carcinoma | Immune infiltration | Concomitant CIS | Main PDO morphology |
| --- | --- | --- | --- | --- | --- | --- | --- | --- |
|  |  |  | Type | Grade |  |  |  |  |
| BLCa34 | TUR-B | Ta | papillary | Low | N/A | none | No | Hollow |
| BLCa35 | TUR-B | Ta | papillary | High | N/A | Peritumoral weak | No | Solid |
| BLCa57 | Cystectomy | Ta | papillary | Low | N/A | Peritumoral weak | No | Hollow |
| BLCa60 | Cystectomy | Ta | papillary | Low | N/A | Peritumoral weak | No | Mixed |
| BLCa69 | TUR-B | Ta | papillary | High | N/A | none | No | Mixed |
| BLCa81 (BLCa69) | TUR-B | Ta/Tis | papillary | High | N/A | none | Yes | Hollow |
| BLCa61 | Cystectomy | T1 | papillary | High | N/A | Peritumoral weak | No | Hollow |
| BLCa82 | TUR-B | T1 | CIS | High | Squamous in 70% of tumor | none | Yes | Solid |
| BLCa112 (BLCa57) | Nephroureterectomy | T1 | papillary | High | N/A | Intermediate | No | Mixed |
| BLCa46 | TUR-B | T≥1 | papillary | High | N/A | Peritumoral weak | No | Solid |
| BLCa50 | TUR-B | T≥1 | papillary | High | Glandular (also in non invasive papillary component) in 30% of tumor | Intra- and peritumoral weak | Yes | Solid |
| BLCa22 | Cystectomy | T2a | papillary | High | N/A | Peritumoral weak | No | Solid |
| BLCa40 (BLCa35) | Cystectomy | T2b | papillary | High | Glandular (mucinous/singet ring cells) in 20% of tumor | Intra- and peritumoral weak | Yes | Yes |
| BLCa86 | Cystectomy | T2b | papillary | High | N/A | Intermediate | No | Hollow |
| BLCa47 | Cystectomy | T3a | papillary | High | Squamous in 80% of tumor | Intermediate | No | Solid |
| BLCa98 | Cystectomy | T3a | papillary | High | N/A | Intermediate | Yes | Solid |
| BLCa33 | Cystectomy | yT3b | papillary | High | N/A | Peritumoral weak | No | Mixed |

PDO cultures were successfully established from BLCa samples irrespective of tumor stage or histological pattern, as determined by pathology reports (42 out of 52 tumors samples, **Fig. 1B, Table 1 and 3**). Although no significant difference was observed between NMIBC and MIBC, organoid-forming efficiency was moderately higher in NMIBC (20 out of 22, 91%) vs MIBC (22 out of 30, 73%, p-value= 0.1608; **Fig. 1B, Table 1**). BLCa PDOs formed within 3 to 7 days and failure to generate PDOs was mostly associated with an insufficient number of viable cells in the biopsy (i.e., due to presence of necrotic tissue) or contamination by microorganisms (see **Table 1** for a detailed report).

Proliferative potential of PDO cultures was investigated and compared to the parental tumors (PT). PT were grouped in low- and high-proliferation rate (less or more than 20% of Ki67^+^ nuclei per section) and correlated to a low (2%±2%) or high (11%±8%) percentage of Ki67^+^ cells in the corresponding PDOs (**Fig. 1C-F, Table S2**). To further investigate the proliferative potential of PDO, their viability was measured at 96h post-seeding for a subset of 10 samples. 7 out of 10 samples showed significant proliferation at 96h compared to the day of seeding, whereas the remaining samples did not show significant alterations of cell number (**Fig. 1G**). No significant difference in viability was observed between NMIBC and MIBC organoids (**Fig. 1G**).

Single-cell RNA sequencing (scRNA-Seq) was performed to assess the cellular heterogeneity of PDOs compared to PTs (n=3, **Fig. H-L**). Cells were grouped based on 12 major cell types distinguished according to cell type-specific marker genes (17,43,44). Cell type classification showed that PDO cultures were enriched for epithelial cells (basal and luminal cells), whereas cancer-associated fibroblasts (CAFs), immune cells and endothelial cells were less prevalent during organoid formation (**Fig. H and L**). As expected, proliferating cells were mainly luminal cells (**Fig. I and L**).

Overall, these data demonstrate that BLCa PDOs can be established from different tumor stages. Most importantly, early-passage PDO cultures are proliferative and despite a lower degree of cellular heterogeneity compared to the PTs, PDOs preserve key cytological features of the original samples.

### PDOs preserve key phenotypic and histological features of parental tumors

PDOs were further characterized by morphological analyses and marker expression (**Table 2-3**). PDOs were grouped by three distinct morphological patterns: solid, hollow, or mixed (representative images reported in **Fig. 2A**). Solid morphology was defined by cellular aggregates showing an outer rim but lacking a clearly identifiable luminal space. Hollow morphology was characterized by the presence of a prominent luminal space, delimited either by a thin layer of cells or by an organized layer of epithelium. Mixed morphology included samples presenting a mix of solid and hollow features as well as cultures with more heterogeneous morphologies, i.e., budding and hybrid organoids. Budding organoids were characterized by the presence of budding structures on the surface of the main organoid whereas hybrid organoids presented both hollow and solid phenotypes at different poles of the organoid (see **Fig. S1** for representative images). 29% of the analyzed organoids had a solid morphology, 44% had a hollow morphology and the remaining 27% had a mixed morphology (**Fig. 2B, Table S3**).

**Figure 2.**
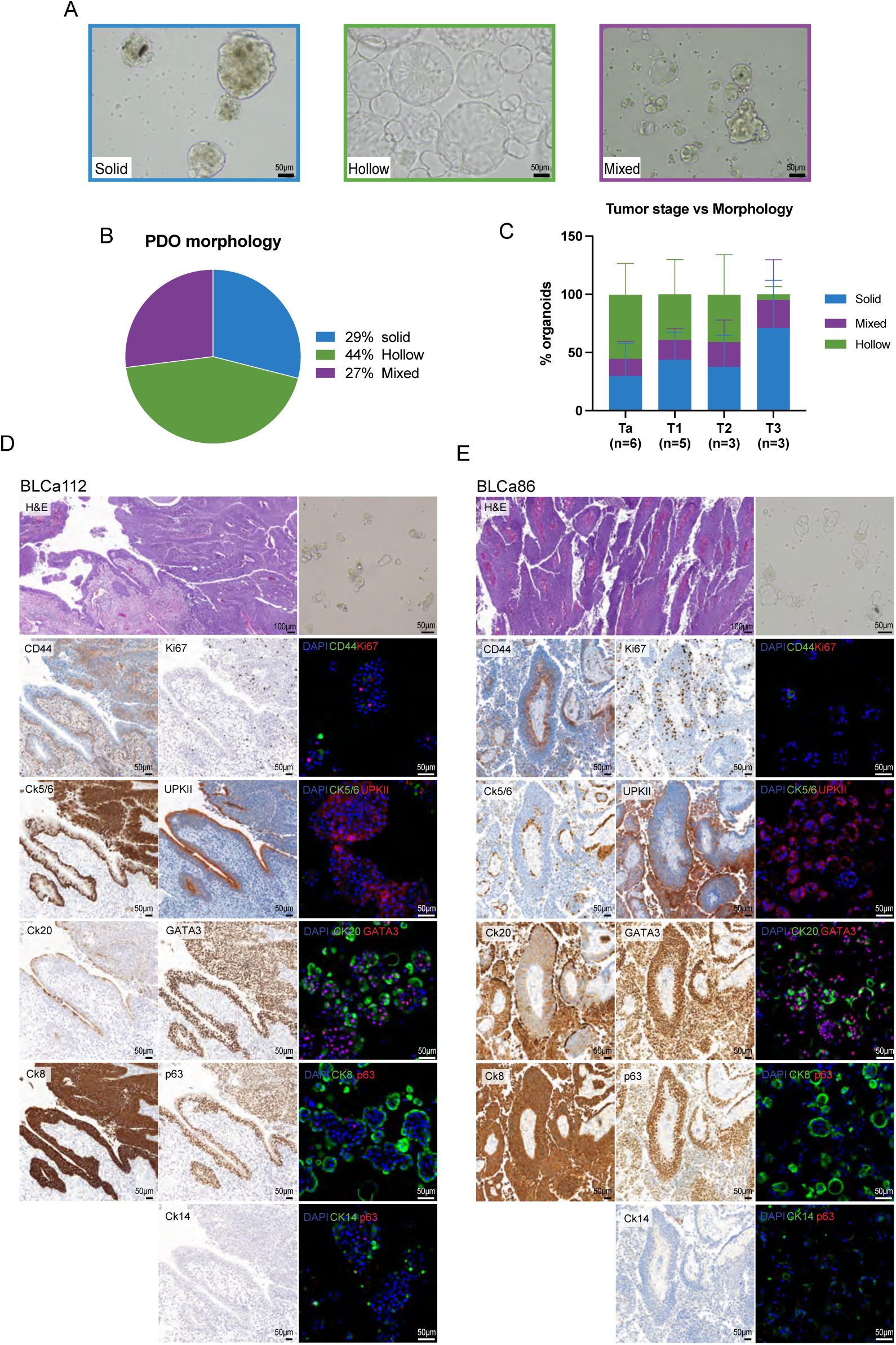
Bladder cancer (BLCa) patient-derived organoids (PDOs) recapitulate original primary tumor (PT) features *in vitro*. (A) Representative brightfield images of BLCa PDOs at passage 1 (p1) with a solid (BLCa50), hollow (BLCa34) or mixed (BLCa69) morphology. (B) % of PDO morphology over the total analyzed samples (n = 929 total counted organoids from 17 samples). (C) Distribution of PDO morphology in samples grouped based on PT stage (mean± SD; n represent the number of patients analysed). Ordinary two-way ANOVA with Turkey’s multiple comparison test between % of organoid morphologies in samples clustered by tumor stages. (D and E) H&E and IHC of PT for indicated markers and brightfield images and whole-mount immunofluorescent staining of PDOs (p1) for indicated markers. Two representative samples non-muscle invasive (BLCa112, E) and muscle invasive (BLCa86, F).

We then investigated the correlation between PDO morphology and PT stage. Although not significant, organoids with predominantly solid morphology originated more frequently from stage T3 tumors (71%±41%) compared to organoids derived from tissue of lower tumor stage (Ta: 30% ± 28%; T1: 44%±23%; T2: 38%±27%; **Fig. 2C, Table S3**). Conversely, stage Ta tumors preferentially gave rise to organoids with hollow morphology (55% ± 27%) compared to higher tumor stages (T3: 5% ± 7%; T2: 41%±34%; T1: 39%±30%; **Fig.2C**). Similar percentages of the mixed morphology organoids were derived from all four tumor stages (Ta: 15%±15%; T1: 17%±10 %; T2: 21%±19%; T3: 24%±34%; **Fig.2C**). We then investigated the correlation between PDO morphology and the presence of concomitant carcinoma *in situ* (CIS) in the PT, an important feature of aggressive disease. In this cohort, 5 out of 17 tumors presented a concomitant CIS, 4 of which generated organoids with a pre-dominant solid morphology, whereas the remaining samples formed mostly hollow organoids (**Table 3** and **Table S3**). PDOs and corresponding PT were evaluated for the expression of basal (CD44, p63, CK 5/6/14) and luminal (uroplakin II (UPKII), GATA3, CK 8/20) markers. The phenotype of PT characterized by predominant expression of either luminal or basal markers was most commonly observed in PDO cultures with PDOs expressing both luminal and basal markers observed less frequently (**Fig. 2D, 2E** and **S2-6)**. Cytosolic markers (CD44, CKs and UPKII) were mainly localized in the outer rim of the organoids (**Fig. 2D, 2E** and **S2-6**). The luminal markers CK20 and GATA3 were mostly expressed in the cytosol and nucleus of the same cells, respectively. In 3 cases, PDOs were positive for both basal and luminal markers and presented basal cells (CD44^+^ or p63^+^) in the core and luminal cells (CK8^+^, CK20^+^ or GATA3^+^) in the outer rim of organoids, suggesting an organized structure (**Fig. 2D** and **S3A**). In 5 cases, both basal and luminal markers could be observed in the outer rim of organoids (**Fig. 2E, S2A, S5A-B** and **S6B**). Interestingly, PDOs derived from two samples longitudinally collected from the same patients (BLCa35 and BLCa40) showed similar histological and morphological patterns (**Fig. S3**).

Basal or luminal marker expression was associated with PDO morphology (**Fig. S7A** and **B**). Solid PDOs were associated with the expression of basal markers, in particular of CD44 (p-value < 0.0001), while hollow organoids were associated with the expression of luminal markers, especially GATA3 (p-value < 0.0001) and UPKII (p-value = 0.0006; **Fig. S7A** and **B**). Mixed organoids, compared to both solid and hollow organoids, were instead associated with the expression of CK8 (p-value = 0.0043 and p-value = 0.0079 compared to solid and hollow organoids, respectively) and CK20 (p-value < 0.0001 and p-value = 0.009 compared to solid and hollow organoids, respectively, **Fig. S7B**).

In addition, basal markers showed a trend for higher positivity in higher tumor stages. In particular, CD44 expression was significantly associated with stage T3 tumors (Ta: p-value = 0.0018; T1: p-value = 0.6788; T2: p-value = 0.0001 compared to T3; **Fig. S7C**). Moreover, PDOs from tumors with a squamous differentiation (BLCa82 and BLCa47), typically associated with advanced stages, predominantly expressed CD44 and showed a comparable staining pattern (**Fig.S4B** and **S6A**). By contrast, the luminal markers were mostly expressed in PDOs derived from lower tumor stages. Notably, CK20 was most highly expressed in organoids derived from stage Ta tumors (T1: p-value = 0.070; T2: p-value = 0.019; T3: p-value = 0.006 compared to Ta; **Fig. S7D**). In parallel, we observed a loss of luminal markers in PDOs from higher stage tumors, such as the significant reduction of GATA3 expression in PDOs derived from stage T3 tumors compared to stage T2 tumors (p-value = 0.0202; **Fig. S7D**).

Taken together, these data indicate that most PDO cultures preserved the predominant phenotype of the PT based on the expression of luminal and basal markers. Moreover, PDO morphology and markers expression were associated with PT histopathological features and stage.

### PDOs retain key genomic features of parental tumors

We determined the genomic landscape of PT and their corresponding matched PDOs (**Table S4**). In terms of tumor cell fractions (i.e., tumor content), values were comparable between PDOs and PTs (**Fig 3A, Table 3, S5** and **S6**; p-value = 0.31), with a trend for higher tumor content in organoids compared to PT (12 out of 15 analyzed pairs). No significant differences in tumor content were detected between NMIBC and MIBC PTs (NMIBC median 0.73±0.18, MIBC median 0.78±0.22, p-value = 0.49) and PDOs (NMIBC median 0.76±0.24, MIBC median 0.75±0.20, p-value = 1).

**Figure 3.**
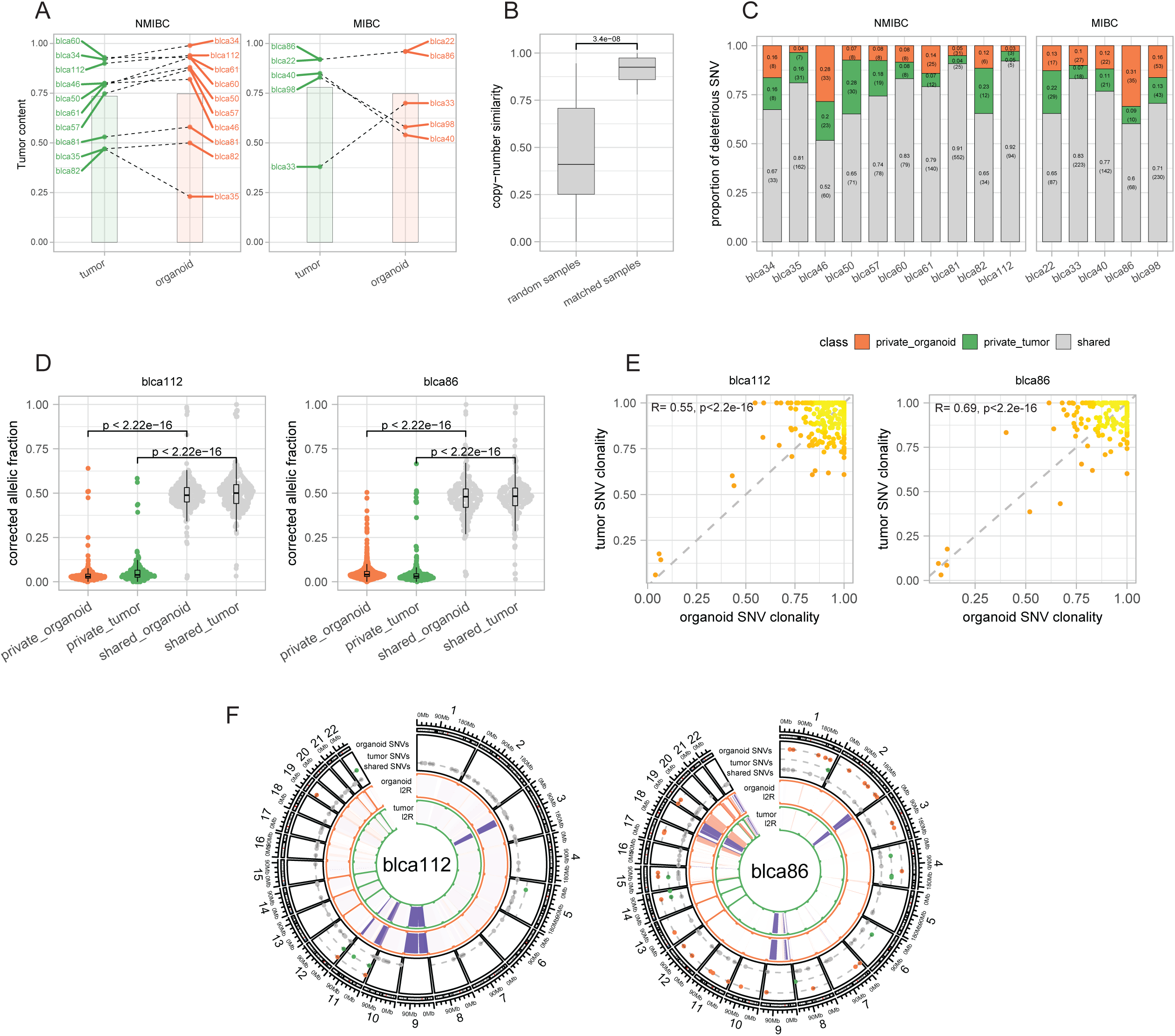
Patient-derived organoids (PDOs) retain key genomic features of parental tumors (PT) (A) Tumor content in PT and matched PDOs of non-muscle invasive bladder cancer (NMIBC, left) and muscle invasive bladder cancer (MIBC, right) samples (n=15). Paired Wilcoxon test, p-value = 0.31. (B) Allele-specific copy-number similarity in randomly paired samples and matched paired ones (PDOs with corresponding PT, n=15). Wilcoxon test between matched and random pairs, p-value = 4.2e-09. (C) Proportion of shared and private deleterious single nucleotide variants (SNVs) between PDOs and PTs. Chi-squared test, p-value < 0.05. (D) Distributions of the tumor content and ploidy corrected allelic fraction (AF) of the shared and private SNVs in PDOs and PTs. Chi-squared test, p-value < 0.05. (E) Clonality of shared point mutations of matched PDOs and PTs (BLCa112 and BLCa86). (F) Copy-number and point mutations profiles between PDOs and PTs for two representative samples (NMIBC BLCa112, MIBC BLCa86).

A genomic similarity metric tailored to copy number changes (see Materials and Methods) indicated that matched PDO/PT pairs were significantly more similar than randomly paired samples (**Fig. 3B**, p-value = 4.2e-09). This is also evident based on the clustering analysis dendrogram (**Fig. S8A**), that further presented a level of similarity determined by the polyploidy feature of a set of tumors.

We further investigated the level of genomic concordance of matched PDOs and PT samples based on deleterious point mutations. Across all sequenced samples, the shared mutations were 73% (± 11.4%) of the detected SNV, with 12.5 % (± 8.1%) and 13.8% (±7.2%) of SNVs exclusively detected in either the PDOs or the TP samples, respectively (**Fig. 3C** and **S8B**).

When comparing the SNVs allelic fraction distributions, we observed significantly higher values for shared mutations (shared fraction) than for mutations found in only one sample of each pair (private fraction; **Fig. 3D** and **S9A**). This result suggests that private point mutations are only present in a small percentage of tumor cells. Nevertheless, several sub-clonal point mutations were also preserved between PTs and PDOs and clonality profiles of shared point mutations were significantly correlated (p-value < 2e-16), hence recapitulating the original tumor heterogeneity (**Fig. 3E** and **S9B**). Side by side analysis of the genomes of paired samples suggest overall concordance (**Fig. 3F**).

We then investigated the correlation between PDO morphology and tumor content, genomic burden and point mutations. No significant tumor content differences were observed in solid (0.64±0.24), hollow (0.87±0.15) and mixed organoids (0.78±0.15; **Fig. S10A, Table S5**). From the genomic point of view, while the mutation load was not significantly different between the three organoid morphologies (hollow: 4.92±0.79, solid: 4.74±0.53, mixed: 5.34±0.75), the genomic burden was significantly higher in solid organoids (0.61+/−0.34) than in organoids that were either hollow (0.06±0.04; p-value = 0.0076) or mixed (0.14±0.07; p-value = 0.0277), reflecting higher load of somatic-copy number aberrations and structural genomic alterations (**Fig. S10C, Table S5**).

Altogether, the data demonstrate that the tumor fractions and the genomic features of PTs are largely maintained in matched PDOs, supporting the preservation of the PT genomic landscape within the models.

### Tumors and their matched organoids recapitulate typical mutational mechanisms of BLCa

NMIBC and MIBC have been proposed to have different genetic backgrounds (11,12). NMIBC predominantly harbors mutations promoting dysregulated growth, whereas MIBC is frequently characterized by mutations causing genetic instability and favoring invasion and epithelial-mesenchymal transition (EMT). We therefore investigated the occurrence of previously identified genomic aberrations of NMIBC (Memorial Sloan Kettering (15)) and MIBC (The Cancer Genome Atlas (45)) patient cohorts.

When comparing measurements of global genomic alterations of our sequenced NMIBC (n=10) and MIBC (n=5) samples, no significant difference in the mutation load (p-value = 0.45) and genomic burden (p-value = 0.45) was observed. Overall, our cohort of PTs and matched PDOs recapitulated the genomic alterations most commonly present in human BLCa (**Fig. 4, Table S6** and **S7**).

**Figure 4.**
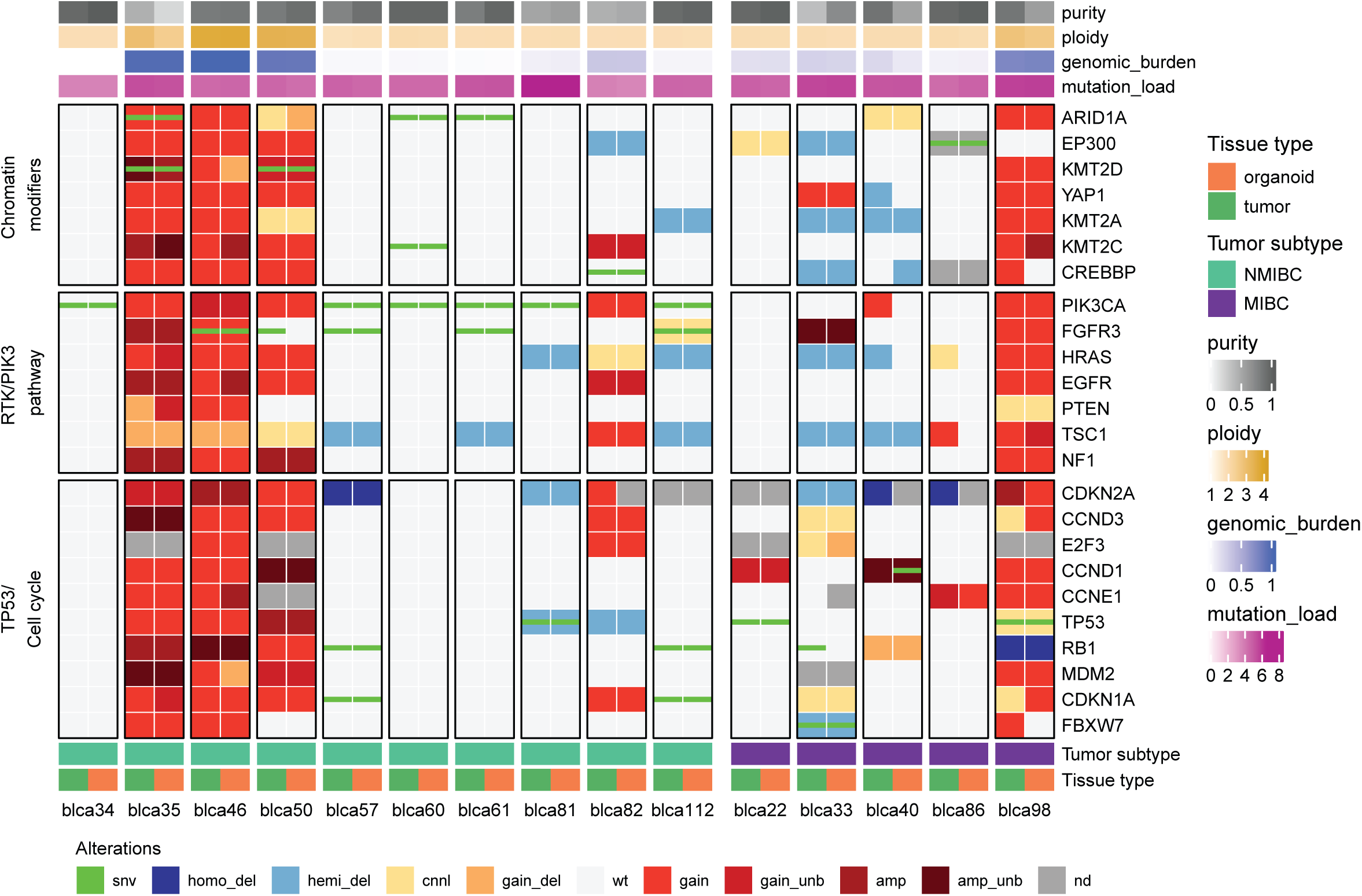
Tumor and matched patient-derived organoids (PDOs) recapitulate typical mutational mechanisms of bladder cancer (BLCa) Mutation heat-map. Samples represented in the columns, with primary tumor on the left and PDOs on the right (n=15). Rows represent genes grouped per pathway. Different types of genomic alterations are indicated in different colors in the bottom and purity, ploidy, genomic burden, and mutation load are reported on the top. Abbreviations: amp, amplification; amp_unb, amplification unbalanced; cnnl, copy number neutral loss; gain_del, gain deletion (gain with LOH); gain_unb, gain unbalanced, homo_del, homo-deletion; hemi_del, hemi-deletion; MIBC, muscle-invasive BLCa; NMIBC, non-muscle invasive BLCa; nd, not determined; snv, single nucleotide variant; wt, wild-type.

In line with previous findings (15), we observed *PIK3/AKT* pathway genes to be frequently mutated in NMIBC with *PIK3CA* and *FGFR3* harboring deleterious point mutations in 60% and 50% of the cases, respectively (**Fig. 4, Table S6**). *FGFR3 and PIK3CA* mutations are considered important driving alterations that support BLCa dysregulated growth especially in NMIBC (14,15). Indeed, no point mutations affecting *FGFR3* and *PIK3CA* genes were identified in the MIBC samples (**Fig. 4, Table S7**). *FGFR3* gene copy number gain instead was observed in 10% of NMIBC and in 20% of MIBC cases, whereas *PIK3CA* gene copy number gain was observed in 40% of both NMIBC and MIBC samples (**Fig. 4, Table S6** and **S7**). Less prevalent driver mutations affecting the *PI3K/AKT* pathway of human BLCa, such as *EGFR* and *HRAS* gene alterations, could be identified in 30% NMIBC and 20% MIBC samples whereas mutations impacting *PTEN* gene were only observed in 20% of NMIBC samples.

According to the central role of tumor suppressor genes in BLCa progression (20), alterations affecting genes with a key role in cell cycle regulation were more prevalent in the MIBC samples (**Fig. 4, Table S6** and **S7**). Whereas 20% of the samples were characterized with *MDM2* gene copy number gain and or point mutations in *RB1* gene in both the NMIBC and MIBC cohorts, 40% of MIBC samples and only 10% of NMIBC harbored deleterious *TP53* point mutations. Moreover, homo-deletion of the *RB1* gene and copy number gain of both *CCND1* and *CCNE1* genes occurred in 20% and 40% of MIBC cohort samples, respectively, but these mutations were not observed in NMIBC cohort samples. Interestingly, *CDKN2A* gene was homo-deleted in 40% of the MIBC and in 10% of NMIBC samples and hemi-deleted in 20% and 10% samples, respectively.

Consistent with previous studies in which mutations affecting epigenetic regulators are believed to be early genomic alterations of BLCa (15,20), we found that chromatin-modifying genes were frequently mutated in both NMIBC and MIBC samples (**Fig. 4, Table S6** and **S7**). Among these, *ARID1A* gene was affected in 30% of the NMIBC samples, whereas *KMT2A* gene was altered in 40% of the MIBC samples. We next performed an SNVs enrichment analysis by using genes harboring at least one deleterious point mutation in the PDOs. This resulted in a total of 16 frequently enriched pathways across the cohort (see Materials and Methods, **Table S8**). Some of these included signaling by *FGFR1, FGFR3* and *ERBB2* and chromatin organization and chromatin modifying enzymes (**Table S8**). Mutations affecting the FGFR signature included genes involved in the activation of FGFR such as FGFs and genes downstream of the FGFR such as *MAPK1&3* and *BRAF*. The signaling by *ERBB2* included mostly mutations directly affecting ERBB receptors such as *ERBB3&4* and downstream signalling mediators such as *AKT1, PIK3CA* and *KRAS*. Finally, the signature of chromatin organization and modifying enzymes included genes known to be relevant for BLCa, such as the *ARID1A* gene, which controls the transcription of specific genes by modifying chromatin structure and the *KMT2D* gene, which is a histone methyltransferase.

Taken together, these data show that the NMIBC and MIBC sample cohorts generated harbor key BLCa mutations, preserve the mutational mechanisms typical of the tumor subtypes and represent the genetic landscape of BLCa.

### BLCa PDOs show heterogenous drug responses to SOC

We next determined drug sensitivity profiles of early passage PDOs with a panel of FDA-approved drugs, including SOC and additional compounds selected based on their clinical relevance, safety in patients and the possibility of drug repurposing.

PDOs showed heterogeneous drug responses to SOC (**Fig. 5A, Table S9**). Effective compounds were selected based on a z-score value (threshold at −1.5, see Material and Methods) and a significant decrease of cell viability compared to vehicle (Adjusted p-value <0.05). Treatment of MIBC PDOs with the MIBC SOC (combination of cisplatin and gemcitabine, cisp/gem combination), significantly reduced PDO viability in 67% of the samples with higher effectiveness than either gemcitabine (p-value = 0.0381) or cisplatin (p-value = 0.039) single treatments. Interestingly, no dose effect of single cisplatin treatment on MIBC PDOs was observed and both methotrexate and vinblastine also had no effect as single compounds. On the other hand, while treatment of NMIBC PDOs with NMIBC SOC (mitomycin C, mmc or epirubicin) showed no significant effect upon the exposure to mitomycin C, epirubicin was widely effective by significantly reducing PDO viability in 50% of the samples. Finally, additional anthracyclines such as doxorubicin significantly reduced PDO viability in 75% of the NMIBC samples whereas daunorubicin was overall less effective (20% of tested NMIBC PDOs).

**Figure 5.**
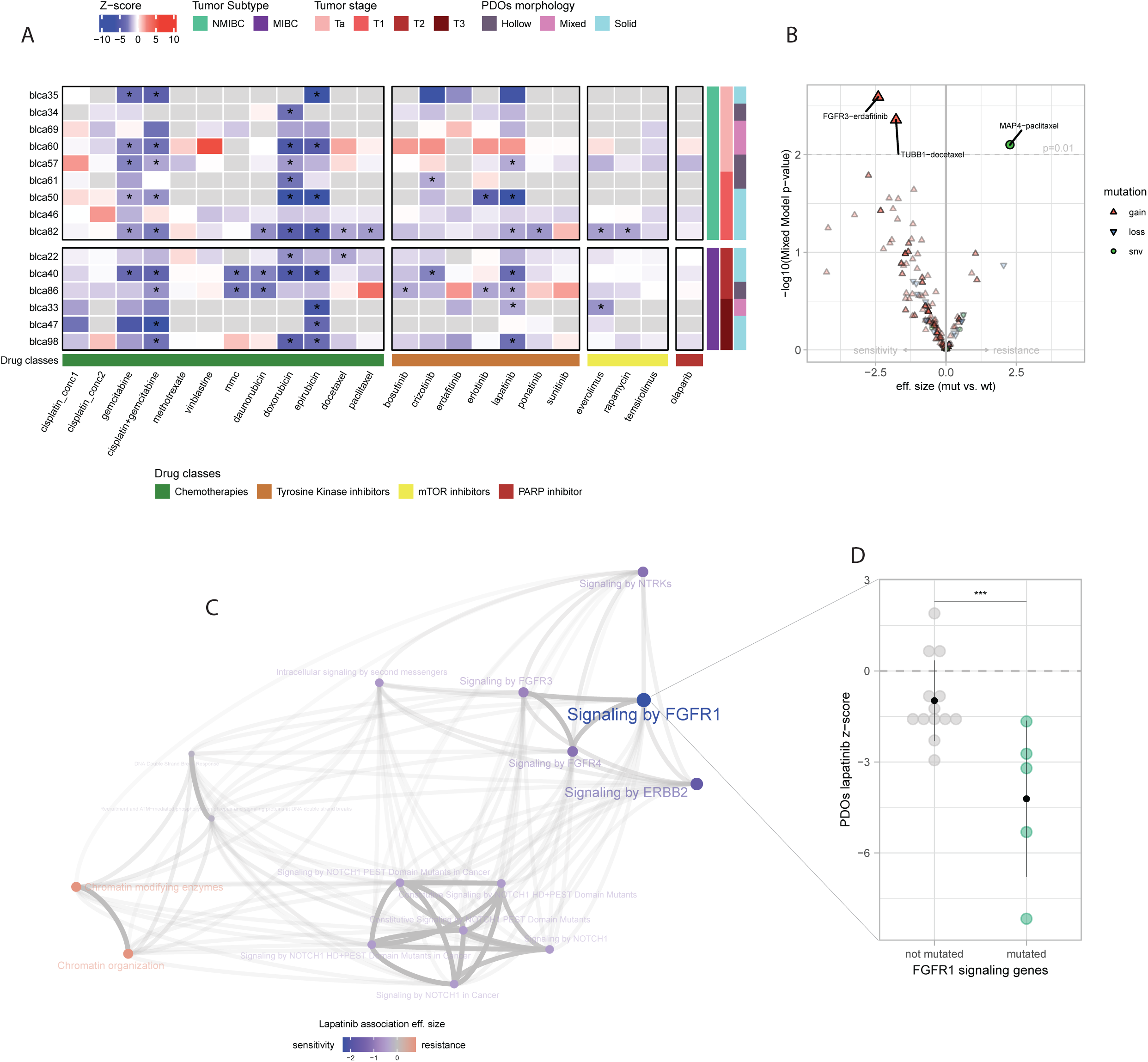
Patient-derived organoid (PDO) drug response and association with gene alterations. (A) Results of PDO drug screen assay on 15 samples. Heatmap reports the average of z-scores normalized to the vehicle values from cell viability assays after 48h exposure of PDOs to drugs (per-replicates data in **Table S8**). Tumor subtype and stage, and PDO morphology are indicated in different colors on the right of the heatmap; not available data are in grey. One-way ANOVA with Dunnet’s multiple comparison test between treatment and vehicle * z-score ≤-1.5 and p-value ≤ 0.05. (B) Genomic association analysis between genomic somatic events and treatment sensitivity/resistance (see Materia and Methods, section “Drugs association analyses”). (C) Association analysis between frequently mutated pathways in PDOs and lapatinib sensitivity. Edges transparency encodes the proportion of shared genes between each term. Node size is proportional to the effect size of the association with lapatinib response (see Materia and Methods, section “Drugs association analyses”). (D) Network depicting frequently enriched terms (see Materia and Methods, section “Drugs association analyses”) showing significant higher sensitivity to lapatinib in PDOs enriching for mutations on FGFR1 signalling genes. *** False discovery rate = 0.02. Abbreviations: conc, concentration; mmc, mitomycin c.

Our cohort covers a wide range of disease stages and therefore might include borderline tumors located between NMIBC and MIBC stages. The strong association of these tumors with tumor progression and poor prognosis complicates disease management (46). Accordingly, and whenever possible, we decided to test the SOC of both NMIBC and MIBC on all samples without distinction between tumor stage (**Fig. 5A**). MIBC PDO viability was frequently reduced by mitomycin C (67%), epirubicin (67%), doxorubicin (75%) and daunorubicin (50%). Interestingly, sample inter-drug analysis comparing viability reduction with respect to the vehicle (i.e., fold-change, see Materials and Methods) showed that compared to the effect of SOC on MIBC, mitomycin C was similarly effective in one sample (BLCa40, SOC: 56%±4%, mmc: 51%±6%). The viability reduction caused by epirubicin instead was significantly higher than SOC in one sample (BLCa40, epirubicin: 95%±1%, p-value = 0.0286) whereas in a second sample (BLCa47, SOC: 47%±18%, epirubicin: 95%) epirubicin effect was moderately higher than SOC but not significantly different. Moreover, epirubicin effect was comparable to SOC in one additional sample (BLCa98, SOC: 28%±54, epirubicin:44%±7%). Similarly, doxorubicin effect compared to SOC was significantly higher in one sample (BLCa40, doxorubicin: 90%±1%, p-value=0.0286), whereas in another sample (BLCa98, SOC: 28%±54, doxorubicin: 57%±10%) despite a high viability reduction caused by doxorubicin, the effects were not significantly different. Similar to MIBC, treatment of NMIBC PDOs with chemotherapeutic drugs (cisp/gem combination) was effective in 62% of the tested samples. Overall, the cisp/gem combination effect was significantly stronger in NMIBC than cisplatin single treatment (p-value = 0.0010) but not compared to gemcitabine alone. Interestingly, treatment with gemcitabine alone was effective in the same 5 NMIBC samples that were sensitive to cisp/gem combination. When tested on NMIBC PDOs, no effect of cisplatin treatment alone was observed at any dose and neither of the cisplatin doses significantly reduced PDO viability in any sample. Finally, additional cytotoxic drugs such as vinblastine and methotrexate were also ineffective on NMIBC PDOs.

Among the microtubule-targeting drugs (taxanes), docetaxel was effective in 25% of tested NMIBC samples and 25% of tested MIBC samples, whereas paclitaxel was effective in 25% of tested NMIBC samples and ineffective on MIBC. In one NMIBC sample (BLCa82) viability decrease with respect to vehicle was comparable in docetaxel (30%±13%) and paclitaxel (34%±1%).

Overall, these results highlight the inter-patient heterogeneity in response to chemotherapies and suggest that PDO-based drug screening may be a useful tool to better stratify patients according to SOC response.

### PDO-based drug screening determines sensitivity to targeted therapies

Next, we used PDOs to screen targeted therapies. All broad-spectrum tyrosine kinase inhibitors (TKIs, i.e., bosutinib, crizotinib and ponatinib) were effective in at least one sample whereas narrow spectrum TKIs either induced a response in PDOs (erlotinib and lapatinib) or failed to reduce the viability of any PDO sample (erdafitinib and sunitinib; **Fig. 5A, Table S9**). Overall, treatments with TKIs targeting similar tyrosine kinases clustered together, showing highly correlated responses across different samples (**Fig. S11A**). Indeed, the FGFR inhibitors ponatinib and erdafitinib were significantly positively correlated (PCC = 0.60, p-value = 0.08, **Fig. S11A**). Across the tested cohort, lapatinib was the most effective treatment, significantly reducing organoid viability in 37% of NMIBC samples and in 67% of MIBC samples. Interestingly, sample inter-drug analysis showed that lapatinib PDO viability reduction was comparable to the one achieved by SOC in two samples (BLCa40 – MIBC, SOC: 56%±4%, lapatinib: 58%±3%; BLCa50 - NMIBC, SOC: 78%±0.03%, lapatinib: 55%±18%; **Fig.5A, Table S9**) and not significantly different in a third one (BLCa98 – MIBC, SOC: 28%±54, lapatinib: 42%12%). Interestingly, one NMIBC sample (BLCa61) that did not respond to SOC epirubicin was significantly sensitive to crizotinib (**Fig.5A, Table S9**).

Furthermore, we tested specific inhibitors of the mTOR signalling pathway (**Fig. 5A, Table S9**). Everolimus was significantly effective in 25% of tested NMIBC samples and in 17% of tested MIBC samples, rapamycin was effective in 33% tested NMIBC samples and ineffective on MIBC, and finally, none of the samples responded to temsirolimus. In one NMIBC sample (BLCa82), we observed a similar viability reduction with respect to vehicle in everolimus (34%±18%) and rapamycin (31%±6%).

Lastly, the PARP inhibitor Olaparib was not effective in any of the samples.

Overall, these findings suggest that the PDO-based drug assay can stratify patients according to the response to targeted-therapies with a potential efficacy on BLCa patients.

### Pharmacogenomic association analysis identifies response biomarkers in PDOs

The comparison between PDO genomic and drug sensitivity profiles revealed significant associations between genomic alterations and drug response. Samples presenting a copy-gain of *FGFR3* (either gain or amplification, see Material and Methods) or *TUBB1* genes were significantly more sensitive to erdafitinib (p-value = 0.0025) and docetaxel (p-value = 0.0044), respectively. Conversely, samples with deleterious mutations in the *MAP4* gene showed increased resistance to paclitaxel (p-value= 0.0079; **Fig. 5B, Table S10**). Given the broad effectiveness of lapatinib treatment on BLCa PDOs and its potential role in personalized therapy for BLCa patients, we investigated the association between lapatinib responses and biological pathways that were found frequently mutated across the PDO cohort (**Fig. 5C, Table S8**). Samples enriched for mutations in genes involved in signaling by *FGFR1, FGFR3, FGFR4, ERBB2* and *NTRKs* signatures, were positively associated with lapatinib response, whereas the genes-sets related to chromatin modification and organization signature were mildly negatively associated. However, by considering a false discovery rate (FDR) of 10%, only the association with the signature of *FGFR1* mediated signaling was statistically significant (FDR = 0.02, **Fig. 5C** and **D, Table S8**).

Next, we compared organoid drug screen data with patient clinical information (**Table 2**). Just few patients received an actual pre- or post-treatment that could allow a comparison between clinical and organoids settings. In case of pre-treated samples, BLCa33 PDOs were derived from a sample treated with cisp/gem combination whereas BLCa57 PDOs from a sample treated with epirubicin. Considering z-scores, organoids derived from these pre-treated samples were not significantly sensitive to the selected treatments (**Fig. 5A, Table S9**).

Finally, we observed an association between PDO morphology and drug sensitivity. Compared to hollow PDOs, a solid morphology was associated with significantly higher sensitivity to docetaxel and erdafitinib (p-value = 0.006, p-value = 0.01, respectively **Fig. S11B** and **C, Table S11**). Tumor stage was not associated with any specific sensitivity.

Taken together, pharmacogenomic analyses highlighted significant associations between genetic variants and drug sensitivity, promoting the identification of novel predictive markers of therapy response with PDOs.

### Clinical application of PDOs for personalized monitoring of tumor reoccurrence and *in vitro* drug sensitivity profiling

Due to a high reoccurrence rate, BLCa patients are often subjected to multiple resections and surgical treatments. In our sample cohort, we included 2 NMIBC patients (patient 1 and patient 2), from whom two samples longitudinally collected during the clinical follow-up were analyzed in detail.

#### Patient 1

Baseline sample (BLCa69) was treatment naïve. Subsequently, the patient underwent local epirubicin treatment (single dose) and experienced a first relapse after 46 days (unsampled; **Fig. 6A**) followed by no treatment and a second relapse after 196 days (relapse; BLCa81) that was followed by local treatment with BCG. The patient then experienced further reoccurrences that were resected 389 days (unsampled), 446 days (sampled, relapse 2), 679 days (unsampled) and 699 days (sampled, relapse 3) after the baseline sample.

**Figure 6.**
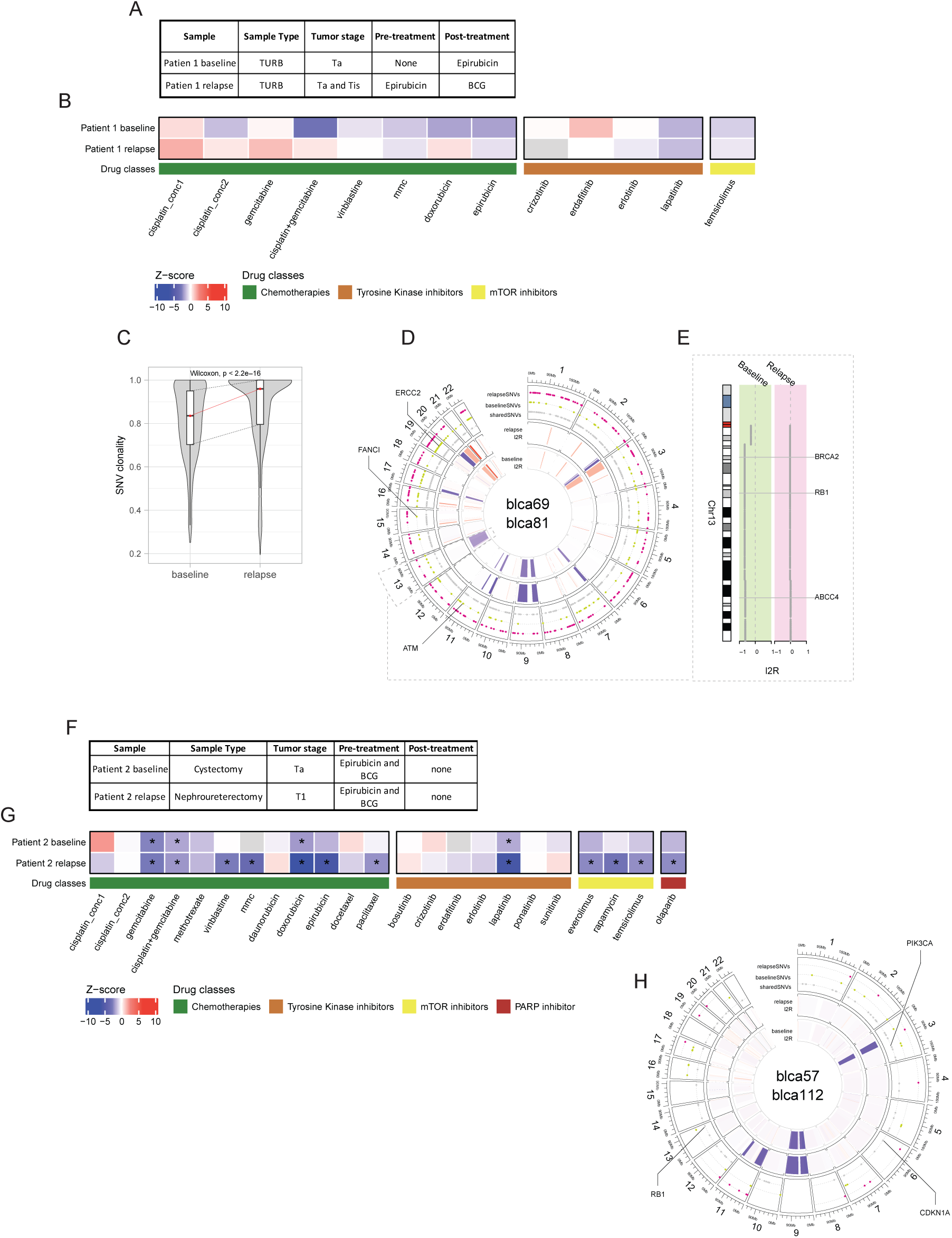
Monitoring tumor recurrence and progression and assessing drug sensitivity *in vitro* with longitudinally patient-derived organoids (PDOs) (A) Patient 1 clinical and pathological information. (B) Results of PDO drug screen assay on patient 1. Heatmap represents the average of z-scores normalized to the vehicle values from cell viability assays after 48h exposure of PDOs to drugs (Single values are listed in **Table S8**). Samples code is indicated on the right side whereas tested drugs on the bottom of the heatmap. Not available data are in grey. One-way ANOVA with Dunnet’s multiple comparison test between treatment and vehicle, * z-score ≤-1.5 and p-value ≤ 0.05. (C) Shared SNV clonality in PDOs baseline and relapse. Paired Wilcoxon test, p< 2.2e^.16^. (D) Copy-number and point mutations profiles between PDO from the baseline and the relapse. Relevant genetic alterations are highlighted. (E) Highlight on chromosome 13 deletion with affected genes. (F) Patient 2 clinical and pathological information. (G) Results of PDO drug screen assay on patient 2. (H) Copy-number and point mutations profiles between PDOs from the baseline and the relapse of patients 2. Relevant genetic alterations are highlighted.

PDOs from both samples (baseline and relapse) were derived and treated with different compounds, including epirubicin (**Fig. 6B**). PDOs from the relapse sample, compared to PDOs from the baseline were significantly less sensitive to epirubicin (z-score: BLCa69, −1.76±0.3; BLCa81, −0.57±0.45; p-value = 0.0286). Remarkably, PDO derived from some of the subsequent relapses (relapse 2 and relapse 3) maintained epirubicin resistance (**Fig. S12A**).

Genomic characterization of patient 1 at baseline and relapse PDOs showed a high mutation load (baseline: 739 SNVs, relapse: 607 SNVs) in both samples. Interestingly, mean clonality of relapse point mutations was significantly higher compared to the one in the baseline (p-value < 2e-16; **Fig. 6C**) suggesting a reduction in tumor heterogeneity. Moreover, SNVs that were lost in the relapse (n=215) were characterized by a significantly lower clonality (i.e., were more sub-clonal; p-value= 0.00017; **Fig. S12B**) compared to those that were conserved.

In contrast to the relapse sample, the baseline PDO was characterized by a large hemi-deletion in chromosome 13 (**Fig. 6D**). This somatic copy number aberration included important genes involved in the DNA damage repair (DDR) pathway such as *BRCA2* and *RB1*, as well as part of the *ABCC4* gene, a multidrug resistance-associated protein also known as *MRP4* (**Fig. 6E**). Additionally, organoids from the baseline sample harbored an *ATM* loss of function (L1348F, Q2733*), genetic alteration in the *FANCI* gene and a point mutation in *ERCC2* (E742K). Both samples had impaired *TP53* gene (R280K).

#### Patient 2

Baseline sample (BLCa57) was obtained from a patient that underwent cystectomy (**Fig. 6F**). This patient was pre-treated with local epirubicin together with 6 cycles of BCG. A relapse occurred in the upper urinary tract 487 days after the surgery. The baseline sample was diagnosed as low-grade Ta whereas the relapse as high-grade T1.

After performing a drug screening on PDOs from baseline and relapsed samples, we observed that organoids from the relapse were significantly more sensitive to epirubicin compared to baseline PDOs (z-score: BLCa57, −1.39±0.68; BLCa112, −5.38±0.12; p-value = 0.0286; **Fig. 6H**). This could be due to the local treatment with epirubicin, while the reoccurrence manifested in the upper urinary tract, a region likely spared from contact with the drug.

While PDOs derived from both samples were sensitive to cisp/gem combination, gemcitabine alone, doxorubicin and lapatinib, PDOs from the relapse were more sensitive to lapatinib and doxorubicin (viability reduction, lapatinib: BLCa57, 47%±7%, BLCa112, 81%±4%; doxorubicin: BLCa57, 57%±8%, BLCa112, 88%±1%) and were sensitive to many additional drugs such as vinblastine, mitomycin c, paclitaxel everolimus, rapamycin, temsirolimus and olaparib.

Genomic characterization of organoids at p1 derived from Patient 2 showed that both samples were characterized by mutation in *CDKN1A* gene (Q29*), a frequently mutated target in BLCa and associated with tumor progression (47), and by *RB1* mutation (S624C; **Fig. 6G**). Moreover, *CDKN2A*, another gene associated with tumor progression, was homo-deleted in both baseline and relapse.

Given the high sensitivity of organoids to *mTOR* inhibition, we investigated the mutational status of genes associated with PI3K-AKT pathway. PDOs from both samples harbored a deleterious point mutation on *PIK3CA* gene (E545K), that is commonly observed in BLCa and results in the expression of a constitutively active PI3K activating downstream AKT and mTOR signaling (48) (**Fig. 6G**). Moreover, in both samples we observed a hemi-deletion of the *TSC1* gene, which forms a complex with *TSC2* that acts as negative regulator of mTORC1 (**Fig. 6G**). These alterations suggested an active mTOR signaling pathway in both baseline and relapse samples.

As shown in the presented cases, longitudinal sampling and PDO generation may be a useful method for detecting variation caused by therapy as well as investigate mechanism at the basis of therapy resistance/sensitivity and/or tumor reoccurrence and progression.

## Discussion

The heterogeneous nature of BLCa is a critical issue that limits therapeutic efficacy and promotes tumor recurrence and progression. To improve response, treatments need to be tailored to each patient to identify more effective compounds, especially for high-risk BLCa patients. This requires the development of reliable customized tumor models with properties that are close to the PT. The majority of BLCa preclinical research has been performed on a small number of cell lines which poorly recapitulate the characteristics of human tumor. Mouse models of BLCa, including genetically engineered, carcinogen-induced and patient-derived xenograft models, have been described but are not suitable for the study of molecular heterogeneity (33,49,50). Over the last few years, personalized medicine approaches have been tested for their feasibility and clinical effectiveness across a variety of cancer types. Organoids may represent an effective tool in this context and several groups developed bladder organoid models (33,34,51–53).

In this study, we established a highly efficient protocol for the generation of BLCa PDOs that cover a wide range of disease stages. The analyses performed on generated BLCa PDOs, confirmed that our PDOs maintain the histopathological and molecular heterogeneity of the PTs in terms of tumor content, deleterious point mutations and tumor cell sub-populations. Organoid genomic profiles were highly comparable to PT profiles and preserved typical BLCa alterations. For instance, alterations affecting the *FGFR3* gene in NMIBC or cell cycle regulators in MIBC that have been proposed in previous studies were confirmed in our cohort (15,20). The concordance of the genomic profiles between PDO and PT has been already demonstrated in previous studies for BLCa (33,54) and upper urinary tract urothelial cancer (UTUC) (55) as well as for other cancer types such as breast (37), ovarian (31), rectal (40), gastrointestinal (39), pancreatic (38,56,57) and prostate cancers (29,32).

Marker analyses confirm that organoids retain the main tumor phenotype. The preservation of tumor sub-clonal mutations in organoids, as shown by the mutation clonality and by the number of high allelic fraction SNVs that are conserved between PTs and PDOs, suggests that the observed similarity of expression markers between organoids and tumor observed in few samples could be caused mostly by biological and technical reasons due to culture conditions rather than through the process of clonal selection. From a biological standpoint, a PDO is a simplified model system composed mostly of tumor cells and lacks the tumor microenvironment components (extracellular matrix (ECM) and other cell types such as CAFs and immune cells). From the technical point of view, instead, the culture conditions used here, and previously reported for prostate organoids and then adapted for bladder organoids, includes few growth factors in the absence of an ECM support (Matrigel) (32). The latter was already reported to control stromal cell overgrowth, thereby increasing drug accessibility and reducing culture time to improve the clinical usability of the tool (32). Furthermore, based on the possibility that the presence of an ECM support might bias organoid cultures towards a more basal phenotype, we decided to opt for a Matrigel-free culture to increase the likelihood of keeping both basal and luminal cells in organoid culture (32–34,58). Our results show that these conditions promote the growth and maintenance of organoids.

Cellular predisposition to acquire a determined morphological phenotype is an intrinsic property of cancer cells. Here we showed how cultured BLCa PDOs can grow accordingly into three different structure types that we described as solid, hollow, and mixed morphologies. The feature of organoids to display morphologies similar to the PT from which they are derived was already observed in different cancer types (e.g., breast, pancreatic, colon and rectal cancers) (36,37,56,59,60). The high tumor fraction conserved in both PTs and PDOs, independently of PDO morphology, suggests no association between organoids’ structure and the prevalence of tumor compared to normal tissue. PDO morphology may rather reflect tumor-specific features. A solid morphology may be associated to more advanced tumors. Solid organoids were indeed mostly derived from high-stage tumors characterized by concomitant CIS. A more advanced phenotype is also supported by the higher genomic burden observed in solid vs. hollow and mixed PDOs. Furthermore, in line with previous analyses of transcriptomic and genomic profiling of aggressive MIBC tumors, solid organoids mostly expressed basal markers compared to hollow organoids (13,19,61). The mixed morphology, defined by the presence of multiple structure types within a single patient-derived culture, instead, was not significantly associated with any specific tumor stage but was linked to luminal marker expression. This morphology might be linked to the diversity of BLCa cell types and originate from intra-sample heterogeneity.

While organoids from healthy tissues were already shown to adopt a hollow morphology compared to solid cancer-derived organoids (55), the observation that different morphologies might also be peculiar to organoids derived from different tumor subtypes is a novel concept. Further experiments to elucidate the functional differences between hollow, solid, and mixed organoids and to understand whether this can be used as criteria to judge PDO quality are necessary. Ultimately, organoid morphologies together with marker expression could be used as a parameter to discriminate between more and less aggressive tumor phenotypes.

We demonstrate the clinical relevance and feasibility of using a BLCa PDO-based drug assay for comparing the SOC with drugs not currently in use for BLCa treatment. Immune-based treatments, including BCG and immune checkpoint inhibitors, that show promising results (62–65) are still challenging to implement in PDO cultures and were not included in the present study. SOC treatments included mitomycin C or epirubicin for NMIBC and the combination of cisp/gem for MIBC. Response to SOC was highly heterogeneous with approximately 50% of NMIBC and 33% of MIBC samples showing no or minimal response. This reflects the inter-patient heterogeneity in the response to chemotherapeutic drugs. Additional chemotherapeutic agents that showed broad effectiveness on NMIBC samples as a single treatment included gemcitabine and doxorubicin. These results are supported by previous studies in which gemcitabine proved to be as effective as intravesical instillation compared to mitomycin c and anthracyclines at reducing both relapse and progression rates but with lower local toxicity (66–71). Similarly, doxorubicin was shown to reduce the risk of reccurrence as an intravesical instillation on NMIBC patients (72). Conversely, cisplatin single treatment was inefficient on NMIBC PDOs, whereas, even if not significant, it showed a more effective trend within MIBC PDOs. In clinical practice, only 30-40% of MIBC patients positively respond to the SOC (24). The heterogeneous responses to SOC observed in the MIBC PDO screening recapitulated these clinical results and additionally provided other chemotherapeutic drugs as candidates for MIBC treatment. These compounds include anthracyclines (doxorubicin and epirubicin) and mitomycin c. Therefore, our findings not only recapitulate current treatment outcomes in real-clinical settings, but also define PDOs as an attractive tool to stratify patients according to a variety of chemotherapeutic drugs that exhibited profound activity in a fraction of patients. In addition to the SOC cisp/gem combination, our study focuses on the assessment of a single agent administration, which in clinical practice might need to be translated into drug combinations that might yield more therapeutic benefits.

We also explored our newly developed PDO-based drug assay to screen for FDA-approved targeted therapies. Drug selection was based on previously reported PDO-based drug screenings from our group (32) as well as additional drugs of relevance for BLCa (73). The latter compounds targeted relevant and frequently altered pathways in BLCa, such as the *FGFR, EGFR/ERBB2* and the *mTOR* pathways. PDO sensitivity to targeted therapies was highly heterogeneous and, in few cases, correlated with the sample-specific genomic background. Overall, TKIs inhibiting similar targets showed a reproducible activity, supporting the biological relevance of our results. In particular, erdafitinib, a drug approved as a second-line treatment for locally advanced or metastatic BLCa, was found to be significantly more effective in patients with somatic copy-number gain alterations in *FGFR3* gene (74,75). Samples that showed higher sensitivity were harboring *FGFR3* copy number gain (BLCa46, BLCa35, BLCa98) or *FGFR3* gene amplification (BLCa33). Within the PDO cohort, we also observed a highly significant sensitivity to lapatinib treatment which is of particular clinical interest in BLCa due to the frequent overexpression of its targets (*EGFR/ERBB2*) (76–78). However, its clinical effectiveness was less evident in different studies, probably due to the insufficient patient stratification or to a limited predictive role of the *EGFR/ERBB2* status (79). Interestingly, in an ongoing trial, trastuzumab, another ERBB2-target drug, is showing positive results in BLCa patients with ERBB2-expressing tumor (80). Samples enriching for mutations in *FGFR1* pathway were also more sensitive to lapatinib activity. Although the correlation was found with respect to the *FGFR1* pathway, the genes that are mutated are not restricted to the *FGFR1* pathway but are also involved in numerous other similar signaling pathways, including lapatinib targets *EGFR/ERBB2*. Therefore, although enrichment for deleterious mutations allows us to formulate hypotheses on the dysfunctionality of a pathway, the integration with other data (e.g., expression) and the availability of a larger cohort would better allow to define this association.

Besides erdafitinib and lapatinib, we identified an association between mutations in *TUBB1* and *MAP4* genes and sensitivity to docetaxel and paclitaxel, respectively (81–84). These microtubule-targeting drugs showed promising results in BLCa both as a single agents and in combination with gemcitabine or carboplatin (85–91).

Understanding the mechanisms involved in therapy resistance and their impact on the evolutionary dynamics of cancer is a central biological question in oncological research. Longitudinal sampling of the same patient at different disease stages can be relevant for the understanding of therapy resistance and clonal selection. In the PDOs from patient 1, we observed decreased sensitivity to epirubicin in the samples collected after epirubicin local treatment. Based on the overall SNV clonality decrease and the loss of sub-clonal SNVs in the relapse compared to baseline, which indicates a lower clonal heterogeneity in the former sample, we hypothesized a drug-induced selection of a pre-existing, epirubicin-resistant population. To support this hypothesis, PDO from the baseline sample were characterized by genetic alterations in key targets of the DDR pathway (*ATM, FANC1, ERCC2, BRCA2* and *RB1*). As the anthracyclines mode of action is based on the induction of double-strand DNA breaks, clones bearing less functional DDR pathways could have a higher susceptibility to these drugs. Sensitivity to DNA damage agents and association with mutations on different DDR genes has been already shown (92–98) in particular, *BRCA2* alterations and the sensitivity to anthracyclines (99,100). While patient 1 underwent pharmacological treatment between the baseline and relapse samples, patient 2 did not receive adjuvant nor neoadjuvant treatment at or following cystectomy. This second study could be particularly relevant for patients diagnosed with UTUCs post-cystectomy, diagnosed in the 2-6% of BLCa cases (101,102). We hypothesize that the UTUC of patient 2 might be caused by a pre-existing malignant transformation in the urothelium as suggested by the “field effect” hypothesis (103). The “field effect” supports that the urothelial carcinoma is often multifocal and the presence of premalignant cells distributed over the epithelium gives rise to cancer upon acquiring additional mutations (103–105). Moreover, many studies claim a clonal origin of UTUCs (106). Detection of UTUCs is complex and often occurs late favoring tumor progression, which would explain the higher tumor stage observed in the UTUC (107). To support this hypothesis, we identified already in baseline sample, a point mutation affecting the *CDKN1A* gene and the homozygous deletion of the *CDKN2A* gene known to be frequently mutated in BLCa and associated with tumor progression from NMIBC (15,47,108,109). Early diagnosis of these UTUCs post-cystectomy is required for timely treatment of patients, improving their clinical outcome. We observed that PDOs from both baseline and relapse patient 2 samples recapitulated a similar drug sensitivity profile. This would suggest that drug response profiles identified at the cystectomy might be informative to choose the best treatment to be applied in an adjuvant setting. Moreover, the observed sensitivity to mTOR inhibitors was supported by the presence of mutations activating the *PIK3-AKT* pathway. Indeed, we identified a mutation affecting the activity of the *PIK3CA* (E545K), commonly observed in BLCa (48,110), together with the hemi-deletion of the mTOR inhibitor, *TSC1* gene (111,112). Therefore, PDOs can aid in the understanding of relevant biological processes such as the “field effect” and tumor progression mechanisms, as well as in finding an effective therapy for this subset of patients diagnosed with UTUCs post-cystectomy.

In summary, with this study we have generated a unique biobank resource of BLCa organoids that encompasses a broad spectrum of disease stages. Moreover, we have demonstrated that PDOs not only retain cancer heterogeneity and mutational burden but can be employed in drug sensitive screens.

## Material and Methods

### Patient clinical characteristics

Fiftytwo bladder cancer samples and matching blood were collected from 41 patients undergoing transurethral resection of the bladder (TUR-B), cystectomies or nephroureterectomy at the Inselspital, University Hospital in Bern. All patients included in this study provided written informed consent (Cantonal Ethical approval KEK 06/03 and 2017-02295).

Clinical details of the patients included in this study are reported in **Table 1**. The patient cohort comprised 37 males and 4 females that at the time of the sampling were at 41 to 91 years of age (median of 68.5 years). A subgroup of 17 (N_NMIBC_ = 11, N_MIBC_ = 6) samples representative of the total patient cohort was selected for further analyses (genomic, morphologic, marker expression and drug screening). Additional clinical details and histopathological evaluation (performed by a certified pathologist) were performed on these samples group and are reported in **Table 2-3**.

### Sample collection

Tumor tissues from TUR-B, cystectomies or nephroureterectomy from patients diagnosed with urothelial BLCa were collected in Dulbecco’s MEM media (Gibco, 61965-026) supplemented with 100 ug/ml Primocin (InVivoGen, ant-pm-1). In case of TUR-B, cold cup biopsies were used for tissue sampling and non-cauterized tissue was selected. For cystectomies and nephroureterectomy, tissue was sampled in the OR immediately after harvesting of the bladder to reduce hypoxic damage of the tissue. Either tumor samples were directly digested for organoid derivation or cryopreserved at −80°C in Fetal Bovine Serum (FBS; Sigma, F7524) with 10%DMSO (Sigma, D2650). Blood was collected in RNA or EDTA blood tubes and stored directly at −80° or white blood cells (WBC) were cryopreserved in FBS/10%DMSO after lysation of erythrocytes in cold EC lysis buffer (150mM NH_4_Cl, 10mM KHCO_3_, 0.1mM EDTA in dH_2_O).

### Tissue digestion and organoid derivation

BLCa tissue was collected in Basis medium (Advanced DMEM F12 Serum Free medium (ThermoFisher, 12634028) containing 2mM GlutaMAX supplement (ThermoFisher, 35050061), 10mM HEPES (ThermoFisher, 15630056) and 100μg/ml Primocin). After mechanical disruption, tumor tissue was washed in Basis medium (200g, 5min) and digested in enzyme mix (5mg/ml collagenase II (ThermoFisher 17101015) dissolved in Basis medium; 15μg/ml DNase I (Roche, 10104159001) and 10μM Y-27632-HCl Rock inhibitor (Selleckchem S1049)). Enzyme mix volume was adjusted so that tumor volume was no more than 1/10 of the total volume. Tissue was incubated at 37°C for 1-2h with mixing every 20min. After digestion tissue was washed with Basis medium (400g, 5min). Pellet was incubated in 5 ml cold EC lysis buffer for 10 min at room temperature and then washed in equal volume of Basis medium (400g, 5min). Cell pellet was suspended in 3-5 ml TrypLE Express (ThermoFisher, 12605028) depending on the pellet volume and incubated at 37°C for 10-15 min with mixing every 5 min. Afterwards cell suspension was passed through 50μm cell strainer (CellTrics, 040042327) and the strainer was extensively washed with 5 ml TrypLE Express and Basis medium. Cells were washed in Basis medium (400g, 4min). Cell pellet was reconstituted in organoid medium and, after determine cell density, cells were seeded in ultra-low attachment (ULA) plates (Corning, 7341582). Generally, 300’000-500’000 cells were seeded per well in 6well plates in 1-1.5 ml medium. Organoids were incubated at 37°C with 5% CO_2_. BLCa organoid media contains the following reagents: Basis medium containing 5% FBS (Gibco, 1027-106), 1x B27 supplement (ThermoFisher, 17504044), 10mM Nicotinamide (Sigma, N0636), 500ng/ml R-Spondin (Peprotch, 120-38), 1.25mM N-acetyl-cysteine (Sigma, A9165), 10μM SB202190 (Selleckchem, S7067), 100ng/ml Noggin (Peprotech, 25038), 10ng/ml Wnt3a (Peprotech, 31520), 50ng/ml hepatocyte growth factor (HGF; Peprotech, 10039), 500nM A83-01 (Tocris, 2939), 50ng/ml epidermal growth factor (EGF; Peprotech, AF-100-15), 10ng/ml fibroblast growth factor 10 (FGF10; Peprotech, 100-26), 10μM Y-27632 Rock inhibitor in basis medium.

Media was stored at 4°C for no longer than one week and it was added to the culture every three days and completely changed after one week.

### DNA isolation from blood, organoids, and tissue samples

DNA was isolated from organoids and blood using the DNeasy Blood and Tissue kit (Qiagen, 69504) according to the manufacturer’s protocol. Snap-frozen tissue was homogenized in 160ul PBS by stainless steel beads (Qiagen, 69989) in TissueLyser MM300 (Qiagen, Germany) at 20Hz for 2x 2min. The lysate was centrifuged at 12’000g for 10min and the supernatant collected for DNA isolation. Subsequently, DNA from tissue samples was extracted with ReliaPrem™ gDNA Tissue MIniprep System (Promega, A2051) according to the manufacturer’s protocol. DNA concentration was assessed with Qubit dsDNA high-sensitivity or broad-range kits (ThermoFIsher, Q33233 and Q33263)

### DNA sequencing

#### Library preparation

Genomic DNA for library preparation is fragmented with Covaris M220 to a target size of 180-220 bp. Libraries for whole exome sequencing are prepared starting from 100 ng gDNA (for the sample derived from FFPE tissue the starting input is 90 ng) with Roche KAPA HyperPrep Kit following the SeqCap EZ HyperCap v2.3 protocol. For the hybridization with Roche SeqCap EZ Human Exome v3.0, up to 11 gDNA samples are multiplexed together mixing 100-200 ng of each library to obtain a combined mass of 1000-1100 μg and then incubated for capture at 47°C for 16-20 hours. Pre- and post-capture libraries are quantified using Qubit dsDNA High Sensitivity Assay and the quality is assessed with Agilent Bioanalyzer High Sensitivity DNA Kit.

### DNA sequencing analysis

#### Genomic analysis pipeline

FASTA files were trimmed using Trimmomatic (113), quality checked performed with FastQC (114), and reads were aligned with BWA algorithm (115) on hg19. Deduplication, realignment around indels and base recalibration were then performed using GATK4 (116). Mutation, copy-number data and samples level statistics were obtained through the recently established SPICE analysis pipeline (45). Briefly, it includes quality control step to assess the similarity between matched samples by running SPIA (117), allele-specific copy number assessment upon data segmentation by running CLONET v2 (118) and mutation and annotation calling via MuTect2 (119) and VEP (120). Sequencing statistics from the pipeline as well as mean coverage are reported in **Table S4**.

#### Allele-Specific Copy Number data analysis

Allele-specific copy-number data analysis was performed on 30 samples, 15 PDO and 15 tissues sample (**Table S4 and S5)** for which data signal was deemed amenable by CLONETv2. The copy-number based similarity (**Figure 3B, Figure S8A**) was assessed exploiting CnA and CnB allele-specific profiles at gene level, using 1 minus the Euclidean-distance between two profiles, followed by a min-max normalization.

#### Deleterious SNVs and SNVs enrichment analysis

First, high quality SNVs were selected by adopting the following filters: minimum tumor coverage ≥ 20, tumor allelic fraction ≥ 0.08, number of tumor alternative reads ≥ 5 and number of normal alternative reads = 0. SNVs were then considered deleterious if their impact on protein function was annotated as medium or high by VEP (120). Chi-squared test was used between number of shared and private SNVs in each sample. Wilcoxon-test instead was used between the allelic fraction of private and shared SNVs in each sample.

To perform the SNVs enrichment analysis, we selected all genes harboring at least one deleterious point mutation and performed an over-representation analysis exploiting Reactome Db gene sets collection (121). For each tested PDO, we selected a subset of enriched terms (Q-value < 0.2) and retained those shortlisted in at least 20% of the study samples (relapse samples BLCa112 and BLCa81 excluded, in **Table S4 and S5**).

### Single cell RNA-sequencing

Organoids were collected collected in Basis medium (100G, 5min) and dissociated into single cells with 1ml TrypLE Express at 37°C for 10 min. Single-cell suspension was counted and cryopreserved in FBS/10%DMSO before scRNA sequencing. To obtain single cells from tumor tissue, tissues were digested as explained in the protocol for organoid derivation, and then cells were counted and cryopreserved in FBS/10%DMSO before analysis.

Slow-frozen single cell suspensions were shipped to the Genomics Facility Basel for scRNA-seq processing. Cryovials were thawed and content transferred into 10 mL Basis medium and incubated for 10 min at room temperature to ensure complete removal of DMSO from the cells. Cells were collected (350g, 5 min), resuspended in 1 mL washing buffer consisting of PBS (Ca^2+^/Mg^2+^-free, Gibco, 10010-015) and 1% BSA (Sigma-Aldrich, A7906) and sequentially filtered through 100 µm (Falcon, 352360) and 40 µm cell strainers (Falcon, 352340). Apoptotic and dead cells were removed by immunomagnetic cell separation using the Annexin Dead Cell Removal Kit (StemCell Technologies, 17899) and EasySep™ Magnet (StemCell Technologies, 18000). Whenever possible and required, repeat dead cell removal was performed to increase cell viability. Next, cells were washed with a resuspension buffer (PBS with 0.05% BSA), spun down and resuspended in a resuspension buffer. Cell numbers and viability were assessed at each step on a Cellometer K2 Image Cytometer (Nexcelom Bioscience, Cellometer K2) using ViaStain AOPI Staining Solution (Nexcelom Bioscience, CS2-0106-5mL) and PD100 cell counting slides (Nexcelom Bioscience, CHT4-PD100-003). Optimal cell concentrations were set according to 10x Genomics protocols (700-1200 cells/µL). Cells were loaded and processed using the 10x Genomics Chromium platform with the Next GEM Single Cell 3’ Reagent Kit v3.1 on 10x Genomics Chromium Single Cell Controller (10x Genomics, PN-120263). 4’000 cells were targeted per sample. Gene expression (GEX) libraries were amplified, pooled and sequenced on the Illumina NovaSeq 6000 platform with addition of 1% PhiX at recommended sequencing depth (20,000-50,000 reads/cell).

### Single-cell RNA sequencing analysis

In the following, we briefly describe the overall steps of the sequencing data analysis. The main processing steps are part of the scRNA-seq analysis workflow scampi (122), enriched with in-house information on selected pathways and marker genes used for cell type classification.

#### Read mapping and quality control

Raw reads were mapped to the GRCh38 reference genome using 10x Genomics Cell Ranger 6.1.1 to infer read counts per gene per cell. Since samples were sequenced on the NovaSeq platform, we performed index-hopping removal using a method developed by Griffiths et al. (123).

After the mapping, the resulting UMI counts were quality controlled and cells and genes filtered to remove known contaminants and potential doublets. Doublet detection was performed using the tool scDblFinder (124). Cells were removed if more than 50% of their reads mapped to mitochondrial genes or if they expressed fewer than 400 different genes. Genes were removed if they are not protein-coding or if they were expressed in less than 20 cells. Subsequently, counts were normalized and corrected for cell-cycle effects and library size using sctransform (125).

#### Unsupervised clustering and cell type annotation

Similar cells were grouped based on unsupervised clustering using Phenograph (126). An automated cell type classification was performed independently for each cell (manuscript in preparation). Cell type annotation was performed using lists of cell type defining, highly expressed marker genes. Marker genes have been selected from previous publications (43,44) as well as based on in-house knowledge on expected tumor markers. Literature-based marker lists have been curated to only select genes that are actually expressed in the cohort.

#### Cohort analysis

For a combined analysis and visualization, all cohort samples were pooled across experiments, followed by a new sctransform normalization to account for library size and additionally including “patient” as a co-variable to the model, to account for the expected patient-specific batch effect. Subsequently, pathway analysis and gene expression analysis and visualization were performed as described above.

### Organoid characterization, proliferation and formation efficiency assays

22 NMIBC and 30 MIBC patient samples were tested to investigate organoid forming efficiency *in vitro* (Table 1–3). Organoid cultures were analyzed at passage 1 (p1), corresponding to a median culture time of 7 days (5 to 16 days of culture). For the subset of 17 samples, the total number of organoids per field and their morphology (solid, hollow or mixed) was manually determined on at least 4 brightfield images per sample (N_NMIBC_ = 11, N_MIBC_ = 6; Table 2 and 3), using cell counter in Fiji (v 2.1.0,(127), **Table S3**).

For the proliferation assay, organoids at p1 were collected and washed in basis medium (100G, 5min) and dissociated into single cells with 1ml TrypLE Express at 37°C for 10 min. Single cells suspension was counted, washed once in basis medium (100g, 5min) and resuspended in BLCa organoid medium. Cells were seeded in ULA 384 well plate (Corning, cat. No. 4588) in 20 ul of BLCa organoids medium at 8’000-10’000 cells per well. After 48h of culture, 20 ul of fresh BLCa organoids medium was added to each well. After a further 48h, CellTiter-Glo 3D assay (Promega, G9682) was used to determine organoids viability, following manufacturer’s instructions. Shortly, 40 ul of CellTiter-Glo 3D reagent were added to each well of the assay. Plates were shaken for 5 min and incubated at 37 °C for 25 min. After incubation, luminescence was measured using Tecan M200 Pro plate reader (Tecan AG).

### Drug screen assay

Organoids were seeded in 348 well ULA plates according to the protocol used for the proliferation assay. After 48h of culture, 20 ul of 2x drugs solutions or vehicle, diluted in complete BLCa medium, were added to organoid cultures. After 48h of drug treatment, CellTiter-Glo 3D assay (Promega, G9682) was used to measure cell viability, following manufacturer’s indications with minor modifications. Shortly, 40 ul of CellTiter-Glo 3D reagent was added per well to the assay plates. Plates were subsequently shaken for 5 min and incubated at 37 °C for 25 min. After incubation, luminescence was measured using Tecan M200 Pro plate reader. Raw counts were normalized independently for each screened sample using the following formula: (X_s_ – X_v_)/SD_v_, X_s_: technical replicates for each drug treatment; X_v_: mean of technical replicates of the matching vehicle conditions; SD_v_: standard deviation of the technical replicates of the matching vehicle conditions (**Table S10)**. Z-scores of cisplatin, gemcitabine as well as their combinations were generated using H_2_O as vehicle, while z-scores of all remaining drugs were generated using DMSO as vehicle. Raw counts were also used to generate fold-changes with respect to the average of vehicle raw values. From fold-change values the actual cell growth inhibition was calculated for each condition as 1 – fold-change.

### Drugs association analyses

Drug screening z-scores were used to perform association analyses with both PDOs genomic and phenotypic information. For the former, for every drug and for every corresponding target gene (**Table S11**), we tested the association between drugs activity and gene mutational status by fitting a random intercept Linear Mixed Model (LMM):

*z* ~ *β*_1_*x* + *β*_0_ *m* + *ϵ*, where *z* is a vector of z-scores for a drug *d* across tested PDOs replicates from different samples, *x* is a binary vector encoding the mutation status of each PDO (0 if *wt* and 1 if *mutated*) and *m* is a vector encoding the PDO sample to which each replicate belongs, treated as a random effect. Estimated *β*_1_ was considered as the effect-size of the resulting association. Corresponding p-values were obtained from a Likelihood Ratio Test comparing the shown model, to a null one (lacking of *β*_1_*x* term).

Three classes of genomic aberrations were considered: copy-gain (one of gain, gain_unb, gain_del, amp, amp_unb, amp_del), copy-loss (one of homo_del, hemi_del) and deleterious SNVs. Only drug-target pairs with at least 3 aberrant and 3 wild-type models were tested (**Table S10**). The same strategy was adopted for the drug-phenotype association analysis, where the binary vector *x* encoded the phenotype information of two compared groups (*solid vs. hallow morphology, MIBC* vs *NMIBC*). SNVs-Enriched-pathway association analysis (**Figure 5C**) was performed by considering, for each frequently reactome term previously identified (see Deleterious SNVs and SNVs enrichment analysis), the presence or absence of enrichment, for each PDO. FDR correction was then performed. The binary matrix reported in **Table S8** reports the groups considered for each term.

### Whole-mount immunofluorescence staining of organoids

Organoids were collected in Basis medium and spin down (100g, 5 min). Organoids were suspended in 1xPBS and collected in a 96Well Round (U) Bottom plate (Sigma, 92097) and spin down (100g, 5 min). Medium was removed and organoids fixed in 4% paraformaldehyde (PFA) for 30 min at room temperature. PFA was removed and organoids washed three times in 1xPBS (100g, 5 min). Organoids were then permeabilized in 0.3% Triton-X (Sigma, T8787) for 10 min at room temperature and then blocked in 10% donkey serum (Lubio, 017-000-121) in PBS/0.05% Tween (Sigma, P4780) for 1h at room temperature gently rocking. After the blocking, the organoids were incubated with primary antibodies in blocking solution overnight at 4°C. After incubation with primary antibodies (1:100), organoids were washed three times (100g, 5min) in PBS/0.05% Tween. Organoids were then incubated with secondary antibodies AlexaFluor 555 donkey anti-rabbit (1:200, ThermoFisher, A21434), AlexaFluor 488 donkey anti-mouse (1:200, ThermoFisher, A21202) and DAPI (1:1000, ThermoFisher, 62248) in blocking solution for 2h at room temperature gently rocking. At the end, organoids were washed three times (100g, 5min) in PBS/0.05% Tween-20, gently rocking and transferred to flat glass bottom, black walled 96-well plates (Corning, cat. No. 4580) in PBS/0.05% Tween-20. The antibodies used are listed in Table 3. Immunofluorescence staining was imaged using a confocal microscope (ZEISS LSM 710 with Airyscan). Image analysis was performed with QuPath v.0.2.3 (Queen’s University, Belfast, Northern Ireland) and Fiji (v 2.1.0, Schindelin et al. 2012).

### Parental tumor histology

FFPE tissue from the tumor samples was sectioned and stained by haematoxylin and eosin (H&E) and immunohistochemistry (IHC). Brightfield images of tissue sections were acquired with slide scanner (3DHistech Pannoramic 250 Flash II). All experiments were performed by the Translational Research Unit (TRU) at the Institute of Pathology, University of Bern or performed in house. Image analysis was processed in QuPath v.0.2.3 (Queen’s University, Belfast, Northern Ireland).

## Acknowledgments

We thank all the patients that participated in our study, as well as all the involved surgical teams and study nurses, in particular Ms. Sofia Bonné and Mr. Anselm Lafita. We would like to acknowledge the Clinical Bioinformatics Unit at the NEXUS for the scRNA sequencing and analysis and the Translational Research Unit (TRU) at the Institute of Pathology, University of Bern for the immunohistochemistry. Finally, we would like to thank Peter Grey for manuscript revision. This project received funding from Swiss National Science Foundation (189149 to M.K.D.J. and 184933 to R.S.), Swiss 3R Competence Center (OC-2019-003 to M.K.D.J.), and the European Research Council (ERC) under the European Union’s Horizon 2020 research and innovation program (648670; SPICE to F.D.).

## Author contributions

M.M. designed experiments, acquired data, interpreted data and wrote manuscript. T.C. performed bioinformatics data analysis, interpreted data and revised the manuscript. M.K. performed initial experiments on clinical samples and revised the manuscript. T.F. performed initial bioinformatics data analysis. F.L.M. provided technical support and revised the manuscript. S.K. performed initial experiments on clinical samples and provided technical support. V.G. and A.R. provided pathological evaluation. I.K. performed immunohistochemistry. P.G. performed sequencing experiments and revised the manuscript. B.K., G.N.T. and R.S.B provided clinical samples, interpreted data, and revised manuscript. F.D. designed genomic experiments, interpreted data and wrote manuscript. M.K.D.J. designed concept of the study and experiments, interpreted data and wrote manuscript.

