## Supplementary material for "Bladder cancer organoids as a functional system to model different disease stages and therapy response": Suplemental figures

Figure S1.

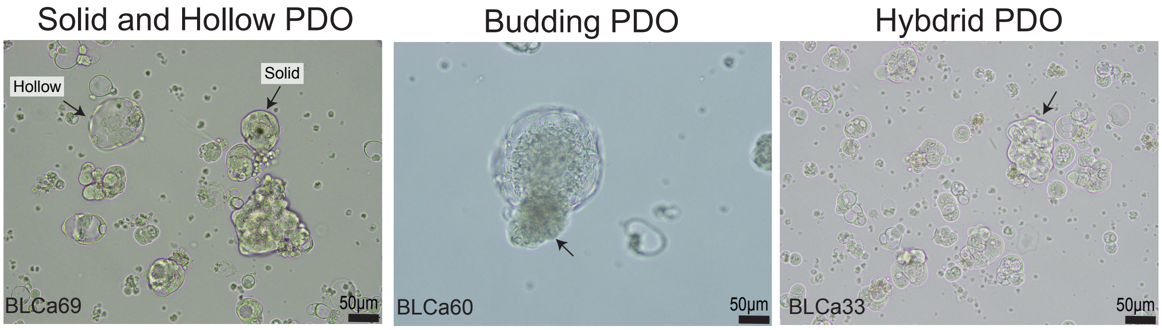
Solid and Hollow PDO
Hollow
Solid
BLCa69
50µm
Budding PDO
BLCa60
50µm
Hybdrid PDO
BLCa33
50µm

Figure S2.

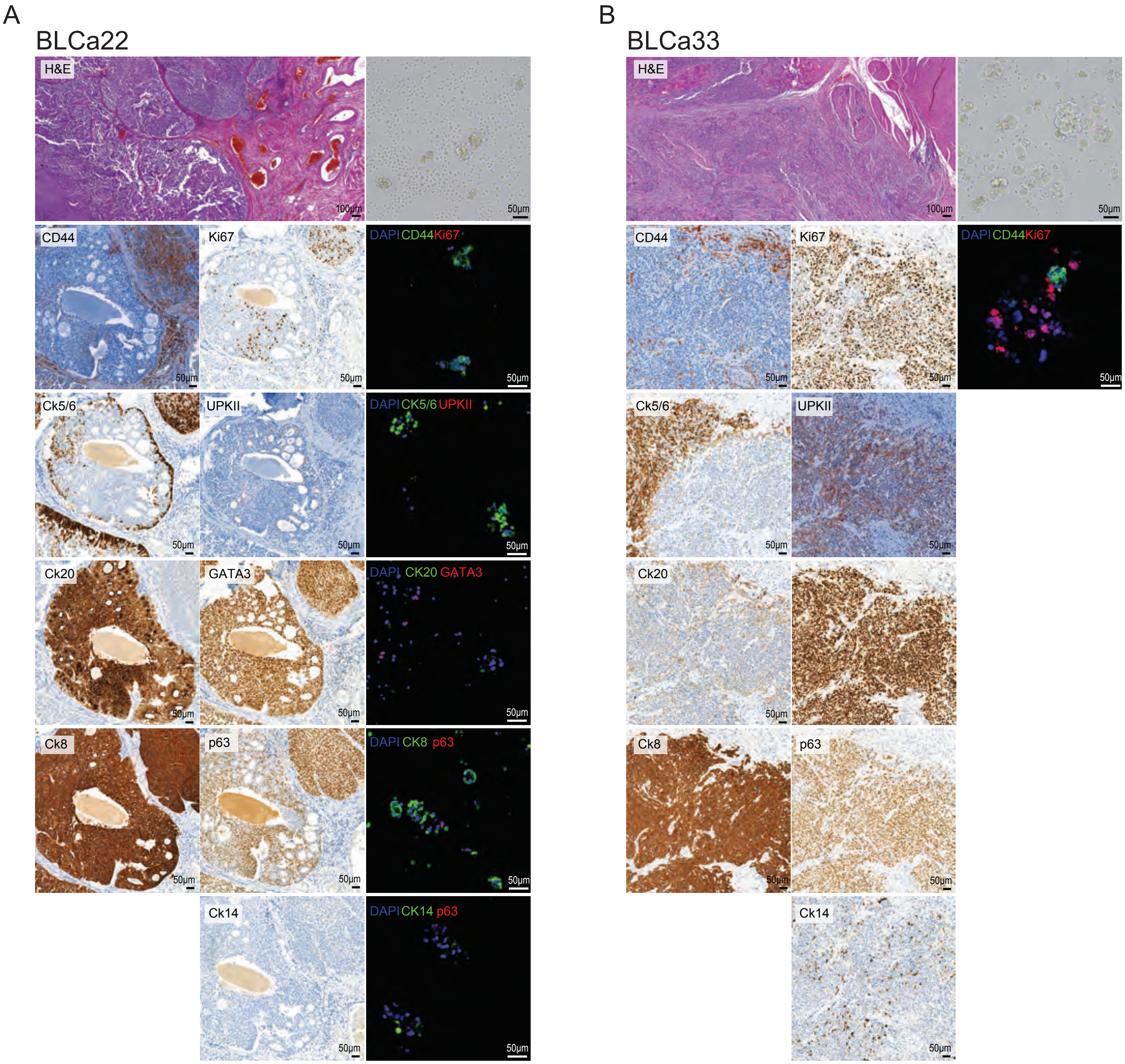
A
BLCa22
H&E
100µm
50µm
CD44
Ki67
DAPI CD44 Ki67
Ck5/6
UPKII
DAPI CK5/6 UPKII
Ck20
GATA3
DAPI CK20 GATA3
Ck8
p63
DAPI CK8 p63
Ck14
DAPI CK14 p63
B
BLCa33
H&E
100µm
50µm
CD44
Ki67
DAPI CD44 Ki67
Ck5/6
UPKII
Ck20
Ck8
p63
Ck14

Figure S3.

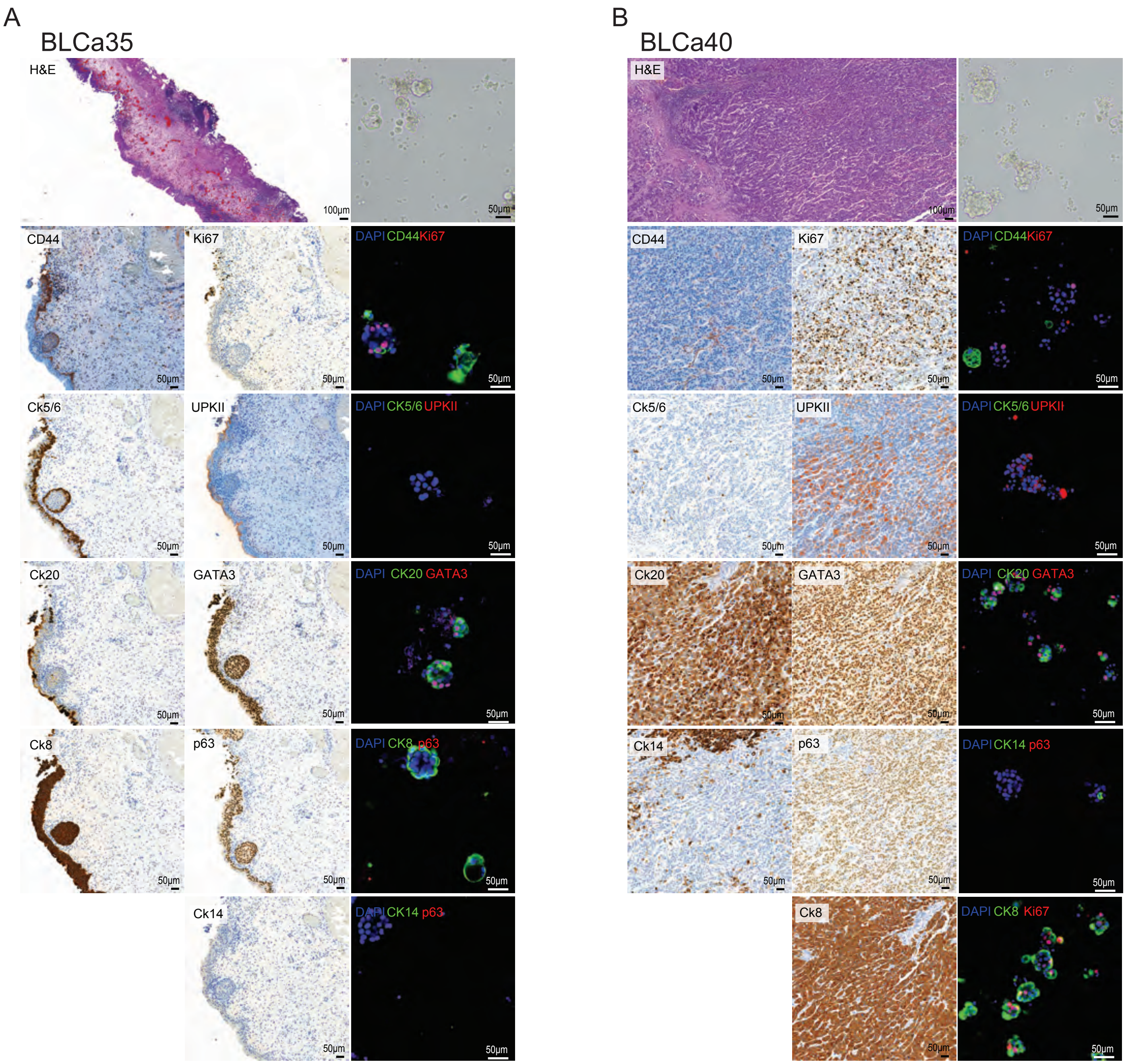
A
BLCa35
H&E
100μm
50μm
CD44
Ki67
DAPI CD44 Ki67
Ck5/6
UPKII
DAPI CK5/6 UPKII
Ck20
GATA3
DAPI CK20 GATA3
Ck8
p63
DAPI CK8 p63
Ck14
DAPI CK14 p63
B
BLCa40
H&E
100μm
50μm
CD44
Ki67
DAPI CD44 Ki67
Ck5/6
UPKII
DAPI CK5/6 UPKII
Ck20
GATA3
DAPI CK20 GATA3
Ck14
p63
DAPI CK14 p63
Ck8
DAPI CK8 Ki67

Figure S4.

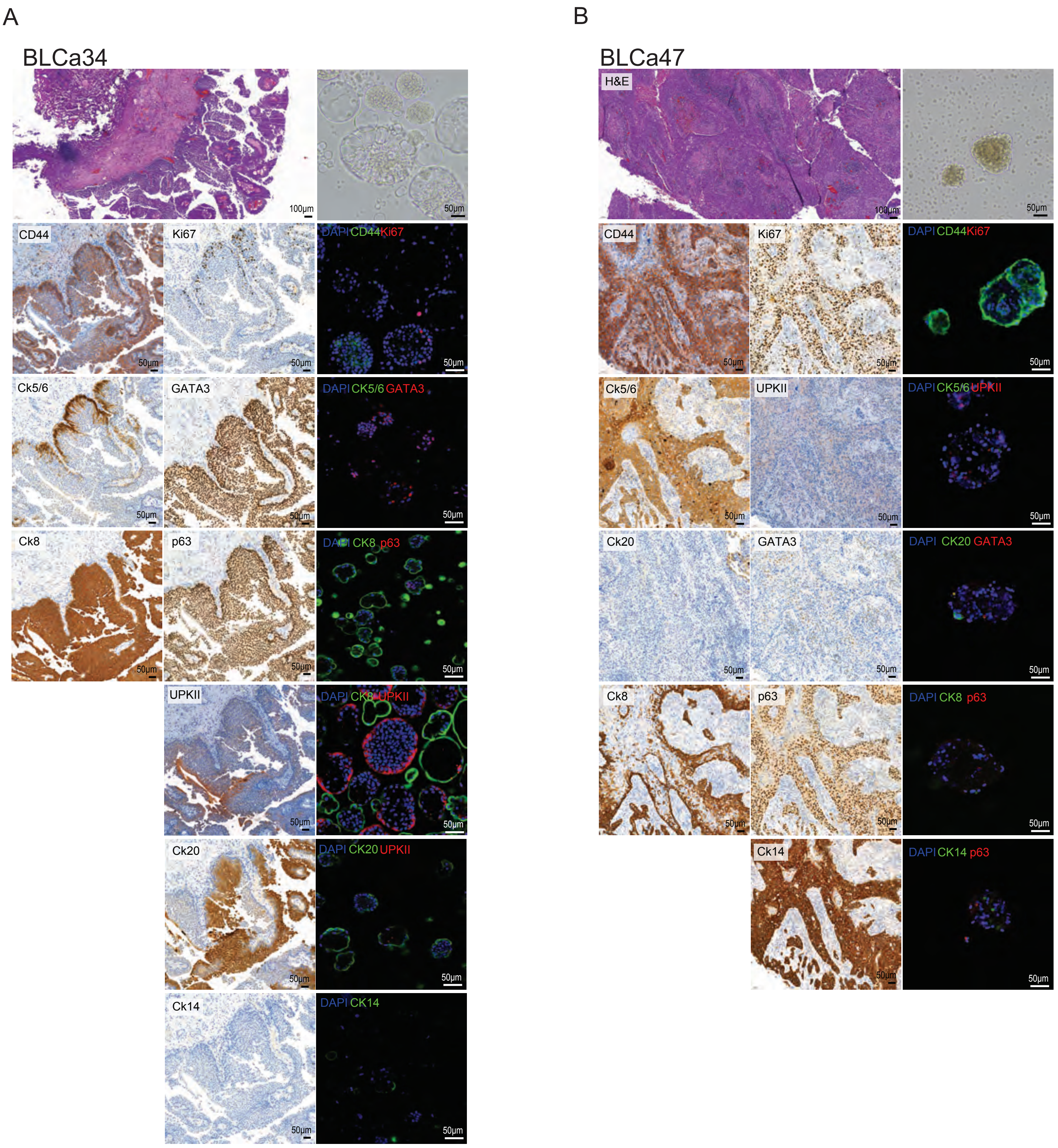
A
BLCa34
100µm
50µm
CD44
Ki67
DAPI CD44 Ki67
Ck5/6
GATA3
DAPI CK5/6 GATA3
Ck8
p63
DAPI CK8 p63
UPKII
DAPI CK8 UPKII
Ck20
DAPI CK20 UPKII
Ck14
DAPI CK14
B
BLCa47
H&E
100µm
50µm
CD44
Ki67
DAPI CD44 Ki67
Ck5/6
UPKII
DAPI CK5/6 UPKII
Ck20
GATA3
DAPI CK20 GATA3
Ck8
p63
DAPI CK8 p63
Ck14
DAPI CK14 p63

Figure S5.

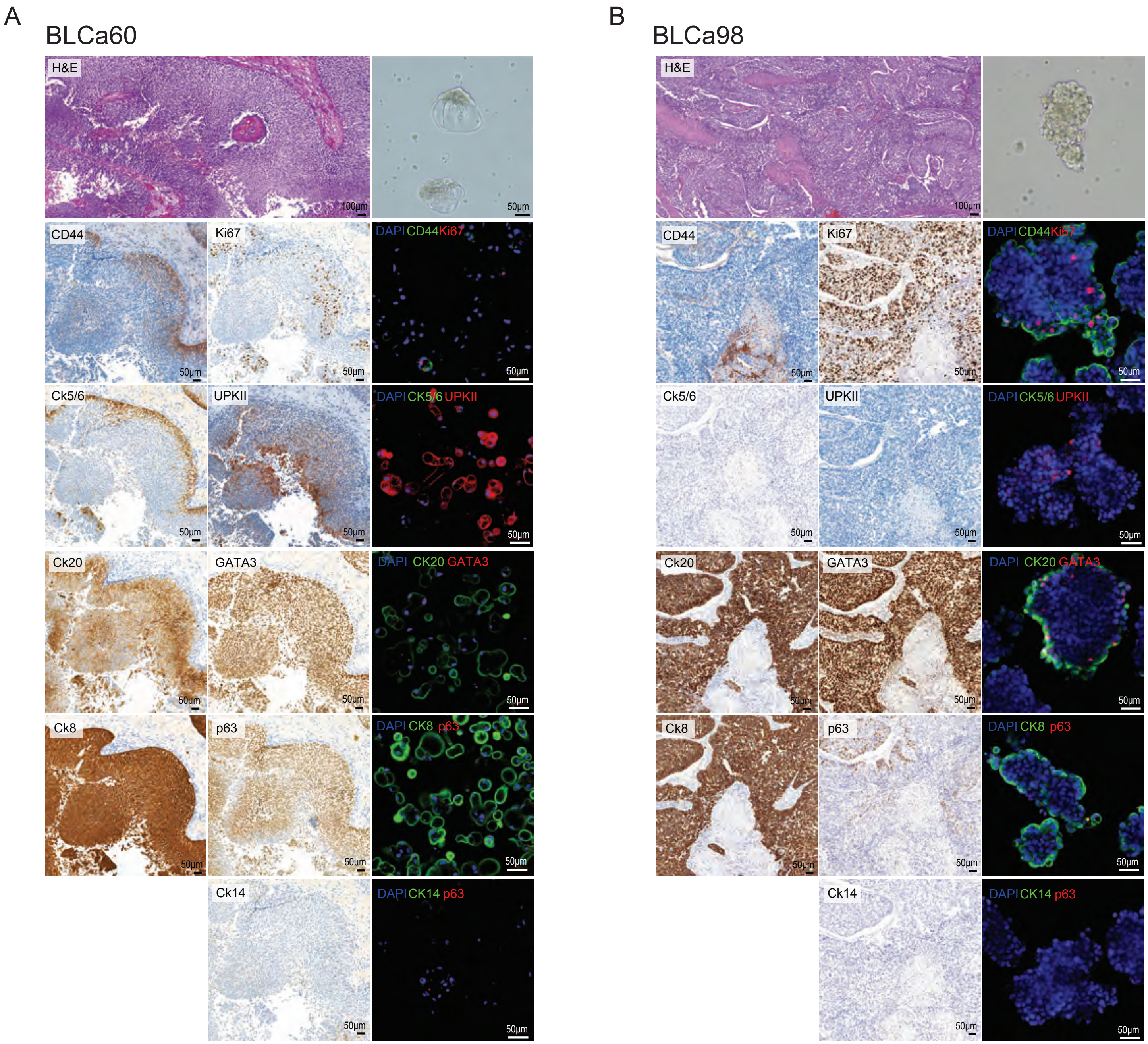
A
BLCa60
H&E
100μm
50μm
CD44
50μm
Ki67
50μm
DAPI CD44 Ki67
50μm
Ck5/6
50μm
UPKII
50μm
DAPI CK5/6 UPKII
50μm
Ck20
50μm
GATA3
50μm
DAPI CK20 GATA3
50μm
Ck8
50μm
p63
50μm
DAPI CK8 p63
50μm
Ck14
50μm
DAPI CK14 p63
50μm
B
BLCa98
H&E
100μm
CD44
50μm
Ki67
50μm
DAPI CD44 Ki67
50μm
Ck5/6
50μm
UPKII
50μm
DAPI CK5/6 UPKII
50μm
Ck20
50μm
GATA3
50μm
DAPI CK20 GATA3
50μm
Ck8
50μm
p63
50μm
DAPI CK8 p63
50μm
Ck14
50μm
DAPI CK14 p63
50μm

Figure S6.

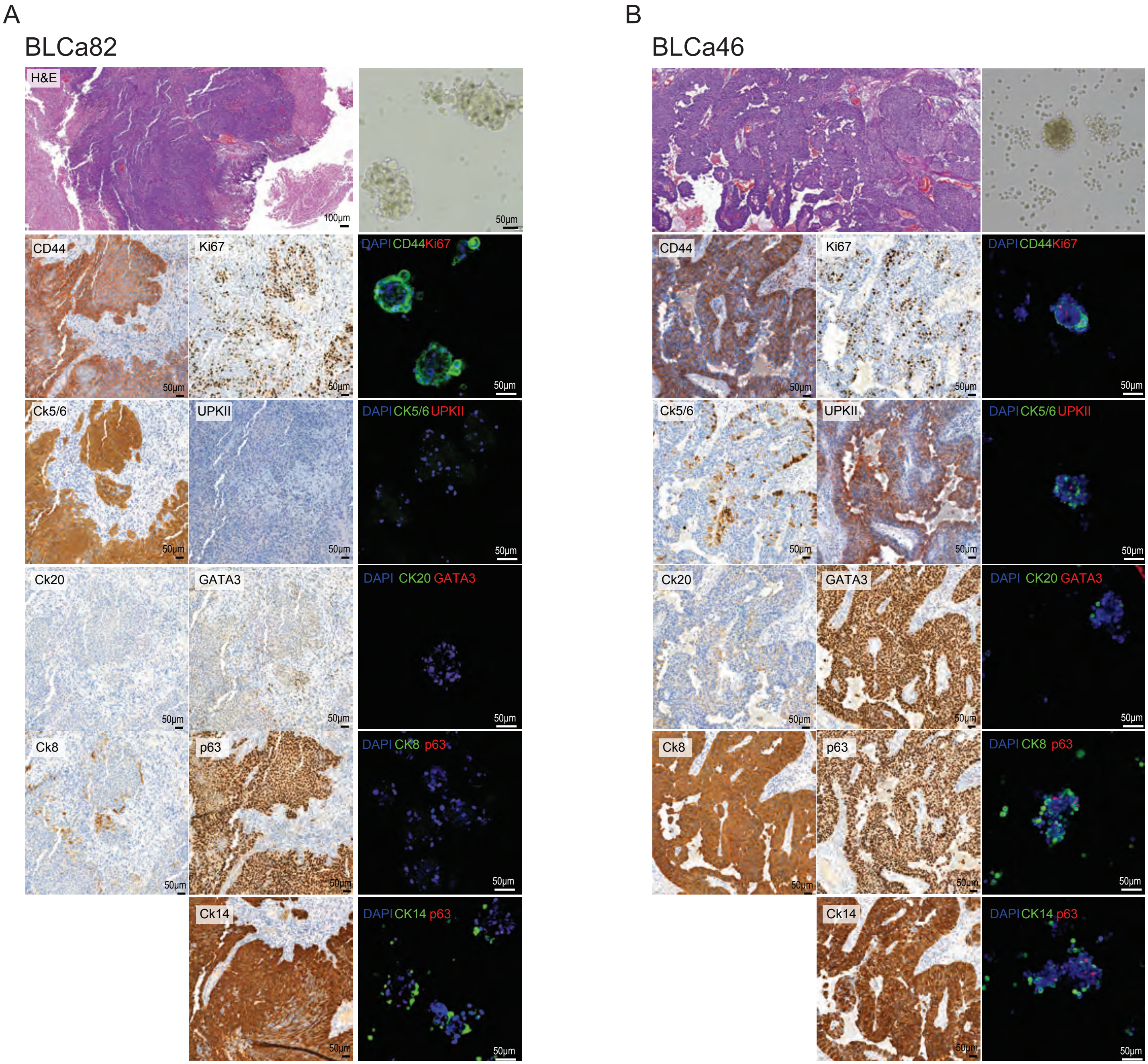
A
BLCa82
H&E
100µm
50µm
CD44
Ki67
DAPI CD44 Ki67
Ck5/6
UPKII
DAPI CK5/6 UPKII
Ck20
GATA3
DAPI CK20 GATA3
Ck8
p63
DAPI CK8 p63
Ck14
DAPI CK14 p63
B
BLCa46
CD44
Ki67
DAPI CD44 Ki67
Ck5/6
UPKII
DAPI CK5/6 UPKII
Ck20
GATA3
DAPI CK20 GATA3
Ck8
p63
DAPI CK8 p63
Ck14
DAPI CK14 p63

Figure S7.

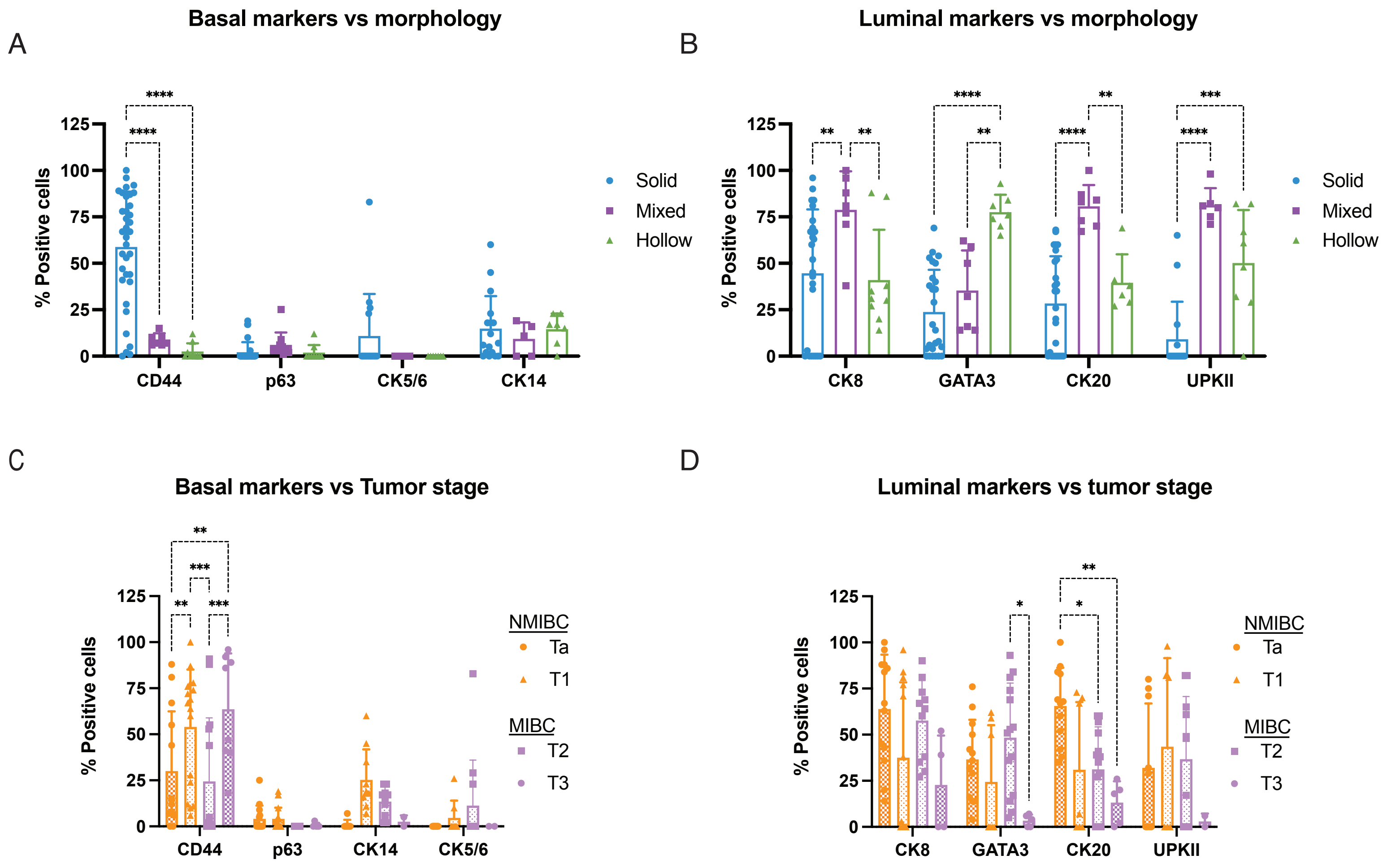
A
Basal markers vs morphology
% Positive cells
125
100
75
50
25
0
****
****
CD44
p63
CK5/6
CK14
Solid
Mixed
Hollow
B
Luminal markers vs morphology
% Positive cells
**
**
****
**
**
****
****
***
****
CK8
GATA3
CK20
UPKII
Solid
Mixed
Hollow
C
Basal markers vs Tumor stage
% Positive cells
**
***
**
***
CD44
p63
CK14
CK5/6
NMIBC
Ta
T1
MIBC
T2
T3
D
Luminal markers vs tumor stage
% Positive cells
*
**
*
CK8
GATA3
CK20
UPKII
NMIBC
Ta
T1
MIBC
T2
T3

Figure S8.

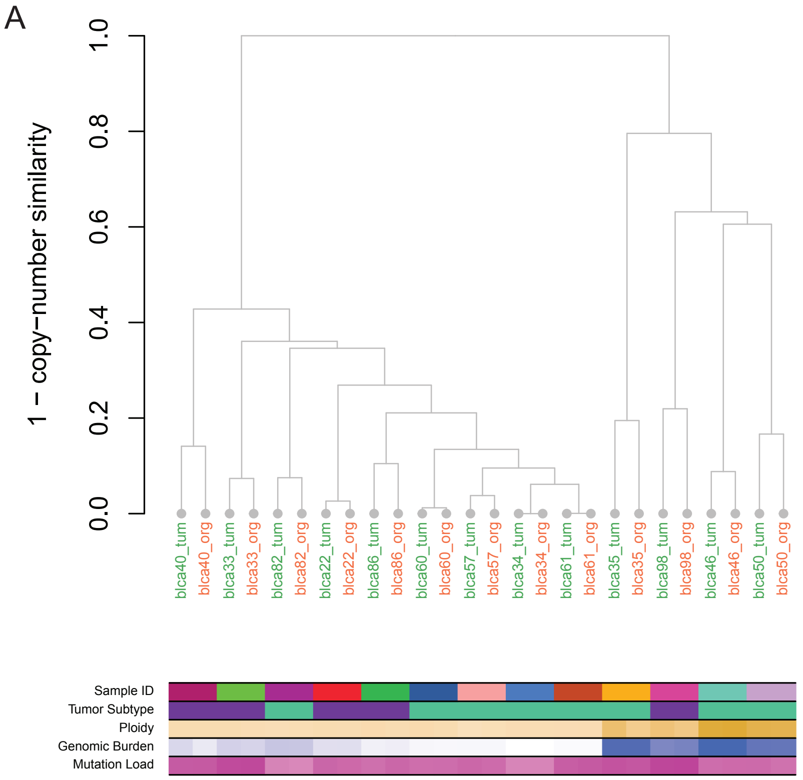
A
1 − copy−number similarity
0.0
0.2
0.4
0.6
0.8
1.0
blca40_tum
blca40_org
blca33_tum
blca33_org
blca82_tum
blca82_org
blca22_tum
blca22_org
blca86_tum
blca86_org
blca60_tum
blca60_org
blca57_tum
blca57_org
blca34_tum
blca34_org
blca61_tum
blca61_org
blca35_tum
blca35_org
blca98_tum
blca98_org
blca46_tum
blca46_org
blca50_tum
blca50_org
Sample ID
Tumor Subtype
Ploidy
Genomic Burden
Mutation Load

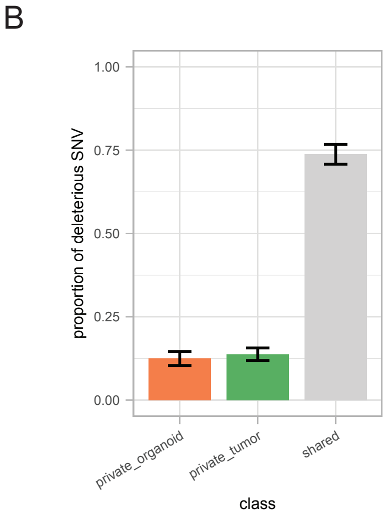
B
proportion of deleterious SNV
0.00
0.25
0.50
0.75
1.00
private_organoid
private_tumor
shared
class

Figure S9.

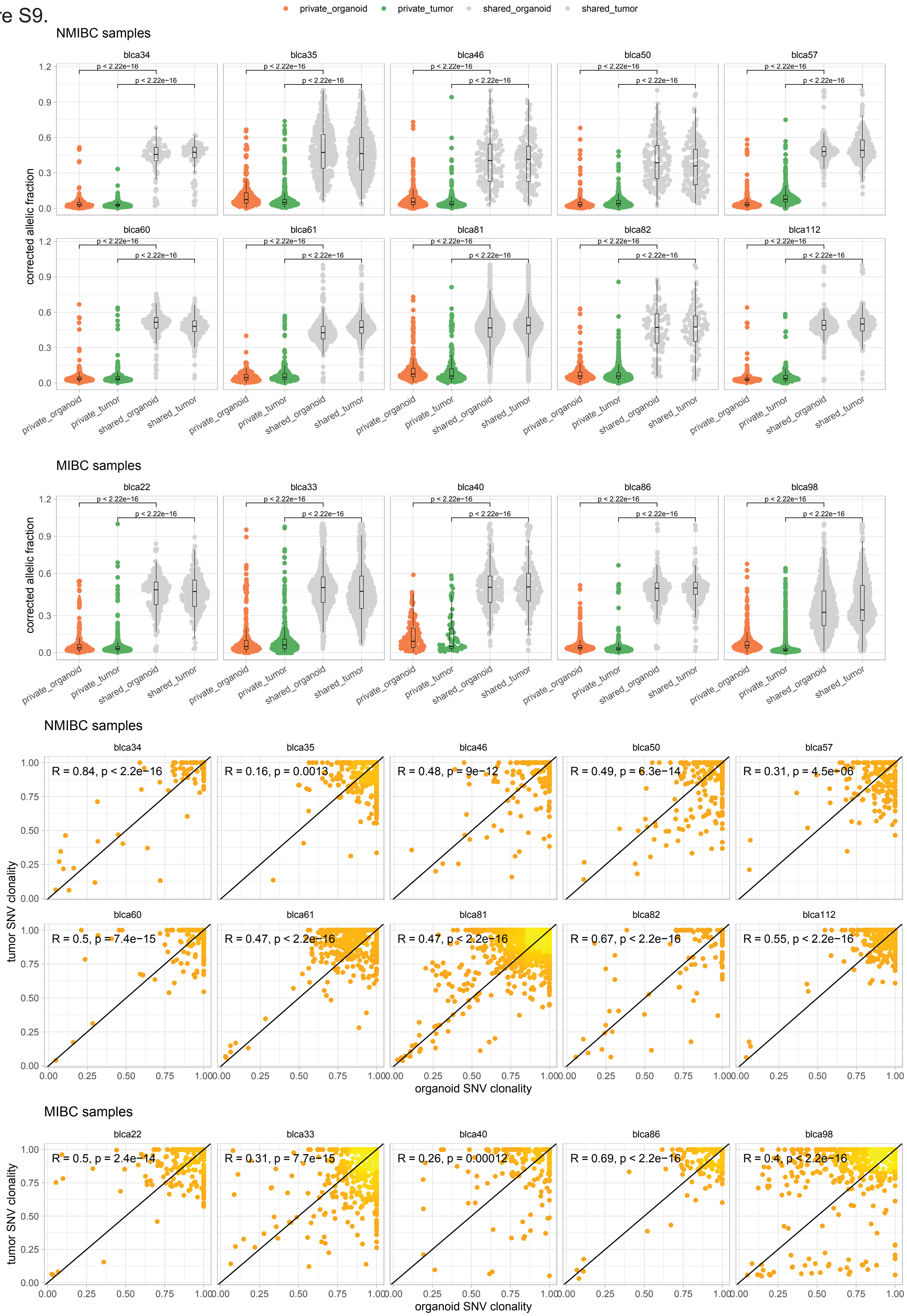

private_organoid
private_tumor
shared_organoid
shared_tumor
NMIBC samples
blca34
blca35
blca46
blca50
blca57
blca60
blca61
blca81
blca82
blca112
p < 2.22e−16
corrected allelic fraction
MIBC samples
blca22
blca33
blca40
blca86
blca98
NMIBC samples
R = 0.84, p < 2.2e−16
R = 0.16, p = 0.0013
R = 0.48, p = 9e−12
R = 0.49, p = 6.3e−14
R = 0.31, p = 4.5e−06
R = 0.5, p = 7.4e−15
R = 0.47, p < 2.2e−16
R = 0.47, p < 2.2e−16
R = 0.67, p < 2.2e−16
R = 0.55, p < 2.2e−16
tumor SNV clonality
organoid SNV clonality
MIBC samples
R = 0.5, p = 2.4e−14
R = 0.31, p = 7.7e−15
R = 0.26, p = 0.00012
R = 0.69, p < 2.2e−16
R = 0.4, p < 2.2e−16
tumor SNV clonality
organoid SNV clonality

Figure S10.

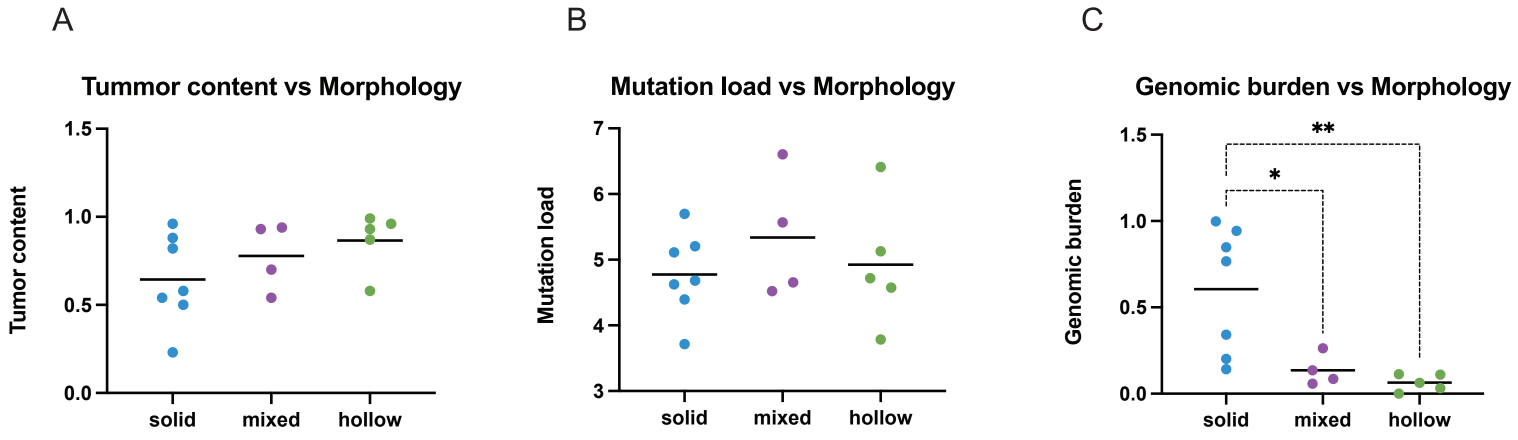
A
Tummor content vs Morphology
Tumor content
1.5
1.0
0.5
0.0
solid
mixed
hollow
B
Mutation load vs Morphology
Mutation load
7
6
5
4
3
solid
mixed
hollow
C
Genomic burden vs Morphology
Genomic burden
1.5
1.0
0.5
0.0
**
*
solid
mixed
hollow

Figure S11.

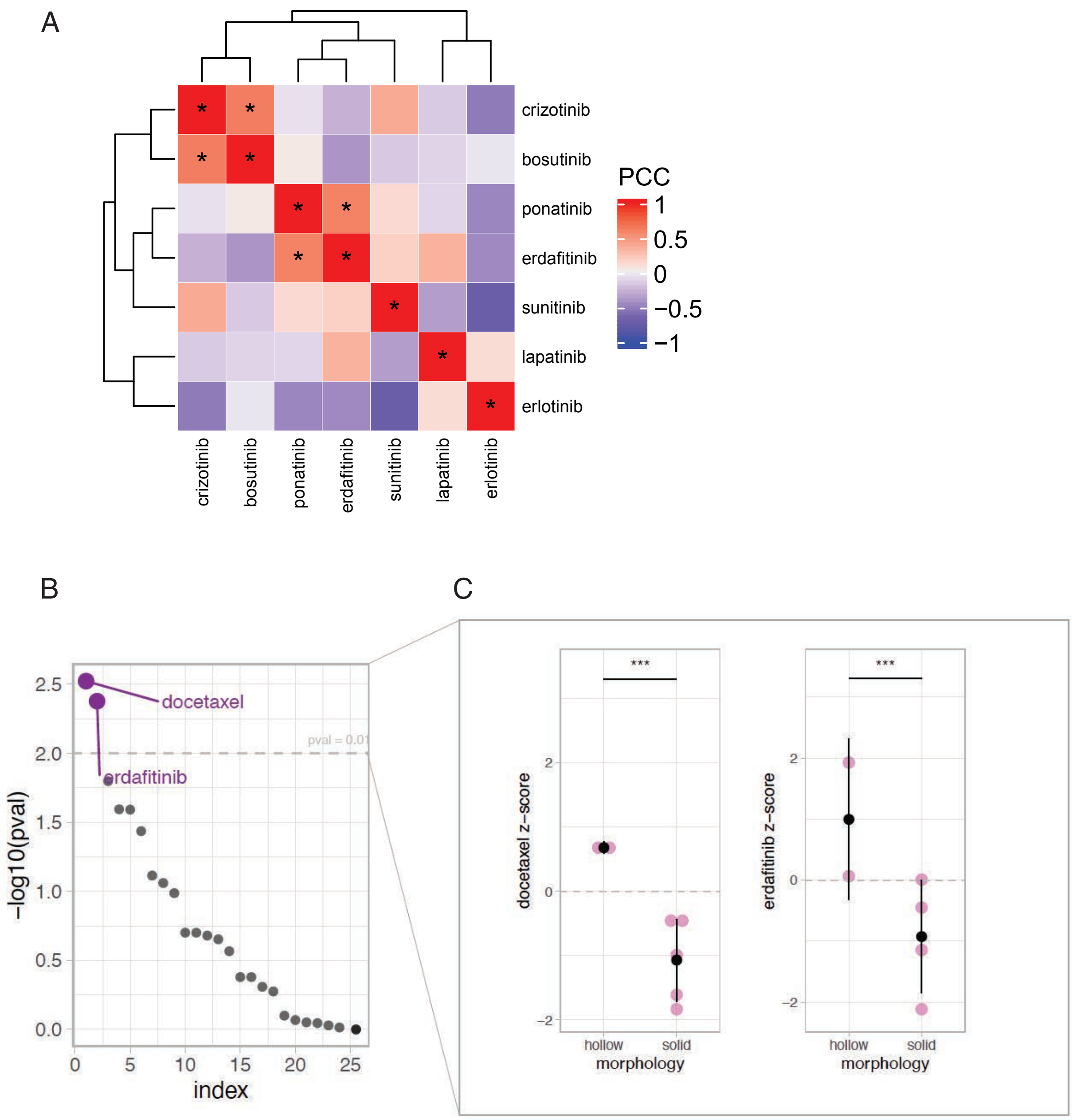

A
crizotinib
bosutinib
ponatinib
erdafitinib
sunitinib
lapatinib
erlotinib
PCC
1
0.5
0
−0.5
−1
B
docetaxel
erdafitinib
pval = 0.01
−log10(pval)
index
C
docetaxel z−score
erdafitinib z−score
***
hollow
solid
morphology

Figure S12.

A

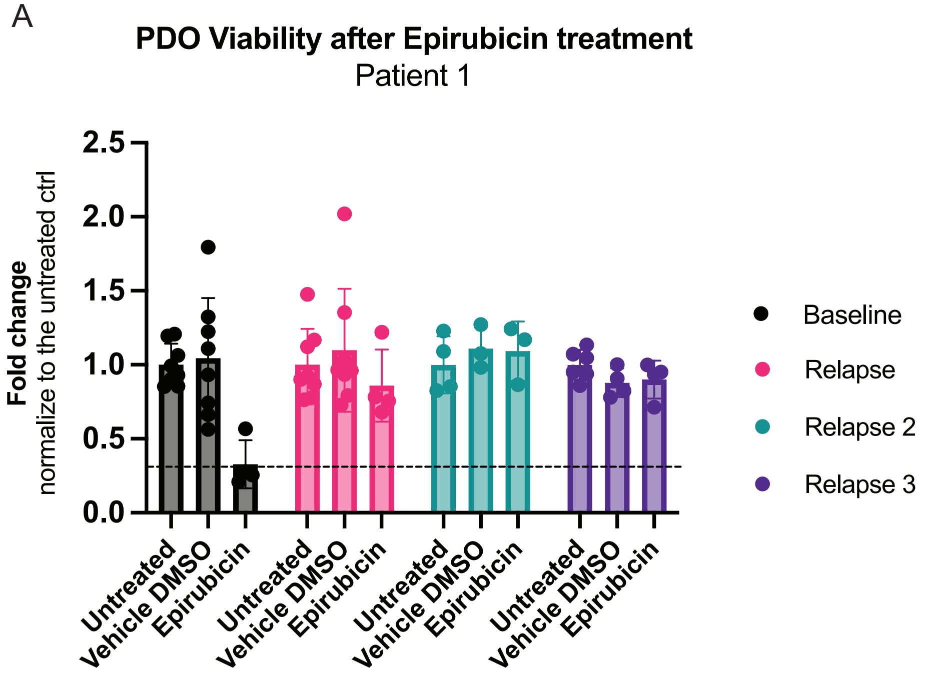


B

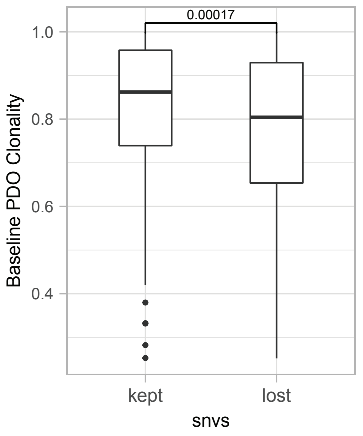

**Figure S1. Bladder cancer organoids characterized with a mixed morphology, Related to Figure 2**
Representative brightfield images of BLCa69 sample characterized with solid and hollow organoids, BLCa60 organoids with budding structures and BLCa33 organoids presenting a mix of solid and hollow features.

**Figure S2. Comparison of markers expression between organoids and parental tumor, Related to Figure 2**
(A-B) H&E and immunohistochemistry of parental tumor for indicated markers and brightfield images and whole mount immunofluorescence staining of organoids at passage 1 for indicated markers. BLCa22 (A) and BLCa33 (B) samples.

**Figure S3. Comparison of markers expression between organoids and parental tumor, Related to Figure 2**
(A-B) H&E and immunohistochemistry of parental tumor for indicated markers and brightfield images and whole mount immunofluorescence staining of organoids at passage 1 for indicated markers. BLCa35 (A) and BLCa40 (B) samples.

**Figure S4. Comparison of markers expression between organoids and parental tumor, Related to Figure 2**
(A-B) H&E and immunohistochemistry of parental tumor for indicated markers and brightfield images and whole mount immunofluorescence staining of organoids at passage 1 for indicated markers. BLCa34 (A) and BLCa47 (B) samples.

**Figure S5. Comparison of markers expression between organoids and parental tumor, Related to Figure 2**
(A-B) H&E and immunohistochemistry of parental tumor for indicated markers and brightfield images and whole mount immunofluorescence staining of organoids at passage 1 for indicated markers. BLCa60 (A) and BLCa98 (B) samples.

**Figure S6. Comparison of markers expression between organoids and parental tumor, Related to Figure 2**
(A-B) H&E and immunohistochemistry of parental tumor for indicated markers and brightfield images and whole mount immunofluorescence staining of organoids at passage 1 for indicated markers. BLCa82 (A) and BLCa46 (B) samples.

**Figure S7. Comparison of organoid markers expression with organoid morphology and primary tumor (PT) stage, Related to Figure 2**
(A-D) Image quantification of basal (A and C) and luminal (B and D) marker expression in PDOs grouped based on morphology (A-D) or PT stage (C-D). Each data point corresponds to one technical replicate (mean±SD), Ordinary two-way ANOVA test between PDO morphologies or PT stages, *p-value ≤ 0.05, ** p-value ≤ 0.01, *** p-value ≤ 0.001, **** p-value ≤ 0.0001.

**Figure S8. Analysis of Whole-Exome Sequencing Data, Related to Figure 3**
(A) Graph representing the clustering analysis performed using copy number similarity. Samples information are reported in the bottom of the graph.
(B) Proportion of single-nucleotide variants (SNV) per-class (shared or private fractions, mean± SD).

Abbreviations: tum, tumor; org, organoids.

**Figure S9. Analysis of Whole-Exome Sequencing Data, Related to Figure 3**

(A) Graphs representing the purity and ploidy corrected allelic fraction distribution for the shared and private point mutations in organoids and primary tumor.

(B) Graphs showing the clonality of sub-clonal point mutations in organoids and primary tumor.

Abbreviations: NMIBC, non-muscle invasive bladder cancer; MIBC, muscle invasive bladder cancer; SNV, single-nucleotide variant.

**Figure S10. Association of genomic features with organoid morphology, Related to Figure 4.**

(A-C) Analysis of tumor content (A), mutation load (B) and genomic burden (C) in solid, hollow and mixed organoids. Each point represents one sample. One-way ANOVA test (multiple comparison) between PDO morphologies, *p-value ≤ 0.05, ** p-value ≤ 0.01.

**Figure S11. Analysis of drug screening data, related to Figure 5**

(A) Correlation matrix showing the Pearson correlation coefficient (PCC) between the different tyrosine-kinases inhibitors tested on organoids. Correlation test, * p ≤ 0.1

(B) Association between drug response and organoid morphology (solid and hollow morphologies). Linear Mixed Model (see Material and methods, section "Drugs association analyses").

(C) Graphs representing the association between docetaxel (left) and erdafitinib (right) response with organoid morphology. Linear Mixed Model, ***p ≤ 0.001, (see Material and methods, section "Drugs association analyses").

**Figure S12. Related to Figure 6**

(A) Organoid longitudinal viability measured in samples derived from patient 1 at different time points (baseline, relapse, relapse 2 and relapse 3). The value of organoid viability at 48h after treatment was normalized to the value measured at time 0 (mean± SD). Each data point corresponds to technical replicates. One-way ANOVA with Dunnet's multiple comparison test between treatment and vehicle.

(B) Clonality of single nucleotide variants (SNVs) preserved and lost between organoids from baseline and relapse. Unpaired Wilcoxon test.
