## Supplemental tables for "Bladder cancer organoids as a functional system to model different disease stages and therapy response": Supplementary Information Tables.pdf

**Supplementary tables**

**Table S1. Cell viability measured in fresh and cryopreserved tissue and cryopreserved single cells.**
Abbreviations: N/A, not available.

**Table S2. Percentage of Ki67 positive cells in parental tumor (PT) and matched derived organoids**.
Abbreviations: PDO patient-derived organoids; N/A, not annotated,

**Table S3. Organoid morphologies quantification over 17 samples**

**Table S4. Sequencing statistics**
Abbreviations: tum, tumor; org, organoids.

**Table S5. Sample sequencing info**
Abbreviations: PDO patient-derived organoids; N/A, not annotated.

**Table S6. Non-muscle invasive sample mutation frequencies**
Abbreviations: amp, amplification; homo_del, homo-deletion; hemi_del, hemi-deletion; msk, Memorial Sloan Kettering; SNV, single nucleotide variants.

**Table S7. Muscle invasive sample mutations frequencies**
Abbreviations: amp, amplification; homo_del, homo-deletion; hemi_del, hemi-deletion; SNV, single nucleotide variants; tcga, The Cancer Genome Atlas.

**Table S8. SNV enrichment tables**
Binary tables showing frequently enriched terms across cohort for each sample.

**Table S9. Drug screening viability (z-scored e fold-change)**

**Table S10. Association analysis table – genomic**

**Table S11. Association analysis table - phenotypic**
