## Supplemental tables for "Bladder cancer organoids as a functional system to model different disease stages and therapy response": Table S1.pdf

| | Cell viability | | |
|---|---|---|---|
| **Sample code** | **Fresh tumor** | **Cryopreserved tumor** | **Cryopreserved cells** |
| BLCa86 | 88% | 99% | 90% |
| BLCa82 | 72% | N/A | N/A |
| BLCa40 | 90% | N/A | N/A |
| BLCa98 | 88% | N/A | 47% |
| BLCa112 | 91% | N/A | N/A |
| BLCa46 | 88% | N/A | 69% |
| BLCa81 | N/A | 99% | N/A |
| BLCa69 | N/A | 90% | N/A |
| BLCa50 | N/A | 82% | N/A |
| BLCa33 | 90% | 82% | 87% |
| BLCa22 | N/A | 90% | 88% |
| BLCa47 | N/A | 60% | N/A |
| BLCa61 | N/A | 79% | N/A |
| BLCa57 | N/A | 92% | N/A |
| BLCa35 | N/A | 78% | N/A |
| BLCa60 | N/A | N/A | 70% |
| BLCa34 | 95% | 92% | N/A |
