## Supplemental tables for "Bladder cancer organoids as a functional system to model different disease stages and therapy response": Table S2.pdf

| | Sample code | % Ki67 in PT | % Ki67 in PDO |
|---|---|---|---|
| PT > 20% Ki67+ nuclei | **BLCa33** | 35% | N/A |
| | **BLCa35** | 28% | 16% |
| | **BLCa40** | 29% | 25% |
| | **BLCa46** | 21% | 4% |
| | **BLCa50** | 48% | N/A |
| | **BLCa57** | 35% | N/A |
| | **BLCa69** | 29% | N/A |
| | **BLCa81** | 36% | N/A |
| | **BLCa82** | 22% | 2% |
| | **BLCa98** | 43% | 7% |
| PT ≤ 20% Ki67+ nuclei | **BLCa22** | 14% | 0% |
| | **BLCa34** | 20% | 1% |
| | **BLCa47** | 17% | 0% |
| | **BLCa60** | 13% | 1% |
| | **BLCa61** | 18% | N/A |
| | **BLCa86** | 18% | 4% |
| | **BLCa112** | 6% | 6% |
