## Supplemental tables for "Bladder cancer organoids as a functional system to model different disease stages and therapy response": Table S3.pdf

| **Model** | **Solid** | **Hollow** | **Mixed** | **Total organoids per picture** |
|---|---|---|---|---|
| BLCa22_1 | 2 | 2 | 0 | 4 |
| BLCa22_2 | 7 | 4 | 0 | 11 |
| BLCa22_3 | 1 | 1 | 1 | 3 |
| BLCa22_4 | 6 | 2 | 0 | 8 |
| BLCa22_5 | 1 | 2 | 0 | 3 |
| BLCa22_6 | 10 | 7 | 0 | 17 |
| BLCa22_7 | 6 | 1 | 0 | 7 |
| BLCa33_1 | 3 | 1 | 25 | 29 |
| BLCa33_2 | 4 | 11 | 22 | 37 |
| BLCa33_3 | 2 | 0 | 17 | 19 |
| BLCa33_4 | 2 | 2 | 26 | 30 |
| BLCa33_5 | 7 | 7 | 10 | 24 |
| BLCa33_6 | 1 | 3 | 16 | 20 |
| BLCa33_7 | 4 | 2 | 15 | 21 |
| BLCa34_1 | 1 | 7 | 0 | 8 |
| BLCa34_2 | 4 | 4 | 2 | 10 |
| BLCa34_3 | 0 | 6 | 1 | 7 |
| BLCa34_4 | 19 | 44 | 6 | 69 |
| BLCa34_5 | 0 | 15 | 3 | 18 |
| BLCa34_6 | 0 | 22 | 0 | 22 |
| BLCa35_1 | 2 | 0 | 1 | 3 |
| BLCa35_2 | 6 | 0 | 0 | 6 |
| BLca35_3 | 6 | 0 | 0 | 6 |
| BLCa35_4 | 3 | 1 | 0 | 4 |
| BLCa35_5 | 5 | 1 | 0 | 6 |
| BLCa35_6 | 3 | 0 | 0 | 3 |
| BLCA35_7 | 3 | 0 | 0 | 3 |
| BLCa40_1 | 7 | 0 | 6 | 13 |
| BLCa40_2 | 7 | 0 | 4 | 11 |
| BLCa40_3 | 6 | 0 | 5 | 11 |
| BLCa40_4 | 3 | 0 | 2 | 5 |
| BLCa40_5 | 5 | 0 | 7 | 12 |
| BLCa40_6 | 6 | 0 | 6 | 12 |
| BLCa40_7 | 2 | 1 | 3 | 6 |
| BLCa46_1 | 2 | 1 | 0 | 3 |
| BCLa46_2 | 1 | 2 | 0 | 3 |
| BLca46_3 | 1 | 0 | 0 | 1 |
| BLCa46_4 | 0 | 0 | 1 | 1 |
| BLCa46_5 | 0 | 1 | 0 | 1 |
| BLCa46_6 | 1 | 1 | 0 | 2 |
| BLCa46_7 | 1 | 0 | 0 | 1 |

| | | | | |
|---|---|---|---|---|
| BLCa47_1 | 1 | 0 | 0 | 1 |
| BLCa47_2 | 2 | 0 | 0 | 2 |
| BLCa47_3 | 2 | 0 | 0 | 2 |
| BLCa47_4 | 1 | 0 | 0 | 1 |
| BLCa47_5 | 1 | 0 | 0 | 1 |
| BLCa47_6 | 3 | 0 | 0 | 3 |
| BLCa50_1 | 3 | 0 | 1 | 4 |
| BLCa50_2 | 1 | 0 | 1 | 2 |
| BLCa50_3 | 2 | 1 | 0 | 3 |
| BLCa50_4 | 1 | 1 | 0 | 2 |
| BLCa50_5 | 1 | 0 | 1 | 2 |
| BLCa50_6 | 2 | 1 | 0 | 3 |
| BLCa50_7 | 0 | 1 | 1 | 2 |
| BLCa57_1 | 0 | 3 | 0 | 3 |
| BLCa57_2 | 0 | 7 | 0 | 7 |
| BLCa57_3 | 1 | 3 | 0 | 4 |
| BLCa57_4 | 1 | 2 | 0 | 3 |
| BLCa57_5 | 1 | 15 | 0 | 16 |
| BLCa57_6 | 0 | 1 | 1 | 2 |
| BLCa60_1 | 1 | 3 | 1 | 5 |
| BLCa60_2 | 0 | 0 | 2 | 2 |
| BLCa60_3 | 0 | 1 | 1 | 2 |
| BLCa60_4 | 0 | 1 | 1 | 2 |
| BLca61_1 | 0 | 7 | 0 | 7 |
| BLca61_2 | 0 | 6 | 0 | 6 |
| BLCa61_3 | 1 | 6 | 0 | 7 |
| BLCa61_4 | 1 | 6 | 0 | 7 |
| BLCa61_5 | 0 | 20 | 1 | 21 |
| BLCa61_6 | 0 | 15 | 1 | 16 |
| BLCa61_7 | 1 | 9 | 0 | 10 |
| BLCa69_1 | 5 | 5 | 1 | 11 |
| BLCa69_2 | 4 | 8 | 3 | 15 |
| BLCa69_3 | 4 | 6 | 4 | 14 |
| BLCa69_4 | 3 | 5 | 2 | 10 |
| BLCa69_5 | 2 | 2 | 2 | 6 |
| BLCa81_1 | 4 | 5 | 0 | 9 |
| BLCa81_2 | 0 | 5 | 2 | 7 |
| BLCa81_3 | 2 | 9 | 2 | 13 |
| BLCa81_4 | 3 | 0 | 1 | 4 |
| BLCa81_5 | 3 | 9 | 0 | 12 |
| BLCa82_1 | 4 | 1 | 0 | 5 |
| BLCa82_2 | 1 | 0 | 0 | 1 |

| | | | | |
|---|---|---|---|---|
| BLCa82_3 | 1 | 0 | 0 | 1 |
| BLCa82_4 | 4 | 0 | 0 | 4 |
| BLCa82_5 | 1 | 0 | 0 | 1 |
| BLCa82_6 | 3 | 0 | 3 | 6 |
| BLCa82_7 | 3 | 0 | 2 | 5 |
| BLCa86_1 | 0 | 11 | 2 | 13 |
| BLCa86_2 | 0 | 5 | 0 | 5 |
| BLCa86_3 | 0 | 14 | 1 | 15 |
| BLCa86_4 | 0 | 6 | 2 | 8 |
| BLCa86_5 | 0 | 4 | 2 | 6 |
| BLCa86_6 | 0 | 9 | 4 | 13 |
| BLCa86_7 | 0 | 13 | 0 | 13 |
| BLCa98_1 | 3 | 0 | 0 | 3 |
| BLCa98_2 | 2 | 0 | 0 | 2 |
| BLCa98_3 | 1 | 0 | 0 | 1 |
| BLCa98_4 | 1 | 0 | 0 | 1 |
| BLCa98_5 | 5 | 0 | 0 | 5 |
| BLCa112_1 | 7 | 13 | 8 | 28 |
| BLCa112_2 | 9 | 6 | 10 | 25 |
| BLCa112_3 | 7 | 9 | 5 | 21 |
| BLCa112_4 | 9 | 4 | 5 | 18 |
| **sum** | **268** | **409** | **252** | **929** |
| **%** | **29%** | **44%** | **27%** | **100%** |
