## Supplemental tables for "Bladder cancer organoids as a functional system to model different disease stages and therapy response": Table S4.pdf

| sample_id | pipeline_id | n_total_reads | n_base_pair_covered | mean_coverage | fraction_of_base_pair_>10x |
|---|---|---|---|---|---|
| blca40_tum | b036 | 79324292 | 35936220 | 87.515631 | 0.972011 |
| blca40_org | b038 | 80315048 | 35936220 | 81.620871 | 0.972212 |
| blca22_tum | s1 | 231937510 | 63373586 | 79.326331 | 0.961547 |
| blca34_tum | s10 | 373793700 | 63373586 | 116.614868 | 0.971433 |
| blca34_org | s11 | 222783958 | 63373586 | 95.338855 | 0.967821 |
| blca35_tum | s13 | 487095876 | 63373586 | 110.648218 | 0.971491 |
| blca35_org | s14 | 284359042 | 63373586 | 102.338638 | 0.967673 |
| blca46_tum | s16 | 256808320 | 63373586 | 84.802871 | 0.963184 |
| blca46_org | s17 | 154421874 | 63373586 | 57.813329 | 0.947061 |
| blca48_tum | s19 | 289118700 | 63373586 | 88.242055 | 0.966725 |
| blca22_org | s2 | 221432548 | 63373586 | 91.23781 | 0.964412 |
| blca48_org | s20 | 293539636 | 63373586 | 115.558042 | 0.971662 |
| blca50_tum | s22 | 199244816 | 63373586 | 69.452154 | 0.956276 |
| blca57_tum | s25 | 228950730 | 63373586 | 44.54912 | 0.954591 |
| blca57_org | s26 | 317153256 | 63373586 | 123.682068 | 0.970404 |
| blca60_tum | s28 | 241866238 | 63373586 | 80.851088 | 0.963963 |
| blca60_org | s29 | 275138718 | 63373586 | 100.520171 | 0.969107 |
| blca61_tum | s31 | 250679080 | 63373586 | 87.671322 | 0.966205 |
| blca61_org | s32 | 299306012 | 63373586 | 114.273704 | 0.970335 |
| blca27_tum | s4 | 283135402 | 63373586 | 101.256708 | 0.970239 |
| blca27_org | s5 | 151799180 | 63373586 | 67.988641 | 0.958774 |
| blca33_tum | s7 | 314207270 | 63373586 | 109.759221 | 0.97165 |
| blca33_org | s8 | 257918676 | 63373586 | 96.232886 | 0.967514 |
| blca100_org | blca100_org | 546538412 | 63373586 | 123.074871 | 0.974151 |
| blca100_tum | blca100_tum | 233895428 | 63373586 | 87.870124 | 0.966368 |
| blca47_org | blca47_org | 358052774 | 63373586 | 75.077445 | 0.960341 |
| blca47_tum | blca47_tum | 213995566 | 63373586 | 78.26234 | 0.962858 |
| blca69_org | blca69_org | 458774310 | 63373586 | 105.8455 | 0.97122 |
| blca81_org | blca81_org | 382530016 | 63373586 | 85.182718 | 0.966888 |
| blca81_tum | blca81_tum | 336610236 | 63373586 | 134.426989 | 0.973036 |
| blca82_org | blca82_org | 463777140 | 63373586 | 120.622539 | 0.971625 |
| blca82_tum | blca82_tum | 347695992 | 63373586 | 124.635257 | 0.972078 |
| blca85_org | blca85_org | 414390762 | 63373586 | 86.828096 | 0.969682 |
| blca85_tum | blca85_tum | 267770342 | 63373586 | 97.451701 | 0.968171 |
| blca86_org | blca86_org | 382075796 | 63373586 | 93.823286 | 0.969996 |
| blca86_tum | blca86_tum | 378310816 | 63373586 | 119.549696 | 0.970317 |
| blca92_org | blca92_org | 385874888 | 63373586 | 75.822649 | 0.965794 |
| blca92_tum | blca92_tum | 632743986 | 63373586 | 215.710548 | 0.975828 |
| blca112_org | blca112_org | 169592752 | 63373586 | 95.508268 | 0.966868 |
| blca112_tum | blca112_tum | 152048076 | 63373586 | 78.049185 | 0.963202 |
| blca50b_org | blca50b_org | 203910138 | 63373586 | 110.280575 | 0.966195 |
| blca98b_org | blca98b_org | 401313720 | 63373586 | 91.159652 | 0.967678 |

| | | | | | |
|---|---|---|---|---|---|
| blca98b_tum | blca98b_tum | 617236362 | 63373586 | 207.112438 | 0.974467 |
