## Supplemental tables for "Bladder cancer organoids as a functional system to model different disease stages and therapy response": Table S5.pdf

| Case id | Tissue type | Tumor subtype | sex | Main PDO morphology | purity | ploidy | Ploidy status | Count deleteroius snvs | Count deleteroius indels | Count deleteroius insertions | Count deleterious deletions | Genomic burden | Spia result | Used in can analyses | Mutation load |
|---|---|---|---|---|---|---|---|---|---|---|---|---|---|---|---|
| blca100 | tumor | NMIBC | m | N/A | 0.49 | 2.22 | diploid | 75 | 16 | 4 | 12 | 0.377946346 | Similar | no | 4.33073334 |
| blca100 | organoid | NMIBC | m | N/A | N/A | 2.27 | N/A | 53 | 50 | 18 | 32 | N/A | Similar | no | 3.98898405 |
| blca112 | tumor | NMIBC | m | N/A | 0.9 | 1.97 | diploid | 107 | 14 | 2 | 12 | 0.084806813 | Similar | yes | 4.68213123 |
| blca112 | organoid | NMIBC | m | Hollow | 0.94 | 1.96 | diploid | 104 | 14 | 2 | 12 | 0.085271448 | Similar | yes | 4.65396035 |
| blca22 | tumor | MIBC | m | N/A | 0.92 | 2.03 | diploid | 118 | 15 | 6 | 9 | 0.19838648 | Similar | yes | 4.77912349 |
| blca22 | organoid | MIBC | m | Solid | 0.96 | 2.02 | diploid | 107 | 22 | 8 | 14 | 0.202446551 | Similar | yes | 4.68213123 |
| blca27 | tumor | MIBC | m | N/A | N/A | 2.06 | N/A | 1 | 174 | 160 | 14 | N/A | Similar | no | 0.69314718 |
| blca27 | organoid | MIBC | m | N/A | N/A | 2.06 | N/A | 6 | 6 | 1 | 5 | N/A | Similar | no | 1.94591015 |
| blca33 | tumor | MIBC | m | N/A | 0.38 | 2.03 | diploid | 250 | 127 | 106 | 21 | 0.284040037 | Similar | yes | 5.52545294 |
| blca33 | organoid | MIBC | m | Mixed | 0.7 | 2 | diploid | 261 | 24 | 2 | 22 | 0.26335074 | Similar | yes | 5.5683445 |
| blca34 | tumor | NMIBC | m | N/A | 0.92 | 2 | diploid | 43 | 21 | 9 | 12 | 0.00235438 | Similar | yes | 3.78418963 |
| blca34 | organoid | NMIBC | m | Hollow | 0.99 | 1.99 | diploid | 43 | 12 | 3 | 9 | 0.001070169 | Similar | yes | 3.78418963 |
| blca35 | tumor | NMIBC | m | N/A | 0.47 | 2.94 | polyploid | 204 | 40 | 8 | 32 | 0.944182885 | Similar | yes | 5.32300998 |
| blca35 | organoid | NMIBC | m | Solid | 0.23 | 2.52 | polyploid | 181 | 35 | 9 | 26 | 0.94343034 | Similar | yes | 5.20400669 |
| blca40 | tumor | MIBC | m | N/A | 0.85 | 2.04 | diploid | 163 | 16 | 4 | 12 | 0.245402814 | Similar | yes | 5.09986643 |
| blca40 | organoid | MIBC | m | Solid | 0.54 | 2.03 | diploid | 165 | 16 | 4 | 12 | 0.141509338 | Similar | yes | 5.11198779 |
| blca46 | tumor | NMIBC | f | N/A | 0.8 | 3.59 | polyploid | 91 | 13 | 3 | 10 | 0.99784745 | Similar | yes | 4.52178858 |
| blca46 | organoid | NMIBC | f | Solid | 0.82 | 3.63 | polyploid | 101 | 28 | 17 | 11 | 0.997746342 | Similar | yes | 4.62497281 |
| blca47 | tumor | MIBC | f | N/A | N/A | 2.03 | N/A | 112 | 36 | 20 | 16 | N/A | Similar | no | 4.72738782 |
| blca47 | organoid | MIBC | f | Solid | N/A | 2.03 | N/A | 89 | 64 | 31 | 33 | N/A | Similar | no | 4.49980967 |
| blca48 | tumor | MIBC | m | N/A | N/A | 1.98 | N/A | 27 | 15 | 5 | 10 | N/A | Similar | no | 3.33220451 |
| blca48 | organoid | MIBC | m | N/A | N/A | 2 | N/A | 1 | 14 | 0 | 14 | N/A | Similar | no | 0.69314718 |
| blca50 | tumor | NMIBC | m | N/A | 0.8 | 3.38 | polyploid | 103 | 12 | 4 | 8 | 0.85502 | Similar | yes | 4.6443909 |
| blca50b | organoid | NMIBC | m | Solid | 0.88 | 3.35 | polyploid | 80 | 12 | 2 | 10 | 0.848643646 | Similar | yes | 4.39444915 |
| blca57 | tumor | NMIBC | m | N/A | 0.75 | 1.92 | diploid | 113 | 16 | 4 | 12 | 0.063373133 | Similar | yes | 4.73619845 |
| blca57 | organoid | NMIBC | m | Hollow | 0.87 | 1.96 | diploid | 96 | 26 | 8 | 18 | 0.063291519 | Similar | yes | 4.57471098 |
| blca60 | tumor | NMIBC | m | N/A | 0.93 | 2.07 | diploid | 92 | 18 | 4 | 14 | 0.066568542 | Similar | yes | 4.53259949 |
| blca60 | organoid | NMIBC | m | Mixed | 0.93 | 2.05 | diploid | 91 | 17 | 5 | 12 | 0.057919751 | Similar | yes | 4.52178858 |
