## Supplemental tables for "Bladder cancer organoids as a functional system to model different disease stages and therapy response": Table S6.pdf

| | snv_inhouse | homo_del_inhouse | hemi_del_inhouse | gain_inhouse | amp_inhouse | snv_msk | homo_del_msk | amp_msk |
|---|---|---|---|---|---|---|---|---|
| PIK3CA | 0.6 | 0 | 0 | 0.4 | 0 | 0.30476191 | 0 | 0 |
| FGFR3 | 0.5 | 0 | 0 | 0.1 | 0.1 | 0.46666667 | 0 | 0.01904762 |
| HRAS | 0 | 0 | 0.2 | 0.3 | 0 | 0.03809524 | 0 | 0 |
| EGFR | 0 | 0 | 0 | 0.3 | 0.1 | 0.02857143 | 0 | 0 |
| PTEN | 0 | 0 | 0 | 0.2 | 0 | 0.01904762 | 0 | 0 |
| TSC1 | 0 | 0 | 0.3 | 0.3 | 0 | 0.11428571 | 0 | 0 |
| NF1 | 0 | 0 | 0 | 0.1 | 0.2 | 0.14285714 | 0 | 0 |
| ARID1A | 0.3 | 0 | 0 | 0.2 | 0 | 0.34285714 | 0 | 0 |
| EP300 | 0 | 0 | 0.1 | 0.3 | 0 | 0.17142857 | 0 | 0 |
| KMT2D | 0.2 | 0 | 0 | 0.2 | 0.1 | 0 | 0 | 0 |
| YAP1 | 0 | 0 | 0 | 0.3 | 0 | 0 | 0.00952381 | 0 |
| KMT2A | 0 | 0 | 0.1 | 0.2 | 0 | 0 | 0 | 0 |
| KMT2C | 0.1 | 0 | 0 | 0.3 | 0.1 | 0 | 0 | 0 |
| CREBBP | 0.1 | 0 | 0 | 0.3 | 0 | 0.22857143 | 0 | 0 |
| CDKN2A | 0 | 0.1 | 0.1 | 0.3 | 0.1 | 0.04761905 | 0.16190476 | 0 |
| CCND3 | 0 | 0 | 0 | 0.3 | 0.1 | 0.01904762 | 0 | 0.00952381 |
| E2F3 | 0 | 0 | 0 | 0.2 | 0 | 0.01904762 | 0 | 0.02857143 |
| CCND1 | 0 | 0 | 0 | 0.2 | 0.1 | 0.00952381 | 0 | 0.06666667 |
| CCNE1 | 0 | 0 | 0 | 0.2 | 0 | 0 | 0 | 0 |
| TP53 | 0.1 | 0 | 0.2 | 0.2 | 0.1 | 0.25714286 | 0 | 0 |
| RB1 | 0.2 | 0 | 0 | 0.1 | 0.2 | 0.03809524 | 0.01904762 | 0 |
| MDM2 | 0 | 0 | 0 | 0.2 | 0.1 | 0.00952381 | 0 | 0.06666667 |
| CDKN1A | 0.2 | 0 | 0 | 0.4 | 0 | 0.16190476 | 0 | 0 |
