## Supplemental tables for "Bladder cancer organoids as a functional system to model different disease stages and therapy response": Table S7.pdf

| | snv_inhouse | homo_del_inhouse | hemi_del_inhouse | gain_inhouse | amp_inhouse | snv_tcga | homo_del_tcga | hemi_del_tcga | gain_tcga | amp_tcga |
|---|---|---|---|---|---|---|---|---|---|---|
| PIK3CA | 0 | 0 | 0 | 0.4 | 0 | 0.03950617 | 0 | 0.00396825 | 0.48015873 | 0.12698413 |
| FGFR3 | 0 | 0 | 0 | 0.2 | 0.2 | 0.02469136 | 0 | 0.06374502 | 0.33864542 | 0.03187251 |
| HRAS | 0 | 0 | 0.4 | 0.2 | 0 | 0 | 0.0078125 | 0.19140625 | 0.2265625 | 0.00390625 |
| EGFR | 0 | 0 | 0 | 0.2 | 0 | 0.00246914 | 0 | 0.01219512 | 0.45528455 | 0.05284553 |
| PTEN | 0 | 0 | 0 | 0 | 0 | 0.00493827 | 0.01181102 | 0.08661417 | 0.31889764 | 0.00787402 |
| TSC1 | 0 | 0 | 0.4 | 0.4 | 0 | 0.00740741 | 0 | 0.1627907 | 0.34496124 | 0.02325581 |
| NF1 | 0 | 0 | 0 | 0.2 | 0 | 0.00493827 | 0 | 0.0122449 | 0.47346939 | 0.08163265 |
| ARID1A | 0 | 0 | 0 | 0.2 | 0 | 0.04444444 | 0 | 0.03937008 | 0.42913386 | 0.02755906 |
| EP300 | 0.2 | 0 | 0.2 | 0 | 0 | 0.02222222 | 0 | 0.10116732 | 0.33463035 | 0.01167315 |
| KMT2D | 0 | 0 | 0 | 0.2 | 0 | 0.03950617 | 0 | 0.03137255 | 0.48235294 | 0.01568627 |
| YAP1 | 0 | 0 | 0.2 | 0.4 | 0 | 0.00493827 | 0 | 0.06719368 | 0.33596838 | 0.03162055 |
| KMT2A | 0 | 0 | 0.4 | 0.2 | 0 | 0.01975309 | 0 | 0.09448819 | 0.33858268 | 0.01574803 |
| KMT2C | 0 | 0 | 0 | 0.2 | 0 | 0.03703704 | 0 | 0.02745098 | 0.44705882 | 0.02745098 |
| CREBBP | 0 | 0 | 0.2 | 0.2 | 0 | 0.00987654 | 0.00772201 | 0.1003861 | 0.29343629 | 0.01544402 |
| CDKN2A | 0 | 0.4 | 0.2 | 0 | 0.2 | 0.00246914 | 0.22222222 | 0.14814815 | 0.25514403 | 0.02880658 |
| CCND3 | 0 | 0 | 0 | 0 | 0 | 0 | 0 | 0.01937984 | 0.41860465 | 0.03488372 |
| E2F3 | 0 | 0 | 0 | 0 | 0 | 0 | 0 | 0.01769912 | 0.40265487 | 0.06637168 |
| CCND1 | 0 | 0 | 0 | 0.4 | 0.2 | 0.00246914 | 0 | 0.02155172 | 0.40517241 | 0.06465517 |
| CCNE1 | 0 | 0 | 0 | 0.4 | 0 | 0 | 0 | 0.01293103 | 0.40517241 | 0.09482759 |
| TP53 | 0.4 | 0 | 0 | 0 | 0 | 0.0691358 | 0.00387597 | 0.18217054 | 0.29069767 | 0.00775194 |
| RB1 | 0.2 | 0.2 | 0 | 0.2 | 0 | 0.01975309 | 0.01219512 | 0.13821138 | 0.32113821 | 0.0203252 |
| MDM2 | 0 | 0 | 0 | 0.2 | 0 | 0 | 0 | 0.01639344 | 0.50409836 | 0.03278689 |
| CDKN1A | 0 | 0 | 0 | 0 | 0 | 0.00246914 | 0 | 0.02 | 0.424 | 0.036 |
