## Supplemental tables for "Bladder cancer organoids as a functional system to model different disease stages and therapy response": Table S9.pdf

| drug | raw_count | model | z_score | fold_change | viability reduction (1- fold change) |
|---|---|---|---|---|---|
| untreated | 56615 | BLCa48 | 0.456866816 | 1.10598493 | |
| untreated | 24601 | BLCa48 | -2.239029148 | 0.48058527 | |
| untreated | 27805 | BLCa48 | -1.969220606 | 0.54317603 | |
| untreated | 50322 | BLCa48 | -0.073066129 | 0.98304996 | |
| untreated | 47754 | BLCa48 | -0.289317171 | 0.93288359 | |
| untreated | 67199 | BLCa48 | 1.348144472 | 1.31274541 | |
| dmso | 38877 | BLCa48 | -1.036848515 | 0.75946968 | 0.24053032 |
| dmso | 42506 | BLCa48 | -0.731250763 | 0.8303629 | 0.1696371 |
| dmso | 62711 | BLCa48 | 0.970210409 | 1.22507147 | -0.2250715 |
| dmso | 55393 | BLCa48 | 0.353962309 | 1.08211293 | -0.0821129 |
| dmso | 41300 | BLCa48 | -0.832807911 | 0.80680346 | 0.19319654 |
| dmso | 66351 | BLCa48 | 1.276734471 | 1.29617957 | -0.2961796 |
| h2o | 42642 | BLCa48 | -0.450680575 | 0.91581511 | 0.08418489 |
| h2o | 48000 | BLCa48 | 0.165357621 | 1.03088798 | -0.030888 |
| h2o | 58593 | BLCa48 | 1.383292039 | 1.25839207 | -0.2583921 |
| h2o | 48672 | BLCa48 | 0.24262109 | 1.04532041 | -0.0453204 |
| h2o | 34902 | BLCa48 | -1.340590175 | 0.74958442 | 0.25041558 |
| bosutinib | 44920 | BLCa48 | -0.527968047 | 0.87752085 | 0.12247915 |
| bosutinib | 45402 | BLCa48 | -0.487378872 | 0.88693682 | 0.11306318 |
| bosutinib | 52246 | BLCa48 | 0.088953732 | 1.02063568 | -0.0206357 |
| cisplatin_6_um | 64252 | BLCa48 | 2.033937831 | 1.37992947 | -0.3799295 |
| cisplatin_6_um | 55526 | BLCa48 | 1.030662486 | 1.19252263 | -0.1925226 |
| cisplatin_6_um | 48425 | BLCa48 | 0.214222166 | 1.04001564 | -0.0400156 |
| epirubicin | 9170 | BLCa48 | -3.538472224 | 0.17913772 | 0.82086228 |
| epirubicin | 12475 | BLCa48 | -3.260158481 | 0.24370153 | 0.75629847 |
| epirubicin | 10932 | BLCa48 | -3.390094367 | 0.21355873 | 0.78644127 |
| everolimus | 30392 | BLCa48 | -1.751369577 | 0.59371358 | 0.40628642 |
| everolimus | 32135 | BLCa48 | -1.604591709 | 0.62776342 | 0.37223658 |
| everolimus | 37709 | BLCa48 | -1.135205687 | 0.73665258 | 0.26334742 |
| erlotinib | 22628 | BLCa48 | -2.405175295 | 0.44204234 | 0.55795766 |
| erlotinib | 19652 | BLCa48 | -2.655783978 | 0.38390561 | 0.61609439 |
| erlotinib | 32175 | BLCa48 | -1.601223312 | 0.62854482 | 0.37145518 |
| docetaxel | 56796 | BLCa48 | 0.472108809 | 1.1095208 | -0.1095208 |
| docetaxel | 35760 | BLCa48 | -1.299330796 | 0.69857849 | 0.30142151 |
| docetaxel | 67438 | BLCa48 | 1.368270639 | 1.31741432 | -0.3174143 |
| lapatinib | 24957 | BLCa48 | -2.209050421 | 0.4875398 | 0.5124602 |
| lapatinib | 40995 | BLCa48 | -0.858491932 | 0.80084522 | 0.19915478 |
| lapatinib | 45015 | BLCa48 | -0.519968106 | 0.8793767 | 0.1206233 |
| crizotinib | 45918 | BLCa48 | -0.44392656 | 0.89701698 | 0.10298302 |
| crizotinib | 46351 | BLCa48 | -0.40746367 | 0.90547571 | 0.09452429 |
| crizotinib | 55442 | BLCa48 | 0.358088595 | 1.08307015 | -0.0830702 |
| gemcitabine | 32463 | BLCa48 | -1.621015177 | 0.69720243 | 0.30279757 |
| gemcitabine | 35371 | BLCa48 | -1.286666712 | 0.75965706 | 0.24034294 |
| gemcitabine | 28729 | BLCa48 | -2.050333323 | 0.61700793 | 0.38299207 |
| cisplatin+gemcitabine | 21412 | BLCa48 | -2.891608329 | 0.45986195 | 0.54013805 |
| cisplatin+gemcitabine | 29818 | BLCa48 | -1.925125112 | 0.6403962 | 0.3596038 |
| cisplatin+gemcitabine | 31497 | BLCa48 | -1.732081414 | 0.67645581 | 0.32354419 |
| untreated | 19908 | BLCa92 | -0.118171079 | 0.97749354 | |
| untreated | 21933 | BLCa92 | 0.403882776 | 1.07692213 | |
| untreated | 18931 | BLCa92 | -0.370045952 | 0.92952231 | |
| untreated | 18411 | BLCa92 | -0.504104226 | 0.90399003 | |
| untreated | 22901 | BLCa92 | 0.653437409 | 1.12445145 | |
| untreated | 18667 | BLCa92 | -0.438106306 | 0.91655977 | |
| untreated | 19409 | BLCa92 | -0.246815462 | 0.95299237 | |
| dmso | 17744 | BLCa92 | -0.676059743 | 0.87123997 | 0.12876003 |
| dmso | 17751 | BLCa92 | -0.674255112 | 0.87158368 | 0.12841632 |
| dmso | 21776 | BLCa92 | 0.363407489 | 1.06921335 | -0.0692133 |
| dmso | 24945 | BLCa92 | 1.180389547 | 1.22481296 | -0.224813 |
| dmso | 23497 | BLCa92 | 0.807088815 | 1.15371538 | -0.1537154 |
| dmso | 24501 | BLCa92 | 1.065924406 | 1.20301232 | -0.2030123 |
| dmso | 14164 | BLCa92 | -1.598999398 | 0.69546004 | 0.30453996 |
| dmso | 18553 | BLCa92 | -0.467496005 | 0.91096231 | 0.08903769 |
| h2o | 22690 | BLCa92 | 1.028375796 | 1.17834277 | -0.1783428 |
| h2o | 18535 | BLCa92 | -0.21586523 | 0.96256427 | 0.03743573 |
| h2o | 12507 | BLCa92 | -2.020987952 | 0.64951666 | 0.35048334 |

| | | | | | |
|---|---|---|---|---|---|
| h2o | 21811 | BLCa92 | 0.765153687 | 1.13269432 | -0.1326943 |
| h2o | 20670 | BLCa92 | 0.423474022 | 1.07343962 | -0.0734396 |
| h2o | 18780 | BLCa92 | -0.142498431 | 0.97528767 | 0.02471233 |
| h2o | 19798 | BLCa92 | 0.162348107 | 1.0281547 | -0.0281547 |
| cisplatin_6_um | 19918 | BLCa92 | 0.198282866 | 1.03438657 | -0.0343866 |
| cisplatin_6_um | 18551 | BLCa92 | -0.211073929 | 0.96339518 | 0.03660482 |
| cisplatin_6_um | 16488 | BLCa92 | -0.828852326 | 0.85625895 | 0.14374105 |
| cisplatin_6_um | 16724 | BLCa92 | -0.758180633 | 0.86851496 | 0.13148504 |
| cisplatin_6_um | 17118 | BLCa92 | -0.640194841 | 0.88897627 | 0.11102373 |
| bosutinib | 16055 | BLCa92 | -1.111491328 | 0.78830916 | 0.21169084 |
| bosutinib | 15924 | BLCa92 | -1.145263701 | 0.78187699 | 0.21812301 |
| bosutinib | 16565 | BLCa92 | -0.980011098 | 0.81335044 | 0.18664956 |
| bosutinib | 26976 | BLCa92 | 1.703990229 | 1.32453615 | -0.3245362 |
| bosutinib | 21830 | BLCa92 | 0.377328925 | 1.07186478 | -0.0718648 |
| cisplatin_14_um | 18941 | BLCa92 | -0.094285963 | 0.98364876 | 0.01635124 |
| cisplatin_14_um | 17889 | BLCa92 | -0.409314016 | 0.92901603 | 0.07098397 |
| cisplatin_14_um | 21643 | BLCa92 | 0.714845025 | 1.1239697 | -0.1239697 |
| cisplatin_14_um | 15182 | BLCa92 | -1.219942285 | 0.78843543 | 0.21156457 |
| cisplatin_14_um | 22630 | BLCa92 | 1.010408417 | 1.17522683 | -0.1752268 |
| crizotinib | 11179 | BLCa92 | -2.368545451 | 0.54889493 | 0.45110507 |
| crizotinib | 20205 | BLCa92 | -0.041603181 | 0.9920764 | 0.0079236 |
| crizotinib | 10827 | BLCa92 | -2.45929259 | 0.53161154 | 0.46838846 |
| crizotinib | 13081 | BLCa92 | -1.878201534 | 0.64228416 | 0.35771584 |
| crizotinib | 17512 | BLCa92 | -0.735870357 | 0.85984865 | 0.14015135 |
| daunorubicin | 15557 | BLCa92 | -1.239877906 | 0.76385709 | 0.23614291 |
| daunorubicin | 14547 | BLCa92 | -1.500260323 | 0.71426555 | 0.28573445 |
| daunorubicin | 15067 | BLCa92 | -1.366202049 | 0.73979783 | 0.26020217 |
| daunorubicin | 19200 | BLCa92 | -0.300696576 | 0.94273036 | 0.05726964 |
| daunorubicin | 12307 | BLCa92 | -2.077742118 | 0.60428034 | 0.39571966 |
| docetaxel | 22970 | BLCa92 | 0.671225911 | 1.12783939 | -0.1278394 |
| docetaxel | 22834 | BLCa92 | 0.636164516 | 1.12116172 | -0.1211617 |
| docetaxel | 19835 | BLCa92 | -0.136990799 | 0.9739092 | 0.0260908 |
| docetaxel | 25225 | BLCa92 | 1.252574772 | 1.23856111 | -0.2385611 |
| doxorubicin | 2804 | BLCa92 | -4.527657075 | 0.13767791 | 0.86232209 |
| doxorubicin | 1725 | BLCa92 | -4.805827993 | 0.08469843 | 0.91530157 |
| doxorubicin | 1973 | BLCa92 | -4.741892509 | 0.09687536 | 0.90312464 |
| doxorubicin | 2053 | BLCa92 | -4.721268159 | 0.10080341 | 0.89919659 |
| doxorubicin | 2360 | BLCa92 | -4.642122216 | 0.11587727 | 0.88412273 |
| epirubicin | 1673 | BLCa92 | -4.81923382 | 0.0821452 | 0.9178548 |
| epirubicin | 2014 | BLCa92 | -4.731322529 | 0.09888849 | 0.90111151 |
| epirubicin | 1905 | BLCa92 | -4.759423206 | 0.09353653 | 0.90646347 |
| epirubicin | 1762 | BLCa92 | -4.796289231 | 0.08651515 | 0.91348485 |
| epirubicin | 2842 | BLCa92 | -4.517860508 | 0.13954373 | 0.86045627 |
| erlotinib | 18105 | BLCa92 | -0.582992364 | 0.88896527 | 0.11103473 |
| erlotinib | 20505 | BLCa92 | 0.035738131 | 1.00680656 | -0.0068066 |
| erlotinib | 18453 | BLCa92 | -0.493276442 | 0.90605226 | 0.09394774 |
| erlotinib | 21927 | BLCa92 | 0.40233595 | 1.07662753 | -0.0766275 |
| erlotinib | 22872 | BLCa92 | 0.645961082 | 1.12302754 | -0.1230275 |
| everolimus | 16584 | BLCa92 | -0.975112815 | 0.81428335 | 0.18571665 |
| everolimus | 14503 | BLCa92 | -1.511603715 | 0.71210512 | 0.28789488 |
| everolimus | 18850 | BLCa92 | -0.390928106 | 0.92554517 | 0.07445483 |
| everolimus | 15088 | BLCa92 | -1.360788157 | 0.74082894 | 0.25917106 |
| everolimus | 14256 | BLCa92 | -1.575281395 | 0.69997729 | 0.30002271 |
| gemcitabine | 17329 | BLCa92 | -0.577009557 | 0.89993397 | 0.10006603 |
| gemcitabine | 15451 | BLCa92 | -1.139388534 | 0.80240521 | 0.19759479 |
| gemcitabine | 16192 | BLCa92 | -0.917491397 | 0.840887 | 0.159113 |
| gemcitabine | 16071 | BLCa92 | -0.953725613 | 0.8346032 | 0.1653968 |
| gemcitabine | 15894 | BLCa92 | -1.006729382 | 0.82541119 | 0.17458881 |
| lapatinib | 18280 | BLCa92 | -0.537876599 | 0.89755786 | 0.10244214 |
| lapatinib | 25449 | BLCa92 | 1.310322951 | 1.24955963 | -0.2495596 |
| lapatinib | 24845 | BLCa92 | 1.15460911 | 1.2199029 | -0.2199029 |
| lapatinib | 22515 | BLCa92 | 0.553924921 | 1.10549865 | -0.1054986 |
| lapatinib | 23560 | BLCa92 | 0.823330491 | 1.15680871 | -0.1568087 |
| methotrexate | 29278 | BLCa92 | 2.297455895 | 1.4375656 | -0.4375656 |
| methotrexate | 26680 | BLCa92 | 1.627680134 | 1.31000239 | -0.3100024 |
| methotrexate | 32795 | BLCa92 | 3.204153875 | 1.61025219 | -0.6102522 |
| methotrexate | 28172 | BLCa92 | 2.012324259 | 1.3832604 | -0.3832604 |

| | | | | | |
|---|---|---|---|---|---|
| methotrexate | 23554 | BLCa92 | 0.821783664 | 1.15651411 | -0.1565141 |
| olaparib | 28241 | BLCa92 | 2.03011276 | 1.38664834 | -0.3866483 |
| olaparib | 24055 | BLCa92 | 0.950943655 | 1.18111348 | -0.1811135 |
| olaparib | 18904 | BLCa92 | -0.37700667 | 0.9281966 | 0.0718034 |
| olaparib | 19185 | BLCa92 | -0.304563641 | 0.94199385 | 0.05800615 |
| olaparib | 28470 | BLCa92 | 2.089149962 | 1.39789236 | -0.3978924 |
| paclitaxel | 20202 | BLCa92 | -0.042376594 | 0.9919291 | 0.0080709 |
| paclitaxel | 14752 | BLCa92 | -1.447410426 | 0.72433116 | 0.27566884 |
| paclitaxel | 20618 | BLCa92 | 0.064870025 | 1.01235492 | -0.0123549 |
| paclitaxel | 23083 | BLCa92 | 0.700357805 | 1.13338775 | -0.1333878 |
| paclitaxel | 21982 | BLCa92 | 0.41651519 | 1.07932806 | -0.0793281 |
| ponatinib | 33350 | BLCa92 | 3.347235302 | 1.63750299 | -0.637503 |
| ponatinib | 13512 | BLCa92 | -1.767087849 | 0.66344649 | 0.33655351 |
| ponatinib | 17155 | BLCa92 | -0.827906518 | 0.84231975 | 0.15768025 |
| ponatinib | 12468 | BLCa92 | -2.036235614 | 0.61218553 | 0.38781447 |
| ponatinib | 18916 | BLCa92 | -0.373913017 | 0.92878581 | 0.07121419 |
| rapamycin | 16307 | BLCa92 | -1.046524626 | 0.8006825 | 0.1993175 |
| rapamycin | 13554 | BLCa92 | -1.756260065 | 0.66550871 | 0.33449129 |
| rapamycin | 14700 | BLCa92 | -1.460816254 | 0.72177793 | 0.27822207 |
| rapamycin | 16934 | BLCa92 | -0.884881285 | 0.83146854 | 0.16853146 |
| rapamycin | 14094 | BLCa92 | -1.617045704 | 0.692023 | 0.307977 |
| sunitinib | 14610 | BLCa92 | -1.484018647 | 0.71735888 | 0.28264112 |
| sunitinib | 10839 | BLCa92 | -2.456198938 | 0.53220075 | 0.46779925 |
| sunitinib | 8853 | BLCa92 | -2.968198423 | 0.43468708 | 0.56531292 |
| sunitinib | 12842 | BLCa92 | -1.939816779 | 0.63054913 | 0.36945087 |
| sunitinib | 14221 | BLCa92 | -1.584304548 | 0.69825877 | 0.30174123 |
| temsirolimus | 13799 | BLCa92 | -1.693097994 | 0.67753834 | 0.32246166 |
| temsirolimus | 16508 | BLCa92 | -0.994705948 | 0.81055171 | 0.18944829 |
| temsirolimus | 25864 | BLCa92 | 1.417311766 | 1.26993635 | -0.2699364 |
| temsirolimus | 14568 | BLCa92 | -1.494846431 | 0.71529666 | 0.28470334 |
| temsirolimus | 16050 | BLCa92 | -1.11278035 | 0.78806366 | 0.21193634 |
| vinblastine | 18822 | BLCa92 | -0.398146628 | 0.92417035 | 0.07582965 |
| vinblastine | 16987 | BLCa92 | -0.871217653 | 0.83407086 | 0.16592914 |
| vinblastine | 18728 | BLCa92 | -0.42238024 | 0.9195549 | 0.0804451 |
| vinblastine | 26566 | BLCa92 | 1.598290436 | 1.30440493 | -0.3044049 |
| mmc | 15935 | BLCa92 | -1.142427853 | 0.7824171 | 0.2175829 |
| mmc | 14861 | BLCa92 | -1.41930975 | 0.72968312 | 0.27031688 |
| mmc | 13604 | BLCa92 | -1.743369847 | 0.66796374 | 0.33203626 |
| mmc | 14018 | BLCa92 | -1.636638836 | 0.68829136 | 0.31170864 |
| erdafitinib | 21846 | BLCa92 | 0.381453795 | 1.07265039 | -0.0726504 |
| erdafitinib | 20305 | BLCa92 | -0.015822743 | 0.99698645 | 0.00301355 |
| erdafitinib | 19780 | BLCa92 | -0.151170039 | 0.97120867 | 0.02879133 |
| erdafitinib | 29904 | BLCa92 | 2.458841433 | 1.46830253 | -0.4683025 |
| cisplatin+gemcitabine | 12450 | BLCa92 | -2.038056962 | 0.64655652 | 0.35344348 |
| cisplatin+gemcitabine | 8962 | BLCa92 | -3.08256062 | 0.46541683 | 0.53458317 |
| cisplatin+gemcitabine | 14419 | BLCa92 | -1.44842746 | 0.74881112 | 0.25118888 |
| cisplatin+gemcitabine | 12930 | BLCa92 | -1.894317927 | 0.671484 | 0.328516 |
| untreated | 23240.6 | BLCa77 | 24.27622629 | 4.77889796 | |
| untreated | 3983.6 | BLCa77 | -1.161896778 | 0.81913625 | |
| untreated | 4804.6 | BLCa77 | -0.07737172 | 0.98795613 | |
| untreated | 4127.6 | BLCa77 | -0.971675574 | 0.84874656 | |
| untreated | 10480.6 | BLCa77 | 7.420514064 | 2.15509573 | |
| untreated | 6275.6 | BLCa77 | 1.865790716 | 1.29043364 | |
| untreated | 7431.6 | BLCa77 | 3.392844269 | 1.5281386 | |
| dmso | 4394.6 | BLCa77 | -0.618973758 | 0.903649 | 0.096351 |
| dmso | 4087.6 | BLCa77 | -1.024514797 | 0.84052147 | 0.15947853 |
| dmso | 4862.6 | BLCa77 | -0.000754846 | 0.9998825 | 0.0001175 |
| dmso | 5242.6 | BLCa77 | 0.501217775 | 1.07802081 | -0.0780208 |
| dmso | 4724.6 | BLCa77 | -0.183050166 | 0.97150595 | 0.02849405 |
| dmso | 4377.6 | BLCa77 | -0.641430428 | 0.90015334 | 0.09984666 |
| dmso | 6352.6 | BLCa77 | 1.967506221 | 1.30626693 | -0.3062669 |
| h2o | 5393.6 | BLCa77 | -1.259218942 | 0.91843476 | 0.08156524 |
| h2o | 6442.6 | BLCa77 | 1.498444252 | 1.09706093 | -0.0970609 |
| h2o | 5741.6 | BLCa77 | -0.344379293 | 0.97769302 | 0.02230698 |
| h2o | 5835.6 | BLCa77 | -0.097267434 | 0.99369955 | 0.00630045 |
| h2o | 5949.6 | BLCa77 | 0.202421416 | 1.01311174 | -0.0131117 |
| bosutinib | 6138.6 | BLCa77 | 1.684816377 | 1.26226272 | -0.2622627 |

| | | | | | |
|---|---|---|---|---|---|
| bosutinib | 6704.6 | BLCa77 | 2.432491386 | 1.37864768 | -0.3786477 |
| bosutinib | 7563.6 | BLCa77 | 3.567213706 | 1.55528139 | -0.5552814 |
| bosutinib | 5647.6 | BLCa77 | 1.036214911 | 1.1612998 | -0.1612998 |
| bosutinib | 5607.6 | BLCa77 | 0.983375688 | 1.15307471 | -0.1530747 |
| cisplatin_14_um | 4043.6 | BLCa77 | -4.808165854 | 0.68855362 | 0.31144638 |
| cisplatin_14_um | 4089.6 | BLCa77 | -4.687238774 | 0.69638661 | 0.30361339 |
| cisplatin_14_um | 2966.6 | BLCa77 | -7.639436835 | 0.50515955 | 0.49484045 |
| cisplatin_14_um | 2721.6 | BLCa77 | -8.283504979 | 0.46344038 | 0.53655962 |
| cisplatin_14_um | 3796.6 | BLCa77 | -5.457491697 | 0.64649389 | 0.35350611 |
| mmc | 6766.6 | BLCa77 | 2.514392182 | 1.39139656 | -0.3913966 |
| mmc | 5508.6 | BLCa77 | 0.85259861 | 1.13271763 | -0.1327176 |
| mmc | 6298.6 | BLCa77 | 1.89617327 | 1.29516306 | -0.2951631 |
| mmc | 2766.6 | BLCa77 | -2.769530146 | 0.56888803 | 0.43111197 |
| mmc | 4077.6 | BLCa77 | -1.037724603 | 0.8384652 | 0.1615348 |
| crizotinib | 4579.6 | BLCa77 | -0.374592351 | 0.94169002 | 0.05830998 |
| crizotinib | 4624.6 | BLCa77 | -0.315148225 | 0.95094324 | 0.04905676 |
| crizotinib | 4728.6 | BLCa77 | -0.177766244 | 0.97232846 | 0.02767154 |
| crizotinib | 4950.6 | BLCa77 | 0.115491445 | 1.01797769 | -0.0179777 |
| daunorubicin | 4470.6 | BLCa77 | -0.518579234 | 0.91927666 | 0.08072334 |
| daunorubicin | 4913.6 | BLCa77 | 0.066615164 | 1.01036948 | -0.0103695 |
| daunorubicin | 3638.6 | BLCa77 | -1.617635079 | 0.74819489 | 0.25180511 |
| daunorubicin | 3240.6 | BLCa77 | -2.14338535 | 0.66635529 | 0.33364471 |
| daunorubicin | 4591.6 | BLCa77 | -0.358740584 | 0.94415755 | 0.05584245 |
| docetaxel | 6972.6 | BLCa77 | 2.786514182 | 1.43375575 | -0.4337558 |
| docetaxel | 5794.6 | BLCa77 | 1.230399056 | 1.19152699 | -0.191527 |
| docetaxel | 4648.6 | BLCa77 | -0.283444691 | 0.95587829 | 0.04412171 |
| docetaxel | 4748.6 | BLCa77 | -0.151346632 | 0.97644101 | 0.02355899 |
| docetaxel | 3912.6 | BLCa77 | -1.255686399 | 0.80453672 | 0.19546328 |
| doxorubicin | 4537.6 | BLCa77 | -0.430073535 | 0.93305368 | 0.06694632 |
| doxorubicin | 5111.6 | BLCa77 | 0.328169319 | 1.05108365 | -0.0510837 |
| doxorubicin | 3219.6 | BLCa77 | -2.171125942 | 0.66203712 | 0.33796288 |
| doxorubicin | 4311.6 | BLCa77 | -0.728615147 | 0.88658195 | 0.11341805 |
| doxorubicin | 5936.6 | BLCa77 | 1.417978299 | 1.22072604 | -0.220726 |
| epirubicin | 5204.6 | BLCa77 | 0.451020513 | 1.07020698 | -0.070207 |
| epirubicin | 7440.6 | BLCa77 | 3.404733095 | 1.52998925 | -0.5299892 |
| epirubicin | 8161.6 | BLCa77 | 4.357160094 | 1.67824641 | -0.6782464 |
| epirubicin | 3773.6 | BLCa77 | -1.4393027 | 0.77595455 | 0.22404545 |
| epirubicin | 2895.6 | BLCa77 | -2.599123651 | 0.59541393 | 0.40458607 |
| erlotinib | 4984.6 | BLCa77 | 0.160404785 | 1.02496901 | -0.024969 |
| erlotinib | 10005.6 | BLCa77 | 6.793048288 | 2.05742285 | -1.0574228 |
| erlotinib | 7004.6 | BLCa77 | 2.828785561 | 1.44033582 | -0.4403358 |
| erlotinib | 6052.6 | BLCa77 | 1.571212047 | 1.24457879 | -0.2445788 |
| erlotinib | 7798.6 | BLCa77 | 3.877644143 | 1.60360376 | -0.6036038 |
| everolimus | 8210.6 | BLCa77 | 4.421888143 | 1.68832214 | -0.6883221 |
| everolimus | 6586.6 | BLCa77 | 2.276615678 | 1.35438368 | -0.3543837 |
| everolimus | 5309.6 | BLCa77 | 0.589723474 | 1.09179783 | -0.0917978 |
| everolimus | 7873.6 | BLCa77 | 3.976717687 | 1.6190258 | -0.6190258 |
| everolimus | 5724.6 | BLCa77 | 1.137930416 | 1.17713309 | -0.1771331 |
| gemcitabine | 6934.6 | BLCa77 | 2.791838238 | 1.18083983 | -0.1808398 |
| gemcitabine | 5591.6 | BLCa77 | -0.738706728 | 0.95215067 | 0.04784933 |
| gemcitabine | 6732.6 | BLCa77 | 2.260810626 | 1.1464428 | -0.1464428 |
| gemcitabine | 5037.6 | BLCa77 | -2.195089387 | 0.85781426 | 0.14218574 |
| gemcitabine | 6477.6 | BLCa77 | 1.590453987 | 1.10302081 | -0.1030208 |
| lapatinib | 6736.6 | BLCa77 | 2.474762765 | 1.38522775 | -0.3852277 |
| lapatinib | 4284.6 | BLCa77 | -0.764281623 | 0.88103002 | 0.11896998 |
| lapatinib | 6712.6 | BLCa77 | 2.443059231 | 1.3802927 | -0.3802927 |
| lapatinib | 6009.6 | BLCa77 | 1.514409882 | 1.23573682 | -0.2357368 |
| lapatinib | 4295.6 | BLCa77 | -0.749750836 | 0.88329191 | 0.11670809 |
| methotrexate | 5088.6 | BLCa77 | 0.297786766 | 1.04635423 | -0.0463542 |
| methotrexate | 6829.6 | BLCa77 | 2.597613959 | 1.40435107 | -0.4043511 |
| methotrexate | 6091.6 | BLCa77 | 1.622730289 | 1.25259825 | -0.2525982 |
| methotrexate | 7574.6 | BLCa77 | 3.581744493 | 1.55754328 | -0.5575433 |
| olaparib | 6662.6 | BLCa77 | 2.377010202 | 1.37001134 | -0.3700113 |
| olaparib | 6903.6 | BLCa77 | 2.695366522 | 1.41956748 | -0.4195675 |
| olaparib | 6962.6 | BLCa77 | 2.773304376 | 1.43169948 | -0.4316995 |
| olaparib | 5513.6 | BLCa77 | 0.859203513 | 1.13374576 | -0.1337458 |
| paclitaxel | 6697.6 | BLCa77 | 2.423244522 | 1.37720829 | -0.3772083 |

| | | | | | |
|---|---|---|---|---|---|
| paclitaxel | 6647.6 | BLCa77 | 2.357195493 | 1.36692693 | -0.3669269 |
| paclitaxel | 7283.6 | BLCa77 | 3.197339143 | 1.49770579 | -0.4977058 |
| paclitaxel | 5654.6 | BLCa77 | 1.045461775 | 1.16273919 | -0.1627392 |
| paclitaxel | 6148.6 | BLCa77 | 1.698026183 | 1.26431899 | -0.264319 |
| ponatinib | 6432.6 | BLCa77 | 2.073184668 | 1.3227171 | -0.3227171 |
| ponatinib | 7171.6 | BLCa77 | 3.049389318 | 1.47467555 | -0.4746755 |
| ponatinib | 5840.6 | BLCa77 | 1.291164163 | 1.20098584 | -0.2009858 |
| ponatinib | 6661.6 | BLCa77 | 2.375689221 | 1.36980571 | -0.3698057 |
| ponatinib | 8822.6 | BLCa77 | 5.230328259 | 1.81416595 | -0.8141659 |
| rapamycin | 8745.6 | BLCa77 | 5.128612754 | 1.79833266 | -0.7983327 |
| rapamycin | 5971.6 | BLCa77 | 1.46421262 | 1.22792299 | -0.227923 |
| rapamycin | 5260.6 | BLCa77 | 0.524995426 | 1.0817221 | -0.0817221 |
| rapamycin | 4643.6 | BLCa77 | -0.290049594 | 0.95485016 | 0.04514984 |
| rapamycin | 5604.6 | BLCa77 | 0.979412746 | 1.15245783 | -0.1524578 |
| sunitinib | 7883.6 | BLCa77 | 3.989927493 | 1.62108207 | -0.6210821 |
| sunitinib | 7417.6 | BLCa77 | 3.374350541 | 1.52525982 | -0.5252598 |
| sunitinib | 7268.6 | BLCa77 | 3.177524435 | 1.49462138 | -0.4946214 |
| sunitinib | 7429.6 | BLCa77 | 3.390202308 | 1.52772735 | -0.5277274 |
| sunitinib | 6941.6 | BLCa77 | 2.745563784 | 1.42738131 | -0.4273813 |
| temsirolimus | 6368.6 | BLCa77 | 1.988641911 | 1.30955696 | -0.309557 |
| temsirolimus | 5699.6 | BLCa77 | 1.104905901 | 1.17199241 | -0.1719924 |
| temsirolimus | 6764.6 | BLCa77 | 2.511750221 | 1.39098531 | -0.3909853 |
| temsirolimus | 5310.6 | BLCa77 | 0.591044455 | 1.09200345 | -0.0920035 |
| temsirolimus | 5755.6 | BLCa77 | 1.178880814 | 1.18350753 | -0.1835075 |
| vinblastine | 4733.6 | BLCa77 | -0.171161341 | 0.9733566 | 0.0266434 |
| vinblastine | 7649.6 | BLCa77 | 3.680818036 | 1.57296532 | -0.5729653 |
| vinblastine | 6264.6 | BLCa77 | 1.85125993 | 1.28817174 | -0.2881717 |
| vinblastine | 6081.6 | BLCa77 | 1.609520484 | 1.25054197 | -0.250542 |
| vinblastine | 7231.6 | BLCa77 | 3.128648153 | 1.48701318 | -0.4870132 |
| cisplatin+gemcitabine | 2447.6 | BLCa77 | -9.00380976 | 0.41678303 | 0.58321697 |
| cisplatin+gemcitabine | 2246.6 | BLCa77 | -9.532208522 | 0.38255628 | 0.61744372 |
| cisplatin+gemcitabine | 1705.6 | BLCa77 | -10.95441614 | 0.29043354 | 0.70956646 |
| cisplatin+gemcitabine | 2346.6 | BLCa77 | -9.269323566 | 0.39958451 | 0.60041549 |
| cisplatin+gemcitabine | 1932.6 | BLCa77 | -10.35766729 | 0.32908763 | 0.67091237 |
| cisplatin_6_um | 12061.6 | BLCa77 | 16.26994996 | 2.05387733 | -1.0538773 |
| cisplatin_6_um | 6105.6 | BLCa77 | 0.612521949 | 1.03967578 | -0.0396758 |
| cisplatin_6_um | 10853.6 | BLCa77 | 13.09429968 | 1.84817628 | -0.8481763 |
| cisplatin_6_um | 7836.6 | BLCa77 | 5.163060545 | 1.33443449 | -0.3344345 |
| cisplatin_6_um | 7215.6 | BLCa77 | 3.530544966 | 1.22868917 | -0.2286892 |
| untreated | 4504 | BLCa85 | 0.242545519 | 1.0444058 | |
| untreated | 4510 | BLCa85 | 0.250144856 | 1.0457971 | |
| untreated | 5048 | BLCa85 | 0.931552111 | 1.17055072 | |
| untreated | 4795 | BLCa85 | 0.611113383 | 1.11188406 | |
| untreated | 3548 | BLCa85 | -0.968282242 | 0.82272464 | |
| untreated | 5233 | BLCa85 | 1.165865014 | 1.21344928 | |
| untreated | 4188 | BLCa85 | -0.157686251 | 0.97113043 | |
| untreated | 5009 | BLCa85 | 0.882156418 | 1.16150725 | |
| untreated | 3758 | BLCa85 | -0.702305432 | 0.87142029 | |
| dmso | 4886 | BLCa85 | 0.726370001 | 1.13298551 | -0.1329855 |
| dmso | 3704 | BLCa85 | -0.770699469 | 0.85889855 | 0.14110145 |
| dmso | 3466 | BLCa85 | -1.072139853 | 0.80371014 | 0.19628986 |
| dmso | 4469 | BLCa85 | 0.198216051 | 1.03628986 | -0.0362899 |
| dmso | 5490 | BLCa85 | 1.491369967 | 1.27304348 | -0.2730435 |
| dmso | 3060 | BLCa85 | -1.586361684 | 0.70956522 | 0.29043478 |
| dmso | 3739 | BLCa85 | -0.726370001 | 0.86701449 | 0.13298551 |
| dmso | 4447 | BLCa85 | 0.170351814 | 1.03118841 | -0.0311884 |
| dmso | 4708 | BLCa85 | 0.500922991 | 1.09171014 | -0.0917101 |
| dmso | 5156 | BLCa85 | 1.068340184 | 1.1955942 | -0.1955942 |
| h2o | 2571 | BLCa85 | -1.479319122 | 0.70388767 | 0.29611233 |
| h2o | 4122 | BLCa85 | 0.642060842 | 1.12852003 | -0.12852 |
| h2o | 3858 | BLCa85 | 0.280974891 | 1.05624218 | -0.0562422 |
| h2o | 4869 | BLCa85 | 1.663769954 | 1.33303348 | -0.3330335 |
| h2o | 3352 | BLCa85 | -0.411106516 | 0.91770964 | 0.08229036 |
| h2o | 3204 | BLCa85 | -0.613533489 | 0.87719024 | 0.12280976 |
| h2o | 3592 | BLCa85 | -0.08284656 | 0.98341677 | 0.01658323 |
| bosutinib | 4530 | BLCa85 | 0.275475981 | 1.05043478 | -0.0504348 |
| bosutinib | 5076 | BLCa85 | 0.967015686 | 1.17704348 | -0.1770435 |

| | | | | | |
|---|---|---|---|---|---|
| bosutinib | 2696 | BLCa85 | -2.047388154 | 0.62515942 | 0.37484058 |
| bosutinib | 3508 | BLCa85 | -1.018944491 | 0.81344928 | 0.18655072 |
| bosutinib | 3881 | BLCa85 | -0.546519015 | 0.89994203 | 0.10005797 |
| cisplatin_6_um | 3471 | BLCa85 | -0.248344288 | 0.95028942 | 0.04971058 |
| cisplatin_6_um | 3728 | BLCa85 | 0.103167415 | 1.02065081 | -0.0206508 |
| cisplatin_6_um | 3341 | BLCa85 | -0.426151764 | 0.91469806 | 0.08530194 |
| cisplatin_6_um | 3403 | BLCa85 | -0.341351275 | 0.9316724 | 0.0683276 |
| cisplatin_6_um | 3557 | BLCa85 | -0.130717804 | 0.97383448 | 0.02616552 |
| cisplatin_14_um | 3524 | BLCa85 | -0.175853548 | 0.96479975 | 0.03520025 |
| cisplatin_14_um | 3402 | BLCa85 | -0.342719025 | 0.93139862 | 0.06860138 |
| cisplatin_14_um | 3753 | BLCa85 | 0.13736116 | 1.02749531 | -0.0274953 |
| cisplatin_14_um | 3737 | BLCa85 | 0.115477163 | 1.02311483 | -0.0231148 |
| crizotinib | 2774 | BLCa85 | -1.948596768 | 0.64324638 | 0.35675362 |
| crizotinib | 2868 | BLCa85 | -1.829540482 | 0.66504348 | 0.33495652 |
| crizotinib | 3018 | BLCa85 | -1.639557046 | 0.69982609 | 0.30017391 |
| crizotinib | 2990 | BLCa85 | -1.675020621 | 0.69333333 | 0.30666667 |
| crizotinib | 2611 | BLCa85 | -2.155045434 | 0.60544928 | 0.39455072 |
| daunorubicin | 3366 | BLCa85 | -1.198795477 | 0.78052174 | 0.21947826 |
| daunorubicin | 4105 | BLCa85 | -0.262810419 | 0.95188406 | 0.04811594 |
| daunorubicin | 3222 | BLCa85 | -1.381179574 | 0.74713043 | 0.25286957 |
| daunorubicin | 3189 | BLCa85 | -1.42297593 | 0.73947826 | 0.26052174 |
| daunorubicin | 2715 | BLCa85 | -2.023323586 | 0.62956522 | 0.37043478 |
| docetaxel | 4591 | BLCa85 | 0.352735911 | 1.06457971 | -0.0645797 |
| docetaxel | 4768 | BLCa85 | 0.576916365 | 1.10562319 | -0.1056232 |
| docetaxel | 3242 | BLCa85 | -1.35584845 | 0.75176812 | 0.24823188 |
| docetaxel | 3446 | BLCa85 | -1.097470978 | 0.79907246 | 0.20092754 |
| docetaxel | 3357 | BLCa85 | -1.210194483 | 0.77843478 | 0.22156522 |
| doxorubicin | 3925 | BLCa85 | -0.490790541 | 0.91014493 | 0.08985507 |
| doxorubicin | 3422 | BLCa85 | -1.127868327 | 0.79350725 | 0.20649275 |
| doxorubicin | 4395 | BLCa85 | 0.104490889 | 1.01913043 | -0.0191304 |
| doxorubicin | 5182 | BLCa85 | 1.101270646 | 1.20162319 | -0.2016232 |
| doxorubicin | 3097 | BLCa85 | -1.539499104 | 0.71814493 | 0.28185507 |
| epirubicin | 4101 | BLCa85 | -0.267876644 | 0.95095652 | 0.04904348 |
| epirubicin | 3461 | BLCa85 | -1.078472634 | 0.80255072 | 0.19744928 |
| epirubicin | 3435 | BLCa85 | -1.111403096 | 0.79652174 | 0.20347826 |
| epirubicin | 2453 | BLCa85 | -2.355161319 | 0.56881159 | 0.43118841 |
| epirubicin | 2812 | BLCa85 | -1.900467631 | 0.65205797 | 0.34794203 |
| erlotinib | 4443 | BLCa85 | 0.165285589 | 1.03026087 | -0.0302609 |
| erlotinib | 4182 | BLCa85 | -0.165285589 | 0.96973913 | 0.03026087 |
| erlotinib | 3915 | BLCa85 | -0.503456103 | 0.90782609 | 0.09217391 |
| erlotinib | 3144 | BLCa85 | -1.479970961 | 0.72904348 | 0.27095652 |
| erlotinib | 4084 | BLCa85 | -0.2894081 | 0.94701449 | 0.05298551 |
| everolimus | 3448 | BLCa85 | -1.094937865 | 0.79953623 | 0.20046377 |
| everolimus | 2427 | BLCa85 | -2.388091781 | 0.56278261 | 0.43721739 |
| everolimus | 2940 | BLCa85 | -1.738348433 | 0.68173913 | 0.31826087 |
| everolimus | 3120 | BLCa85 | -1.51036831 | 0.72347826 | 0.27652174 |
| everolimus | 3250 | BLCa85 | -1.345716 | 0.75362319 | 0.24637681 |
| gemcitabine | 3123 | BLCa85 | -0.724321224 | 0.85501408 | 0.14498592 |
| gemcitabine | 3815 | BLCa85 | 0.222161649 | 1.04446965 | -0.0444696 |
| gemcitabine | 3799 | BLCa85 | 0.200277652 | 1.04008917 | -0.0400892 |
| gemcitabine | 3974 | BLCa85 | 0.439633869 | 1.08800063 | -0.0880006 |
| gemcitabine | 3489 | BLCa85 | -0.223724791 | 0.95521746 | 0.04478254 |
| lapatinib | 1955 | BLCa85 | -2.985906324 | 0.45333333 | 0.54666667 |
| lapatinib | 3511 | BLCa85 | -1.015144822 | 0.81414493 | 0.18585507 |
| lapatinib | 2322 | BLCa85 | -2.521080186 | 0.53843478 | 0.46156522 |
| lapatinib | 3069 | BLCa85 | -1.574962678 | 0.71165217 | 0.28834783 |
| lapatinib | 2505 | BLCa85 | -2.289300395 | 0.58086957 | 0.41913043 |
| methotrexate | 4730 | BLCa85 | 0.528787228 | 1.09681159 | -0.0968116 |
| methotrexate | 3136 | BLCa85 | -1.490103411 | 0.72718841 | 0.27281159 |
| methotrexate | 4551 | BLCa85 | 0.302073662 | 1.05530435 | -0.0553043 |
| methotrexate | 4012 | BLCa85 | -0.380600149 | 0.93031884 | 0.06968116 |
| methotrexate | 3120 | BLCa85 | -1.51036831 | 0.72347826 | 0.27652174 |
| olaparib | 2280 | BLCa85 | -2.574275548 | 0.52869565 | 0.47130435 |
| olaparib | 2782 | BLCa85 | -1.938464318 | 0.64510145 | 0.35489855 |
| olaparib | 4353 | BLCa85 | 0.051295528 | 1.0093913 | -0.0093913 |
| olaparib | 3391 | BLCa85 | -1.167131571 | 0.78631884 | 0.21368116 |
| olaparib | 3966 | BLCa85 | -0.438861735 | 0.91965217 | 0.08034783 |

| | | | | | |
|---|---|---|---|---|---|
| paclitaxel | 4862 | BLCa85 | 0.695972651 | 1.12742029 | -0.1274203 |
| paclitaxel | 2573 | BLCa85 | -2.203174571 | 0.59663768 | 0.40336232 |
| paclitaxel | 4914 | BLCa85 | 0.761833575 | 1.13947826 | -0.1394783 |
| paclitaxel | 5384 | BLCa85 | 1.357115006 | 1.24846377 | -0.2484638 |
| paclitaxel | 4131 | BLCa85 | -0.229879957 | 0.95791304 | 0.04208696 |
| ponatinib | 3447 | BLCa85 | -1.096204422 | 0.79930435 | 0.20069565 |
| ponatinib | 3710 | BLCa85 | -0.763100132 | 0.86028986 | 0.13971014 |
| ponatinib | 5326 | BLCa85 | 1.283654744 | 1.23501449 | -0.2350145 |
| ponatinib | 4844 | BLCa85 | 0.673174639 | 1.12324638 | -0.1232464 |
| ponatinib | 3495 | BLCa85 | -1.035409722 | 0.81043478 | 0.18956522 |
| rapamycin | 3533 | BLCa85 | -0.987280585 | 0.81924638 | 0.18075362 |
| rapamycin | 4716 | BLCa85 | 0.511055441 | 1.09356522 | -0.0935652 |
| rapamycin | 4684 | BLCa85 | 0.470525641 | 1.08614493 | -0.0861449 |
| rapamycin | 5273 | BLCa85 | 1.216527264 | 1.22272464 | -0.2227246 |
| rapamycin | 2974 | BLCa85 | -1.695285521 | 0.68962319 | 0.31037681 |
| sunitinib | 2994 | BLCa85 | -1.669954396 | 0.69426087 | 0.30573913 |
| sunitinib | 3813 | BLCa85 | -0.632644839 | 0.88417391 | 0.11582609 |
| sunitinib | 3956 | BLCa85 | -0.451527298 | 0.91733333 | 0.08266667 |
| sunitinib | 4402 | BLCa85 | 0.113356783 | 1.02075362 | -0.0207536 |
| sunitinib | 2671 | BLCa85 | -2.07905206 | 0.61936232 | 0.38063768 |
| temsirolimus | 3092 | BLCa85 | -1.545831885 | 0.71698551 | 0.28301449 |
| temsirolimus | 5248 | BLCa85 | 1.184863358 | 1.21692754 | -0.2169275 |
| temsirolimus | 3709 | BLCa85 | -0.764366688 | 0.86005797 | 0.13994203 |
| temsirolimus | 4033 | BLCa85 | -0.354002468 | 0.93518841 | 0.06481159 |
| temsirolimus | 3467 | BLCa85 | -1.070873297 | 0.80394203 | 0.19605797 |
| vinblastine | 3671 | BLCa85 | -0.812495825 | 0.85124638 | 0.14875362 |
| vinblastine | 3065 | BLCa85 | -1.580028903 | 0.71072464 | 0.28927536 |
| vinblastine | 2516 | BLCa85 | -2.275368276 | 0.58342029 | 0.41657971 |
| vinblastine | 4002 | BLCa85 | -0.393265711 | 0.928 | 0.072 |
| vinblastine | 3202 | BLCa85 | -1.406510699 | 0.74249275 | 0.25750725 |
| mmc | 2875 | BLCa85 | -1.820674588 | 0.66666667 | 0.33333333 |
| mmc | 3430 | BLCa85 | -1.117735877 | 0.79536232 | 0.20463768 |
| mmc | 3849 | BLCa85 | -0.587048815 | 0.89252174 | 0.10747826 |
| mmc | 4082 | BLCa85 | -0.291941212 | 0.94655072 | 0.05344928 |
| mmc | 4380 | BLCa85 | 0.085492546 | 1.01565217 | -0.0156522 |
| erdafitinib | 3890 | BLCa85 | -0.535120009 | 0.90202899 | 0.09797101 |
| erdafitinib | 3967 | BLCa85 | -0.437595179 | 0.91988406 | 0.08011594 |
| erdafitinib | 4704 | BLCa85 | 0.495856766 | 1.09078261 | -0.0907826 |
| erdafitinib | 3931 | BLCa85 | -0.483191204 | 0.91153623 | 0.08846377 |
| erdafitinib | 3664 | BLCa85 | -0.821361718 | 0.84962319 | 0.15037681 |
| cisplatin+gemcitabine | 3027 | BLCa85 | -0.855625206 | 0.82873123 | 0.17126877 |
| cisplatin+gemcitabine | 3750 | BLCa85 | 0.133257911 | 1.02667397 | -0.026674 |
| cisplatin+gemcitabine | 3307 | BLCa85 | -0.472655258 | 0.90538955 | 0.09461045 |
| cisplatin+gemcitabine | 2717 | BLCa85 | -1.279627649 | 0.74385951 | 0.25614049 |
| cisplatin+gemcitabine | 3652 | BLCa85 | -0.000781571 | 0.99984355 | 0.00015645 |
| untreated | 9489 | BLCa100 | -1.555865031 | 0.77480199 | |
| untreated | 14511 | BLCa100 | 1.277185798 | 1.1848616 | |
| untreated | 13677 | BLCa100 | 0.806703044 | 1.11676329 | |
| untreated | 15521 | BLCa100 | 1.846955081 | 1.26733077 | |
| untreated | 14136 | BLCa100 | 1.065637797 | 1.15424186 | |
| untreated | 12763 | BLCa100 | 0.291090049 | 1.04213277 | |
| untreated | 13700 | BLCa100 | 0.819677988 | 1.1186413 | |
| untreated | 11411 | BLCa100 | -0.47161101 | 0.93173838 | |
| untreated | 8713 | BLCa100 | -1.993628361 | 0.71143954 | |
| untreated | 10170 | BLCa100 | -1.171693861 | 0.83040745 | |
| dmso | 9337 | BLCa100 | -1.641612488 | 0.76239079 | 0.23760921 |
| dmso | 15382 | BLCa100 | 1.768541288 | 1.25598106 | -0.2559811 |
| dmso | 12773 | BLCa100 | 0.296731329 | 1.04294929 | -0.0429493 |
| dmso | 11455 | BLCa100 | -0.446789378 | 0.9353311 | 0.0646689 |
| dmso | 12765 | BLCa100 | 0.292218305 | 1.04229607 | -0.0422961 |
| dmso | 13165 | BLCa100 | 0.517869506 | 1.07495713 | -0.0749571 |
| dmso | 12146 | BLCa100 | -0.056976928 | 0.99175308 | 0.00824692 |
| dmso | 10953 | BLCa100 | -0.729981635 | 0.89434147 | 0.10565853 |
| h2o | 12201 | BLCa100 | 0.087159871 | 1.01691007 | -0.0169101 |
| h2o | 15729 | BLCa100 | 1.602767871 | 1.31095635 | -0.3109564 |
| h2o | 12637 | BLCa100 | 0.2744629 | 1.05324912 | -0.0532491 |
| h2o | 13326 | BLCa100 | 0.570453238 | 1.11067483 | -0.1106748 |

| | | | | | |
|---|---|---|---|---|---|
| h2o | 13350 | BLCa100 | 0.580763497 | 1.11267514 | -0.1126751 |
| h2o | 10773 | BLCa100 | -0.526300511 | 0.89789133 | 0.10210867 |
| h2o | 9060 | BLCa100 | -1.262195212 | 0.75511886 | 0.24488114 |
| h2o | 8164 | BLCa100 | -1.647111529 | 0.68044044 | 0.31955956 |
| h2o | 12743 | BLCa100 | 0.319999875 | 1.06208385 | -0.0620838 |
| bosutinib | 10496 | BLCa100 | -0.987788133 | 0.85702621 | 0.14297379 |
| bosutinib | 7446 | BLCa100 | -2.708378541 | 0.60798563 | 0.39201437 |
| bosutinib | 9137 | BLCa100 | -1.754438088 | 0.74606026 | 0.25393974 |
| bosutinib | 9856 | BLCa100 | -1.348830054 | 0.80476851 | 0.19523149 |
| cisplatin_6_um | 14236 | BLCa100 | 0.961383873 | 1.1865201 | -0.1865201 |
| cisplatin_6_um | 12892 | BLCa100 | 0.384009397 | 1.07450247 | -0.0745025 |
| cisplatin_6_um | 16142 | BLCa100 | 1.780190236 | 1.34537844 | -0.3453784 |
| cisplatin_6_um | 13804 | BLCa100 | 0.77579922 | 1.15051443 | -0.1505144 |
| cisplatin_14_um | 12109 | BLCa100 | 0.047637213 | 1.0092422 | -0.0092422 |
| cisplatin_14_um | 13654 | BLCa100 | 0.711360104 | 1.13801246 | -0.1380125 |
| cisplatin_14_um | 13462 | BLCa100 | 0.628878036 | 1.12200995 | -0.1220099 |
| cisplatin_14_um | 9787 | BLCa100 | -0.949880298 | 0.81571173 | 0.18428827 |
| crizotinib | 7617 | BLCa100 | -2.611912652 | 0.62194823 | 0.37805177 |
| crizotinib | 9516 | BLCa100 | -1.540633575 | 0.77700661 | 0.22299339 |
| crizotinib | 7953 | BLCa100 | -2.422365643 | 0.64938352 | 0.35061648 |
| crizotinib | 10289 | BLCa100 | -1.104562629 | 0.84012411 | 0.15987589 |
| daunorubicin | 11165 | BLCa100 | -0.610386499 | 0.91165183 | 0.08834817 |
| daunorubicin | 11942 | BLCa100 | -0.172059041 | 0.97509594 | 0.02490406 |
| daunorubicin | 9571 | BLCa100 | -1.509606535 | 0.78149751 | 0.21850249 |
| daunorubicin | 9994 | BLCa100 | -1.27098039 | 0.81603658 | 0.18396342 |
| docetaxel | 15775 | BLCa100 | 1.990243593 | 1.28807055 | -0.2880705 |
| docetaxel | 13774 | BLCa100 | 0.86142346 | 1.1246836 | -0.1246836 |
| docetaxel | 15663 | BLCa100 | 1.927061257 | 1.27892545 | -0.2789255 |
| docetaxel | 9532 | BLCa100 | -1.531607527 | 0.77831306 | 0.22168694 |
| doxorubicin | 7436 | BLCa100 | -2.714019821 | 0.6071691 | 0.3928309 |
| doxorubicin | 6433 | BLCa100 | -3.279840207 | 0.5252715 | 0.4747285 |
| doxorubicin | 9294 | BLCa100 | -1.665869992 | 0.75887973 | 0.24112027 |
| doxorubicin | 9020 | BLCa100 | -1.820441064 | 0.7365069 | 0.2634931 |
| epirubicin | 3665 | BLCa100 | -4.841346519 | 0.29925696 | 0.70074304 |
| epirubicin | 7497 | BLCa100 | -2.679608012 | 0.61214991 | 0.38785009 |
| epirubicin | 6527 | BLCa100 | -3.226812175 | 0.53294684 | 0.46705316 |
| epirubicin | 7157 | BLCa100 | -2.871411533 | 0.58438801 | 0.41561199 |
| erlotinib | 13528 | BLCa100 | 0.722647971 | 1.10459704 | -0.104597 |
| erlotinib | 13804 | BLCa100 | 0.8783473 | 1.12713318 | -0.1271332 |
| erlotinib | 12207 | BLCa100 | -0.02256512 | 0.99673389 | 0.00326611 |
| erlotinib | 10759 | BLCa100 | -0.839422468 | 0.87850086 | 0.12149914 |
| everolimus | 10256 | BLCa100 | -1.123178853 | 0.83742957 | 0.16257043 |
| everolimus | 9866 | BLCa100 | -1.343188774 | 0.80558504 | 0.19441496 |
| everolimus | 13023 | BLCa100 | 0.43776333 | 1.06336246 | -0.0633625 |
| everolimus | 12650 | BLCa100 | 0.227343585 | 1.03290602 | -0.032906 |
| gemcitabine | 12624 | BLCa100 | 0.268878177 | 1.05216562 | -0.0521656 |
| gemcitabine | 13268 | BLCa100 | 0.54553678 | 1.10584073 | -0.1058407 |
| gemcitabine | 9236 | BLCa100 | -1.186586649 | 0.76978784 | 0.23021216 |
| gemcitabine | 6669 | BLCa100 | -2.289354715 | 0.55583749 | 0.44416251 |
| lapatinib | 9333 | BLCa100 | -1.643869 | 0.76206418 | 0.23793582 |
| lapatinib | 5487 | BLCa100 | -3.813505298 | 0.44802809 | 0.55197191 |
| lapatinib | 4901 | BLCa100 | -4.144084307 | 0.40017964 | 0.59982036 |
| lapatinib | 11049 | BLCa100 | -0.675825347 | 0.90218013 | 0.09781987 |
| methotrexate | 11202 | BLCa100 | -0.589513763 | 0.91467298 | 0.08532702 |
| methotrexate | 8310 | BLCa100 | -2.220971946 | 0.67853352 | 0.32146648 |
| methotrexate | 13939 | BLCa100 | 0.95450458 | 1.13815628 | -0.1381563 |
| methotrexate | 11538 | BLCa100 | -0.399966754 | 0.94210827 | 0.05789173 |
| olaparib | 7874 | BLCa100 | -2.466931755 | 0.64293296 | 0.35706704 |
| olaparib | 13945 | BLCa100 | 0.957889348 | 1.1386462 | -0.1386462 |
| olaparib | 11700 | BLCa100 | -0.308578017 | 0.955336 | 0.044664 |
| olaparib | 12846 | BLCa100 | 0.337912674 | 1.04890994 | -0.0489099 |
| paclitaxel | 14458 | BLCa100 | 1.247287014 | 1.18053401 | -0.180534 |
| paclitaxel | 10684 | BLCa100 | -0.881732068 | 0.87237691 | 0.12762309 |
| paclitaxel | 11560 | BLCa100 | -0.387555938 | 0.94390463 | 0.05609537 |
| paclitaxel | 12327 | BLCa100 | 0.04513024 | 1.00653221 | -0.0065322 |
| ponatinib | 13349 | BLCa100 | 0.621669059 | 1.08998122 | -0.0899812 |
| ponatinib | 10806 | BLCa100 | -0.812908452 | 0.88233853 | 0.11766147 |

| | | | | | |
|---|---|---|---|---|---|
| ponatinib | 9871 | BLCa100 | -1.340368134 | 0.8059933 | 0.1940067 |
| ponatinib | 11035 | BLCa100 | -0.683723139 | 0.90103699 | 0.09896301 |
| rapamycin | 13062 | BLCa100 | 0.459764322 | 1.06654691 | -0.0665469 |
| rapamycin | 7665 | BLCa100 | -2.584834508 | 0.62586756 | 0.37413244 |
| rapamycin | 11461 | BLCa100 | -0.44340461 | 0.93582102 | 0.06417898 |
| rapamycin | 13022 | BLCa100 | 0.437199202 | 1.0632808 | -0.0632808 |
| sunitinib | 7960 | BLCa100 | -2.418416747 | 0.64995509 | 0.35004491 |
| sunitinib | 5372 | BLCa100 | -3.878380018 | 0.43863803 | 0.56136197 |
| sunitinib | 9654 | BLCa100 | -1.462783911 | 0.78827468 | 0.21172532 |
| sunitinib | 11671 | BLCa100 | -0.32493773 | 0.95296807 | 0.04703193 |
| temsirolimus | 12758 | BLCa100 | 0.288269409 | 1.0417245 | -0.0417245 |
| temsirolimus | 9110 | BLCa100 | -1.769669544 | 0.74385564 | 0.25614436 |
| temsirolimus | 10426 | BLCa100 | -1.027277093 | 0.85131053 | 0.14868947 |
| temsirolimus | 13412 | BLCa100 | 0.657209123 | 1.09512534 | -0.0951253 |
| vinblastine | 11163 | BLCa100 | -0.611514755 | 0.91148853 | 0.08851147 |
| vinblastine | 10178 | BLCa100 | -1.167180837 | 0.83106067 | 0.16893933 |
| vinblastine | 12135 | BLCa100 | -0.063182336 | 0.9908549 | 0.0091451 |
| vinblastine | 13823 | BLCa100 | 0.889065732 | 1.12868458 | -0.1286846 |
| mmc | 11971 | BLCa100 | -0.155699329 | 0.97746387 | 0.02253613 |
| mmc | 9104 | BLCa100 | -1.773054312 | 0.74336572 | 0.25663428 |
| mmc | 11675 | BLCa100 | -0.322681218 | 0.95329468 | 0.04670532 |
| mmc | 9561 | BLCa100 | -1.515247815 | 0.78068098 | 0.21931902 |
| erdafitinib | 11522 | BLCa100 | -0.408992802 | 0.94080183 | 0.05919817 |
| erdafitinib | 16438 | BLCa100 | 2.364260459 | 1.34220625 | -0.3422063 |
| erdafitinib | 9073 | BLCa100 | -1.79054228 | 0.74083449 | 0.25916551 |
| erdafitinib | 10966 | BLCa100 | -0.722647971 | 0.89540296 | 0.10459704 |
| cisplatin+gemcitabine | 10354 | BLCa100 | -0.70630044 | 0.86296917 | 0.13703083 |
| cisplatin+gemcitabine | 8066 | BLCa100 | -1.689211751 | 0.67227249 | 0.32772751 |
| cisplatin+gemcitabine | 9196 | BLCa100 | -1.203770413 | 0.76645398 | 0.23354602 |
| cisplatin+gemcitabine | 10552 | BLCa100 | -0.621240808 | 0.87947177 | 0.12052823 |
| untreated | 14859 | BLCa82 | -1.108270271 | 0.79586965 | |
| untreated | 16553 | BLCa82 | -0.615659548 | 0.88660275 | |
| untreated | 14056 | BLCa82 | -1.341780549 | 0.7528598 | |
| untreated | 16952 | BLCa82 | -0.499631402 | 0.90797377 | |
| untreated | 19862 | BLCa82 | 0.346588907 | 1.0638376 | |
| untreated | 23642 | BLCa82 | 1.445802918 | 1.2662999 | |
| untreated | 28319 | BLCa82 | 2.805862158 | 1.51680682 | |
| dmso | 15778 | BLCa82 | -0.8410275 | 0.84509262 | 0.15490738 |
| dmso | 21810 | BLCa82 | 0.913062159 | 1.16817531 | -0.1681753 |
| dmso | 15289 | BLCa82 | -0.983227407 | 0.81890107 | 0.18109893 |
| dmso | 17290 | BLCa82 | -0.401341896 | 0.92607754 | 0.07392246 |
| dmso | 24835 | BLCa82 | 1.792724165 | 1.33019871 | -0.3301987 |
| dmso | 17954 | BLCa82 | -0.20825245 | 0.96164235 | 0.03835765 |
| dmso | 17735 | BLCa82 | -0.271937072 | 0.94991239 | 0.05008761 |
| h2o | 15657 | BLCa82 | -0.389288206 | 0.90817865 | 0.09182135 |
| h2o | 13572 | BLCa82 | -0.902027252 | 0.78723898 | 0.21276102 |
| h2o | 14024 | BLCa82 | -0.790872313 | 0.81345708 | 0.18654292 |
| h2o | 14160 | BLCa82 | -0.757427464 | 0.82134571 | 0.17865429 |
| h2o | 17532 | BLCa82 | 0.071808058 | 1.01693735 | -0.0169374 |
| h2o | 22895 | BLCa82 | 1.390666334 | 1.32801624 | -0.3280162 |
| h2o | 22840 | BLCa82 | 1.377140843 | 1.32482599 | -0.324826 |
| bosutinib | 13189 | BLCa82 | -1.593901858 | 0.70642202 | 0.29357798 |
| bosutinib | 16009 | BLCa82 | -0.77385331 | 0.85746532 | 0.14253468 |
| bosutinib | 13091 | BLCa82 | -1.622399999 | 0.701173 | 0.298827 |
| bosutinib | 16144 | BLCa82 | -0.734595667 | 0.86469612 | 0.13530388 |
| cisplatin_6_um | 18144 | BLCa82 | 0.222309879 | 1.05243619 | -0.0524362 |
| cisplatin_6_um | 16756 | BLCa82 | -0.119024316 | 0.97192575 | 0.02807425 |
| cisplatin_6_um | 13456 | BLCa82 | -0.930553741 | 0.78051044 | 0.21948956 |
| cisplatin_6_um | 16418 | BLCa82 | -0.202144602 | 0.95232019 | 0.04767981 |
| cisplatin_14_um | 18951 | BLCa82 | 0.420765711 | 1.09924594 | -0.0992459 |
| cisplatin_14_um | 18930 | BLCa82 | 0.415601433 | 1.09802784 | -0.0980278 |
| cisplatin_14_um | 17887 | BLCa82 | 0.159108951 | 1.037529 | -0.037529 |
| cisplatin_14_um | 15458 | BLCa82 | -0.43822589 | 0.89663573 | 0.10336427 |
| mmc | 17164 | BLCa82 | -0.437982363 | 0.9193288 | 0.0806712 |
| mmc | 21392 | BLCa82 | 0.791508864 | 1.14578663 | -0.1457866 |
| mmc | 16000 | BLCa82 | -0.776470486 | 0.85698327 | 0.14301673 |
| mmc | 19306 | BLCa82 | 0.184905577 | 1.03405743 | -0.0340574 |

| | | | | | |
|---|---|---|---|---|---|
| crizotinib | 15878 | BLCa82 | -0.811947764 | 0.85044877 | 0.14955123 |
| crizotinib | 13779 | BLCa82 | -1.422331417 | 0.73802328 | 0.26197672 |
| crizotinib | 13057 | BLCa82 | -1.632287109 | 0.69935191 | 0.30064809 |
| crizotinib | 19367 | BLCa82 | 0.202644216 | 1.03732468 | -0.0373247 |
| daunorubicin | 13353 | BLCa82 | -1.546211091 | 0.7152061 | 0.2847939 |
| daunorubicin | 14625 | BLCa82 | -1.176316853 | 0.78333627 | 0.21666373 |
| daunorubicin | 10495 | BLCa82 | -2.377309938 | 0.56212746 | 0.43787254 |
| daunorubicin | 7404 | BLCa82 | -3.27616457 | 0.39656901 | 0.60343099 |
| docetaxel | 12125 | BLCa82 | -1.903310246 | 0.64943263 | 0.35056737 |
| docetaxel | 10550 | BLCa82 | -2.361316084 | 0.56507334 | 0.43492666 |
| docetaxel | 12615 | BLCa82 | -1.760819541 | 0.67567774 | 0.32432226 |
| docetaxel | 17041 | BLCa82 | -0.473750437 | 0.91274074 | 0.08725926 |
| doxorubicin | 3148 | BLCa82 | -4.513798122 | 0.16861146 | 0.83138854 |
| doxorubicin | 3579 | BLCa82 | -4.388464461 | 0.19169644 | 0.80830356 |
| doxorubicin | 2720 | BLCa82 | -4.638259391 | 0.14568716 | 0.85431284 |
| doxorubicin | 3363 | BLCa82 | -4.45127669 | 0.18012717 | 0.81987283 |
| epirubicin | 1759 | BLCa82 | -4.917715651 | 0.0942146 | 0.9057854 |
| epirubicin | 2142 | BLCa82 | -4.806340263 | 0.11472863 | 0.88527137 |
| epirubicin | 2359 | BLCa82 | -4.743237237 | 0.12635147 | 0.87364853 |
| epirubicin | 254 | BLCa82 | -5.355365674 | 0.01360461 | 0.98639539 |
| erlotinib | 16942 | BLCa82 | -0.502539376 | 0.90743816 | 0.09256184 |
| erlotinib | 15457 | BLCa82 | -0.934373451 | 0.8278994 | 0.1721006 |
| erlotinib | 19198 | BLCa82 | 0.153499462 | 1.0282728 | -0.0282728 |
| erlotinib | 13042 | BLCa82 | -1.636649069 | 0.69854848 | 0.30145152 |
| everolimus | 8841 | BLCa82 | -2.858288767 | 0.47353682 | 0.52646318 |
| everolimus | 17780 | BLCa82 | -0.25885119 | 0.95232265 | 0.04767735 |
| everolimus | 12136 | BLCa82 | -1.900111475 | 0.65002181 | 0.34997819 |
| everolimus | 10870 | BLCa82 | -2.268260929 | 0.58221301 | 0.41778699 |
| gemcitabine | 7606 | BLCa82 | -2.369174086 | 0.44118329 | 0.55881671 |
| gemcitabine | 9636 | BLCa82 | -1.869960531 | 0.55893271 | 0.44106729 |
| gemcitabine | 11153 | BLCa82 | -1.496902913 | 0.64692575 | 0.35307425 |
| gemcitabine | 8751 | BLCa82 | -2.087597967 | 0.50759861 | 0.49240139 |
| lapatinib | 11859 | BLCa82 | -1.980662343 | 0.63518528 | 0.36481472 |
| lapatinib | 16095 | BLCa82 | -0.748844737 | 0.8620716 | 0.1379284 |
| lapatinib | 10074 | BLCa82 | -2.499735626 | 0.53957809 | 0.46042191 |
| lapatinib | 13477 | BLCa82 | -1.510152219 | 0.72184772 | 0.27815228 |
| methotrexate | 18972 | BLCa82 | 0.087779259 | 1.01616791 | -0.0161679 |
| methotrexate | 23218 | BLCa82 | 1.322504838 | 1.24358984 | -0.2435898 |
| methotrexate | 26229 | BLCa82 | 2.198095681 | 1.40486338 | -0.4048634 |
| methotrexate | 14854 | BLCa82 | -1.109724258 | 0.79560184 | 0.20439816 |
| olaparib | 22029 | BLCa82 | 0.976746781 | 1.17990527 | -0.1799053 |
| olaparib | 17072 | BLCa82 | -0.464735719 | 0.91440114 | 0.08559886 |
| olaparib | 14388 | BLCa82 | -1.245235826 | 0.7706422 | 0.2293578 |
| olaparib | 16379 | BLCa82 | -0.666258288 | 0.87728306 | 0.12271694 |
| paclitaxel | 12490 | BLCa82 | -1.79716921 | 0.66898256 | 0.33101744 |
| paclitaxel | 12121 | BLCa82 | -1.904473435 | 0.64921839 | 0.35078161 |
| paclitaxel | 12184 | BLCa82 | -1.886153202 | 0.65259276 | 0.34740724 |
| ponatinib | 12247 | BLCa82 | -1.867832968 | 0.65596713 | 0.34403287 |
| ponatinib | 10957 | BLCa82 | -2.242961559 | 0.58687285 | 0.41312715 |
| ponatinib | 13720 | BLCa82 | -1.439488461 | 0.73486315 | 0.26513685 |
| ponatinib | 11037 | BLCa82 | -2.219697771 | 0.59115777 | 0.40884223 |
| rapamycin | 12092 | BLCa82 | -1.912906559 | 0.6476651 | 0.3523349 |
| rapamycin | 13544 | BLCa82 | -1.490668796 | 0.72543633 | 0.27456367 |
| rapamycin | 14341 | BLCa82 | -1.258903302 | 0.76812481 | 0.23187519 |
| rapamycin | 11259 | BLCa82 | -2.155140757 | 0.60304841 | 0.39695159 |
| sunitinib | 26639 | BLCa82 | 2.317322598 | 1.42682358 | -0.4268236 |
| sunitinib | 22074 | BLCa82 | 0.989832662 | 1.18231554 | -0.1823155 |
| sunitinib | 24289 | BLCa82 | 1.633948808 | 1.30095416 | -0.3009542 |
| sunitinib | 20731 | BLCa82 | 0.599291811 | 1.11038251 | -0.1103825 |
| temsirolimus | 18140 | BLCa82 | -0.154164142 | 0.97160478 | 0.02839522 |
| temsirolimus | 15372 | BLCa82 | -0.959091227 | 0.82334667 | 0.17665333 |
| temsirolimus | 13927 | BLCa82 | -1.379293408 | 0.74595037 | 0.25404963 |
| temsirolimus | 16497 | BLCa82 | -0.6319442 | 0.88360331 | 0.11639669 |
| vinblastine | 17751 | BLCa82 | -0.267284314 | 0.95076937 | 0.04923063 |
| vinblastine | 20318 | BLCa82 | 0.479192502 | 1.08826162 | -0.0882616 |
| vinblastine | 16374 | BLCa82 | -0.667712275 | 0.87701525 | 0.12298475 |
| vinblastine | 12442 | BLCa82 | -1.811127484 | 0.66641161 | 0.33358839 |

| | | | | | |
|---|---|---|---|---|---|
| cisplatin+gemcitabine | 6519 | BLCa82 | -2.636486961 | 0.37813225 | 0.62186775 |
| cisplatin+gemcitabine | 6733 | BLCa82 | -2.583860507 | 0.39054524 | 0.60945476 |
| cisplatin+gemcitabine | 7075 | BLCa82 | -2.499756548 | 0.41038283 | 0.58961717 |
| cisplatin+gemcitabine | 8776 | BLCa82 | -2.081450017 | 0.50904872 | 0.49095128 |
| erdafitinib | 16655 | BLCa82 | -0.585998217 | 0.89206602 | 0.10793398 |
| erdafitinib | 17599 | BLCa82 | -0.311485512 | 0.94262803 | 0.05737197 |
| erdafitinib | 21110 | BLCa82 | 0.709504009 | 1.1306823 | -0.1306823 |
| erdafitinib | 13612 | BLCa82 | -1.470894576 | 0.72907851 | 0.27092149 |
| untreated | 17594 | BLCa112 | 0.55713771 | 1.08233766 | |
| untreated | 23007 | BLCa112 | 2.81034025 | 1.41533151 | |
| untreated | 20687 | BLCa112 | 1.844622553 | 1.27261107 | |
| untreated | 18787 | BLCa112 | 1.05373306 | 1.15572796 | |
| untreated | 17262 | BLCa112 | 0.418940177 | 1.06191388 | |
| untreated | 15225 | BLCa112 | -0.428976611 | 0.93660287 | |
| untreated | 16258 | BLCa112 | 0.001017519 | 1.00015038 | |
| untreated | 14390 | BLCa112 | -0.77655173 | 0.88523582 | |
| untreated | 17735 | BLCa112 | 0.615830035 | 1.09101162 | |
| untreated | 19600 | BLCa112 | 1.392150511 | 1.20574163 | |
| dmso | 19618 | BLCa112 | 1.399643149 | 1.20684894 | -0.2068489 |
| dmso | 19201 | BLCa112 | 1.226063718 | 1.18119617 | -0.1811962 |
| dmso | 15214 | BLCa112 | -0.433555445 | 0.93592618 | 0.06407382 |
| dmso | 18049 | BLCa112 | 0.74653493 | 1.11032809 | -0.1103281 |
| dmso | 15535 | BLCa112 | -0.299936746 | 0.95567327 | 0.04432673 |
| dmso | 15364 | BLCa112 | -0.371116801 | 0.94515379 | 0.05484621 |
| dmso | 11892 | BLCa112 | -1.816363285 | 0.73156528 | 0.26843472 |
| dmso | 16401 | BLCa112 | 0.060542359 | 1.00894737 | -0.0089474 |
| dmso | 15026 | BLCa112 | -0.511811879 | 0.9243609 | 0.0756391 |
| h2o | 17070 | BLCa112 | -0.595332254 | 0.92311852 | 0.07688148 |
| h2o | 18937 | BLCa112 | 0.186486258 | 1.02408292 | -0.0240829 |
| h2o | 17337 | BLCa112 | -0.48352425 | 0.93755746 | 0.06244254 |
| h2o | 22120 | BLCa112 | 1.519388414 | 1.19621451 | -0.1962145 |
| h2o | 14574 | BLCa112 | -1.640548647 | 0.7881388 | 0.2118612 |
| h2o | 17424 | BLCa112 | -0.447092429 | 0.94226228 | 0.05773772 |
| h2o | 19888 | BLCa112 | 0.584723754 | 1.07551149 | -0.0755115 |
| h2o | 21572 | BLCa112 | 1.289909815 | 1.16657954 | -0.1665795 |
| h2o | 17503 | BLCa112 | -0.41401066 | 0.94653447 | 0.05346553 |
| bosutinib | 16332 | BLCa112 | 0.031820583 | 1.00470267 | -0.0047027 |
| bosutinib | 19680 | BLCa112 | 1.425451121 | 1.21066302 | -0.210663 |
| bosutinib | 19071 | BLCa112 | 1.171950226 | 1.17319891 | -0.1731989 |
| bosutinib | 14923 | BLCa112 | -0.554686415 | 0.91802461 | 0.08197539 |
| cisplatin_6_um | 15499 | BLCa112 | -1.253198822 | 0.83816133 | 0.16183867 |
| cisplatin_6_um | 14706 | BLCa112 | -1.58527278 | 0.79527715 | 0.20472285 |
| cisplatin_6_um | 18560 | BLCa112 | 0.028615032 | 1.00369536 | -0.0036954 |
| cisplatin_6_um | 15840 | BLCa112 | -1.110402833 | 0.85660207 | 0.14339793 |
| cisplatin_14_um | 20284 | BLCa112 | 0.750551355 | 1.09692654 | -0.0969265 |
| cisplatin_14_um | 16227 | BLCa112 | -0.948344041 | 0.87753042 | 0.12246958 |
| cisplatin_14_um | 18197 | BLCa112 | -0.123393602 | 0.98406489 | 0.01593511 |
| cisplatin_14_um | 18667 | BLCa112 | 0.073421985 | 1.00948175 | -0.0094817 |
| crizotinib | 17791 | BLCa112 | 0.639140462 | 1.0944566 | -0.0944566 |
| crizotinib | 17650 | BLCa112 | 0.580448137 | 1.08578264 | -0.0857826 |
| crizotinib | 14875 | BLCa112 | -0.574666781 | 0.91507177 | 0.08492823 |
| crizotinib | 11154 | BLCa112 | -2.123561414 | 0.68616541 | 0.31383459 |
| daunorubicin | 19964 | BLCa112 | 1.543668288 | 1.22813397 | -0.228134 |
| daunorubicin | 16060 | BLCa112 | -0.081401492 | 0.98796992 | 0.01203008 |
| daunorubicin | 15845 | BLCa112 | -0.170896882 | 0.97474368 | 0.02525632 |
| daunorubicin | 19468 | BLCa112 | 1.337204504 | 1.19762133 | -0.1976213 |
| docetaxel | 15812 | BLCa112 | -0.184633383 | 0.9727136 | 0.0272864 |
| docetaxel | 13394 | BLCa112 | -1.191144328 | 0.82396446 | 0.17603554 |
| docetaxel | 13893 | BLCa112 | -0.983431771 | 0.85466165 | 0.14533835 |
| docetaxel | 13643 | BLCa112 | -1.087496178 | 0.8392823 | 0.1607177 |
| doxorubicin | 1921 | BLCa112 | -5.966868093 | 0.11817498 | 0.88182502 |
| doxorubicin | 1908 | BLCa112 | -5.972279442 | 0.11737526 | 0.88262474 |
| doxorubicin | 2199 | BLCa112 | -5.851148472 | 0.13527683 | 0.86472317 |
| doxorubicin | 1851 | BLCa112 | -5.996006127 | 0.11386876 | 0.88613124 |
| epirubicin | 3172 | BLCa112 | -5.446129801 | 0.19513329 | 0.80486671 |
| epirubicin | 3634 | BLCa112 | -5.253818776 | 0.22355434 | 0.77644566 |
| epirubicin | 3592 | BLCa112 | -5.271301597 | 0.22097061 | 0.77902939 |

| | | | | | |
|---|---|---|---|---|---|
| epirubicin | 2906 | BLCa112 | -5.55685433 | 0.17876965 | 0.82123035 |
| erlotinib | 15653 | BLCa112 | -0.250818346 | 0.96293233 | 0.03706767 |
| erlotinib | 10152 | BLCa112 | -2.540651558 | 0.62452495 | 0.37547505 |
| erlotinib | 18509 | BLCa112 | 0.938013439 | 1.13862611 | -0.1386261 |
| erlotinib | 11893 | BLCa112 | -1.815947027 | 0.73162679 | 0.26837321 |
| everolimus | 9227 | BLCa112 | -2.925689863 | 0.56762133 | 0.43237867 |
| everolimus | 13537 | BLCa112 | -1.131619487 | 0.83276145 | 0.16723855 |
| everolimus | 9723 | BLCa112 | -2.71922608 | 0.59813397 | 0.40186603 |
| everolimus | 11415 | BLCa112 | -2.014918173 | 0.70222146 | 0.29777854 |
| gemcitabine | 14768 | BLCa112 | -1.559309873 | 0.79863001 | 0.20136999 |
| gemcitabine | 12712 | BLCa112 | -2.420273377 | 0.68744479 | 0.31255521 |
| gemcitabine | 11785 | BLCa112 | -2.808460715 | 0.63731411 | 0.36268589 |
| gemcitabine | 10427 | BLCa112 | -3.377132134 | 0.56387562 | 0.43612438 |
| lapatinib | 3648 | BLCa112 | -5.24799117 | 0.22441558 | 0.77558442 |
| lapatinib | 3533 | BLCa112 | -5.295860797 | 0.21734108 | 0.78265892 |
| lapatinib | 2202 | BLCa112 | -5.8498997 | 0.13546138 | 0.86453862 |
| lapatinib | 2999 | BLCa112 | -5.51814237 | 0.18449077 | 0.81550923 |
| methotrexate | 10552 | BLCa112 | -2.374148506 | 0.64913192 | 0.35086808 |
| methotrexate | 11536 | BLCa112 | -1.964551 | 0.70966507 | 0.29033493 |
| methotrexate | 13077 | BLCa112 | -1.323097996 | 0.80446343 | 0.19553657 |
| methotrexate | 16872 | BLCa112 | 0.256599702 | 1.03792208 | -0.0379221 |
| olaparib | 11241 | BLCa112 | -2.087347001 | 0.69151743 | 0.30848257 |
| olaparib | 9448 | BLCa112 | -2.833696928 | 0.58121668 | 0.41878332 |
| olaparib | 12358 | BLCa112 | -1.62238723 | 0.7602324 | 0.2397676 |
| olaparib | 11350 | BLCa112 | -2.041974919 | 0.69822283 | 0.30177717 |
| paclitaxel | 10537 | BLCa112 | -2.380392371 | 0.64820916 | 0.35179084 |
| paclitaxel | 11923 | BLCa112 | -1.803459298 | 0.73347232 | 0.26652768 |
| paclitaxel | 10010 | BLCa112 | -2.599760141 | 0.61578947 | 0.38421053 |
| paclitaxel | 12810 | BLCa112 | -1.434238782 | 0.78803828 | 0.21196172 |
| ponatinib | 17459 | BLCa112 | 0.50094293 | 1.07403281 | -0.0740328 |
| ponatinib | 15301 | BLCa112 | -0.397341031 | 0.9412782 | 0.0587218 |
| ponatinib | 12791 | BLCa112 | -1.442147677 | 0.78686945 | 0.21313055 |
| ponatinib | 20124 | BLCa112 | 1.610269508 | 1.23797676 | -0.2379768 |
| rapamycin | 10692 | BLCa112 | -2.315872438 | 0.65774436 | 0.34225564 |
| rapamycin | 10602 | BLCa112 | -2.353335625 | 0.65220779 | 0.34779221 |
| rapamycin | 8150 | BLCa112 | -3.373999329 | 0.50136705 | 0.49863295 |
| rapamycin | 8311 | BLCa112 | -3.306981851 | 0.51127136 | 0.48872864 |
| sunitinib | 18835 | BLCa112 | 1.073713426 | 1.15868079 | -0.1586808 |
| sunitinib | 16976 | BLCa112 | 0.299890496 | 1.04431989 | -0.0443199 |
| sunitinib | 18640 | BLCa112 | 0.992543188 | 1.14668489 | -0.1466849 |
| sunitinib | 16966 | BLCa112 | 0.295727919 | 1.04370472 | -0.0437047 |
| temsirolimus | 11181 | BLCa112 | -2.112322458 | 0.68782638 | 0.31217362 |
| temsirolimus | 9227 | BLCa112 | -2.925689863 | 0.56762133 | 0.43237867 |
| temsirolimus | 12296 | BLCa112 | -1.648195203 | 0.75641832 | 0.24358168 |
| temsirolimus | 12139 | BLCa112 | -1.713547651 | 0.74676008 | 0.25323992 |
| vinblastine | 7918 | BLCa112 | -3.470571098 | 0.48709501 | 0.51290499 |
| vinblastine | 8766 | BLCa112 | -3.11758463 | 0.53926179 | 0.46073821 |
| vinblastine | 16914 | BLCa112 | 0.274082523 | 1.04050581 | -0.0405058 |
| vinblastine | 7539 | BLCa112 | -3.628332739 | 0.4637799 | 0.5362201 |
| mmc | 6607 | BLCa112 | -4.016284849 | 0.40644566 | 0.59355434 |
| mmc | 8217 | BLCa112 | -3.346110068 | 0.50548872 | 0.49451128 |
| mmc | 10353 | BLCa112 | -2.456983774 | 0.63688995 | 0.36311005 |
| mmc | 8112 | BLCa112 | -3.389817119 | 0.49902939 | 0.50097061 |
| cisplatin+gemcitabine | 14020 | BLCa112 | -1.872539786 | 0.75817936 | 0.24182064 |
| cisplatin+gemcitabine | 11961 | BLCa112 | -2.734759559 | 0.64683191 | 0.35316809 |
| cisplatin+gemcitabine | 13677 | BLCa112 | -2.016173289 | 0.73963046 | 0.26036954 |
| cisplatin+gemcitabine | 15597 | BLCa112 | -1.212160679 | 0.84346102 | 0.15653898 |
| erdafitinib | 12609 | BLCa112 | -1.517906566 | 0.77567327 | 0.22432673 |
| erdafitinib | 13207 | BLCa112 | -1.268984504 | 0.8124607 | 0.1875393 |
| erdafitinib | 16218 | BLCa112 | -0.015632786 | 0.99768968 | 0.00231032 |
| erdafitinib | 16068 | BLCa112 | -0.078071431 | 0.98846206 | 0.01153794 |
| untreated | 149 | BLCa46 | 0.153197398 | 1.09257562 | |
| untreated | 182 | BLCa46 | 0.553634161 | 1.33455545 | |
| untreated | 129 | BLCa46 | -0.089491549 | 0.94592117 | |
| untreated | 78 | BLCa46 | -0.708348365 | 0.57195234 | |
| untreated | 133 | BLCa46 | -0.04095376 | 0.97525206 | |
| untreated | 49 | BLCa46 | -1.060247339 | 0.35930339 | |

| | | | | | |
|---|---|---|---|---|---|
| untreated | 140 | BLCa46 | 0.043987372 | 1.02658112 | |
| untreated | 48 | BLCa46 | -1.072381786 | 0.35197067 | |
| dmso | 51 | BLCa46 | -1.035978444 | 0.37396884 | 0.62603116 |
| dmso | 104 | BLCa46 | -0.392852734 | 0.76260312 | 0.23739688 |
| dmso | 69 | BLCa46 | -0.817558391 | 0.50595784 | 0.49404216 |
| dmso | 137 | BLCa46 | 0.00758403 | 1.00458295 | -0.004583 |
| dmso | 305 | BLCa46 | 2.046171187 | 2.23648029 | -1.2364803 |
| dmso | 130 | BLCa46 | -0.077357102 | 0.9532539 | 0.0467461 |
| dmso | 93 | BLCa46 | -0.526331655 | 0.68194317 | 0.31805683 |
| dmso | 202 | BLCa46 | 0.796323108 | 1.4812099 | -0.4812099 |
| h2o | 183 | BLCa46 | 1.842373063 | 1.94164456 | -0.9416446 |
| h2o | 112 | BLCa46 | 0.368474613 | 1.18832891 | -0.1883289 |
| h2o | 39 | BLCa46 | -1.146942104 | 0.4137931 | 0.5862069 |
| h2o | 127 | BLCa46 | 0.679861609 | 1.34748011 | -0.3474801 |
| h2o | 100 | BLCa46 | 0.119365015 | 1.06100796 | -0.061008 |
| h2o | 90 | BLCa46 | -0.088226316 | 0.95490716 | 0.04509284 |
| h2o | 41 | BLCa46 | -1.105423838 | 0.43501326 | 0.56498674 |
| h2o | 62 | BLCa46 | -0.669482043 | 0.65782493 | 0.34217507 |
| bosutinib | 126 | BLCa46 | -0.125894891 | 0.92392301 | 0.07607699 |
| bosutinib | 263 | BLCa46 | 1.536524398 | 1.92850596 | -0.928506 |
| bosutinib | 53 | BLCa46 | -1.011709549 | 0.38863428 | 0.61136572 |
| bosutinib | 140 | BLCa46 | 0.043987372 | 1.02658112 | -0.0265811 |
| cisplatin_6_um | 158 | BLCa46 | 1.323394736 | 1.67639257 | -0.6763926 |
| cisplatin_6_um | 90 | BLCa46 | -0.088226316 | 0.95490716 | 0.04509284 |
| cisplatin_6_um | 75 | BLCa46 | -0.399613312 | 0.79575597 | 0.20424403 |
| cisplatin_6_um | 46 | BLCa46 | -1.001628172 | 0.48806366 | 0.51193634 |
| cisplatin_14_um | 110 | BLCa46 | 0.326956346 | 1.16710875 | -0.1671088 |
| cisplatin_14_um | 324 | BLCa46 | 4.769410832 | 3.43766578 | -2.4376658 |
| cisplatin_14_um | 255 | BLCa46 | 3.337030647 | 2.70557029 | -1.7055703 |
| cisplatin_14_um | 95 | BLCa46 | 0.01556935 | 1.00795756 | -0.0079576 |
| crizotinib | 74 | BLCa46 | -0.756886155 | 0.54262145 | 0.45737855 |
| crizotinib | 126 | BLCa46 | -0.125894891 | 0.92392301 | 0.07607699 |
| crizotinib | 52 | BLCa46 | -1.023843997 | 0.38130156 | 0.61869844 |
| crizotinib | 86 | BLCa46 | -0.611272786 | 0.63061412 | 0.36938588 |
| daunorubicin | 125 | BLCa46 | -0.138029339 | 0.91659028 | 0.08340972 |
| daunorubicin | 143 | BLCa46 | 0.080390714 | 1.04857929 | -0.0485793 |
| daunorubicin | 82 | BLCa46 | -0.659810576 | 0.60128323 | 0.39871677 |
| daunorubicin | 56 | BLCa46 | -0.975306207 | 0.41063245 | 0.58936755 |
| docetaxel | 115 | BLCa46 | -0.259373812 | 0.84326306 | 0.15673694 |
| docetaxel | 79 | BLCa46 | -0.696213918 | 0.57928506 | 0.42071494 |
| docetaxel | 87 | BLCa46 | -0.599138339 | 0.63794684 | 0.36205316 |
| docetaxel | 124 | BLCa46 | -0.150163786 | 0.90925756 | 0.09074244 |
| doxorubicin | 55 | BLCa46 | -0.987440655 | 0.40329973 | 0.59670027 |
| doxorubicin | 25 | BLCa46 | -1.351474076 | 0.18331806 | 0.81668194 |
| doxorubicin | 41 | BLCa46 | -1.157322918 | 0.30064161 | 0.69935839 |
| doxorubicin | 88 | BLCa46 | -0.587003891 | 0.64527956 | 0.35472044 |
| epirubicin | 24 | BLCa46 | -1.363608523 | 0.17598533 | 0.82401467 |
| epirubicin | 68 | BLCa46 | -0.829692839 | 0.49862511 | 0.50137489 |
| epirubicin | 79 | BLCa46 | -0.696213918 | 0.57928506 | 0.42071494 |
| epirubicin | 71 | BLCa46 | -0.793289497 | 0.52062328 | 0.47937672 |
| erlotinib | 98 | BLCa46 | -0.465659418 | 0.71860678 | 0.28139322 |
| erlotinib | 150 | BLCa46 | 0.165331845 | 1.09990834 | -0.0999083 |
| erlotinib | 40 | BLCa46 | -1.169457365 | 0.29330889 | 0.70669111 |
| erlotinib | 122 | BLCa46 | -0.174432681 | 0.89459212 | 0.10540788 |
| everolimus | 63 | BLCa46 | -0.890365076 | 0.4619615 | 0.5380385 |
| everolimus | 177 | BLCa46 | 0.492961924 | 1.29789184 | -0.2978918 |
| everolimus | 142 | BLCa46 | 0.068256266 | 1.04124656 | -0.0412466 |
| everolimus | 159 | BLCa46 | 0.274541872 | 1.16590284 | -0.1659028 |
| gemcitabine | 25 | BLCa46 | -1.437569968 | 0.26525199 | 0.73474801 |
| gemcitabine | 39 | BLCa46 | -1.146942104 | 0.4137931 | 0.5862069 |
| gemcitabine | 13 | BLCa46 | -1.686679565 | 0.13793103 | 0.86206897 |
| lapatinib | 47 | BLCa46 | -1.084516233 | 0.34463795 | 0.65536205 |
| lapatinib | 92 | BLCa46 | -0.538466102 | 0.67461045 | 0.32538955 |
| lapatinib | 84 | BLCa46 | -0.635541681 | 0.61594867 | 0.38405133 |
| lapatinib | 57 | BLCa46 | -0.96317176 | 0.41796517 | 0.58203483 |
| methotrexate | 119 | BLCa46 | -0.210836023 | 0.87259395 | 0.12740605 |
| methotrexate | 63 | BLCa46 | -0.890365076 | 0.4619615 | 0.5380385 |

| | | | | | |
|---|---|---|---|---|---|
| methotrexate | 237 | BLCa46 | 1.221028766 | 1.73785518 | -0.7378552 |
| methotrexate | 143 | BLCa46 | 0.080390714 | 1.04857929 | -0.0485793 |
| olaparib | 9 | BLCa46 | -1.545625233 | 0.0659945 | 0.9340055 |
| olaparib | 151 | BLCa46 | 0.177466293 | 1.10724106 | -0.1072411 |
| olaparib | 59 | BLCa46 | -0.938902865 | 0.43263061 | 0.56736939 |
| olaparib | 91 | BLCa46 | -0.550600549 | 0.66727773 | 0.33272227 |
| paclitaxel | 130 | BLCa46 | -0.077357102 | 0.9532539 | 0.0467461 |
| paclitaxel | 40 | BLCa46 | -1.169457365 | 0.29330889 | 0.70669111 |
| paclitaxel | 154 | BLCa46 | 0.213869635 | 1.12923923 | -0.1292392 |
| paclitaxel | 64 | BLCa46 | -0.878230628 | 0.46929423 | 0.53070577 |
| ponatinib | 75 | BLCa46 | -0.744751707 | 0.54995417 | 0.45004583 |
| ponatinib | 122 | BLCa46 | -0.174432681 | 0.89459212 | 0.10540788 |
| ponatinib | 32 | BLCa46 | -1.266532944 | 0.23464711 | 0.76535289 |
| ponatinib | 106 | BLCa46 | -0.368583839 | 0.77726856 | 0.22273144 |
| rapamycin | 227 | BLCa46 | 1.099684293 | 1.66452796 | -0.664528 |
| rapamycin | 169 | BLCa46 | 0.395886345 | 1.23923006 | -0.2392301 |
| rapamycin | 50 | BLCa46 | -1.048112891 | 0.36663611 | 0.63336389 |
| rapamycin | 130 | BLCa46 | -0.077357102 | 0.9532539 | 0.0467461 |
| sunitinib | 35 | BLCa46 | -1.230129602 | 0.25664528 | 0.74335472 |
| sunitinib | 126 | BLCa46 | -0.125894891 | 0.92392301 | 0.07607699 |
| sunitinib | 38 | BLCa46 | -1.19372626 | 0.27864345 | 0.72135655 |
| sunitinib | 71 | BLCa46 | -0.793289497 | 0.52062328 | 0.47937672 |
| temsirolimus | 68 | BLCa46 | -0.829692839 | 0.49862511 | 0.50137489 |
| temsirolimus | 43 | BLCa46 | -1.133054023 | 0.31530706 | 0.68469294 |
| temsirolimus | 115 | BLCa46 | -0.259373812 | 0.84326306 | 0.15673694 |
| temsirolimus | 62 | BLCa46 | -0.902499523 | 0.45462878 | 0.54537122 |
| vinblastine | 45 | BLCa46 | -1.108785128 | 0.3299725 | 0.6700275 |
| vinblastine | 42 | BLCa46 | -1.14518847 | 0.30797434 | 0.69202566 |
| vinblastine | 41 | BLCa46 | -1.157322918 | 0.30064161 | 0.69935839 |
| mmc | 29 | BLCa46 | -1.302936286 | 0.21264895 | 0.78735105 |
| mmc | 120 | BLCa46 | -0.198701576 | 0.87992667 | 0.12007333 |
| mmc | 61 | BLCa46 | -0.91463397 | 0.44729606 | 0.55270394 |
| mmc | 56 | BLCa46 | -0.975306207 | 0.41063245 | 0.58936755 |
| erdafitinib | 16 | BLCa46 | -1.460684102 | 0.11732356 | 0.88267644 |
| erdafitinib | 68 | BLCa46 | -0.829692839 | 0.49862511 | 0.50137489 |
| erdafitinib | 90 | BLCa46 | -0.562734997 | 0.659945 | 0.340055 |
| erdafitinib | 11 | BLCa46 | -1.521356339 | 0.08065995 | 0.91934005 |
| cisplatin+gemcitabine | 205 | BLCa46 | 2.299073992 | 2.17506631 | -1.1750663 |
| cisplatin+gemcitabine | 74 | BLCa46 | -0.420372445 | 0.78514589 | 0.21485411 |
| cisplatin+gemcitabine | 22 | BLCa46 | -1.499847367 | 0.23342175 | 0.76657825 |
| cisplatin+gemcitabine | 183 | BLCa46 | 1.842373063 | 1.94164456 | -0.9416446 |
| untreated | 4065 | BLCa81 | -0.802430032 | 0.69585313 | |
| untreated | 4624 | BLCa81 | -0.549969985 | 0.79154363 | |
| untreated | 6215 | BLCa81 | 0.168570147 | 1.06389353 | |
| untreated | 4788 | BLCa81 | -0.475902995 | 0.81961741 | |
| untreated | 4123 | BLCa81 | -0.776235609 | 0.70578166 | |
| untreated | 4918 | BLCa81 | -0.417191356 | 0.84187101 | |
| untreated | 7852 | BLCa81 | 0.907885168 | 1.34411777 | |
| untreated | 5970 | BLCa81 | 0.05792129 | 1.02195404 | |
| dmso | 5113 | BLCa81 | -0.329123898 | 0.87525142 | 0.12474858 |
| dmso | 7199 | BLCa81 | 0.61297209 | 1.2323362 | -0.2323362 |
| dmso | 3872 | BLCa81 | -0.889594234 | 0.66281508 | 0.33718492 |
| dmso | 10745 | BLCa81 | 2.214444944 | 1.83934609 | -0.8393461 |
| dmso | 5484 | BLCa81 | -0.161569914 | 0.93875979 | 0.06124021 |
| dmso | 4220 | BLCa81 | -0.732427694 | 0.72238627 | 0.27761373 |
| dmso | 4967 | BLCa81 | -0.395061585 | 0.85025891 | 0.14974109 |
| dmso | 5134 | BLCa81 | -0.31963971 | 0.87884624 | 0.12115376 |
| h2o | 3808 | BLCa81 | -0.526781371 | 0.85799583 | 0.14200417 |
| h2o | 4204 | BLCa81 | -0.195792997 | 0.94722019 | 0.05277981 |
| h2o | 3704 | BLCa81 | -0.613707611 | 0.83456317 | 0.16543683 |
| h2o | 4432 | BLCa81 | -0.005223933 | 0.99859179 | 0.00140821 |
| h2o | 3785 | BLCa81 | -0.546005443 | 0.85281361 | 0.14718639 |
| h2o | 6705 | BLCa81 | 1.894615903 | 1.51073058 | -0.5107306 |
| h2o | 5750 | BLCa81 | 1.09639899 | 1.29555568 | -0.2955557 |
| h2o | 3118 | BLCa81 | -1.103503538 | 0.70252915 | 0.29747085 |
| cisplatin_6_um | 4820 | BLCa81 | 0.319077808 | 1.08601363 | -0.0860136 |
| cisplatin_6_um | 7031 | BLCa81 | 2.167096231 | 1.58418295 | -0.584183 |

| | | | | | |
|---|---|---|---|---|---|
| cisplatin_6_um | 5854 | BLCa81 | 1.18332523 | 1.31898834 | -0.3189883 |
| cisplatin_6_um | 7880 | BLCa81 | 2.876715246 | 1.77547457 | -0.7754746 |
| cisplatin_14_um | 5017 | BLCa81 | 0.483736166 | 1.1304005 | -0.1304005 |
| cisplatin_14_um | 5044 | BLCa81 | 0.506303555 | 1.13648397 | -0.136484 |
| cisplatin_14_um | 5016 | BLCa81 | 0.482900337 | 1.13017518 | -0.1301752 |
| gemcitabine | 5690 | BLCa81 | 1.046249236 | 1.28203684 | -0.2820368 |
| gemcitabine | 7695 | BLCa81 | 2.722086839 | 1.73379147 | -0.7337915 |
| gemcitabine | 5718 | BLCa81 | 1.069652455 | 1.28834563 | -0.2883456 |
| gemcitabine | 4983 | BLCa81 | 0.455317972 | 1.12273982 | -0.1227398 |
| cisplatin+gemcitabine | 3768 | BLCa81 | -0.56021454 | 0.84898327 | 0.15101673 |
| cisplatin+gemcitabine | 7117 | BLCa81 | 2.238977545 | 1.60355996 | -0.60356 |
| cisplatin+gemcitabine | 4716 | BLCa81 | 0.232151568 | 1.06258097 | -0.062581 |
| cisplatin+gemcitabine | 4633 | BLCa81 | 0.162777742 | 1.04387991 | -0.0438799 |
| mmc | 4110 | BLCa81 | -0.782106773 | 0.7035563 | 0.2964437 |
| mmc | 6354 | BLCa81 | 0.231346438 | 1.08768776 | -0.0876878 |
| mmc | 4483 | BLCa81 | -0.613649532 | 0.76740703 | 0.23259297 |
| mmc | 3680 | BLCa81 | -0.976306808 | 0.62994822 | 0.37005178 |
| vinblastine | 4612 | BLCa81 | -0.555389521 | 0.78948945 | 0.21051055 |
| vinblastine | 6593 | BLCa81 | 0.339285528 | 1.12860016 | -0.1286002 |
| vinblastine | 6080 | BLCa81 | 0.107600369 | 1.04078401 | -0.040784 |
| vinblastine | 6405 | BLCa81 | 0.254379466 | 1.09641803 | -0.096418 |
| erdafitinib | 6498 | BLCa81 | 0.296380869 | 1.11233791 | -0.1123379 |
| erdafitinib | 6366 | BLCa81 | 0.236765974 | 1.08974194 | -0.0897419 |
| erdafitinib | 5600 | BLCa81 | -0.109181067 | 0.95861685 | 0.04138315 |
| erdafitinib | 4895 | BLCa81 | -0.4275788 | 0.83793384 | 0.16206616 |
| erlotinib | 6135 | BLCa81 | 0.132439908 | 1.050199 | -0.050199 |
| erlotinib | 4379 | BLCa81 | -0.660618843 | 0.74960414 | 0.25039586 |
| erlotinib | 4188 | BLCa81 | -0.746879789 | 0.71690846 | 0.28309154 |
| erlotinib | 4923 | BLCa81 | -0.414933216 | 0.84272692 | 0.15727308 |
| lapatinib | 2578 | BLCa81 | -1.474000854 | 0.44130612 | 0.55869388 |
| lapatinib | 2783 | BLCa81 | -1.381417116 | 0.47639834 | 0.52360166 |
| lapatinib | 2399 | BLCa81 | -1.554842264 | 0.41066461 | 0.58933539 |
| lapatinib | 4704 | BLCa81 | -0.513839746 | 0.80523816 | 0.19476184 |
| temsirolimus | 3775 | BLCa81 | -0.933402149 | 0.64621047 | 0.35378953 |
| temsirolimus | 3714 | BLCa81 | -0.960951457 | 0.63576839 | 0.36423161 |
| temsirolimus | 4844 | BLCa81 | -0.450611828 | 0.82920358 | 0.17079642 |
| temsirolimus | 7046 | BLCa81 | 0.543873008 | 1.20614542 | -0.2061454 |
| doxorubicin | 5146 | BLCa81 | -0.314220174 | 0.88090042 | 0.11909958 |
| doxorubicin | 6120 | BLCa81 | 0.125665488 | 1.04763127 | -0.0476313 |
| doxorubicin | 9320 | BLCa81 | 1.570875058 | 1.59541233 | -0.5954123 |
| doxorubicin | 8205 | BLCa81 | 1.067309849 | 1.40454487 | -0.4045449 |
| epirubicin | 3614 | BLCa81 | -1.006114256 | 0.61865023 | 0.38134977 |
| epirubicin | 4163 | BLCa81 | -0.758170489 | 0.71262892 | 0.28737108 |
| epirubicin | 6491 | BLCa81 | 0.293219473 | 1.11113964 | -0.1111396 |
| epirubicin | 4015 | BLCa81 | -0.825011432 | 0.68729405 | 0.31270595 |
| untreated | 7226 | BLCa69 | -0.2785635 | 0.89149343 | |
| untreated | 7006 | BLCa69 | -0.348243967 | 0.86435137 | |
| untreated | 9366 | BLCa69 | 0.399237399 | 1.15551169 | |
| untreated | 6624 | BLCa69 | -0.469234594 | 0.81722287 | |
| untreated | 6638 | BLCa69 | -0.464800383 | 0.8189501 | |
| untreated | 7702 | BLCa69 | -0.12780031 | 0.95021899 | |
| untreated | 9278 | BLCa69 | 0.371365212 | 1.14465486 | |
| untreated | 8257 | BLCa69 | 0.047984503 | 1.01869101 | |
| dmso | 4372 | BLCa69 | -1.182509185 | 0.53938684 | 0.46061316 |
| dmso | 10280 | BLCa69 | 0.688728063 | 1.26827463 | -0.2682746 |
| dmso | 13931 | BLCa69 | 1.845107073 | 1.71870952 | -0.7187095 |
| dmso | 7240 | BLCa69 | -0.274129289 | 0.89322065 | 0.10677935 |
| dmso | 8613 | BLCa69 | 0.160740166 | 1.06261181 | -0.0626118 |
| dmso | 9496 | BLCa69 | 0.44041222 | 1.17155018 | -0.1715502 |
| dmso | 5144 | BLCa69 | -0.937994095 | 0.63463081 | 0.36536919 |
| dmso | 5768 | BLCa69 | -0.740354954 | 0.71161557 | 0.28838443 |
| h2o | 7025 | BLCa69 | 1.199628108 | 1.11342984 | -0.1134298 |
| h2o | 6598 | BLCa69 | 0.483874216 | 1.04575232 | -0.0457523 |
| h2o | 5223 | BLCa69 | -1.820953891 | 0.82782122 | 0.17217878 |
| h2o | 6364 | BLCa69 | 0.091634378 | 1.00866441 | -0.0086644 |
| h2o | 6307 | BLCa69 | -0.003911223 | 0.99963018 | 0.00036982 |
| h2o | 6339 | BLCa69 | 0.049728412 | 1.00470203 | -0.004702 |

| | | | | | |
|---|---|---|---|---|---|
| cisplatin_6_um | 5750 | BLCa69 | -0.937576137 | 0.91134827 | 0.08865173 |
| cisplatin_6_um | 5131 | BLCa69 | -1.975167844 | 0.81323964 | 0.18676036 |
| cisplatin_6_um | 9853 | BLCa69 | 5.940030935 | 1.56165469 | -0.5616547 |
| cisplatin_6_um | 6139 | BLCa69 | -0.285519312 | 0.97300296 | 0.02699704 |
| cisplatin_14_um | 5980 | BLCa69 | -0.552041253 | 0.9478022 | 0.0521978 |
| cisplatin_14_um | 5427 | BLCa69 | -1.479001212 | 0.86015427 | 0.13984573 |
| cisplatin_14_um | 5209 | BLCa69 | -1.844421232 | 0.82560228 | 0.17439772 |
| cisplatin_14_um | 5779 | BLCa69 | -0.888965217 | 0.91594463 | 0.08405537 |
| gemcitabine | 8205 | BLCa69 | 3.177589684 | 1.30045435 | -0.3004544 |
| gemcitabine | 5460 | BLCa69 | -1.423685337 | 0.86538462 | 0.13461538 |
| gemcitabine | 4808 | BLCa69 | -2.51659292 | 0.76204565 | 0.23795435 |
| gemcitabine | 7326 | BLCa69 | 1.704175934 | 1.16113694 | -0.1611369 |
| cisplatin+gemcitabine | 4335 | BLCa69 | -3.309453789 | 0.68707735 | 0.31292265 |
| cisplatin+gemcitabine | 4794 | BLCa69 | -2.540060261 | 0.75982671 | 0.24017329 |
| cisplatin+gemcitabine | 4582 | BLCa69 | -2.895422849 | 0.7262257 | 0.2737743 |
| mmc | 2979 | BLCa69 | -1.623713228 | 0.36752822 | 0.63247178 |
| mmc | 8657 | BLCa69 | 0.17467626 | 1.06804022 | -0.0680402 |
| mmc | 3352 | BLCa69 | -1.505573165 | 0.41354636 | 0.58645364 |
| mmc | 4543 | BLCa69 | -1.128348459 | 0.56048362 | 0.43951638 |
| vinblastine | 6783 | BLCa69 | -0.418874621 | 0.83683918 | 0.16316082 |
| vinblastine | 4310 | BLCa69 | -1.202146407 | 0.53173771 | 0.46826229 |
| vinblastine | 8068 | BLCa69 | -0.011877352 | 0.99537351 | 0.00462649 |
| vinblastine | 6365 | BLCa69 | -0.551267507 | 0.78526926 | 0.21473074 |
| erdafitinib | 16848 | BLCa69 | 2.76900671 | 2.07858861 | -1.0785886 |
| erdafitinib | 20745 | BLCa69 | 4.003301151 | 2.55937327 | -1.5593733 |
| erdafitinib | 5715 | BLCa69 | -0.757141612 | 0.7050768 | 0.2949232 |
| erdafitinib | 5838 | BLCa69 | -0.718183896 | 0.72025168 | 0.27974832 |
| erlotinib | 4870 | BLCa69 | -1.024777948 | 0.6008266 | 0.3991734 |
| erlotinib | 8930 | BLCa69 | 0.261143384 | 1.10172105 | -0.1017211 |
| erlotinib | 11420 | BLCa69 | 1.04979957 | 1.40891987 | -0.4089199 |
| erlotinib | 8089 | BLCa69 | -0.005226035 | 0.99796435 | 0.00203565 |
| lapatinib | 3585 | BLCa69 | -1.431775217 | 0.44229227 | 0.55770773 |
| lapatinib | 3940 | BLCa69 | -1.319336283 | 0.48608969 | 0.51391031 |
| lapatinib | 3358 | BLCa69 | -1.503672789 | 0.4142866 | 0.5857134 |
| temsirolimus | 5209 | BLCa69 | -0.917406684 | 0.64265005 | 0.35734995 |
| temsirolimus | 5682 | BLCa69 | -0.767593682 | 0.70100549 | 0.29899451 |
| temsirolimus | 4833 | BLCa69 | -1.036496935 | 0.5962618 | 0.4037382 |
| temsirolimus | 5431 | BLCa69 | -0.847092759 | 0.67003886 | 0.32996114 |
| doxorubicin | 2422 | BLCa69 | -1.8001315 | 0.29880945 | 0.70119055 |
| doxorubicin | 2938 | BLCa69 | -1.636699133 | 0.36246993 | 0.63753007 |
| doxorubicin | 2195 | BLCa69 | -1.872029072 | 0.27080378 | 0.72919622 |
| doxorubicin | 3070 | BLCa69 | -1.594890854 | 0.37875517 | 0.62124483 |
| epirubicin | 1630 | BLCa69 | -2.050981178 | 0.20109802 | 0.79890198 |
| epirubicin | 4410 | BLCa69 | -1.170473468 | 0.54407501 | 0.45592499 |
| epirubicin | 2155 | BLCa69 | -1.884698247 | 0.26586885 | 0.73413115 |
| epirubicin | 1979 | BLCa69 | -1.94044262 | 0.2441552 | 0.7558448 |
| crizotinib | 8565 | BLCa69 | 0.145537156 | 1.0566899 | -0.0566899 |
| crizotinib | 13263 | BLCa69 | 1.633531839 | 1.63629634 | -0.6362963 |
| crizotinib | 5612 | BLCa69 | -0.789764739 | 0.69236938 | 0.30763062 |
| crizotinib | 6760 | BLCa69 | -0.426159397 | 0.8340016 | 0.1659984 |
| untreated | 3532 | BLCa22 | -0.169374501 | 0.95809033 | |
| untreated | 4472 | BLCa22 | 0.861124083 | 1.21307473 | |
| untreated | 4387 | BLCa22 | 0.767940701 | 1.19001763 | |
| untreated | 4102 | BLCa22 | 0.4555023 | 1.11270853 | |
| untreated | 3402 | BLCa22 | -0.311890263 | 0.92282653 | |
| untreated | 4378 | BLCa22 | 0.758074225 | 1.18757629 | |
| untreated | 2251 | BLCa22 | -1.573702892 | 0.61060627 | |
| untreated | 1824 | BLCa22 | -2.041812355 | 0.49477824 | |
| dmso | 3108 | BLCa22 | -0.63419514 | 0.84307609 | 0.15692391 |
| dmso | 3937 | BLCa22 | 0.27461691 | 1.06795063 | -0.0679506 |
| dmso | 2918 | BLCa22 | -0.842487407 | 0.79153669 | 0.20846331 |
| dmso | 5817 | BLCa22 | 2.33561408 | 1.57791944 | -0.5779194 |
| dmso | 3429 | BLCa22 | -0.282290836 | 0.93015055 | 0.06984945 |
| dmso | 3335 | BLCa22 | -0.385340694 | 0.90465211 | 0.09534789 |
| dmso | 3381 | BLCa22 | -0.33491204 | 0.91713007 | 0.08286993 |
| dmso | 3567 | BLCa22 | -0.131004873 | 0.96758443 | 0.03241557 |
| h2o | 7170 | BLCa22 | 1.792969841 | 1.56905654 | -0.5690565 |

| | | | | | |
|---|---|---|---|---|---|
| h2o | 5405 | BLCa22 | 0.575994686 | 1.18281041 | -0.1828104 |
| h2o | 2572 | BLCa22 | -1.377371101 | 0.56284706 | 0.43715294 |
| h2o | 3511 | BLCa22 | -0.729926529 | 0.76833438 | 0.23166562 |
| h2o | 4835 | BLCa22 | 0.182977213 | 1.05807369 | -0.0580737 |
| h2o | 3541 | BLCa22 | -0.709241399 | 0.77489947 | 0.22510053 |
| h2o | 4072 | BLCa22 | -0.343114596 | 0.89110157 | 0.10889843 |
| h2o | 5451 | BLCa22 | 0.607711885 | 1.19287688 | -0.1928769 |
| bosutinib | 4134 | BLCa22 | 0.490583103 | 1.12138885 | -0.1213889 |
| bosutinib | 2537 | BLCa22 | -1.260168216 | 0.68818663 | 0.31181337 |
| bosutinib | 3152 | BLCa22 | -0.585959036 | 0.85501153 | 0.14498847 |
| bosutinib | 2299 | BLCa22 | -1.521081688 | 0.62362675 | 0.37637325 |
| cisplatin_6_um | 3680 | BLCa22 | -0.613400296 | 0.80531772 | 0.19468228 |
| cisplatin_6_um | 5171 | BLCa22 | 0.414650671 | 1.1316027 | -0.1316027 |
| cisplatin_6_um | 4282 | BLCa22 | -0.198318685 | 0.9370572 | 0.0629428 |
| cisplatin_6_um | 3228 | BLCa22 | -0.925056256 | 0.7064037 | 0.2935963 |
| cisplatin_14_um | 5975 | BLCa22 | 0.969012158 | 1.30754712 | -0.3075471 |
| cisplatin_14_um | 2875 | BLCa22 | -1.168451287 | 0.62915447 | 0.37084553 |
| cisplatin_14_um | 4269 | BLCa22 | -0.207282241 | 0.93421233 | 0.06578767 |
| cisplatin_14_um | 4536 | BLCa22 | -0.023184583 | 0.99264163 | 0.00735837 |
| crizotinib | 2372 | BLCa22 | -1.441053606 | 0.64342873 | 0.35657127 |
| crizotinib | 2888 | BLCa22 | -0.87537566 | 0.78339889 | 0.21660111 |
| crizotinib | 2443 | BLCa22 | -1.363218075 | 0.66268819 | 0.33731181 |
| crizotinib | 3783 | BLCa22 | 0.105790546 | 1.02617659 | -0.0261766 |
| daunorubicin | 4802 | BLCa22 | 1.222894863 | 1.30259053 | -0.3025905 |
| daunorubicin | 2500 | BLCa22 | -1.300730394 | 0.67815001 | 0.32184999 |
| daunorubicin | 3330 | BLCa22 | -0.39082207 | 0.90329581 | 0.09670419 |
| daunorubicin | 2544 | BLCa22 | -1.252494291 | 0.69008545 | 0.30991455 |
| docetaxel | 2492 | BLCa22 | -1.309500595 | 0.67597993 | 0.32402007 |
| docetaxel | 2289 | BLCa22 | -1.532044439 | 0.62091415 | 0.37908585 |
| docetaxel | 2148 | BLCa22 | -1.686619226 | 0.58266649 | 0.41733351 |
| docetaxel | 1901 | BLCa22 | -1.957399174 | 0.51566527 | 0.48433473 |
| doxorubicin | 2203 | BLCa22 | -1.626324096 | 0.59758579 | 0.40241421 |
| doxorubicin | 1699 | BLCa22 | -2.178846742 | 0.46087074 | 0.53912926 |
| doxorubicin | 1652 | BLCa22 | -2.230371671 | 0.44812152 | 0.55187848 |
| doxorubicin | 1667 | BLCa22 | -2.213927545 | 0.45219042 | 0.54780958 |
| epirubicin | 3132 | BLCa22 | -0.607884538 | 0.84958633 | 0.15041367 |
| epirubicin | 3152 | BLCa22 | -0.585959036 | 0.85501153 | 0.14498847 |
| epirubicin | 2141 | BLCa22 | -1.694293152 | 0.58076767 | 0.41923233 |
| epirubicin | 1547 | BLCa22 | -2.345480555 | 0.41963922 | 0.58036078 |
| erlotinib | 2736 | BLCa22 | -1.042009473 | 0.74216737 | 0.25783263 |
| erlotinib | 3637 | BLCa22 | -0.054265617 | 0.98657263 | 0.01342737 |
| erlotinib | 4015 | BLCa22 | 0.360126367 | 1.08910891 | -0.0891089 |
| erlotinib | 4908 | BLCa22 | 1.339100023 | 1.33134409 | -0.3313441 |
| everolimus | 3757 | BLCa22 | 0.077287394 | 1.01912383 | -0.0191238 |
| everolimus | 2830 | BLCa22 | -0.938959615 | 0.76766581 | 0.23233419 |
| everolimus | 2837 | BLCa22 | -0.931285689 | 0.76956463 | 0.23043537 |
| everolimus | 3386 | BLCa22 | -0.329430665 | 0.91848637 | 0.08151363 |
| gemcitabine | 4268 | BLCa22 | -0.207971746 | 0.93399349 | 0.06600651 |
| gemcitabine | 2595 | BLCa22 | -1.361512502 | 0.5678803 | 0.4321197 |
| gemcitabine | 3468 | BLCa22 | -0.759575215 | 0.75892442 | 0.24107558 |
| gemcitabine | 2393 | BLCa22 | -1.500792378 | 0.52367536 | 0.47632464 |
| lapatinib | 3175 | BLCa22 | -0.560744709 | 0.86125051 | 0.13874949 |
| lapatinib | 2459 | BLCa22 | -1.345677673 | 0.66702835 | 0.33297165 |
| lapatinib | 1779 | BLCa22 | -2.091144735 | 0.48257154 | 0.51742846 |
| lapatinib | 1972 | BLCa22 | -1.879563642 | 0.53492473 | 0.46507527 |
| methotrexate | 4455 | BLCa22 | 0.842487407 | 1.20846331 | -0.2084633 |
| methotrexate | 4486 | BLCa22 | 0.876471935 | 1.21687237 | -0.2168724 |
| methotrexate | 3413 | BLCa22 | -0.299831237 | 0.92581039 | 0.07418961 |
| methotrexate | 4995 | BLCa22 | 1.434475956 | 1.35494371 | -0.3549437 |
| olaparib | 4154 | BLCa22 | 0.512508605 | 1.12681405 | -0.1268141 |
| olaparib | 3141 | BLCa22 | -0.598018062 | 0.85202767 | 0.14797233 |
| olaparib | 4361 | BLCa22 | 0.739437548 | 1.18296487 | -0.1829649 |
| olaparib | 3100 | BLCa22 | -0.64296534 | 0.84090601 | 0.15909399 |
| paclitaxel | 3437 | BLCa22 | -0.273520635 | 0.93232063 | 0.06767937 |
| paclitaxel | 3772 | BLCa22 | 0.09373152 | 1.02319273 | -0.0231927 |
| paclitaxel | 1801 | BLCa22 | -2.067026683 | 0.48853926 | 0.51146074 |
| paclitaxel | 2565 | BLCa22 | -1.229472514 | 0.69578191 | 0.30421809 |

| | | | | | |
|---|---|---|---|---|---|
| ponatinib | 2781 | BLCa22 | -0.992677094 | 0.75437407 | 0.24562593 |
| ponatinib | 3824 | BLCa22 | 0.150737825 | 1.03729825 | -0.0372983 |
| ponatinib | 3429 | BLCa22 | -0.282290836 | 0.93015055 | 0.06984945 |
| ponatinib | 2592 | BLCa22 | -1.199873086 | 0.70310593 | 0.29689407 |
| rapamycin | 2902 | BLCa22 | -0.860027808 | 0.78719653 | 0.21280347 |
| rapamycin | 3558 | BLCa22 | -0.140871349 | 0.96514309 | 0.03485691 |
| rapamycin | 4324 | BLCa22 | 0.69887537 | 1.17292825 | -0.1729283 |
| rapamycin | 3449 | BLCa22 | -0.260365334 | 0.93557575 | 0.06442425 |
| sunitinib | 3284 | BLCa22 | -0.441250724 | 0.89081785 | 0.10918215 |
| sunitinib | 3300 | BLCa22 | -0.423710322 | 0.89515801 | 0.10484199 |
| sunitinib | 3313 | BLCa22 | -0.409458746 | 0.89868439 | 0.10131561 |
| temsirolimus | 4219 | BLCa22 | 0.583766486 | 1.14444595 | -0.144446 |
| temsirolimus | 3550 | BLCa22 | -0.14964155 | 0.96297301 | 0.03702699 |
| temsirolimus | 3530 | BLCa22 | -0.171567052 | 0.95754781 | 0.04245219 |
| temsirolimus | 2526 | BLCa22 | -1.272227242 | 0.68520277 | 0.31479723 |
| vinblastine | 3742 | BLCa22 | 0.060843268 | 1.01505493 | -0.0150549 |
| vinblastine | 3105 | BLCa22 | -0.637483965 | 0.84226231 | 0.15773769 |
| vinblastine | 2165 | BLCa22 | -1.66798255 | 0.58727791 | 0.41272209 |
| vinblastine | 2933 | BLCa22 | -0.82604328 | 0.79560559 | 0.20439441 |
| cisplatin+gemcitabine | 3142 | BLCa22 | -0.984353629 | 0.68758377 | 0.31241623 |
| cisplatin+gemcitabine | 3065 | BLCa22 | -1.037445463 | 0.67073338 | 0.32926662 |
| cisplatin+gemcitabine | 2000 | BLCa22 | -1.771767582 | 0.43767268 | 0.56232732 |
| cisplatin+gemcitabine | 2270 | BLCa22 | -1.585601411 | 0.49675849 | 0.50324151 |
| untreated | 70068 | BLCa98 | 1.996085685 | 1.2743923 | |
| untreated | 51100 | BLCa98 | -0.513558916 | 0.92940353 | |
| untreated | 42659 | BLCa98 | -1.630382518 | 0.77587916 | |
| untreated | 51031 | BLCa98 | -0.522688264 | 0.92814856 | |
| untreated | 44644 | BLCa98 | -1.367748369 | 0.81198221 | |
| untreated | 39153 | BLCa98 | -2.094259256 | 0.71211226 | |
| dmso | 48688 | BLCa98 | -0.832689176 | 0.88553423 | 0.11446577 |
| dmso | 45632 | BLCa98 | -1.237026687 | 0.82995189 | 0.17004811 |
| dmso | 52662 | BLCa98 | -0.306891641 | 0.95781308 | 0.04218692 |
| dmso | 66022 | BLCa98 | 1.460761873 | 1.20080391 | -0.2008039 |
| dmso | 60362 | BLCa98 | 0.711890699 | 1.09786019 | -0.0978602 |
| dmso | 56523 | BLCa98 | 0.203954932 | 1.0280367 | -0.0280367 |
| h2o | 42095 | BLCa98 | -1.514516835 | 0.85120366 | 0.14879634 |
| h2o | 54513 | BLCa98 | 1.041339665 | 1.10230823 | -0.1023082 |
| h2o | 52300 | BLCa98 | 0.585862903 | 1.05755912 | -0.0575591 |
| h2o | 50709 | BLCa98 | 0.258405366 | 1.02538749 | -0.0253875 |
| h2o | 52225 | BLCa98 | 0.570426501 | 1.05604255 | -0.0560425 |
| h2o | 44879 | BLCa98 | -0.9415176 | 0.90749896 | 0.09250104 |
| bosutinib | 60685 | BLCa98 | 0.754626633 | 1.10373489 | -0.1037349 |
| bosutinib | 44465 | BLCa98 | -1.391431751 | 0.80872657 | 0.19127343 |
| bosutinib | 35852 | BLCa98 | -2.531012568 | 0.65207388 | 0.34792612 |
| bosutinib | 53355 | BLCa98 | -0.215201231 | 0.97041732 | 0.02958268 |
| cisplatin_6_um | 48527 | BLCa98 | -0.190691017 | 0.98126523 | 0.01873477 |
| cisplatin_6_um | 38003 | BLCa98 | -2.356726917 | 0.76845926 | 0.23154074 |
| cisplatin_6_um | 45199 | BLCa98 | -0.875655619 | 0.91396969 | 0.08603031 |
| cisplatin_6_um | 43556 | BLCa98 | -1.213815728 | 0.88074656 | 0.11925344 |
| cisplatin_14_um | 60947 | BLCa98 | 2.365577121 | 1.23241024 | -0.2324102 |
| cisplatin_14_um | 42693 | BLCa98 | -1.391437258 | 0.86329582 | 0.13670418 |
| cisplatin_14_um | 59796 | BLCa98 | 2.128679808 | 1.20913585 | -0.2091359 |
| cisplatin_14_um | 59169 | BLCa98 | 1.999631489 | 1.19645728 | -0.1964573 |
| crizotinib | 42026 | BLCa98 | -1.714134365 | 0.7643662 | 0.2356338 |
| crizotinib | 36287 | BLCa98 | -2.473457981 | 0.65998563 | 0.34001437 |
| crizotinib | 51906 | BLCa98 | -0.406917544 | 0.944063 | 0.055937 |
| crizotinib | 51255 | BLCa98 | -0.49305096 | 0.93222266 | 0.06777734 |
| daunorubicin | 57108 | BLCa98 | 0.281355928 | 1.03867665 | -0.0386766 |
| daunorubicin | 57474 | BLCa98 | 0.329781167 | 1.04533343 | -0.0453334 |
| daunorubicin | 68065 | BLCa98 | 1.731069967 | 1.23796186 | -0.2379619 |
| daunorubicin | 51401 | BLCa98 | -0.473733788 | 0.9348781 | 0.0651219 |
| docetaxel | 40237 | BLCa98 | -1.950835872 | 0.73182798 | 0.26817202 |
| docetaxel | 43298 | BLCa98 | -1.545836814 | 0.78750125 | 0.21249875 |
| docetaxel | 51715 | BLCa98 | -0.432188638 | 0.94058911 | 0.05941089 |
| docetaxel | 54258 | BLCa98 | -0.095725847 | 0.98684103 | 0.01315897 |
| doxorubicin | 33465 | BLCa98 | -2.846835094 | 0.60865928 | 0.39134072 |
| doxorubicin | 21421 | BLCa98 | -4.440369445 | 0.38960378 | 0.61039622 |

| | | | | | |
|---|---|---|---|---|---|
| doxorubicin | 20200 | BLCa98 | -4.601919216 | 0.36739631 | 0.63260369 |
| doxorubicin | 19533 | BLCa98 | -4.690169582 | 0.35526495 | 0.64473505 |
| epirubicin | 32351 | BLCa98 | -2.994227759 | 0.58839792 | 0.41160208 |
| epirubicin | 25091 | BLCa98 | -3.954793966 | 0.4563535 | 0.5436465 |
| epirubicin | 31351 | BLCa98 | -3.126537154 | 0.57020998 | 0.42979002 |
| epirubicin | 35414 | BLCa98 | -2.588964083 | 0.64410756 | 0.35589244 |
| erlotinib | 47295 | BLCa98 | -1.016996163 | 0.86019843 | 0.13980157 |
| erlotinib | 57257 | BLCa98 | 0.301070028 | 1.04138665 | -0.0413866 |
| erlotinib | 65078 | BLCa98 | 1.335861805 | 1.1836345 | -0.1836345 |
| erlotinib | 47081 | BLCa98 | -1.045310374 | 0.85630621 | 0.14369379 |
| everolimus | 53340 | BLCa98 | -0.217185872 | 0.9701445 | 0.0298555 |
| everolimus | 59075 | BLCa98 | 0.541608508 | 1.07445232 | -0.0744523 |
| everolimus | 59393 | BLCa98 | 0.583682895 | 1.08023608 | -0.0802361 |
| everolimus | 44698 | BLCa98 | -1.360603662 | 0.81296436 | 0.18703564 |
| gemcitabine | 40160 | BLCa98 | -1.912776001 | 0.81207599 | 0.18792401 |
| gemcitabine | 38716 | BLCa98 | -2.20997819 | 0.78287684 | 0.21712316 |
| gemcitabine | 37780 | BLCa98 | -2.402624485 | 0.76394997 | 0.23605003 |
| gemcitabine | 45108 | BLCa98 | -0.89438512 | 0.91212958 | 0.08787042 |
| lapatinib | 36015 | BLCa98 | -2.509446137 | 0.65503851 | 0.34496149 |
| lapatinib | 22404 | BLCa98 | -4.31030931 | 0.40748252 | 0.59251748 |
| lapatinib | 29452 | BLCa98 | -3.377792695 | 0.53567109 | 0.46432891 |
| lapatinib | 40378 | BLCa98 | -1.932180247 | 0.73439248 | 0.26560752 |
| methotrexate | 63280 | BLCa98 | 1.097969513 | 1.15093259 | -0.1509326 |
| methotrexate | 48198 | BLCa98 | -0.89752078 | 0.87662214 | 0.12337786 |
| methotrexate | 62429 | BLCa98 | 0.985374218 | 1.13545465 | -0.1354547 |
| methotrexate | 43815 | BLCa98 | -1.477432857 | 0.79690441 | 0.20309559 |
| olaparib | 55652 | BLCa98 | 0.088713449 | 1.01219501 | -0.012195 |
| olaparib | 56221 | BLCa98 | 0.163997495 | 1.02254395 | -0.0225439 |
| olaparib | 39615 | BLCa98 | -2.033132315 | 0.72051508 | 0.27948492 |
| olaparib | 50448 | BLCa98 | -0.599824641 | 0.91754499 | 0.08245501 |
| paclitaxel | 56752 | BLCa98 | 0.234253784 | 1.03220174 | -0.0322017 |
| paclitaxel | 62375 | BLCa98 | 0.978229511 | 1.1344725 | -0.1344725 |
| paclitaxel | 51175 | BLCa98 | -0.503635711 | 0.93076762 | 0.06923238 |
| paclitaxel | 42073 | BLCa98 | -1.707915823 | 0.76522103 | 0.23477897 |
| ponatinib | 49979 | BLCa98 | -0.661877748 | 0.90901485 | 0.09098515 |
| ponatinib | 48613 | BLCa98 | -0.842612381 | 0.88417013 | 0.11582987 |
| ponatinib | 39016 | BLCa98 | -2.112385643 | 0.70962051 | 0.29037949 |
| ponatinib | 59754 | BLCa98 | 0.631446587 | 1.08680192 | -0.0868019 |
| rapamycin | 64553 | BLCa98 | 1.266399372 | 1.17408583 | -0.1740858 |
| rapamycin | 55256 | BLCa98 | 0.036318929 | 1.00499259 | -0.0049926 |
| rapamycin | 37596 | BLCa98 | -2.300264984 | 0.68379364 | 0.31620636 |
| rapamycin | 39769 | BLCa98 | -2.012756669 | 0.72331602 | 0.27668398 |
| sunitinib | 48635 | BLCa98 | -0.839701574 | 0.88457026 | 0.11542974 |
| sunitinib | 82528 | BLCa98 | 3.644660744 | 1.50101398 | -0.501014 |
| sunitinib | 44848 | BLCa98 | -1.340757252 | 0.81569255 | 0.18430745 |
| sunitinib | 56565 | BLCa98 | 0.209511927 | 1.0288006 | -0.0288006 |
| temsirolimus | 38351 | BLCa98 | -2.20037139 | 0.69752553 | 0.30247447 |
| temsirolimus | 50744 | BLCa98 | -0.560661061 | 0.92292862 | 0.07707138 |
| temsirolimus | 62033 | BLCa98 | 0.932979698 | 1.12825223 | -0.1282522 |
| temsirolimus | 57586 | BLCa98 | 0.344599819 | 1.04737048 | -0.0473705 |
| vinblastine | 55381 | BLCa98 | 0.052857603 | 1.00726608 | -0.0072661 |
| vinblastine | 61576 | BLCa98 | 0.872514304 | 1.11994034 | -0.1199403 |
| vinblastine | 65701 | BLCa98 | 1.418290558 | 1.19496558 | -0.1949656 |
| vinblastine | 38259 | BLCa98 | -2.212543855 | 0.69585224 | 0.30414776 |
| mmc | 52889 | BLCa98 | -0.276857409 | 0.96194174 | 0.03805826 |
| mmc | 63248 | BLCa98 | 1.093735612 | 1.15035057 | -0.1503506 |
| mmc | 70051 | BLCa98 | 1.993836425 | 1.2740831 | -0.2740831 |
| mmc | 70750 | BLCa98 | 2.086320692 | 1.28679647 | -0.2867965 |
| cisplatin+gemcitabine | 33401 | BLCa98 | -3.303904531 | 0.67540215 | 0.32459785 |
| cisplatin+gemcitabine | 38196 | BLCa98 | -2.31700391 | 0.77236192 | 0.22763808 |
| cisplatin+gemcitabine | 35242 | BLCa98 | -2.924992322 | 0.71262904 | 0.28737096 |
| erdafitinib | 51887 | BLCa98 | -0.409431422 | 0.94371743 | 0.05628257 |
| erdafitinib | 33407 | BLCa98 | -2.854509038 | 0.60760438 | 0.39239562 |
| erdafitinib | 36098 | BLCa98 | -2.498464457 | 0.65654811 | 0.34345189 |
| erdafitinib | 41909 | BLCa98 | -1.729614564 | 0.76223821 | 0.23776179 |
| untreated | 4178 | BLCa40 | 1.265314694 | 1.2327211 | |
| untreated | 5387 | BLCa40 | 3.204795474 | 1.58943719 | |

| | | | | | |
|---|---|---|---|---|---|
| untreated | 4113 | BLCa40 | 1.161041534 | 1.21354282 | |
| untreated | 3091 | BLCa40 | -0.478453385 | 0.91200118 | |
| untreated | 2754 | BLCa40 | -1.019069616 | 0.81256915 | |
| untreated | 3244 | BLCa40 | -0.233010408 | 0.95714391 | |
| untreated | 3074 | BLCa40 | -0.505724827 | 0.90698532 | |
| dmso | 3932 | BLCa40 | 0.870680888 | 1.16013867 | -0.1601387 |
| dmso | 3769 | BLCa40 | 0.609195886 | 1.11204544 | -0.1120454 |
| dmso | 4013 | BLCa40 | 1.000621287 | 1.18403777 | -0.1840378 |
| dmso | 3622 | BLCa40 | 0.373378124 | 1.06867301 | -0.068673 |
| dmso | 2962 | BLCa40 | -0.685395503 | 0.87393966 | 0.12606034 |
| dmso | 3742 | BLCa40 | 0.565882419 | 1.10407907 | -0.1040791 |
| dmso | 2735 | BLCa40 | -1.049549463 | 0.80696319 | 0.19303681 |
| dmso | 2339 | BLCa40 | -1.684813639 | 0.69012318 | 0.30987682 |
| h2o | 2673 | BLCa40 | -1.038558138 | 0.8446832 | 0.1553168 |
| h2o | 3437 | BLCa40 | 0.575802834 | 1.08611155 | -0.0861116 |
| h2o | 2504 | BLCa40 | -1.395661547 | 0.79127824 | 0.20872176 |
| h2o | 3531 | BLCa40 | 0.774428398 | 1.11581608 | -0.1158161 |
| h2o | 3925 | BLCa40 | 1.606965339 | 1.24032233 | -0.2403223 |
| h2o | 2854 | BLCa40 | -0.656098275 | 0.90188023 | 0.09811977 |
| h2o | 3148 | BLCa40 | -0.034865126 | 0.99478591 | 0.00521409 |
| h2o | 3244 | BLCa40 | 0.167986515 | 1.02512245 | -0.0251225 |
| bosutinib | 2596 | BLCa40 | -1.272533605 | 0.76595117 | 0.23404883 |
| bosutinib | 3116 | BLCa40 | -0.438348324 | 0.91937744 | 0.08062256 |
| bosutinib | 2392 | BLCa40 | -1.599790908 | 0.70576086 | 0.29423914 |
| bosutinib | 2600 | BLCa40 | -1.266116795 | 0.76713137 | 0.23286863 |
| cisplatin_6_um | 2653 | BLCa40 | -1.080818897 | 0.83836309 | 0.16163691 |
| cisplatin_6_um | 2723 | BLCa40 | -0.932906242 | 0.86048349 | 0.13951651 |
| cisplatin_6_um | 3850 | BLCa40 | 1.448487495 | 1.2166219 | -0.2166219 |
| cisplatin_6_um | 1934 | BLCa40 | -2.600093162 | 0.611155 | 0.388845 |
| cisplatin_14_um | 2965 | BLCa40 | -0.421551065 | 0.93695687 | 0.06304313 |
| cisplatin_14_um | 3013 | BLCa40 | -0.320125245 | 0.95212514 | 0.04787486 |
| cisplatin_14_um | 2463 | BLCa40 | -1.482296102 | 0.77832201 | 0.22167799 |
| cisplatin_14_um | 3981 | BLCa40 | 1.725295463 | 1.25801864 | -0.2580186 |
| crizotinib | 1868 | BLCa40 | -2.440393 | 0.55115439 | 0.44884561 |
| crizotinib | 1920 | BLCa40 | -2.356974471 | 0.56649701 | 0.43350299 |
| crizotinib | 2060 | BLCa40 | -2.132386126 | 0.60780409 | 0.39219591 |
| crizotinib | 1946 | BLCa40 | -2.315265207 | 0.57416833 | 0.42583167 |
| daunorubicin | 1199 | BLCa40 | -3.513604449 | 0.35376558 | 0.64623442 |
| daunorubicin | 1265 | BLCa40 | -3.407727086 | 0.37323892 | 0.62676108 |
| daunorubicin | 1879 | BLCa40 | -2.422746773 | 0.55439994 | 0.44560006 |
| daunorubicin | 3418 | BLCa40 | 0.046120821 | 1.0084827 | -0.0084827 |
| docetaxel | 3099 | BLCa40 | -0.465619765 | 0.91436158 | 0.08563842 |
| docetaxel | 3204 | BLCa40 | -0.297178507 | 0.94534189 | 0.05465811 |
| docetaxel | 3016 | BLCa40 | -0.59876857 | 0.88987239 | 0.11012761 |
| docetaxel | 3079 | BLCa40 | -0.497703815 | 0.90846057 | 0.09153943 |
| doxorubicin | 311 | BLCa40 | -4.938136237 | 0.09176071 | 0.90823929 |
| doxorubicin | 382 | BLCa40 | -4.824237862 | 0.1127093 | 0.8872907 |
| doxorubicin | 251 | BLCa40 | -5.034388385 | 0.07405768 | 0.92594232 |
| doxorubicin | 353 | BLCa40 | -4.870759734 | 0.10415284 | 0.89584716 |
| epirubicin | 137 | BLCa40 | -5.217267466 | 0.04042192 | 0.95957808 |
| epirubicin | 174 | BLCa40 | -5.157911975 | 0.05133879 | 0.94866121 |
| epirubicin | 241 | BLCa40 | -5.05043041 | 0.07110718 | 0.92889282 |
| epirubicin | 178 | BLCa40 | -5.151495165 | 0.05251899 | 0.94748101 |
| erlotinib | 2622 | BLCa40 | -1.230824341 | 0.77362248 | 0.22637752 |
| erlotinib | 5701 | BLCa40 | 3.708515048 | 1.68208306 | -0.6820831 |
| erlotinib | 3392 | BLCa40 | 0.004411557 | 1.00081139 | -0.0008114 |
| erlotinib | 2772 | BLCa40 | -0.990193971 | 0.81788006 | 0.18211994 |
| everolimus | 3679 | BLCa40 | 0.464817664 | 1.08549089 | -0.0854909 |
| everolimus | 3400 | BLCa40 | 0.017245176 | 1.00317179 | -0.0031718 |
| everolimus | 3140 | BLCa40 | -0.399847464 | 0.92645866 | 0.07354134 |
| everolimus | 3277 | BLCa40 | -0.180071727 | 0.96688058 | 0.03311942 |
| gemcitabine | 1697 | BLCa40 | -3.100883149 | 0.53626165 | 0.46373835 |
| gemcitabine | 1281 | BLCa40 | -3.979906925 | 0.40480329 | 0.59519671 |
| gemcitabine | 1500 | BLCa40 | -3.51715162 | 0.47400853 | 0.52599147 |
| gemcitabine | 1893 | BLCa40 | -2.686727717 | 0.59819877 | 0.40180123 |
| lapatinib | 1333 | BLCa40 | -3.298641318 | 0.39330235 | 0.60669765 |
| lapatinib | 1434 | BLCa40 | -3.136616869 | 0.42310246 | 0.57689754 |

| | | | | | |
|---|---|---|---|---|---|
| lapatinib | 1611 | BLCa40 | -2.852673033 | 0.4753264 | 0.5246736 |
| lapatinib | 1348 | BLCa40 | -3.274578281 | 0.39772811 | 0.60227189 |
| methotrexate | 3391 | BLCa40 | 0.002807354 | 1.00051634 | -0.0005163 |
| methotrexate | 4170 | BLCa40 | 1.252481074 | 1.2303607 | -0.2303607 |
| methotrexate | 2290 | BLCa40 | -1.763419559 | 0.67566571 | 0.32433429 |
| methotrexate | 4029 | BLCa40 | 1.026288527 | 1.18875857 | -0.1887586 |
| olaparib | 2612 | BLCa40 | -1.246866366 | 0.77067198 | 0.22932802 |
| olaparib | 3299 | BLCa40 | -0.144779272 | 0.97337169 | 0.02662831 |
| olaparib | 3595 | BLCa40 | 0.330064657 | 1.06070665 | -0.0607066 |
| olaparib | 2968 | BLCa40 | -0.675770288 | 0.87570997 | 0.12429003 |
| paclitaxel | 2398 | BLCa40 | -1.590165693 | 0.70753116 | 0.29246884 |
| paclitaxel | 2563 | BLCa40 | -1.325472287 | 0.7562145 | 0.2437855 |
| paclitaxel | 2841 | BLCa40 | -0.879504001 | 0.83823855 | 0.16176145 |
| paclitaxel | 3100 | BLCa40 | -0.464015563 | 0.91465663 | 0.08534337 |
| ponatinib | 2990 | BLCa40 | -0.640477834 | 0.88220108 | 0.11779892 |
| ponatinib | 3020 | BLCa40 | -0.59235176 | 0.89105259 | 0.10894741 |
| ponatinib | 3004 | BLCa40 | -0.618019 | 0.88633178 | 0.11366822 |
| rapamycin | 3215 | BLCa40 | -0.279532279 | 0.94858745 | 0.05141255 |
| rapamycin | 3396 | BLCa40 | 0.010828367 | 1.00199159 | -0.0019916 |
| rapamycin | 4118 | BLCa40 | 1.169062546 | 1.21501807 | -0.2150181 |
| rapamycin | 2662 | BLCa40 | -1.166656243 | 0.7854245 | 0.2145755 |
| sunitinib | 3935 | BLCa40 | 0.875493495 | 1.16102383 | -0.1610238 |
| sunitinib | 3772 | BLCa40 | 0.614008493 | 1.11293059 | -0.1129306 |
| sunitinib | 3076 | BLCa40 | -0.502516422 | 0.90757542 | 0.09242458 |
| sunitinib | 2314 | BLCa40 | -1.7249187 | 0.68274692 | 0.31725308 |
| temsirolimus | 3064 | BLCa40 | -0.521766852 | 0.90403482 | 0.09596518 |
| temsirolimus | 3004 | BLCa40 | -0.618019 | 0.88633178 | 0.11366822 |
| temsirolimus | 2433 | BLCa40 | -1.534018607 | 0.71785793 | 0.28214207 |
| temsirolimus | 4454 | BLCa40 | 1.708074574 | 1.31415505 | -0.314155 |
| vinblastine | 2783 | BLCa40 | -0.972547744 | 0.82112562 | 0.17887438 |
| vinblastine | 3569 | BLCa40 | 0.288355393 | 1.05303533 | -0.0530353 |
| vinblastine | 4198 | BLCa40 | 1.297398743 | 1.23862211 | -0.2386221 |
| vinblastine | 3316 | BLCa40 | -0.117507831 | 0.97838755 | 0.02161245 |
| mmc | 1803 | BLCa40 | -2.54466616 | 0.5319761 | 0.4680239 |
| mmc | 1601 | BLCa40 | -2.868715058 | 0.47237589 | 0.52762411 |
| mmc | 1330 | BLCa40 | -3.303453926 | 0.3924172 | 0.6075828 |
| mmc | 1878 | BLCa40 | -2.424350975 | 0.55410489 | 0.44589511 |
| erdafitinib | 2953 | BLCa40 | -0.699833325 | 0.87128421 | 0.12871579 |
| erdafitinib | 3514 | BLCa40 | 0.200124257 | 1.03680755 | -0.0368076 |
| erdafitinib | 3306 | BLCa40 | -0.133549855 | 0.97543704 | 0.02456296 |
| erdafitinib | 3482 | BLCa40 | 0.148789779 | 1.02736594 | -0.0273659 |
| cisplatin+gemcitabine | 1555 | BLCa40 | -3.400934534 | 0.49138884 | 0.50861116 |
| cisplatin+gemcitabine | 1321 | BLCa40 | -3.895385408 | 0.41744351 | 0.58255649 |
| cisplatin+gemcitabine | 1202 | BLCa40 | -4.14683692 | 0.37983884 | 0.62016116 |
| cisplatin+gemcitabine | 1451 | BLCa40 | -3.620690478 | 0.45852425 | 0.54147575 |
| untreated | 10362 | BLCa86 | -1.659229478 | 0.82864843 | |
| untreated | 12026 | BLCa86 | -0.370687988 | 0.96171839 | |
| untreated | 10666 | BLCa86 | -1.423822859 | 0.85295929 | |
| untreated | 12667 | BLCa86 | 0.125679257 | 1.01297912 | |
| untreated | 11199 | BLCa86 | -1.011086913 | 0.89558326 | |
| untreated | 14363 | BLCa86 | 1.439000391 | 1.14860812 | |
| untreated | 13195 | BLCa86 | 0.534543384 | 1.05520324 | |
| untreated | 15207 | BLCa86 | 2.092563503 | 1.21610275 | |
| untreated | 14024 | BLCa86 | 1.176491037 | 1.12149832 | |
| untreated | 13236 | BLCa86 | 0.566292303 | 1.05848201 | |
| dmso | 10400 | BLCa86 | -1.62980365 | 0.83168729 | 0.16831271 |
| dmso | 11416 | BLCa86 | -0.843049952 | 0.91293674 | 0.08706326 |
| dmso | 11077 | BLCa86 | -1.105559306 | 0.88582693 | 0.11417307 |
| dmso | 11858 | BLCa86 | -0.500781119 | 0.94828345 | 0.05171655 |
| dmso | 12472 | BLCa86 | -0.025321699 | 0.99738498 | 0.00261502 |
| dmso | 13203 | BLCa86 | 0.540738295 | 1.055843 | -0.055843 |
| dmso | 13207 | BLCa86 | 0.54383575 | 1.05616288 | -0.0561629 |
| dmso | 13251 | BLCa86 | 0.577907761 | 1.05968156 | -0.0596816 |
| dmso | 13620 | BLCa86 | 0.863648031 | 1.08919046 | -0.0891905 |
| dmso | 14543 | BLCa86 | 1.578385889 | 1.16300271 | -0.1630027 |
| h2o | 9874 | BLCa86 | -1.154840577 | 0.74171449 | 0.25828551 |
| h2o | 11238 | BLCa86 | -0.69671978 | 0.84417536 | 0.15582464 |

| | | | | | |
|---|---|---|---|---|---|
| h2o | 11825 | BLCa86 | -0.499566622 | 0.88826958 | 0.11173042 |
| h2o | 11708 | BLCa86 | -0.538862908 | 0.87948078 | 0.12051922 |
| h2o | 9436 | BLCa86 | -1.301949748 | 0.70881284 | 0.29118716 |
| h2o | 16491 | BLCa86 | 1.067582671 | 1.23876987 | -0.2387699 |
| h2o | 14924 | BLCa86 | 0.541281141 | 1.12106006 | -0.1210601 |
| h2o | 15318 | BLCa86 | 0.673612221 | 1.15065653 | -0.1506565 |
| h2o | 18496 | BLCa86 | 1.740993373 | 1.38938133 | -0.3893813 |
| h2o | 13814 | BLCa86 | 0.168470228 | 1.03767916 | -0.0376792 |
| bosutinib | 9974 | BLCa86 | -1.959682662 | 0.79762009 | 0.20237991 |
| bosutinib | 10570 | BLCa86 | -1.498161791 | 0.84528217 | 0.15471783 |
| bosutinib | 10645 | BLCa86 | -1.440084501 | 0.85127992 | 0.14872008 |
| bosutinib | 10023 | BLCa86 | -1.921738832 | 0.80153862 | 0.19846138 |
| bosutinib | 11029 | BLCa86 | -1.142728772 | 0.88198837 | 0.11801163 |
| cisplatin_6_um | 10380 | BLCa86 | -0.984892539 | 0.77972417 | 0.22027583 |
| cisplatin_6_um | 10941 | BLCa86 | -0.796471889 | 0.82186533 | 0.17813467 |
| cisplatin_6_um | 10615 | BLCa86 | -0.905964103 | 0.79737688 | 0.20262312 |
| cisplatin_6_um | 11284 | BLCa86 | -0.681269959 | 0.84763078 | 0.15236922 |
| cisplatin_6_um | 11365 | BLCa86 | -0.654064838 | 0.85371533 | 0.14628467 |
| cisplatin_14_um | 9780 | BLCa86 | -1.186411951 | 0.73465341 | 0.26534659 |
| cisplatin_14_um | 9774 | BLCa86 | -1.188427146 | 0.7342027 | 0.2657973 |
| cisplatin_14_um | 11058 | BLCa86 | -0.757175604 | 0.83065413 | 0.16934587 |
| cisplatin_14_um | 10807 | BLCa86 | -0.841477891 | 0.81179953 | 0.18820047 |
| cisplatin_14_um | 10723 | BLCa86 | -0.869690609 | 0.80548962 | 0.19451038 |
| mmc | 9095 | BLCa86 | -2.640348509 | 0.72732653 | 0.27267347 |
| mmc | 8335 | BLCa86 | -3.228865055 | 0.66654938 | 0.33345062 |
| mmc | 10057 | BLCa86 | -1.89541046 | 0.8042576 | 0.1957424 |
| mmc | 9325 | BLCa86 | -2.462244817 | 0.74571961 | 0.25428039 |
| mmc | 8679 | BLCa86 | -2.962483881 | 0.69405903 | 0.30594097 |
| crizotinib | 10107 | BLCa86 | -1.856692266 | 0.8082561 | 0.1917439 |
| crizotinib | 11105 | BLCa86 | -1.083877118 | 0.88806609 | 0.11193391 |
| crizotinib | 11411 | BLCa86 | -0.846921771 | 0.91253689 | 0.08746311 |
| crizotinib | 9927 | BLCa86 | -1.996077764 | 0.79386151 | 0.20613849 |
| crizotinib | 11112 | BLCa86 | -1.07845657 | 0.88862588 | 0.11137412 |
| daunorubicin | 8535 | BLCa86 | -3.07399228 | 0.68254336 | 0.31745664 |
| daunorubicin | 9944 | BLCa86 | -1.982913578 | 0.795221 | 0.204779 |
| daunorubicin | 9398 | BLCa86 | -2.405716254 | 0.75155741 | 0.24844259 |
| daunorubicin | 9239 | BLCa86 | -2.528840111 | 0.7388422 | 0.2611578 |
| daunorubicin | 9034 | BLCa86 | -2.687584705 | 0.72244836 | 0.27755164 |
| docetaxel | 11566 | BLCa86 | -0.726895371 | 0.92493223 | 0.07506777 |
| docetaxel | 13471 | BLCa86 | 0.748267814 | 1.07727494 | -0.0772749 |
| docetaxel | 12154 | BLCa86 | -0.271569411 | 0.97195455 | 0.02804545 |
| docetaxel | 13739 | BLCa86 | 0.955797332 | 1.09870689 | -0.0987069 |
| docetaxel | 14113 | BLCa86 | 1.245409422 | 1.12861564 | -0.1286156 |
| doxorubicin | 11541 | BLCa86 | -0.746254468 | 0.92293298 | 0.07706702 |
| doxorubicin | 13597 | BLCa86 | 0.845837662 | 1.08735116 | -0.0873512 |
| doxorubicin | 9397 | BLCa86 | -2.406490618 | 0.75147744 | 0.24852256 |
| doxorubicin | 10942 | BLCa86 | -1.210098429 | 0.87503099 | 0.12496901 |
| doxorubicin | 9858 | BLCa86 | -2.049508871 | 0.78834358 | 0.21165642 |
| epirubicin | 10313 | BLCa86 | -1.697173308 | 0.8247299 | 0.1752701 |
| epirubicin | 10944 | BLCa86 | -1.208549702 | 0.87519093 | 0.12480907 |
| epirubicin | 11110 | BLCa86 | -1.080005298 | 0.88846594 | 0.11153406 |
| epirubicin | 11323 | BLCa86 | -0.915065793 | 0.90549953 | 0.09450047 |
| erlotinib | 9871 | BLCa86 | -2.039442141 | 0.78938319 | 0.21061681 |
| erlotinib | 10573 | BLCa86 | -1.4958387 | 0.84552208 | 0.15447792 |
| erlotinib | 10749 | BLCa86 | -1.359550658 | 0.85959679 | 0.14040321 |
| erlotinib | 11072 | BLCa86 | -1.109431125 | 0.88542708 | 0.11457292 |
| erlotinib | 9539 | BLCa86 | -2.296530948 | 0.76283317 | 0.23716683 |
| everolimus | 10809 | BLCa86 | -1.313088825 | 0.86439499 | 0.13560501 |
| everolimus | 11340 | BLCa86 | -0.901901607 | 0.90685902 | 0.09314098 |
| everolimus | 10433 | BLCa86 | -1.604249642 | 0.83432629 | 0.16567371 |
| everolimus | 11584 | BLCa86 | -0.712956821 | 0.92637168 | 0.07362832 |
| everolimus | 12823 | BLCa86 | 0.246480022 | 1.02545443 | -0.0254544 |
| gemcitabine | 9369 | BLCa86 | -1.324452749 | 0.70377993 | 0.29622007 |
| gemcitabine | 10442 | BLCa86 | -0.964068867 | 0.78438148 | 0.21561852 |
| gemcitabine | 9751 | BLCa86 | -1.196152056 | 0.73247499 | 0.26752501 |
| gemcitabine | 10310 | BLCa86 | -1.008403137 | 0.77446591 | 0.22553409 |
| gemcitabine | 11236 | BLCa86 | -0.697391512 | 0.84402512 | 0.15597488 |

| | | | | | |
|---|---|---|---|---|---|
| lapatinib | 9688 | BLCa86 | -2.18115073 | 0.77474869 | 0.22525131 |
| lapatinib | 8885 | BLCa86 | -2.802964923 | 0.71053284 | 0.28946716 |
| lapatinib | 11053 | BLCa86 | -1.124144039 | 0.88390765 | 0.11609235 |
| lapatinib | 10663 | BLCa86 | -1.426145951 | 0.85271938 | 0.14728062 |
| lapatinib | 10872 | BLCa86 | -1.264303901 | 0.86943309 | 0.13056691 |
| methotrexate | 11967 | BLCa86 | -0.416375456 | 0.95700017 | 0.04299983 |
| methotrexate | 12239 | BLCa86 | -0.205748482 | 0.97875199 | 0.02124801 |
| methotrexate | 12134 | BLCa86 | -0.287056689 | 0.97035515 | 0.02964485 |
| methotrexate | 13319 | BLCa86 | 0.630564504 | 1.06511952 | -0.0651195 |
| methotrexate | 11675 | BLCa86 | -0.642489708 | 0.93364895 | 0.06635105 |
| olaparib | 11384 | BLCa86 | -0.867829596 | 0.9103777 | 0.0896223 |
| olaparib | 11999 | BLCa86 | -0.391595812 | 0.95955921 | 0.04044079 |
| olaparib | 13827 | BLCa86 | 1.023941354 | 1.10574424 | -0.1057442 |
| olaparib | 10830 | BLCa86 | -1.296827184 | 0.86607436 | 0.13392564 |
| olaparib | 15475 | BLCa86 | 2.300093022 | 1.23753469 | -0.2375347 |
| paclitaxel | 14809 | BLCa86 | 1.78436668 | 1.18427471 | -0.1842747 |
| paclitaxel | 21126 | BLCa86 | 6.676023286 | 1.68944477 | -0.6894448 |
| paclitaxel | 17909 | BLCa86 | 4.184894696 | 1.4321815 | -0.4321815 |
| paclitaxel | 15449 | BLCa86 | 2.279959561 | 1.23545547 | -0.2354555 |
| paclitaxel | 15872 | BLCa86 | 2.60751548 | 1.26928275 | -0.2692827 |
| ponatinib | 11375 | BLCa86 | -0.874798871 | 0.90965797 | 0.09034203 |
| ponatinib | 12603 | BLCa86 | 0.076119969 | 1.00786104 | -0.007861 |
| ponatinib | 14715 | BLCa86 | 1.711576476 | 1.17675754 | -0.1767575 |
| ponatinib | 13309 | BLCa86 | 0.622820866 | 1.06431982 | -0.0643198 |
| ponatinib | 17339 | BLCa86 | 3.743507287 | 1.38659864 | -0.3865986 |
| rapamycin | 9990 | BLCa86 | -1.94729284 | 0.79889961 | 0.20110039 |
| rapamycin | 11232 | BLCa86 | -0.985532905 | 0.89822227 | 0.10177773 |
| rapamycin | 11734 | BLCa86 | -0.596802239 | 0.93836717 | 0.06163283 |
| rapamycin | 11866 | BLCa86 | -0.494586208 | 0.9489232 | 0.0510768 |
| rapamycin | 13319 | BLCa86 | 0.630564504 | 1.06511952 | -0.0651195 |
| sunitinib | 14077 | BLCa86 | 1.217532323 | 1.12573672 | -0.1257367 |
| sunitinib | 13709 | BLCa86 | 0.932566416 | 1.09630779 | -0.0963078 |
| sunitinib | 15384 | BLCa86 | 2.229625909 | 1.23025742 | -0.2302574 |
| sunitinib | 16564 | BLCa86 | 3.143375283 | 1.32462194 | -0.3246219 |
| sunitinib | 15497 | BLCa86 | 2.317129027 | 1.23929403 | -0.239294 |
| temsirolimus | 14010 | BLCa86 | 1.165649943 | 1.12037874 | -0.1203787 |
| temsirolimus | 13175 | BLCa86 | 0.519056106 | 1.05360384 | -0.0536038 |
| temsirolimus | 13099 | BLCa86 | 0.460204452 | 1.04752613 | -0.0475261 |
| temsirolimus | 11277 | BLCa86 | -0.950686531 | 0.90182092 | 0.09817908 |
| temsirolimus | 11170 | BLCa86 | -1.033543466 | 0.89326413 | 0.10673587 |
| vinblastine | 13512 | BLCa86 | 0.780016733 | 1.08055371 | -0.0805537 |
| vinblastine | 13662 | BLCa86 | 0.896171314 | 1.0925492 | -0.0925492 |
| vinblastine | 12794 | BLCa86 | 0.224023469 | 1.0231353 | -0.0231353 |
| vinblastine | 12302 | BLCa86 | -0.156963558 | 0.98379009 | 0.01620991 |
| vinblastine | 12235 | BLCa86 | -0.208845937 | 0.97843211 | 0.02156789 |
| cisplatin+gemcitabine | 7190 | BLCa86 | -2.05630408 | 0.54009795 | 0.45990205 |
| cisplatin+gemcitabine | 7591 | BLCa86 | -1.921621939 | 0.57022025 | 0.42977975 |
| cisplatin+gemcitabine | 7450 | BLCa86 | -1.968979001 | 0.55962862 | 0.44037138 |
| cisplatin+gemcitabine | 7804 | BLCa86 | -1.850082548 | 0.58622037 | 0.41377963 |
| cisplatin+gemcitabine | 6735 | BLCa86 | -2.209122967 | 0.50591929 | 0.49408071 |
| erdafitinib | 14285 | BLCa86 | 1.378600009 | 1.14237047 | -0.1423705 |
| erdafitinib | 16663 | BLCa86 | 3.220037307 | 1.33253897 | -0.332539 |
| erdafitinib | 14387 | BLCa86 | 1.457585124 | 1.1505274 | -0.1505274 |
| erdafitinib | 16114 | BLCa86 | 2.794911539 | 1.28863547 | -0.2886355 |
| erdafitinib | 14991 | BLCa86 | 1.925300906 | 1.19882924 | -0.1988292 |
| untreated | 26475 | BLCa61 | 1.178610599 | 1.18269403 | |
| untreated | 21211 | BLCa61 | -0.338433869 | 0.94754006 | |
| untreated | 21525 | BLCa61 | -0.247941475 | 0.9615671 | |
| untreated | 26385 | BLCa61 | 1.152673288 | 1.17867354 | |
| untreated | 23074 | BLCa61 | 0.198468457 | 1.03076419 | |
| untreated | 20450 | BLCa61 | -0.557748238 | 0.91354458 | |
| dmso | 18976 | BLCa61 | -0.982543745 | 0.8476979 | 0.1523021 |
| dmso | 18758 | BLCa61 | -1.045369674 | 0.83795938 | 0.16204062 |
| dmso | 20938 | BLCa61 | -0.417110377 | 0.93534457 | 0.06465543 |
| dmso | 27607 | BLCa61 | 1.504844326 | 1.23326285 | -0.2332629 |
| dmso | 24624 | BLCa61 | 0.645166581 | 1.10000596 | -0.100006 |
| dmso | 23409 | BLCa61 | 0.29501289 | 1.04572935 | -0.0457293 |

| | | | | | |
|---|---|---|---|---|---|
| h2o | 23431 | BLCa61 | -0.028600692 | 0.9955458 | 0.0044542 |
| h2o | 21054 | BLCa61 | -0.67709525 | 0.89455086 | 0.10544914 |
| h2o | 18502 | BLCa61 | -1.373333411 | 0.78612045 | 0.21387955 |
| h2o | 22962 | BLCa61 | -0.156553552 | 0.97561874 | 0.02438126 |
| h2o | 26694 | BLCa61 | 0.861612913 | 1.13418546 | -0.1341855 |
| h2o | 28572 | BLCa61 | 1.373969992 | 1.21397868 | -0.2139787 |
| cisplatin_6_um | 21446 | BLCa61 | -0.570149576 | 0.91120632 | 0.08879368 |
| cisplatin_6_um | 25242 | BLCa61 | 0.465477407 | 1.0724923 | -0.0724923 |
| cisplatin_6_um | 21023 | BLCa61 | -0.685552688 | 0.89323372 | 0.10676628 |
| gemcitabine | 17564 | BLCa61 | -1.62923913 | 0.74626633 | 0.25373367 |
| gemcitabine | 17314 | BLCa61 | -1.697444279 | 0.73564423 | 0.26435577 |
| gemcitabine | 16733 | BLCa61 | -1.855953046 | 0.71095847 | 0.28904153 |
| crizotinib | 17463 | BLCa61 | -1.418578752 | 0.780109 | 0.219891 |
| crizotinib | 16430 | BLCa61 | -1.716281438 | 0.73396271 | 0.26603729 |
| crizotinib | 16907 | BLCa61 | -1.578813692 | 0.75527131 | 0.24472869 |
| doxorubicin | 12564 | BLCa61 | -2.830433017 | 0.56126035 | 0.43873965 |
| doxorubicin | 11414 | BLCa61 | -3.161854206 | 0.50988743 | 0.49011257 |
| doxorubicin | 14351 | BLCa61 | -2.315433309 | 0.6410894 | 0.3589106 |
| erlotinib | 23534 | BLCa61 | 0.331036932 | 1.05131336 | -0.0513134 |
| erlotinib | 24989 | BLCa61 | 0.750356784 | 1.11631128 | -0.1163113 |
| erlotinib | 24684 | BLCa61 | 0.662458121 | 1.10268628 | -0.1026863 |
| epirubicin | 21840 | BLCa61 | -0.157160888 | 0.97563881 | 0.02436119 |
| epirubicin | 16688 | BLCa61 | -1.641927814 | 0.74548812 | 0.25451188 |
| epirubicin | 22652 | BLCa61 | 0.07685129 | 1.01191256 | -0.0119126 |
| cisplatin+gemcitabine | 27021 | BLCa61 | 0.950825248 | 1.14807917 | -0.1480792 |
| cisplatin+gemcitabine | 30268 | BLCa61 | 1.836673723 | 1.28603902 | -0.286039 |
| cisplatin+gemcitabine | 13649 | BLCa61 | -2.697331764 | 0.57992423 | 0.42007577 |
| untreated | 158562 | BLCa60 | -0.635322232 | 0.8991732 | |
| untreated | 135825 | BLCa60 | -1.447770016 | 0.77023625 | |
| untreated | 117353 | BLCa60 | -2.107819076 | 0.66548525 | |
| untreated | 143794 | BLCa60 | -1.163018449 | 0.81542684 | |
| untreated | 169002 | BLCa60 | -0.262275882 | 0.95837634 | |
| untreated | 194293 | BLCa60 | 0.641432474 | 1.10179651 | |
| untreated | 154007 | BLCa60 | -0.798083356 | 0.87334271 | |
| dmso | 194219 | BLCa60 | 0.638788276 | 1.10137687 | -0.1013769 |
| dmso | 148311 | BLCa60 | -1.001615157 | 0.84104184 | 0.15895816 |
| dmso | 221931 | BLCa60 | 1.629004795 | 1.25852605 | -0.258526 |
| dmso | 154678 | BLCa60 | -0.774106909 | 0.87714782 | 0.12285218 |
| dmso | 197154 | BLCa60 | 0.743662897 | 1.11802066 | -0.1180207 |
| dmso | 149266 | BLCa60 | -0.967490707 | 0.84645745 | 0.15354255 |
| dmso | 154770 | BLCa60 | -0.770819527 | 0.87766953 | 0.12233047 |
| dmso | 190407 | BLCa60 | 0.502576333 | 1.07975978 | -0.0797598 |
| h2o | 148319 | BLCa60 | -1.719721228 | 0.65569922 | 0.34430078 |
| h2o | 205429 | BLCa60 | -0.458648635 | 0.90817519 | 0.09182481 |
| h2o | 256125 | BLCa60 | 0.660793427 | 1.13229568 | -0.1322957 |
| h2o | 205294 | BLCa60 | -0.461629633 | 0.90757837 | 0.09242163 |
| h2o | 216464 | BLCa60 | -0.214979644 | 0.9569595 | 0.0430405 |
| h2o | 304312 | BLCa60 | 1.724833087 | 1.34532421 | -0.3453242 |
| h2o | 229593 | BLCa60 | 0.074927939 | 1.01500112 | -0.0150011 |
| h2o | 244062 | BLCa60 | 0.394424688 | 1.07896671 | -0.0789667 |
| bosutinib | 278000 | BLCa60 | 3.632485237 | 1.57648206 | -0.5764821 |
| bosutinib | 245557 | BLCa60 | 2.473218691 | 1.39250434 | -0.3925043 |
| bosutinib | 230043 | BLCa60 | 1.918866097 | 1.30452757 | -0.3045276 |
| bosutinib | 183641 | BLCa60 | 0.260810853 | 1.04139116 | -0.0413912 |
| cisplatin_6_um | 244440 | BLCa60 | 0.402771482 | 1.0806378 | -0.0806378 |
| cisplatin_6_um | 224046 | BLCa60 | -0.04755796 | 0.99047855 | 0.00952145 |
| cisplatin_6_um | 242551 | BLCa60 | 0.361059591 | 1.07228677 | -0.0722868 |
| cisplatin_6_um | 158522 | BLCa60 | -1.494424018 | 0.70080537 | 0.29919463 |
| cisplatin_14_um | 217465 | BLCa60 | -0.192876096 | 0.96138479 | 0.03861521 |
| cisplatin_14_um | 226058 | BLCa60 | -0.003130048 | 0.99937334 | 0.00062666 |
| cisplatin_14_um | 128349 | BLCa60 | -2.160688128 | 0.56741442 | 0.43258558 |
| cisplatin_14_um | 191729 | BLCa60 | -0.761164736 | 0.84760925 | 0.15239075 |
| crizotinib | 265284 | BLCa60 | 3.178111923 | 1.50437219 | -0.5043722 |
| crizotinib | 191096 | BLCa60 | 0.527195963 | 1.08366697 | -0.083667 |
| crizotinib | 262373 | BLCa60 | 3.074094881 | 1.48786449 | -0.4878645 |
| crizotinib | 241737 | BLCa60 | 2.336720888 | 1.37084189 | -0.3708419 |
| daunorubicin | 156266 | BLCa60 | -0.717363844 | 0.88615304 | 0.11384696 |

| | | | | | |
|---|---|---|---|---|---|
| daunorubicin | 176406 | BLCa60 | 0.002286874 | 1.00036293 | -0.0003629 |
| daunorubicin | 233387 | BLCa60 | 2.038355273 | 1.32349072 | -0.3234907 |
| daunorubicin | 137740 | BLCa60 | -1.379342453 | 0.78109583 | 0.21890417 |
| docetaxel | 188716 | BLCa60 | 0.442152829 | 1.07017046 | -0.0701705 |
| docetaxel | 214384 | BLCa60 | 1.359332304 | 1.21572853 | -0.2157285 |
| docetaxel | 262672 | BLCa60 | 3.084778871 | 1.48956006 | -0.4895601 |
| docetaxel | 241369 | BLCa60 | 2.323571362 | 1.36875503 | -0.368755 |
| doxorubicin | 33963 | BLCa60 | -5.087544665 | 0.19259734 | 0.80740266 |
| doxorubicin | 33900 | BLCa60 | -5.089795806 | 0.19224008 | 0.80775992 |
| doxorubicin | 23403 | BLCa60 | -5.464878904 | 0.1327137 | 0.8672863 |
| doxorubicin | 46586 | BLCa60 | -4.636494465 | 0.26417983 | 0.73582017 |
| epirubicin | 63042 | BLCa60 | -4.048481943 | 0.3574985 | 0.6425015 |
| epirubicin | 50406 | BLCa60 | -4.499996663 | 0.28584228 | 0.71415772 |
| epirubicin | 44857 | BLCa60 | -4.698275801 | 0.25437502 | 0.74562498 |
| epirubicin | 51654 | BLCa60 | -4.455402616 | 0.29291944 | 0.70708056 |
| erlotinib | 234092 | BLCa60 | 2.063546621 | 1.32748863 | -0.3274886 |
| erlotinib | 226480 | BLCa60 | 1.791551524 | 1.28432251 | -0.2843225 |
| erlotinib | 269038 | BLCa60 | 3.312251387 | 1.52566036 | -0.5256604 |
| erlotinib | 219804 | BLCa60 | 1.553001961 | 1.24646426 | -0.2464643 |
| everolimus | 128092 | BLCa60 | -1.724088735 | 0.72638396 | 0.27361604 |
| everolimus | 205803 | BLCa60 | 1.052712502 | 1.1670674 | -0.1670674 |
| everolimus | 172505 | BLCa60 | -0.137105253 | 0.97824115 | 0.02175885 |
| everolimus | 271006 | BLCa60 | 3.382572768 | 1.5368205 | -0.5368205 |
| gemcitabine | 119771 | BLCa60 | -2.350102954 | 0.52949219 | 0.47050781 |
| gemcitabine | 154214 | BLCa60 | -1.589550979 | 0.68176026 | 0.31823974 |
| gemcitabine | 93506 | BLCa60 | -2.93007269 | 0.413378 | 0.586622 |
| gemcitabine | 133923 | BLCa60 | -2.03760603 | 0.59205636 | 0.40794364 |
| lapatinib | 217546 | BLCa60 | 1.472318181 | 1.23365959 | -0.2336596 |
| lapatinib | 305058 | BLCa60 | 4.59933276 | 1.72992254 | -0.7299225 |
| lapatinib | 179179 | BLCa60 | 0.101372844 | 1.01608806 | -0.0160881 |
| lapatinib | 241551 | BLCa60 | 2.33007466 | 1.36978712 | -0.3697871 |
| methotrexate | 207632 | BLCa60 | 1.118067078 | 1.17743929 | -0.1774393 |
| methotrexate | 156367 | BLCa60 | -0.71375487 | 0.88672579 | 0.11327421 |
| methotrexate | 240276 | BLCa60 | 2.284515839 | 1.36255685 | -0.3625568 |
| methotrexate | 211623 | BLCa60 | 1.260675123 | 1.20007145 | -0.2000715 |
| olaparib | 248219 | BLCa60 | 2.568338363 | 1.4076 | -0.4076 |
| olaparib | 171497 | BLCa60 | -0.173123522 | 0.97252498 | 0.02747502 |
| olaparib | 150048 | BLCa60 | -0.939547963 | 0.85089202 | 0.14910798 |
| olaparib | 190877 | BLCa60 | 0.519370565 | 1.08242506 | -0.0824251 |
| paclitaxel | 160275 | BLCa60 | -0.574112616 | 0.90888728 | 0.09111272 |
| paclitaxel | 223098 | BLCa60 | 1.670704516 | 1.26514387 | -0.2651439 |
| paclitaxel | 156379 | BLCa60 | -0.713326081 | 0.88679384 | 0.11320616 |
| paclitaxel | 203550 | BLCa60 | 0.972207385 | 1.15429109 | -0.1542911 |
| ponatinib | 154260 | BLCa60 | -0.789043056 | 0.87477742 | 0.12522258 |
| ponatinib | 185093 | BLCa60 | 0.312694311 | 1.04962516 | -0.0496252 |
| ponatinib | 198371 | BLCa60 | 0.787149238 | 1.12492203 | -0.124922 |
| ponatinib | 180758 | BLCa60 | 0.157794318 | 1.02504225 | -0.0250422 |
| sunitinib | 155379 | BLCa60 | -0.74905849 | 0.88112304 | 0.11887696 |
| sunitinib | 178446 | BLCa60 | 0.075180989 | 1.01193136 | -0.0119314 |
| sunitinib | 203870 | BLCa60 | 0.983641756 | 1.15610575 | -0.1561057 |
| sunitinib | 181983 | BLCa60 | 0.201566519 | 1.03198898 | -0.031989 |
| temsirolimus | 140944 | BLCa60 | -1.264855815 | 0.79926506 | 0.20073494 |
| temsirolimus | 124949 | BLCa60 | -1.836395697 | 0.70856064 | 0.29143936 |
| temsirolimus | 136950 | BLCa60 | -1.407571056 | 0.77661589 | 0.22338411 |
| temsirolimus | 215439 | BLCa60 | 1.397029996 | 1.22171122 | -0.2217112 |
| vinblastine | 308416 | BLCa60 | 4.71932219 | 1.74896508 | -0.7489651 |
| vinblastine | 332728 | BLCa60 | 5.588048518 | 1.88683354 | -0.8868335 |
| vinblastine | 351087 | BLCa60 | 6.244059815 | 1.99094373 | -0.9909437 |
| vinblastine | 311381 | BLCa60 | 4.825268782 | 1.765779 | -0.765779 |
| cisplatin+gemcitabine | 134452 | BLCa60 | -2.025924934 | 0.594395 | 0.405605 |
| cisplatin+gemcitabine | 70308 | BLCa60 | -3.442318567 | 0.31082262 | 0.68917738 |
| cisplatin+gemcitabine | 98051 | BLCa60 | -2.829712422 | 0.43347086 | 0.56652914 |
| cisplatin+gemcitabine | 85191 | BLCa60 | -3.11368009 | 0.37661845 | 0.62338155 |
| untreated | 59990 | BLCa57 | -0.341767339 | 0.89749152 | |
| untreated | 64989 | BLCa57 | -0.092419623 | 0.97227999 | |
| untreated | 46831 | BLCa57 | -0.998131929 | 0.70062386 | |
| untreated | 38421 | BLCa57 | -1.417618684 | 0.5748045 | |

| | | | | | |
|---|---|---|---|---|---|
| untreated | 71830 | BLCa57 | 0.248806166 | 1.07462604 | |
| untreated | 57716 | BLCa57 | -0.455193365 | 0.86347092 | |
| untreated | 42271 | BLCa57 | -1.225582536 | 0.63240313 | |
| dmso | 76164 | BLCa57 | 0.464984002 | 1.13946565 | -0.1394656 |
| dmso | 91393 | BLCa57 | 1.224599197 | 1.36730193 | -0.3673019 |
| dmso | 78309 | BLCa57 | 0.57197557 | 1.17155632 | -0.1715563 |
| dmso | 82127 | BLCa57 | 0.762415574 | 1.22867621 | -0.2286762 |
| dmso | 55804 | BLCa57 | -0.550563005 | 0.83486609 | 0.16513391 |
| dmso | 40388 | BLCa57 | -1.31950567 | 0.60423216 | 0.39576784 |
| dmso | 43708 | BLCa57 | -1.153905667 | 0.65390164 | 0.34609836 |
| h2o | 69281 | BLCa57 | 0.665803305 | 1.16636644 | -0.1663664 |
| h2o | 74128 | BLCa57 | 0.992371673 | 1.24796714 | -0.2479671 |
| h2o | 61503 | BLCa57 | 0.141757757 | 1.03542147 | -0.0354215 |
| h2o | 77145 | BLCa57 | 1.195643133 | 1.29875924 | -0.2987592 |
| h2o | 41359 | BLCa57 | -1.215451489 | 0.69629118 | 0.30370882 |
| h2o | 43607 | BLCa57 | -1.06399168 | 0.73413694 | 0.26586306 |
| h2o | 48770 | BLCa57 | -0.716132698 | 0.82105759 | 0.17894241 |
| bosutinib | 72495 | BLCa57 | 0.281976047 | 1.08457489 | -0.0845749 |
| bosutinib | 68907 | BLCa57 | 0.103008332 | 1.03089595 | -0.030896 |
| bosutinib | 62563 | BLCa57 | -0.213427336 | 0.93598536 | 0.06401464 |
| bosutinib | 67801 | BLCa57 | 0.047841584 | 1.01434943 | -0.0143494 |
| cisplatin_6_um | 89270 | BLCa57 | 2.012569369 | 1.50288725 | -0.5028873 |
| cisplatin_6_um | 115268 | BLCa57 | 3.764193971 | 1.94057139 | -0.9405714 |
| cisplatin_6_um | 78843 | BLCa57 | 1.310046494 | 1.32734558 | -0.3273456 |
| cisplatin_6_um | 79986 | BLCa57 | 1.38705653 | 1.34658833 | -0.3465883 |
| cisplatin_14_um | 61739 | BLCa57 | 0.157658342 | 1.0393946 | -0.0393946 |
| cisplatin_14_um | 71031 | BLCa57 | 0.783710184 | 1.19582821 | -0.1958282 |
| cisplatin_14_um | 66111 | BLCa57 | 0.452223414 | 1.11299854 | -0.1129985 |
| cisplatin_14_um | 30445 | BLCa57 | -1.950786165 | 0.51255072 | 0.48744928 |
| crizotinib | 81623 | BLCa57 | 0.737276296 | 1.22113603 | -0.221136 |
| crizotinib | 87910 | BLCa57 | 1.050868832 | 1.31519386 | -0.3151939 |
| crizotinib | 82214 | BLCa57 | 0.766755092 | 1.22997779 | -0.2299778 |
| crizotinib | 75005 | BLCa57 | 0.407173639 | 1.12212621 | -0.1221262 |
| daunorubicin | 51136 | BLCa57 | -0.7834006 | 0.76502961 | 0.23497039 |
| daunorubicin | 71486 | BLCa57 | 0.231647612 | 1.06947956 | -0.0694796 |
| daunorubicin | 57305 | BLCa57 | -0.475693847 | 0.85732208 | 0.14267792 |
| daunorubicin | 60671 | BLCa57 | -0.307799386 | 0.90767975 | 0.09232025 |
| docetaxel | 62335 | BLCa57 | -0.224799867 | 0.93257433 | 0.06742567 |
| docetaxel | 75373 | BLCa57 | 0.425529302 | 1.12763174 | -0.1276317 |
| docetaxel | 82909 | BLCa57 | 0.801421358 | 1.24037547 | -0.2403755 |
| docetaxel | 102814 | BLCa57 | 1.794273183 | 1.53816791 | -0.5381679 |
| doxorubicin | 37800 | BLCa57 | -1.448593865 | 0.56551391 | 0.43448609 |
| doxorubicin | 24218 | BLCa57 | -2.126057493 | 0.36231788 | 0.63768212 |
| doxorubicin | 28657 | BLCa57 | -1.904642308 | 0.42872836 | 0.57127164 |
| doxorubicin | 24454 | BLCa57 | -2.114285926 | 0.3658486 | 0.6341514 |
| epirubicin | 44788 | BLCa57 | -1.100035787 | 0.67005918 | 0.32994082 |
| epirubicin | 53851 | BLCa57 | -0.647977706 | 0.80564787 | 0.19435213 |
| epirubicin | 40037 | BLCa57 | -1.337013381 | 0.59898096 | 0.40101904 |
| epirubicin | 16673 | BLCa57 | -2.502398464 | 0.24943951 | 0.75056049 |
| erlotinib | 46976 | BLCa57 | -0.990899399 | 0.70279316 | 0.29720684 |
| erlotinib | 68176 | BLCa57 | 0.066546404 | 1.01995969 | -0.0199597 |
| erlotinib | 72814 | BLCa57 | 0.297887613 | 1.08934735 | -0.0893474 |
| erlotinib | 44746 | BLCa57 | -1.102130726 | 0.66943083 | 0.33056917 |
| everolimus | 47608 | BLCa57 | -0.959375543 | 0.71224831 | 0.28775169 |
| everolimus | 34829 | BLCa57 | -1.596785916 | 0.52106571 | 0.47893429 |
| everolimus | 28257 | BLCa57 | -1.924594115 | 0.42274409 | 0.57725591 |
| everolimus | 41126 | BLCa57 | -1.282694585 | 0.61527315 | 0.38472685 |
| gemcitabine | 21861 | BLCa57 | -2.529136252 | 0.3680365 | 0.6319635 |
| gemcitabine | 27229 | BLCa57 | -2.167465321 | 0.45840839 | 0.54159161 |
| gemcitabine | 26971 | BLCa57 | -2.184848164 | 0.45406488 | 0.54593512 |
| gemcitabine | 23650 | BLCa57 | -2.408601734 | 0.39815485 | 0.60184515 |
| lapatinib | 29126 | BLCa57 | -1.881248813 | 0.43574492 | 0.56425508 |
| lapatinib | 33571 | BLCa57 | -1.659534351 | 0.50224517 | 0.49775483 |
| lapatinib | 36658 | BLCa57 | -1.505556276 | 0.54842881 | 0.45157119 |
| lapatinib | 41189 | BLCa57 | -1.279552176 | 0.61621567 | 0.38378433 |
| methotrexate | 33891 | BLCa57 | -1.643572905 | 0.50703259 | 0.49296741 |
| methotrexate | 47586 | BLCa57 | -0.960472892 | 0.71191918 | 0.28808082 |

| | | | | | |
|---|---|---|---|---|---|
| methotrexate | 43096 | BLCa57 | -1.184431933 | 0.6447457 | 0.3552543 |
| methotrexate | 49145 | BLCa57 | -0.882710722 | 0.73524289 | 0.26475711 |
| olaparib | 44317 | BLCa57 | -1.12352904 | 0.6630127 | 0.3369873 |
| olaparib | 29435 | BLCa57 | -1.865836042 | 0.44036778 | 0.55963222 |
| olaparib | 45944 | BLCa57 | -1.042375063 | 0.68735373 | 0.31264627 |
| olaparib | 39501 | BLCa57 | -1.363748804 | 0.59096204 | 0.40903796 |
| paclitaxel | 90679 | BLCa57 | 1.18898522 | 1.35662 | -0.35662 |
| paclitaxel | 64570 | BLCa57 | -0.113319142 | 0.96601146 | 0.03398854 |
| paclitaxel | 33060 | BLCa57 | -1.685022786 | 0.49460026 | 0.50539974 |
| paclitaxel | 56984 | BLCa57 | -0.491705173 | 0.8525197 | 0.1474803 |
| ponatinib | 63907 | BLCa57 | -0.146389263 | 0.95609253 | 0.04390747 |
| ponatinib | 61767 | BLCa57 | -0.253131433 | 0.92407666 | 0.07592334 |
| ponatinib | 67310 | BLCa57 | 0.023350741 | 1.00700374 | -0.0070037 |
| ponatinib | 66579 | BLCa57 | -0.013111188 | 0.99606748 | 0.00393252 |
| rapamycin | 68464 | BLCa57 | 0.080911705 | 1.02426837 | -0.0242684 |
| rapamycin | 56483 | BLCa57 | -0.516694812 | 0.8450244 | 0.1549756 |
| rapamycin | 51636 | BLCa57 | -0.75846084 | 0.77250995 | 0.22749005 |
| rapamycin | 53952 | BLCa57 | -0.642939874 | 0.8071589 | 0.1928411 |
| sunitinib | 58693 | BLCa57 | -0.406461075 | 0.87808751 | 0.12191249 |
| sunitinib | 59257 | BLCa57 | -0.378329026 | 0.88652534 | 0.11347466 |
| sunitinib | 59709 | BLCa57 | -0.355783483 | 0.89328757 | 0.10671243 |
| sunitinib | 44091 | BLCa57 | -1.134801811 | 0.65963158 | 0.34036842 |
| temsirolimus | 55784 | BLCa57 | -0.551560596 | 0.83456688 | 0.16543312 |
| temsirolimus | 44251 | BLCa57 | -1.126821088 | 0.66202529 | 0.33797471 |
| temsirolimus | 29158 | BLCa57 | -1.879652669 | 0.43622367 | 0.56377633 |
| temsirolimus | 55448 | BLCa57 | -0.568320114 | 0.82954009 | 0.17045991 |
| vinblastine | 86691 | BLCa57 | 0.990065698 | 1.29695678 | -0.2969568 |
| vinblastine | 68893 | BLCa57 | 0.102310019 | 1.0306865 | -0.0306865 |
| vinblastine | 58962 | BLCa57 | -0.393043484 | 0.88211194 | 0.11788806 |
| vinblastine | 57301 | BLCa57 | -0.475893365 | 0.85726224 | 0.14273776 |
| cisplatin+gemcitabine | 31274 | BLCa57 | -1.894931991 | 0.52650718 | 0.47349282 |
| cisplatin+gemcitabine | 34760 | BLCa57 | -1.660061488 | 0.58519504 | 0.41480496 |
| cisplatin+gemcitabine | 34783 | BLCa57 | -1.658511854 | 0.58558225 | 0.41441775 |
| cisplatin+gemcitabine | 38224 | BLCa57 | -1.426673242 | 0.64351252 | 0.35648748 |
| untreated | 17550 | BLCa50 | -0.602902672 | 0.95087593 | |
| untreated | 16439 | BLCa50 | -1.341680101 | 0.89068087 | |
| untreated | 17511 | BLCa50 | -0.628836353 | 0.94876287 | |
| untreated | 18559 | BLCa50 | 0.068048206 | 1.00554452 | |
| dmso | 17738 | BLCa50 | -0.47788903 | 0.96106195 | 0.03893805 |
| dmso | 20185 | BLCa50 | 1.149283219 | 1.09364277 | -0.0936428 |
| dmso | 17447 | BLCa50 | -0.671394189 | 0.94529529 | 0.05470471 |
| epirubicin | 4857 | BLCa50 | -9.04331843 | 0.26315694 | 0.73684306 |
| epirubicin | 3599 | BLCa50 | -9.879845887 | 0.19499729 | 0.80500271 |
| epirubicin | 3628 | BLCa50 | -9.860561868 | 0.19656854 | 0.80343146 |
| crizotinib | 18192 | BLCa50 | -0.175994383 | 0.9856601 | 0.0143399 |
| crizotinib | 17564 | BLCa50 | -0.593593146 | 0.95163446 | 0.04836554 |
| crizotinib | 13765 | BLCa50 | -3.119799674 | 0.74580098 | 0.25419902 |
| erlotinib | 14620 | BLCa50 | -2.551253588 | 0.7921257 | 0.2078743 |
| erlotinib | 11406 | BLCa50 | -4.688454899 | 0.61798808 | 0.38201192 |
| erlotinib | 11079 | BLCa50 | -4.905898841 | 0.6002709 | 0.3997291 |
| lapatinib | 5709 | BLCa50 | -8.476767242 | 0.30931913 | 0.69068087 |
| lapatinib | 6170 | BLCa50 | -8.170217832 | 0.33429655 | 0.66570345 |
| lapatinib | 13006 | BLCa50 | -3.624509007 | 0.70467762 | 0.29532238 |
| untreated | 46697 | BLCa50 | 0.065791798 | 1.00804382 | |
| untreated | 51273 | BLCa50 | 0.873744207 | 1.10682551 | |
| untreated | 44546 | BLCa50 | -0.313995272 | 0.96161038 | |
| untreated | 48276 | BLCa50 | 0.344584817 | 1.04212955 | |
| untreated | 50308 | BLCa50 | 0.703360886 | 1.08599414 | |
| untreated | 46771 | BLCa50 | 0.078857462 | 1.00964125 | |
| untreated | 42246 | BLCa50 | -0.720090233 | 0.9119605 | |
| untreated | 45831 | BLCa50 | -0.087111783 | 0.98934956 | |
| h2o | 48977 | BLCa50 | 0.468355498 | 1.05726197 | -0.057262 |
| h2o | 47334 | BLCa50 | 0.178262446 | 1.02179468 | -0.0217947 |
| h2o | 43458 | BLCa50 | -0.506095845 | 0.93812383 | 0.06187617 |
| h2o | 48679 | BLCa50 | 0.415739716 | 1.05082907 | -0.0508291 |
| h2o | 53150 | BLCa50 | 1.205153008 | 1.14734414 | -0.1473441 |
| h2o | 43306 | BLCa50 | -0.532933425 | 0.93484262 | 0.06515738 |

| | | | | | |
|---|---|---|---|---|---|
| h2o | 35009 | BLCa50 | -1.997876856 | 0.75573605 | 0.24426395 |
| h2o | 50682 | BLCa50 | 0.769395458 | 1.09406765 | -0.0940676 |
| cisplatin_6_um | 48117 | BLCa50 | 0.316511295 | 1.03869723 | -0.0386972 |
| cisplatin_6_um | 51563 | BLCa50 | 0.924947485 | 1.11308571 | -0.1130857 |
| cisplatin_6_um | 47132 | BLCa50 | 0.142596714 | 1.01743413 | -0.0174341 |
| cisplatin_6_um | 55375 | BLCa50 | 1.598005742 | 1.19537501 | -0.195375 |
| cisplatin_14_um | 48290 | BLCa50 | 0.347056699 | 1.04243177 | -0.0424318 |
| cisplatin_14_um | 47533 | BLCa50 | 0.213398488 | 1.02609048 | -0.0260905 |
| cisplatin_14_um | 48713 | BLCa50 | 0.421742859 | 1.05156303 | -0.051563 |
| cisplatin_14_um | 50143 | BLCa50 | 0.674227987 | 1.0824323 | -0.0824323 |
| gemcitabine | 34647 | BLCa50 | -2.061792672 | 0.74792159 | 0.25207841 |
| gemcitabine | 34783 | BLCa50 | -2.0377801 | 0.7508574 | 0.2491426 |
| gemcitabine | 41567 | BLCa50 | -0.839976528 | 0.89730299 | 0.10269701 |
| gemcitabine | 35671 | BLCa50 | -1.880992133 | 0.77002658 | 0.22997342 |
| cisplatin+gemcitabine | 35140 | BLCa50 | -1.9747471 | 0.75856393 | 0.24143607 |
| cisplatin+gemcitabine | 37177 | BLCa50 | -1.615088215 | 0.80253646 | 0.19746354 |
| cisplatin+gemcitabine | 35293 | BLCa50 | -1.947732957 | 0.76186673 | 0.23813327 |
| cisplatin+gemcitabine | 31447 | BLCa50 | -2.626794357 | 0.67884348 | 0.32115652 |
| doxorubicin | 11216 | BLCa50 | -6.198840946 | 0.24211875 | 0.75788125 |
| doxorubicin | 11175 | BLCa50 | -6.20608003 | 0.24123369 | 0.75876631 |
| doxorubicin | 13617 | BLCa50 | -5.774913119 | 0.29394892 | 0.70605108 |
| untreated | 4728 | BLCa47 | -1.365733244 | 0.61844343 | 0.38155657 |
| untreated | 5765 | BLCa47 | -0.880212033 | 0.75408764 | 0.24591236 |
| untreated | 7923 | BLCa47 | 0.130159013 | 1.03636364 | -0.0363636 |
| dmso | 9479 | BLCa47 | 0.85867493 | 1.23989536 | -0.2398954 |
| dmso | 8156 | BLCa47 | 0.239249122 | 1.06684107 | -0.0668411 |
| dmso | 5300 | BLCa47 | -1.097924051 | 0.69326357 | 0.30673643 |
| h2o | 8069 | BLCa47 | -1.128161977 | 0.9256271 | 0.0743729 |
| h2o | 8919 | BLCa47 | 0.350919278 | 1.02313399 | -0.023134 |
| h2o | 9164 | BLCa47 | 0.777242699 | 1.05123891 | -0.0512389 |
| cisplatin_6_um | 8228 | BLCa47 | -0.851486778 | 0.94386663 | 0.05613337 |
| cisplatin_6_um | 7213 | BLCa47 | -2.617683806 | 0.82743194 | 0.17256806 |
| cisplatin_6_um | 5718 | BLCa47 | -5.219126721 | 0.65593454 | 0.34406546 |
| gemcitabine | 4969 | BLCa47 | -6.522458321 | 0.57001377 | 0.42998623 |
| gemcitabine | 5365 | BLCa47 | -5.833380466 | 0.6154405 | 0.3845595 |
| gemcitabine | 6854 | BLCa47 | -3.242378125 | 0.78624962 | 0.21375038 |
| cisplatin+gemcitabine | 6819 | BLCa47 | -3.303281471 | 0.78223463 | 0.21776537 |
| cisplatin+gemcitabine | 3431 | BLCa47 | -9.198725346 | 0.39358366 | 0.60641634 |
| cisplatin+gemcitabine | 3657 | BLCa47 | -8.805463741 | 0.41950902 | 0.58049098 |
| epirubicin | 324 | BLCa47 | -3.427676751 | 0.04238064 | 0.95761936 |
| epirubicin | 413 | BLCa47 | -3.386007139 | 0.05402224 | 0.94597776 |
| epirubicin | 347 | BLCa47 | -3.416908199 | 0.04538914 | 0.95461086 |
| lapatinib | 9491 | BLCa47 | 0.864293304 | 1.24146501 | -0.241465 |
| lapatinib | 8680 | BLCa47 | 0.484584816 | 1.1353826 | -0.1353826 |
| lapatinib | 5465 | BLCa47 | -1.0206714 | 0.7148463 | 0.2851537 |
| everolimus | 5563 | BLCa47 | -0.974788006 | 0.72766514 | 0.27233486 |
| everolimus | 8997 | BLCa47 | 0.633003547 | 1.17684761 | -0.1768476 |
| everolimus | 4352 | BLCa47 | -1.541775651 | 0.56926095 | 0.43073905 |
| ponatinib | 5862 | BLCa47 | -0.834796837 | 0.76677567 | 0.23322433 |
| ponatinib | 9828 | BLCa47 | 1.022075993 | 1.28554611 | -0.2855461 |
| ponatinib | 5691 | BLCa47 | -0.914858677 | 0.74440811 | 0.25559189 |
| untreated | 12933 | BLCa35 | 4.689583334 | 1.21001092 | |
| untreated | 10792 | BLCa35 | 0.216581589 | 1.00969905 | |
| untreated | 10909 | BLCa35 | 0.46101933 | 1.02064556 | |
| dmso | 11119 | BLCa35 | 0.899753737 | 1.04029315 | -0.0402932 |
| dmso | 10773 | BLCa35 | 0.176886571 | 1.00792141 | -0.0079214 |
| dmso | 10173 | BLCa35 | -1.076640308 | 0.95178544 | 0.04821456 |
| h2o | 12825 | BLCa35 | 0.851962065 | 1.1174523 | -0.1174523 |
| h2o | 11871 | BLCa35 | 0.249015619 | 1.03432953 | -0.0343295 |
| h2o | 9735 | BLCa35 | -1.100977684 | 0.84821818 | 0.15178182 |
| cisplatin_6_um | 10540 | BLCa35 | -0.592202118 | 0.91835846 | 0.08164154 |
| cisplatin_6_um | 11668 | BLCa35 | 0.120715693 | 1.01664198 | -0.016642 |
| cisplatin_6_um | 12095 | BLCa35 | 0.39058795 | 1.05384682 | -0.0538468 |
| gemcitabine | 7784 | BLCa35 | -2.334047409 | 0.67822602 | 0.32177398 |
| gemcitabine | 7382 | BLCa35 | -2.588119183 | 0.64319944 | 0.35680056 |
| gemcitabine | 4268 | BLCa35 | -4.556227396 | 0.37187418 | 0.62812582 |
| cisplatin+gemcitabine | 4848 | BLCa35 | -4.189656181 | 0.42241004 | 0.57758996 |

| | | | | | |
|---|---|---|---|---|---|
| cisplatin+gemcitabine | 5910 | BLCa35 | -3.518451646 | 0.51494293 | 0.48505707 |
| cisplatin+gemcitabine | 4112 | BLCa35 | -4.654822413 | 0.35828178 | 0.64171822 |
| epirubicin | 3737 | BLCa35 | -14.52280529 | 0.34963356 | 0.65036644 |
| epirubicin | 4181 | BLCa35 | -13.5951954 | 0.39117418 | 0.60882582 |
| epirubicin | 2999 | BLCa35 | -16.06464336 | 0.28058631 | 0.71941369 |
| lapatinib | 8846 | BLCa35 | -3.849023922 | 0.82763137 | 0.17236863 |
| lapatinib | 6584 | BLCa35 | -8.574820255 | 0.61599875 | 0.38400125 |
| lapatinib | 8058 | BLCa35 | -5.495322556 | 0.75390613 | 0.24609387 |
| lapatinib | 8235 | BLCa35 | -5.125532127 | 0.77046624 | 0.22953376 |
| crizotinib | 6854 | BLCa35 | -8.010733159 | 0.64125994 | 0.35874006 |
| crizotinib | 10834 | BLCa35 | 0.30432847 | 1.01362857 | -0.0136286 |
| crizotinib | 1976 | BLCa35 | -18.20190668 | 0.18487447 | 0.81512553 |
| erdafitinib | 9209 | BLCa35 | -3.09064016 | 0.86159364 | 0.13840636 |
| erdafitinib | 10019 | BLCa35 | -1.398378874 | 0.9373772 | 0.0626228 |
| erdafitinib | 9767 | BLCa35 | -1.924860163 | 0.91380009 | 0.08619991 |
| untreated | 94292 | BLCa34 | 4.020383749 | 2.12831143 | |
| untreated | 31144 | BLCa34 | -1.058385157 | 0.70296665 | |
| untreated | 68679 | BLCa34 | 1.960421312 | 1.55018772 | |
| untreated | 37230 | BLCa34 | -0.568909836 | 0.84033677 | |
| untreated | 35833 | BLCa34 | -0.681265574 | 0.80880439 | |
| untreated | 28609 | BLCa34 | -1.262266185 | 0.64574791 | |
| dmso | 57799 | BLCa34 | 1.085381632 | 1.30460985 | -0.3046099 |
| dmso | 51585 | BLCa34 | 0.585611727 | 1.16435058 | -0.1643506 |
| dmso | 56579 | BLCa34 | 0.987261374 | 1.27707263 | -0.2770726 |
| dmso | 36881 | BLCa34 | -0.596978664 | 0.83245931 | 0.16754069 |
| dmso | 32837 | BLCa34 | -0.922223192 | 0.74118019 | 0.25881981 |
| dmso | 30141 | BLCa34 | -1.139052877 | 0.68032744 | 0.31967256 |
| h2o | 60926 | BLCa34 | 0.664638384 | 1.28490866 | -0.2849087 |
| h2o | 61708 | BLCa34 | 0.703111389 | 1.30140078 | -0.3014008 |
| h2o | 63695 | BLCa34 | 0.800868245 | 1.34330593 | -0.3433059 |
| h2o | 29208 | BLCa34 | -0.895830643 | 0.61598681 | 0.38401319 |
| h2o | 21546 | BLCa34 | -1.272787376 | 0.45439783 | 0.54560217 |
| bosutinib | 38695 | BLCa34 | -0.4510851 | 0.87340401 | 0.12659599 |
| bosutinib | 40902 | BLCa34 | -0.273583944 | 0.9232193 | 0.0767807 |
| bosutinib | 31851 | BLCa34 | -1.001523663 | 0.71892469 | 0.28107531 |
| cisplatin_14_um | 38867 | BLCa34 | -0.420625071 | 0.81969184 | 0.18030816 |
| cisplatin_14_um | 33177 | BLCa34 | -0.700562922 | 0.69969167 | 0.30030833 |
| cisplatin_14_um | 37710 | BLCa34 | -0.477547407 | 0.7952911 | 0.2047089 |
| crizotinib | 30653 | BLCa34 | -1.097874539 | 0.69188404 | 0.30811596 |
| crizotinib | 29408 | BLCa34 | -1.198005459 | 0.66378253 | 0.33621747 |
| crizotinib | 29115 | BLCa34 | -1.221570406 | 0.65716908 | 0.34283092 |
| daunorubicin | 49229 | BLCa34 | 0.396127031 | 1.11117214 | -0.1111721 |
| daunorubicin | 50712 | BLCa34 | 0.515399444 | 1.14464567 | -0.1446457 |
| daunorubicin | 44308 | BLCa34 | 0.000348515 | 1.00009781 | -9.781E-05 |
| doxorubicin | 7421 | BLCa34 | -2.96634162 | 0.16750307 | 0.83249693 |
| doxorubicin | 7602 | BLCa34 | -2.951784434 | 0.17158851 | 0.82841149 |
| doxorubicin | 6036 | BLCa34 | -3.077732241 | 0.13624155 | 0.86375845 |
| erlotinib | 41374 | BLCa34 | -0.235622664 | 0.93387304 | 0.06612696 |
| erlotinib | 30024 | BLCa34 | -1.148462771 | 0.67768657 | 0.32231343 |
| erlotinib | 29421 | BLCa34 | -1.196959915 | 0.66407596 | 0.33592404 |
| gemcitabine | 36369 | BLCa34 | -0.543522215 | 0.76700987 | 0.23299013 |
| gemcitabine | 31429 | BLCa34 | -0.786561404 | 0.66282694 | 0.33717306 |
| gemcitabine | 32369 | BLCa34 | -0.740315081 | 0.68265122 | 0.31734878 |
| lapatinib | 36195 | BLCa34 | -0.652151202 | 0.81697527 | 0.18302473 |
| lapatinib | 22471 | BLCa34 | -1.755923681 | 0.50720407 | 0.49279593 |
| lapatinib | 23638 | BLCa34 | -1.662066024 | 0.533545 | 0.466455 |
| untreated | 6759 | BLCa33 | -0.463998439 | 0.93589034 | |
| untreated | 5497 | BLCa33 | -1.728719886 | 0.7611465 | |
| untreated | 5989 | BLCa33 | -1.23565891 | 0.82927167 | |
| untreated | 7348 | BLCa33 | 0.126271713 | 1.01744669 | |
| dmso | 8042 | BLCa33 | 0.821768293 | 1.11354196 | -0.113542 |
| dmso | 7513 | BLCa33 | 0.291627529 | 1.04029355 | -0.0402935 |
| dmso | 6111 | BLCa33 | -1.113395822 | 0.8461645 | 0.1538355 |
| h2o | 5859 | BLCa33 | -1.0193627 | 0.92228985 | 0.07771015 |
| h2o | 6827 | BLCa33 | 0.979441676 | 1.07466681 | -0.0746668 |
| h2o | 6372 | BLCa33 | 0.039921024 | 1.00304334 | -0.0030433 |
| cisplatin_6_um | 5864 | BLCa33 | -1.009038298 | 0.92307692 | 0.07692308 |

| | | | | | |
|---|---|---|---|---|---|
| cisplatin_6_um | 5497 | BLCa33 | -1.766849461 | 0.86530591 | 0.13469409 |
| cisplatin_6_um | 5079 | BLCa33 | -2.629969533 | 0.79950677 | 0.20049323 |
| gemcitabine | 5877 | BLCa33 | -0.98219485 | 0.92512331 | 0.07487669 |
| gemcitabine | 5622 | BLCa33 | -1.508739392 | 0.88498268 | 0.11501732 |
| gemcitabine | 5014 | BLCa33 | -2.764186769 | 0.78927485 | 0.21072515 |
| cisplatin+gemcitabine | 3640 | BLCa33 | -5.60133265 | 0.57298772 | 0.42701228 |
| cisplatin+gemcitabine | 7193 | BLCa33 | 1.735187959 | 1.13228041 | -0.1322804 |
| cisplatin+gemcitabine | 4873 | BLCa33 | -3.055334927 | 0.76707944 | 0.23292056 |
| epirubicin | 1519 | BLCa33 | -5.715298265 | 0.21032955 | 0.78967045 |
| epirubicin | 2533 | BLCa33 | -4.69911162 | 0.35073387 | 0.64926613 |
| epirubicin | 1455 | BLCa33 | -5.779436279 | 0.20146774 | 0.79853226 |
| everolimus | 5806 | BLCa33 | -1.419053541 | 0.80393243 | 0.19606757 |
| everolimus | 4896 | BLCa33 | -2.331015915 | 0.67792855 | 0.32207145 |
| everolimus | 4990 | BLCa33 | -2.236813209 | 0.69094434 | 0.30905566 |
| lapatinib | 6088 | BLCa33 | -1.13644542 | 0.84297978 | 0.15702022 |
| lapatinib | 4902 | BLCa33 | -2.325002977 | 0.67875935 | 0.32124065 |
| lapatinib | 5985 | BLCa33 | -1.239667535 | 0.82871781 | 0.17128219 |
| erdafitinib | 5405 | BLCa33 | -1.82091828 | 0.74840764 | 0.25159236 |
| erdafitinib | 5865 | BLCa33 | -1.35992631 | 0.81210191 | 0.18789809 |
| erdafitinib | 5880 | BLCa33 | -1.344893963 | 0.8141789 | 0.1858211 |
