## Supplemental tables for "Bladder cancer organoids as a functional system to model different disease stages and therapy response": Table S10.pdf

| mut | drug | gene | beta | pval | fdr |
|---|---|---|---|---|---|
| gain | bosutinib | ABL1 | -0.64243666 | 0.50403 | 0.921373782 |
| gain | bosutinib | SRC | -0.416807645 | 0.54531 | 0.921373782 |
| gain | bosutinib | LYN | 1 | 0.10297 | 0.575777647 |
| gain | bosutinib | HCK | -0.416807645 | 0.54531 | 0.921373782 |
| gain | bosutinib | CSK | 0.077152588 | 0.92471 | 0.982244706 |
| gain | bosutinib | ABL2 | -0.235480009 | 0.76279 | 0.982244706 |
| gain | bosutinib | FGR | -0.039792801 | 0.96476 | 0.982244706 |
| gain | bosutinib | TEC | 0.077152588 | 0.92471 | 0.982244706 |
| gain | bosutinib | LCK | -0.039792801 | 0.96476 | 0.982244706 |
| gain | bosutinib | CAMK1D | -0.085926402 | 0.90777 | 0.982244706 |
| gain | bosutinib | FRK | -0.482002569 | 0.53145 | 0.921373782 |
| gain | bosutinib | FYN | -0.482002569 | 0.53145 | 0.921373782 |
| gain | bosutinib | STK10 | -0.138581735 | 0.86811 | 0.982244706 |
| gain | bosutinib | EPHB4 | -0.317057462 | 0.66406 | 0.975338125 |
| gain | bosutinib | YES1 | -0.121340675 | 0.88116 | 0.982244706 |
| gain | bosutinib | MAP4K2 | 0.077152588 | 0.92471 | 0.982244706 |
| gain | bosutinib | PTK2 | 1 | 0.10297 | 0.575777647 |
| gain | bosutinib | SYK | -0.553302395 | 0.48748 | 0.921373782 |
| gain | bosutinib | MAP4K5 | -0.085926402 | 0.90777 | 0.982244706 |
| gain | bosutinib | FER | -0.138581735 | 0.86811 | 0.982244706 |
| gain | bosutinib | STK4 | -0.416807645 | 0.54531 | 0.921373782 |
| gain | bosutinib | STK24 | -0.298556634 | 0.68281 | 0.982244706 |
| gain | bosutinib | TBK1 | 0.077152588 | 0.92471 | 0.982244706 |
| gain | bosutinib | PTK2B | 0.41408893 | 0.4592 | 0.921373782 |
| gain | bosutinib | TNK2 | -0.317057462 | 0.66406 | 0.975338125 |
| gain | bosutinib | TXK | 0.077152588 | 0.92471 | 0.982244706 |
| gain | bosutinib | CDK2 | -0.039792801 | 0.96476 | 0.982244706 |
| gain | bosutinib | MAP2K1 | 0.077152588 | 0.92471 | 0.982244706 |
| gain | bosutinib | MAP2K2 | -0.537455377 | 0.45121 | 0.921373782 |
| gain | bosutinib | MAP3K2 | 0.077152588 | 0.92471 | 0.982244706 |
| gain | bosutinib | ALK | 0.077152588 | 0.92471 | 0.982244706 |
| gain | bosutinib | TAOK3 | 0.077152588 | 0.92471 | 0.982244706 |
| gain | bosutinib | PRKAA1 | -0.317057462 | 0.66406 | 0.975338125 |
| gain | bosutinib | TNK2 | -0.317057462 | 0.66406 | 0.975338125 |
| gain | bosutinib | AAK1 | 0.077152588 | 0.92471 | 0.982244706 |
| gain | bosutinib | GAK | 0.077152588 | 0.92471 | 0.982244706 |
| gain | bosutinib | BMP2K | 0.077152588 | 0.92471 | 0.982244706 |
| gain | bosutinib | MAP3K3 | -0.617134383 | 0.35995 | 0.870653636 |
| gain | bosutinib | STK35 | -0.085926402 | 0.90777 | 0.982244706 |
| gain | bosutinib | MAPK14 | -0.317057462 | 0.66406 | 0.975338125 |
| gain | bosutinib | TNIK | -0.317057462 | 0.66406 | 0.975338125 |
| gain | bosutinib | MAP4K4 | 0.077152588 | 0.92471 | 0.982244706 |
| gain | bosutinib | MAP3K1 | -0.138581735 | 0.86811 | 0.982244706 |
| gain | bosutinib | MAP4K1 | -0.617134383 | 0.35995 | 0.870653636 |
| gain | bosutinib | MAP2K5 | 0.077152588 | 0.92471 | 0.982244706 |
| gain | bosutinib | MAP3K4 | -0.263511744 | 0.7621 | 0.982244706 |
| gain | crizotinib | ROS1 | -2 | 0.09246 | 0.575777647 |
| gain | erlotinib | EGFR | -1 | 0.0985 | 0.575777647 |
| gain | erlotinib | ERBB2 | -1 | 0.10334 | 0.575777647 |
| gain | erlotinib | NR1I2 | -1 | 0.0985 | 0.575777647 |
| gain | erlotinib | ERBB2 | -1 | 0.10334 | 0.575777647 |
| gain | erlotinib | AAK1 | -1 | 0.10334 | 0.575777647 |
| gain | erlotinib | ABL1 | -0.658963403 | 0.40743 | 0.870653636 |
| gain | erlotinib | ABL2 | -1 | 0.27759 | 0.870653636 |
| gain | erlotinib | BMP2K | -1 | 0.10334 | 0.575777647 |
| gain | erlotinib | EPHA6 | -1 | 0.10334 | 0.575777647 |
| gain | erlotinib | GAK | 0.17063733 | 0.87508 | 0.982244706 |
| gain | erlotinib | MAPK9 | -2 | 0.04591 | 0.575777647 |
| gain | erlotinib | LCK | -1 | 0.13099 | 0.631438974 |
| gain | erlotinib | MKNK2 | -2 | 0.02831 | 0.575777647 |
| gain | erlotinib | RIPK2 | -0.033950045 | 0.97348 | 0.982244706 |
| gain | erlotinib | SRC | -0.849629866 | 0.28215 | 0.870653636 |
| gain | erlotinib | STK10 | -0.204122613 | 0.84808 | 0.982244706 |
| gain | erlotinib | ABCG2 | -1 | 0.10334 | 0.575777647 |
| gain | erlotinib | ABCB1 | -1 | 0.0985 | 0.575777647 |
| gain | lapatinib | EGFR | -2 | 0.03746 | 0.575777647 |

| | | | | | |
|---|---|---|---|---|---|
| gain | lapatinib | ERBB2 | -3 | 0.0163 | 0.575777647 |
| gain | lapatinib | ERBB4 | -3 | 0.04143 | 0.575777647 |
| gain | lapatinib | CYP3A4 | -2 | 0.03746 | 0.575777647 |
| gain | lapatinib | CYP2C8 | -1 | 0.3906 | 0.870653636 |
| gain | lapatinib | ABCB1 | -2 | 0.03746 | 0.575777647 |
| gain | lapatinib | ABCG2 | -3 | 0.0163 | 0.575777647 |
| gain | lapatinib | STK10 | -0.715158942 | 0.6222 | 0.959231475 |
| gain | lapatinib | SLK | -1 | 0.3906 | 0.870653636 |
| gain | ponatinib | FGFR1 | -0.861262407 | 0.07521 | 0.575777647 |
| gain | ponatinib | FGFR3 | -0.363642136 | 0.57992 | 0.921373782 |
| gain | ponatinib | FGFR4 | -1 | 0.05126 | 0.575777647 |
| gain | ponatinib | SRC | -1 | 0.0228 | 0.575777647 |
| gain | ponatinib | RET | -0.363642136 | 0.57992 | 0.921373782 |
| gain | ponatinib | TEK | -0.355524751 | 0.25471 | 0.870653636 |
| gain | ponatinib | FLT3 | -0.392836918 | 0.50568 | 0.921373782 |
| gain | ponatinib | LCK | -0.415608677 | 0.55795 | 0.921373782 |
| gain | ponatinib | LYN | -0.083834936 | 0.87592 | 0.982244706 |
| gain | sunitinib | FLT3 | -0.731514334 | 0.26495 | 0.870653636 |
| gain | sunitinib | PDGFRB | 0.029369326 | 0.97702 | 0.982244706 |
| gain | sunitinib | CSF1R | 0.029369326 | 0.97702 | 0.982244706 |
| gain | sunitinib | RET | -0.617882202 | 0.40754 | 0.870653636 |
| gain | sunitinib | ABCG2 | -0.617882202 | 0.40754 | 0.870653636 |
| gain | sunitinib | ABCB1 | 0.112037265 | 0.88071 | 0.982244706 |
| gain | sunitinib | AAK1 | -0.617882202 | 0.40754 | 0.870653636 |
| gain | sunitinib | ABL1 | -0.093285528 | 0.87109 | 0.982244706 |
| gain | sunitinib | ABL2 | -1.022055757 | 0.14706 | 0.674323902 |
| gain | sunitinib | AURKC | 0.467573805 | 0.4841 | 0.921373782 |
| gain | sunitinib | BMP2K | -0.617882202 | 0.40754 | 0.870653636 |
| gain | sunitinib | CAMK1 | -0.617882202 | 0.40754 | 0.870653636 |
| gain | sunitinib | CAMK1D | -0.839254399 | 0.20351 | 0.721884528 |
| gain | sunitinib | CAMK1G | -1.323224899 | 0.08092 | 0.575777647 |
| gain | sunitinib | CAMK2A | 0.029369326 | 0.97702 | 0.982244706 |
| gain | sunitinib | CAMK2B | 0.112037265 | 0.88071 | 0.982244706 |
| gain | sunitinib | CAMKK1 | -0.423664642 | 0.564 | 0.921373782 |
| gain | sunitinib | CAMKK2 | -1 | 0.40754 | 0.870653636 |
| gain | sunitinib | CLK1 | -0.423664642 | 0.564 | 0.921373782 |
| gain | sunitinib | CLK2 | -1.221021536 | 0.07326 | 0.575777647 |
| gain | sunitinib | CLK4 | 0.029369326 | 0.97702 | 0.982244706 |
| gain | sunitinib | DAPK2 | -0.617882202 | 0.40754 | 0.870653636 |
| gain | sunitinib | DAPK3 | 0.467573805 | 0.4841 | 0.921373782 |
| gain | sunitinib | EPHA5 | -0.617882202 | 0.40754 | 0.870653636 |
| gain | sunitinib | EPHA6 | -0.617882202 | 0.40754 | 0.870653636 |
| gain | sunitinib | EPHA7 | 0.112037265 | 0.88071 | 0.982244706 |
| gain | sunitinib | EPHB1 | 0.112037265 | 0.88071 | 0.982244706 |
| gain | sunitinib | EPHB4 | 0.112037265 | 0.88071 | 0.982244706 |
| gain | sunitinib | FER | 0.029369326 | 0.97702 | 0.982244706 |
| gain | sunitinib | FGFR1 | -0.71074506 | 0.35285 | 0.870653636 |
| gain | sunitinib | FGFR3 | -0.617882202 | 0.40754 | 0.870653636 |
| gain | sunitinib | FGR | -0.733153668 | 0.35527 | 0.870653636 |
| gain | sunitinib | FYN | 0.018380033 | 0.9922 | 0.9922 |
| gain | sunitinib | GAK | -0.617882202 | 0.40754 | 0.870653636 |
| gain | sunitinib | HCK | -0.209727531 | 0.74889 | 0.982244706 |
| gain | sunitinib | INSR | 0.467573805 | 0.4841 | 0.921373782 |
| gain | sunitinib | JAK1 | -0.733153668 | 0.35527 | 0.870653636 |
| gain | sunitinib | JAK2 | 0.454504274 | 0.52492 | 0.921373782 |
| gain | sunitinib | LCK | -0.733153668 | 0.35527 | 0.870653636 |
| gain | sunitinib | LYN | -0.843389847 | 0.18149 | 0.710835833 |
| gain | sunitinib | MAP3K4 | -0.052507343 | 0.94093 | 0.982244706 |
| gain | sunitinib | MAP4K5 | -0.839254399 | 0.20351 | 0.721884528 |
| gain | sunitinib | MARK2 | -0.733153668 | 0.35527 | 0.870653636 |
| gain | sunitinib | MYLK2 | -0.209727531 | 0.74889 | 0.982244706 |
| gain | sunitinib | NEK2 | -1.323224899 | 0.08092 | 0.575777647 |
| gain | sunitinib | NTRK1 | -1.001984362 | 0.10413 | 0.575777647 |
| gain | sunitinib | PHKG2 | -0.617882202 | 0.40754 | 0.870653636 |
| gain | sunitinib | PRKAA1 | 0.112037265 | 0.88071 | 0.982244706 |
| gain | sunitinib | PRKACA | 0.467573805 | 0.4841 | 0.921373782 |
| gain | sunitinib | PTK2 | -0.843389847 | 0.18149 | 0.710835833 |

| | | | | | |
|---|---|---|---|---|---|
| gain | sunitinib | RPS6KA2 | -0.052507343 | 0.94093 | 0.982244706 |
| gain | sunitinib | RPS6KA5 | -0.839254399 | 0.20351 | 0.721884528 |
| gain | sunitinib | SRC | -0.209727531 | 0.74889 | 0.982244706 |
| gain | sunitinib | STK10 | 0.029369326 | 0.97702 | 0.982244706 |
| gain | sunitinib | STK17A | 0.112037265 | 0.88071 | 0.982244706 |
| gain | sunitinib | STK17B | -0.617882202 | 0.40754 | 0.870653636 |
| gain | sunitinib | STK38L | -0.617882202 | 0.40754 | 0.870653636 |
| gain | sunitinib | STK4 | 0 | 0.74889 | 0.982244706 |
| gain | sunitinib | TTK | 0 | 0.564 | 0.921373782 |
| gain | sunitinib | YES1 | -0.288976298 | 0.61177 | 0.958439667 |
| gain | erdafitinib | FGFR1 | -2 | 0.08058 | 0.575777647 |
| gain | erdafitinib | FGFR2 | -2 | 0.07448 | 0.575777647 |
| gain | erdafitinib | FGFR3 | -2.406063562 | 0.00257 | 0.4136 |
| gain | erdafitinib | FGFR4 | -1.585576156 | 0.13014 | 0.631438974 |
| gain | erdafitinib | RET | -2.019125319 | 0.02786 | 0.575777647 |
| gain | erdafitinib | CSF1R | -1.585576156 | 0.13014 | 0.631438974 |
| gain | erdafitinib | PDGFRA | -1.552716054 | 0.16628 | 0.694680889 |
| gain | erdafitinib | PDGFRB | -1.585576156 | 0.13014 | 0.631438974 |
| gain | erdafitinib | KIT | -1.552716054 | 0.16628 | 0.694680889 |
| gain | erdafitinib | ABCB1 | -1.879329675 | 0.03586 | 0.575777647 |
| gain | olaparib | PARP1 | -0.039800738 | 0.92371 | 0.982244706 |
| gain | olaparib | PARP2 | -0.152558496 | 0.68556 | 0.982244706 |
| gain | daunorubicin | TOP2A | 1 | 0.19188 | 0.7214688 |
| gain | daunorubicin | TOP2B | 1 | 0.19188 | 0.7214688 |
| gain | doxorubicin | TOP2A | -1 | 0.49705 | 0.921373782 |
| gain | epirubicin | NOLC1 | -4 | 0.15997 | 0.694680889 |
| gain | epirubicin | TOP2A | -4 | 0.05628 | 0.575777647 |
| gain | gemcitabine | TYMS | 0 | 0.48904 | 0.921373782 |
| gain | gemcitabine | CMPK1 | -0.643562546 | 0.17361 | 0.709536522 |
| gain | gemcitabine | ABCB1 | -0.477408053 | 0.33864 | 0.870653636 |
| gain | gemcitabine | ABCB10 | 0 | 0.53772 | 0.921373782 |
| gain | gemcitabine | DCTD | -0.632459211 | 0.39812 | 0.870653636 |
| gain | gemcitabine | NME1 | -0.159428366 | 0.75591 | 0.982244706 |
| gain | methotrexate | DHFR | 0.274765647 | 0.58275 | 0.921373782 |
| gain | methotrexate | TYMS | 0 | 0.62248 | 0.959231475 |
| gain | methotrexate | MTHFD1 | 0 | 0.72218 | 0.982244706 |
| gain | methotrexate | MTHFR | -0.199732642 | 0.72242 | 0.982244706 |
| gain | vinblastine | TUBA1A | -1.111907617 | 0.44392 | 0.921373782 |
| gain | vinblastine | TUBB | -1.306207669 | 0.30043 | 0.870653636 |
| gain | vinblastine | TUBD1 | -1 | 0.26227 | 0.870653636 |
| gain | vinblastine | TUBG1 | -1 | 0.44392 | 0.921373782 |
| gain | vinblastine | TUBE1 | -1.432238985 | 0.29752 | 0.870653636 |
| gain | vinblastine | JUN | -1.185916701 | 0.44835 | 0.921373782 |
| gain | everolimus | MTOR | 1 | 0.16392 | 0.694680889 |
| gain | temsirolimus | MTOR | 0.015199616 | 0.96517 | 0.982244706 |
| gain | docetaxel | TUBB1 | -1.773809906 | 0.0044 | 0.4136 |
| gain | docetaxel | MAP4 | -1 | 0.51679 | 0.921373782 |
| gain | docetaxel | MAP2 | -0.819171652 | 0.38646 | 0.870653636 |
| gain | docetaxel | MAPT | -0.5812059 | 0.51679 | 0.921373782 |
| gain | docetaxel | NR1I2 | -1.181222524 | 0.1172 | 0.629531429 |
| gain | paclitaxel | TUBB1 | -1.500943402 | 0.13664 | 0.642208 |
| gain | paclitaxel | MAP4 | -0.350701842 | 0.77272 | 0.982244706 |
| gain | paclitaxel | MAP2 | 0.323021707 | 0.58321 | 0.921373782 |
| gain | paclitaxel | MAPT | -0.350701842 | 0.77272 | 0.982244706 |
| gain | paclitaxel | NR1I2 | -1.193482623 | 0.26749 | 0.870653636 |
| loss | bosutinib | ABL1 | -1 | 0.19718 | 0.809017143 |
| loss | bosutinib | SYK | -0.685638745 | 0.49304 | 0.809017143 |
| loss | bosutinib | PTK2B | 0.43659866 | 0.61289 | 0.809017143 |
| loss | bosutinib | STK25 | -1.006865694 | 0.26761 | 0.809017143 |
| loss | erlotinib | ABL1 | -0.7200705 | 0.32041 | 0.809017143 |
| loss | lapatinib | ERBB4 | 0 | 0.64826 | 0.809017143 |
| loss | ponatinib | FGFR1 | -1.05476241 | 0.20863 | 0.809017143 |
| loss | ponatinib | TEK | 0.043620802 | 0.948 | 0.948 |
| loss | sunitinib | ABL1 | 0.560474876 | 0.50621 | 0.809017143 |
| loss | sunitinib | FGFR1 | 0.339672479 | 0.70789 | 0.809017143 |
| loss | sunitinib | JAK2 | 0.560474876 | 0.50621 | 0.809017143 |
| loss | sunitinib | STK16 | 0.620916605 | 0.43389 | 0.809017143 |

| | | | | | |
|---|---|---|---|---|---|
| loss | gemcitabine | DCTD | 0.28660178 | 0.6657 | 0.809017143 |
| loss | methotrexate | ATIC | 0.105621515 | 0.87269 | 0.930869333 |
| loss | docetaxel | MAP2 | -0.711630091 | 0.50059 | 0.809017143 |
| loss | paclitaxel | MAP2 | 2.050496116 | 0.13549 | 0.809017143 |
| snv | ponatinib | FGFR3 | 0.012319085 | 0.98832 | 0.98832 |
| snv | sunitinib | EPHB1 | 0.619308914 | 0.43899 | 0.921675 |
| snv | sunitinib | FGFR3 | -0.506136362 | 0.5917 | 0.921675 |
| snv | erdafitinib | FGFR3 | 0.501686633 | 0.61445 | 0.921675 |
| snv | docetaxel | MAP4 | 0.027653109 | 0.97049 | 0.98832 |
| snv | paclitaxel | MAP4 | 2.276506124 | 0.00791 | 0.04746 |
