## Supplemental tables for "Bladder cancer organoids as a functional system to model different disease stages and therapy response": Table S11.pdf

| pval | beta | drug | fdr | morph |
|---|---|---|---|---|
| 0.53059 | -0.1683335 | untreated | 1 | morphology |
| 1 | -4.19E-16 | dmso | 1 | morphology |
| 1 | -3.59E-16 | h2o | 1 | morphology |
| 0.96839 | -0.0253018 | bosutinib | 1 | morphology |
| 0.20902 | -1 | cisplatin_6_um | 1 | morphology |
| 0.07681 | -3 | epirubicin | 1 | morphology |
| 0.1992 | 0.61903921 | everolimus | 1 | morphology |
| 0.85564 | -0.1569632 | erlotinib | 1 | morphology |
| 0.003 | -2 | docetaxel | 0.078 | morphology |
| 0.49145 | -0.8233104 | lapatinib | 1 | morphology |
| 0.41768 | -0.3855288 | crizotinib | 1 | morphology |
| 0.22243 | -0.8942274 | gemcitabine | 1 | morphology |
| 0.27132 | -1 | cisplatin+gemcitabine | 1 | morphology |
| 0.03661 | 1 | cisplatin_14_um | 0.76881 | morphology |
| 0.93482 | -0.0771532 | daunorubicin | 1 | morphology |
| 0.10294 | -2 | doxorubicin | 1 | morphology |
| 0.02544 | 0.9103173 | methotrexate | 0.58512 | morphology |
| 0.79454 | 0.12332412 | olaparib | 1 | morphology |
| 0.02558 | -3 | paclitaxel | 0.58512 | morphology |
| 0.01589 | -1 | ponatinib | 0.38136 | morphology |
| 0.88867 | 0.06056633 | rapamycin | 1 | morphology |
| 0.4177 | -0.6513619 | sunitinib | 1 | morphology |
| 0.90027 | -0.0447678 | temsirolimus | 1 | morphology |
| 0.087 | -0.6227934 | vinblastine | 1 | morphology |
| 0.19925 | 2 | mmc | 1 | morphology |
| 0.00419 | -3 | erdafitinib | 0.10475 | morphology |
| 0.97779 | 0.00727439 | untreated | 1 | subtype |
| 1 | -2.50E-16 | dmso | 1 | subtype |
| 1 | 4.33E-18 | h2o | 1 | subtype |
| 0.05382 | 0.7000387 | bosutinib | 1 | subtype |
| 0.01217 | 2 | cisplatin_6_um | 0.31642 | subtype |
| 0.70387 | -0.6360095 | epirubicin | 1 | subtype |
| 0.12322 | -0.6589234 | everolimus | 1 | subtype |
| 0.40829 | -0.6496685 | erlotinib | 1 | subtype |
| 0.73636 | 0.21877822 | docetaxel | 1 | subtype |
| 0.61629 | -0.5573422 | lapatinib | 1 | subtype |
| 0.13986 | 0.67249533 | crizotinib | 1 | subtype |
| 0.25036 | 0.81914357 | gemcitabine | 1 | subtype |
| 0.04782 | 2 | cisplatin+gemcitabine | 1 | subtype |
| 0.46718 | 0.42043055 | cisplatin_14_um | 1 | subtype |
| 0.4716 | 0.56806521 | daunorubicin | 1 | subtype |
| 0.88731 | -0.1474654 | doxorubicin | 1 | subtype |
| 0.48618 | -0.3000672 | methotrexate | 1 | subtype |
| 0.07969 | -0.6079831 | olaparib | 1 | subtype |
| 0.31147 | -1 | paclitaxel | 1 | subtype |
| 0.21914 | -0.6925398 | ponatinib | 1 | subtype |
| 0.51621 | -0.2728541 | rapamycin | 1 | subtype |
| 0.51153 | -0.4849737 | sunitinib | 1 | subtype |
| 0.0382 | -0.6706122 | temsirolimus | 0.955 | subtype |
| 0.21095 | -0.4468077 | vinblastine | 1 | subtype |
| 0.48643 | 0.94855928 | mmc | 1 | subtype |
| 0.50082 | -0.8155441 | erdafitinib | 1 | subtype |
| 0.3269 | 0.31570417 | untreated | 1 | growth_pattern |
| 1 | -2.89E-16 | dmso | 1 | growth_pattern |
| 1 | -7.14E-16 | h2o | 1 | growth_pattern |
| 0.44597 | 0.35551246 | bosutinib | 1 | growth_pattern |
| 0.05009 | 1 | cisplatin_6_um | 1 | growth_pattern |
| 0.59515 | 0.96568558 | epirubicin | 1 | growth_pattern |
| 0.53271 | 0.28409849 | everolimus | 1 | growth_pattern |
| 0.70959 | -0.3616739 | erlotinib | 1 | growth_pattern |
| 0.10934 | 1 | docetaxel | 1 | growth_pattern |
| 0.47918 | -0.8513276 | lapatinib | 1 | growth_pattern |
| 0.95126 | 0.03510853 | crizotinib | 1 | growth_pattern |
| 0.08092 | 1 | gemcitabine | 1 | growth_pattern |
| 0.03223 | 3 | cisplatin+gemcitabine | 0.77352 | growth_pattern |
| 0.38101 | -0.616535 | cisplatin_14_um | 1 | growth_pattern |

| | | | | |
|---|---|---|---|---|
| 0.85984 | -0.1583882 | daunorubicin | 1 | growth_pattern |
| 0.23038 | 1 | doxorubicin | 1 | growth_pattern |
| 0.42309 | -0.3701842 | methotrexate | 1 | growth_pattern |
| 0.9357 | 0.02398305 | olaparib | 1 | growth_pattern |
| 0.35665 | 1 | paclitaxel | 1 | growth_pattern |
| 0.12004 | 0.85780942 | ponatinib | 1 | growth_pattern |
| 0.01678 | 0.9575656 | rapamycin | 0.4195 | growth_pattern |
| 0.28119 | -0.8834643 | sunitinib | 1 | growth_pattern |
| 0.69602 | 0.14429355 | temsirolimus | 1 | growth_pattern |
| 0.90212 | 0.03004539 | vinblastine | 1 | growth_pattern |
| 0.01026 | -3 | mmc | 0.26676 | growth_pattern |
| 0.20193 | 1 | erdafitinib | 1 | growth_pattern |
