## Supplementary Information Material and Methods for "Bladder cancer organoids as a functional system to model different disease stages and therapy response"

### KEY RESOURCES TABLES

| REAGENT or RESOURCE | SOURCE | IDENTIFIER |
|---|---|---|
| Antibodies for immunofluorescence | | |
| Mouse monoclonal Anti-CD44 | BD Pharmingen | Cat#550988 |
| Rabbit polyclonal Anti-CK5/6 | BioLegend | Cat#905501 |
| Mouse monoclonal Anti-CK8 | Thermo Fisher Scientific | Cat#MA1-06318 |
| Mouse monoclonal Anti-CK14 | Abcam | Cat#ab9220 |
| Mouse monoclonal Anti-CK20 | Abcam | Cat#Ab854 |
| Rabbit monoclonal Anti-GATA3 | Cell Signalling | Cat#5852 |
| Rabbit monoclonal Anti-Ki67 | Gene Tex | Cat#GTX16667 |
| Rabbit monoclonal Anti-p63 | Abcam | Cat#ab124762-100 |
| Rabbit monoclonal Anti-UPKII | Abcam | Cat#ab213655 |
| Donkey anti-mouse IgG, Alexa Fluor 488 | Thermo Fisher Scientific | Cat#A21202 |
| Donkey anti-rabbit IgG, Alexa Fluor 555 | Thermo Fisher Scientific | Cat#A21434 |
| DAPI | Thermo Fisher Scientific | Cat#62248 |
| Antibodies for immunohistochemistry | | |
| Mouse monoclonal Anti-CD44 | BD Pharmingen | Cat#550988 |
| Mouse monoclonal Anti-CK5/6 | Merck & Cie | Cat#MAB1620 |
| Mouse monoclonal Anti-CK8 | BD Bioscience | Cat# 345779 |
| Mouse monoclonal Anti-CK14 | Biosystems | Cat#NCL-L-LL002 |
| Mouse monoclonal Anti-CK20 | Biosystems | Cat#320M-16 |
| Mouse monoclonal Anti-GATA3 | Biosystems | Cat# 390M-14 |
| Rabbit monoclonal Anti-Ki67 | Biosystems | Cat# RM-9106-S1 |
| Mouse monoclonal Anti-p63 | Biosystems | Cat# NCL-L-p63 |
| Rabbit monoclonal Anti-UPKII | Abcam | Cat#ab213655 |
| Biological samples | | |
| Human bladder tissues and blood | University of Bern, Inselspital | Table 1 |
| Chemicals, Peptides and Recombinant Proteins | | |
| Dulbecco's MEM media | Thermo Fisher Scientific | Cat#61965-026 |
| Primocin | InVivoGen | Cat# ant-pm-1 |
| Fetal Bovine Serum (collection media) | Merck | Cat# F7524 |
| Fetal Bovine Serum (organoid media) | Thermo Fisher Scientific | Cat#1027-106 |
| DMSO | Merck | Cat#D2650 |
| Advanced DMEM F12 Serum Free medium | Thermo Fisher Scientific | Cat#12634028 |
| GlutaMAX | Thermo Fisher Scientific | Cat#35050061 |
| HEPES | Thermo Fisher Scientific | Cat#15630056 |
| Collagenase II | Thermo Fisher Scientific | Cat#17101015 |
| DNase I | Roche | Cat#10104159001 |
| Y-27632-HCl Rock inhibitor | Selleckchem | Cat# S1049 |
| TrypLE Express | Thermo Fisher Scientific | Cat#12605028 |
| 4% PFA | Merck | Cat#P614B |
| Triton-X | Merck | Cat#T8787 |
| Donkey serum | Jackson Immunoresearch | Cat#017-000-121 |
| Tween | Merck | Cat#P4780 |
| B27 supplement | Thermo Fisher Scientific | Cat#17504044 |
| Nicotinamide | Merck | Cat#N0636 |

| | | |
|---|---|---|
| R-Spondin | Peprotch | Cat#120-38 |
| N-acetyl-cysteine | Merck | Cat#A9165 |
| SB202190 | Selleckchem | Cat# S7067 |
| Noggin | Peprotech | Cat#25038 |
| Wnt3a | Peprotech | Cat#31520 |
| Hepatocyte growth factor | Peprotech | Cat#10039 |
| A83-01 | Tocris | Cat#2939 |
| Epidermal growth factor | Peprotech | Cat#AF-100-15 |
| Fibroblast growth factor 10 | Peprotech | Cat#100-26 |
| Bosutinib | Selleckchem | Cat#S1014 |
| Cisplatin | Selleckchem | Cat#S1166 |
| Crizotinib | Selleckchem | Cat#S1068 |
| Daunorubicin | Selleckchem | Cat#S3035 |
| Docetaxel | Selleckchem | Cat#S1148 |
| Doxorubicin | Selleckchem | Cat#S1208 |
| Epirubicin | Selleckchem | Cat#S1223 |
| Erlotinib | Selleckchem | Cat#S7786 |
| Erdafitinib | Selleckchem | Cat#S8401 |
| Everolimus | Selleckchem | Cat#S1120 |
| Gemcitabine | Selleckchem or Sigma | Cat# S1714 G6423 |
| Lapatinib | Selleckchem | Cat#S2111 |
| Methotrexate | Selleckchem | Cat#S1210 |
| Mitomycin C | Selleckchem | Cat#S8146 |
| Olaparib | Selleckchem | Cat#S1060 |
| Paclitaxel | Selleckchem | Cat#S1150 |
| Ponatinib | Selleckchem | Cat#S1490 |
| Rapamycin | Selleckchem | Cat#S1039 |
| Sunitinib | Selleckchem | Cat#S7781 |
| Temsirolimus | Selleckchem | Cat#S1044 |
| Vinblastine | Selleckchem | Cat#S4505 |
| Critical Commercial Assays | | |
| DNeasy Blood and Tissue kit | Qiagen | Cat#69504 |
| ReliaPrem™ gDNA Tissue MIniprep System | Promega | Cat#A2051 |
| Qubit dsDNA high-sensitivity or broad-range kits | Thermo Fisher Scientific | Cat#Q33233 and Q33263 |
| CellTiter-Glo 3D assay | Promega | Cat#G9682 |
| Software and Algorithms | | |
| Prism Version 9 | GraphPad | http://www.graphpad.come/scientificsoftware/prism/ |
| R version 4.0.3 | (R Core Team, 2016) | https://www.r-project.org/ |
| Deposited Data | | |
| Raw data from the single cell RNA-seq | This paper | https://github.com/ETH-NEXUS/scAmpi_single_cell_RNA |

| Compound | Dose in patient | Cmax [uM] | $C_{PDOs}$ [uM] | Reference |
|---|---|---|---|---|
| Bosutinib | 500 mg | 0.377 | 0.37 | Liston and Davis (1) |
| Cisplatin | 80 mg/m$^2$ | 14.4 | 14 | Liston and Davis (1) |
| Cisplatin | 35 mg/m$^2$ | 5.37 | 6 | Protocol Ref: MPHAURCOIG |
| Crizotinib | 250 mg | 0.913 | 0.91 | Liston and Davis (1) |
| Daunorubicin | 50 mg/m$^2$ | 0.310 | 0.31 | Liston and Davis (1) |
| Docetaxel | 100 mg/m$^2$ | 5.47 | 5 | Liston and Davis (1) |
| Doxorubicin | 60 mg/m$^2$ | 6.73 | 6.5 | Liston and Davis (1) |
| Epirubicin | 120 mg/m$^2$ | 16.6 | 16.5 | Liston and Davis (1) |
| Erlotinib | 150 mg | 3.15 | 3 | Liston and Davis (1) |
| Erdafitinib | 8 mg | 3.3 | 0.003 | US Food and Drug Administration. Ref ID4418085 |
| Everolimus | 10 mg | 0.064 | 0.064 | Liston and Davis (1) |
| Gemcitabine | 1250 mg/m$^2$ | 89.3 | 89 | Liston and Davis (1) |
| Lapatinib | 1250 mg | 4.18 | 4 | Liston and Davis (1) |
| Methotrexate | 30 mg | 1.31 | 1 | Liston and Davis (1) |
| Mitomycin C | 15 mg/m$^2$ | 2.18 | 2 | Liston and Davis (1) |
| Olaparib | 400 mg | 13.1 | 13 | Liston and Davis (1) |
| Paclitaxel | 175 mg/m$^2$ | 4.27 | 4 | Liston and Davis (1) |
| Ponatinib | 45 mg | 0.137 | 0.13 | Liston and Davis (1) |
| Rapamycin | 2 mg | 0.016 | 0.016 | Liston and Davis (1) |
| Sunitinib | 50 mg | 0.181 | 0.18 | Liston and Davis (1) |
| Temsirolimus | 25 mg | 0.568 | 0.56 | Liston and Davis (1) |
| Vinblastine | 1 mg/m$^2$ | 0.035 | 0.035 | Liston and Davis (1) |
| Cisplatin + Gemcitabine | 35 mg/m$^2$<br>1250 mg/m$^2$ | 5.37<br>89 | 6<br>89 | Liston and Davis (1) |

### QUANTIFICATION AND STATISTICAL ANALYSIS

Unless specified, all statistical analysis was performed using GraphPad Prism v 9.2.0 and R. In Figure 1B, Fisher's test was used between the PDO forming efficacy measure in NMIBC and MIBC samples. In Figure 1G, two-way ANOVA with Sidak's multiple comparison was used to compare the cell viability measured at 96h to the one measured at time 0 in each sample. Whereas unpaired t test was used to test cell viability difference between NMIBC and MIBC PDO (an average cell viability for each sample was used as replicate). In Figure 2C, two-way ANOVA with Turkey's multiple comparison test was used to compare the % of PDO morphologies in samples clustered by tumor stages. In Figure 3A, Paired Wilcoxon test was used between tumor content measured in PDO and PT. In Figure 3B, Wilcoxon test was used between the copy-number similarity in matched samples and randomly paired ones. In Figure 3C, Chi-squared test was used to compare the proportion of shared and private SNVs between PDOs and PT. In Figure 3D, 3E and S9, Wilcoxon-test was used between the allelic fraction of shared and private SNVs in PDOs and in PT and between the SNV clonality in PT and PDOs. In Figure 5A, 6A and 6G, one-way ANOVA test was used for comparison of each drug treatment to its vehicle (p-value was adjusted within samples controlled for $H_2O$ or DMSO). Effective compounds were selected based on a z-score ≤ to -1.5 and statistically lower compared to the vehicle condition. To test the effect of different compounds with each other in one sample, the viability reductions (fold-changes with respect to the vehicle) from each compound were evaluated with a nonparametric Wilcoxon test, whereas to test their effect between samples an average z-score for each sample was used as replicate and compared with a parametric paired t test. For figure 5B-D a LLM was performed (see "**Drugs association analyses"** section). In Figure 6C, Paired Wilcoxon test was used between the SNV clonality measured in PDO derived from patient 1 in the baseline and relapse. In Figure S7, two-way ANOVA test (multiple comparisons) was used to compare markers expression between three groups (solid, hollow, or mixed morphologies) or four groups (Ta, T1, T2, and T3 tumor-stages). In figure S10. one-way ANOVA test (multiple comparisons) was used for comparison between three groups (solid, hollow, and mixed morphologies). In figure S11A a correlation test was performed between the effect of the different TKIs. Whereas in figure S11B and C a LMM was performed (see "**Drugs association analyses"** section). In Figure S12A, one-way ANOVA test was used for comparison of each drug treatment to its vehicle (p-value was adjusted within samples controlled for $H_2O$ or DMSO). In Figure S12B, a Wilcoxon unpaired test was performed between the clonality of the preserved and lost SNVs in patient 1.

### DATA AND SOFTWARE AVAILABILITY

Whole exome and scRNA sequencing data will be available under the accession number (to be uploaded).
